# Degron-Informed Design of DCAF10 Recruiter for Targeted Protein Degradation

**DOI:** 10.64898/2026.11.20.767674

**Authors:** Chen Zhou, Yuki Aketa, Pedro A S Vaz de Castro, Chenlu Zhang, Xiaokang Jin, Weiguo Cui, Xiaoyu Zhang

**Affiliations:** Department of Chemistry, Northwestern University, Evanston, Illinois 60208; Department of Pathology, Northwestern University, Feinberg School of Medicine, Chicago, Illinois 60611; Chemistry of Life Processes Institute, Northwestern University, Evanston, Illinois 60208; Robert H. Lurie Comprehensive Cancer Center, Northwestern University, Chicago, Illinois 60611; Center for Human Immunobiology, Northwestern University, Chicago, Illinois 60611; International Institute for Nanotechnology, Northwestern University, Evanston, Illinois 60208

## Abstract

Targeted protein degradation (TPD) is an emerging biological and therapeutic strategy that harnesses small molecules or biologics to eliminate proteins of interest by engaging the cellular protein degradation machinery. Here, we report DCAF10 as a TPD-competent E3 ligase. Guided by DCAF10’s endogenous recognition of N-terminal acetyl-glycine degrons, we developed an Ac-Gly-Phe-Gly (Ac-GFG)-based recruiter capable of inducing DCAF10-dependent degradation of FKBP12 and multiple protein kinases. Interestingly, the Ac-GFG-based recruiter also exhibited a minor dependence on a secondary E3 ligase, FBXO22, potentially due to metabolic conversion that generates a terminal amine-containing species. Structure-activity relationship studies further identified an optimized N-propionyl-Gly-Phe-Ile (Pr-GFI) motif that enables potent degradation of FKBP12 (DC_50_ = 1.56 nM, D_max_ = 96%) with exclusive dependence on DCAF10. We further leveraged this recruiting motif to develop a BRD9 degrader that induces potent, selective, and DCAF10-dependent degradation. Our findings establish the Pr-GFI motif as a potent and selective DCAF10 recruiter and provide a new chemical handle for expanding the scope of TPD.

## Introduction

Targeted protein degradation (TPD) has emerged as a promising therapeutic strategy that harnesses the cellular ubiquitin-proteasome system to eliminate disease-associated proteins^1–4^. Various TPD strategies have been developed, including heterobifunctional and monofunctional degraders based on small molecules, peptides, and biologics^4,5^. These approaches typically induce proximity between a protein of interest (POI) and an E3 ligase, leading to POI ubiquitination and subsequent proteasomal degradation. The therapeutic potential of this approach has recently been demonstrated by the U.S. Food and Drug Administration (FDA) approval of vepdegestrant (ARV-471) for the treatment of advanced or metastatic breast cancer^6,7^. Most reported degraders rely on a limited number of E3 ligases, predominantly cereblon (CRBN) and von Hippel-Lindau (VHL)^8^. Notably, substantial progress has been made in expanding the repertoire of E3 ligases for TPD, although challenges remain, including acquired resistance^9,10^ and off-target effects associated with certain degraders^11^. These advances and challenges highlight the importance of identifying additional TPD-competent E3 ligases to further expand the therapeutic scope of TPD.

E3 ligases achieve substrate selectivity by recognizing distinct molecular features, such as short peptide motifs known as degrons^12^. These endogenous recognition elements provide a starting point for the rational development of E3 ligase recruiters for TPD^13–15^. Among various types of degrons, N- and C-terminal degrons provide opportunities for E3 ligase recruitment due to their short, defined recognition motifs^16–21^. However, the charged and polar functional groups frequently present in these degrons can limit cellular permeability. Recent studies identifying F-box protein 31 (FBXO31) as an E3 ligase that recognizes C-terminal amide-bearing proteins (CTAPs), followed by the development of a charge-neutral amidated Ala-Phe motif for FBXO31 recruitment in TPD, have demonstrated a strategy to overcome some of these limitations^16,22^.

A recent study demonstrated that proteins normally undergoing N-myristoylation can instead undergo N-acetylation when N-myristoylation fails^23^. These proteins, bearing an N-terminal acetyl-glycine (Ac-Gly) motif, are recognized by the E3 ligase DCAF10 and targeted for proteasomal degradation (**Figure 1A**)^23^. This N-terminal acetylation provides a potentially favorable molecular feature for degrader development, as it caps the positively charged N-terminal amine. Here, guided by this degron pathway, we developed a cell-permeable DCAF10 recruiter bearing an N-acetyl-Gly-Phe-Gly (Ac-GFG) motif that induces target degradation with high nanomolar potency. We further optimized this recruiter to an N-propionyl-Gly-Phe-Ile (Pr-GFI) motif and achieved single-digit nanomolar degradation potency against FKBP12 and BRD9.

**Figure 1.**
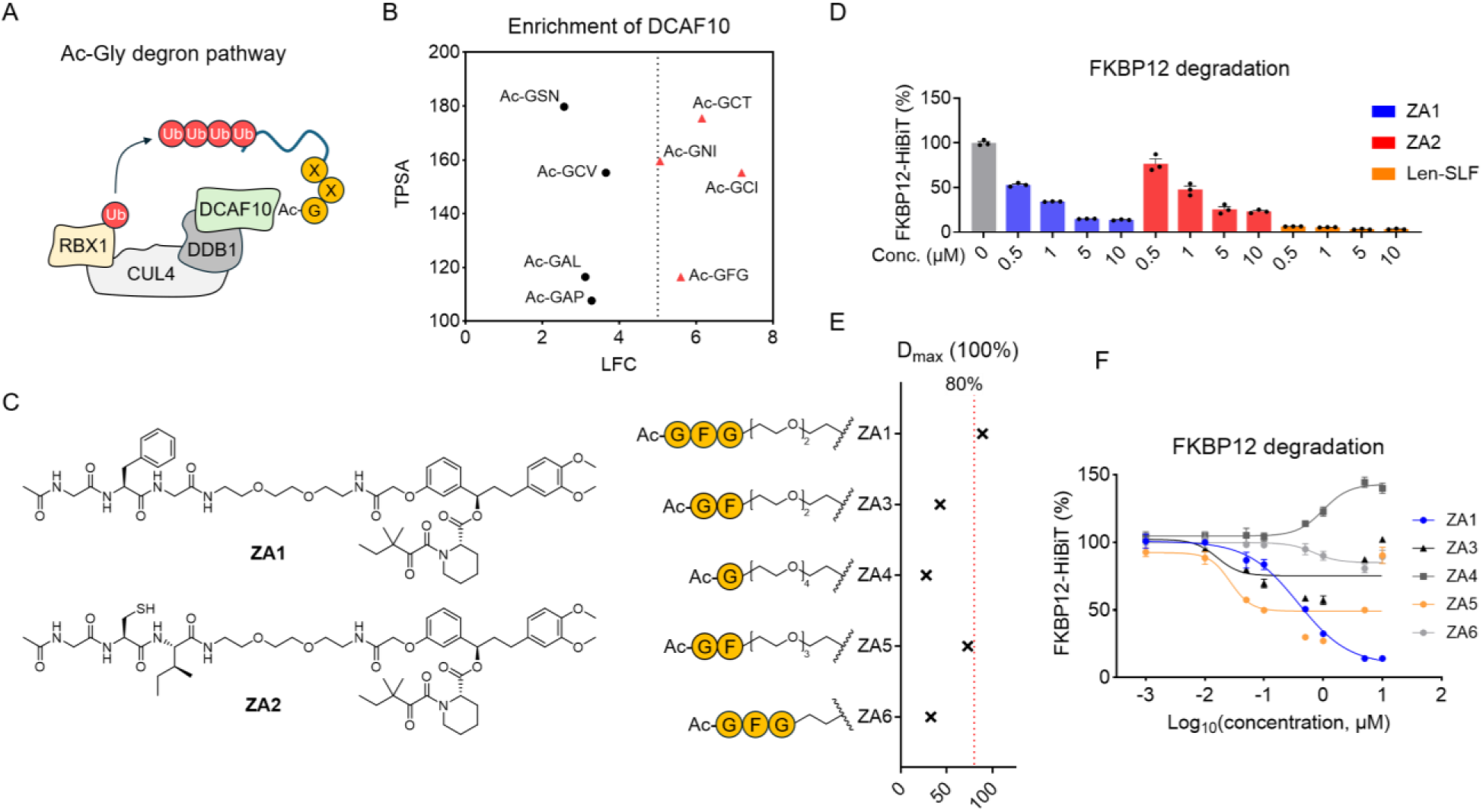
Harnessing the N-terminal acetyl-glycine motif to design bifunctional degraders. **A**. Schematic illustration of DCAF10-mediated recognition and ubiquitination of proteins bearing N-terminal acetyl-glycine (Ac-Gly) degrons. **B**. Analysis of published AP-MS data showing DCAF10 enrichment (log2 fold change, LFC) and calculated topological polar surface area (TPSA) of N-acetylated tripeptides. **C**. Structures of the Ac-GFG- and Ac-GCI-based FKBP12 degraders ZA1 and ZA2. **D**. FKBP12-HiBiT levels in HEK293T cells following treatment with ZA1, ZA2, or Len-SLF for 20 h. Data are presented as mean ± SEM (n = 3). **E**. D_max_ of FKBP12 degradation induced by ZA1 and its analogs ZA3-ZA6. **F**. Concentration-dependent degradation of FKBP12-HiBiT in HEK293T cells treated with ZA1 and ZA3-ZA6 for 20 h. Data are presented as mean ± SEM (n = 3).

## Results

### Harnessing the N-terminal acetyl-glycine motif to design bifunctional degraders

To assess the feasibility of applying the Ac-Gly motif for TPD, we first examined published affinity-purification mass spectrometry (AP-MS) data characterizing DCAF10 enrichment by a panel of Ac-Gly-containing peptides^22^. To select candidate recruiters, we used the reported log2 fold-change (LFC) values as a measure of DCAF10 enrichment and calculated the topological polar surface area (TPSA) of the corresponding N-acetylated tripeptide motifs as a proxy for cell membrane permeability^24^. Based on this analysis, we chose N-acetyl-Gly-Phe-Gly (Ac-GFG) and N-acetyl-Gly-Cys-Ile (Ac-GCI) as two candidate recruiters (**Figure 1B**). We then synthesized two FKBP12-targeting bifunctional compounds, ZA1 and ZA2, by conjugating these tripeptides to SLF (a synthetic ligand for FKBP12) through a PEG2 linker (**Figure 1C**).

Both compounds induced concentration-dependent degradation of endogenously HiBiT-tagged FKBP12 in HEK293T cells (**Figure 1D**). The positive control, the CRBN-recruiting degrader Len-SLF, induced robust FKBP12 degradation in this HiBiT system (**Figures S1A** and **1D**). ZA1 exhibited greater degradation activity than ZA2 at 500 nM, reducing FKBP12-HiBiT levels by 47% versus 23% for ZA2 (**Figure 1D**). In addition, ZA1 showed no obvious cytotoxicity at concentrations up to 10 µM, suggesting that the observed reduction in FKBP12 levels was unlikely to be a secondary effect of nonspecific cell killing (**Figure S1B**).

We next investigated the effects of peptide and linker length on degrader activity. For the initial evaluation, we used the maximum target degradation a degrader can achieve (D_max_). Truncation of the Ac-GFG tripeptide motif in ZA1 (D_max_ = 89%) to the Ac-GF dipeptide (ZA3) or the Ac-G monopeptide with a PEG4 linker to maintain the overall recruiter length (ZA4) reduced D_max_ to 43% and 28%, respectively. Extending the linker of the Ac-GF dipeptide to PEG3 (ZA5) also resulted in a reduced D_max_ of 73%.

Moreover, shortening the PEG2 linker in ZA1 to a two-carbon linker (ZA6) reduced D_max_ to 33% (**Figures 1E** and **1F**). Thus, ZA1 remained the most effective FKBP12 degrader in this series. These results also suggest the importance of retaining the tripeptide motif and an appropriate linker length for maintaining degrader activity based on the ZA1 scaffold.

### Mechanistic characterization of DCAF10-dependent target degradation by ZA1

To further characterize ZA1, we performed a more detailed dose-dependent degradation analysis, which revealed the D_max_ of 88% and the DC_50_ of 450 nM in parental HEK293T cells (**Figure 2A** and **2B**). To investigate DCAF10 dependence, we generated *DCAF10* knockout (KO) and overexpressing (OE) FKBP12-HiBiT HEK293T cells. DCAF10 overexpression enhanced both potency and depth of ZA1-mediated FKBP12 degradation, with the DC_50_ of 1.0 nM and the D_max_ of 97% (**Figure 2A** and **2B**). Conversely, *DCAF10* KO substantially impaired ZA1 activity, increasing the DC_50_ to 2.4 µM and reducing the D_max_ to 62% (**Figure 2A** and **2B**). The Ac-GFG motif contains one chiral center at the L-Phe residue. We further synthesized a D-Phe-containing analog, ZA7 (**Figure 2C**), which showed no FKBP12 degradation at all concentrations tested (**Figure 2A**). This indicates that the naturally occurring L-phenylalanine stereochemistry is essential for degrader activity.

**Figure 2.**
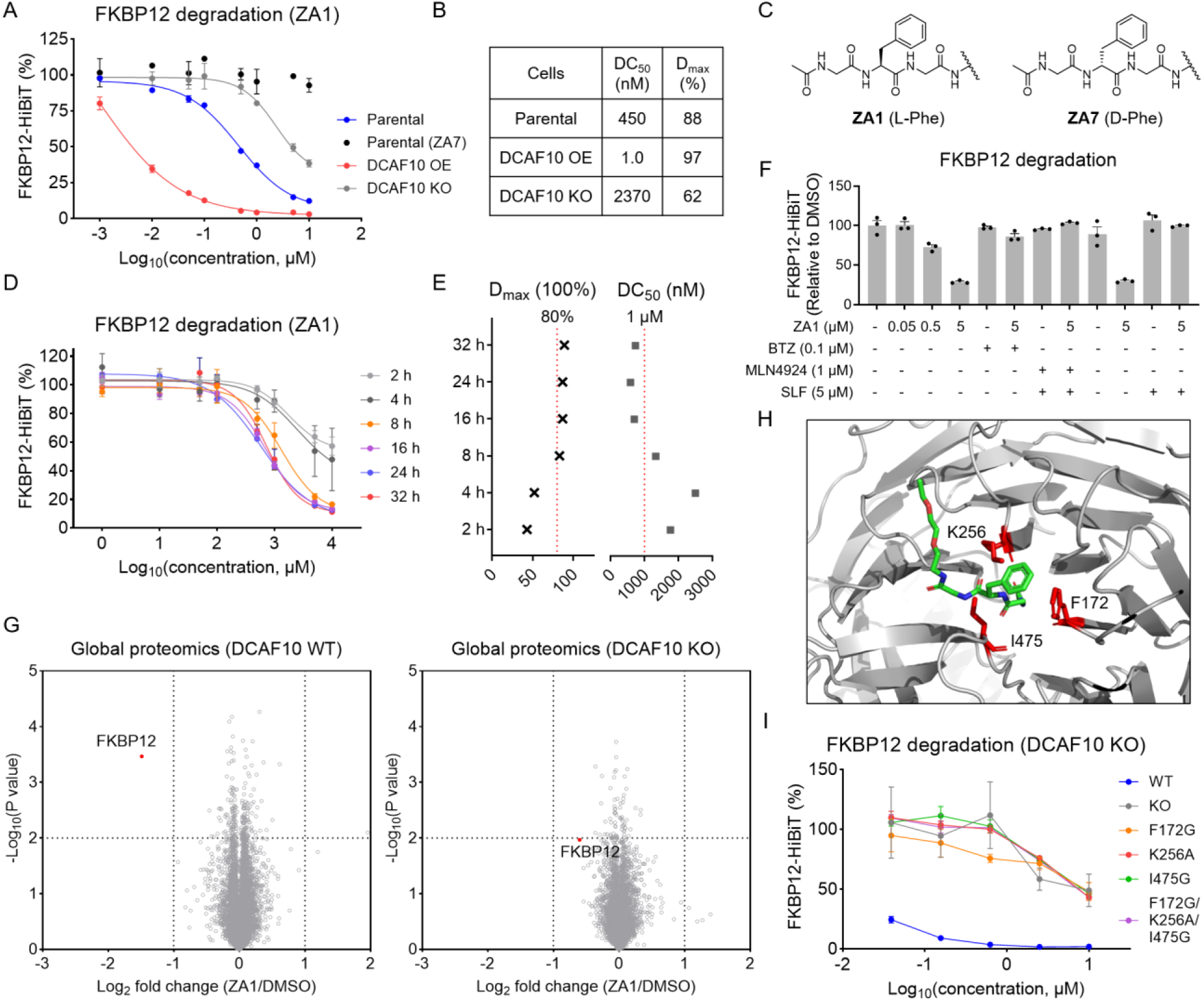
Mechanistic characterization of DCAF10-dependent FKBP12 degradation by ZA1. **A**. Concentration-dependent degradation of FKBP12-HiBiT induced by ZA1 in parental, *DCAF10* KO, and DCAF10 OE HEK293T cells, and by the D-Phe-containing analog ZA7 in parental cells. Cells were treated for 20 h. Data are presented as mean ± SEM (n = 3). **B**. D_max_ and DC_50_ values for ZA1-induced FKBP12 degradation in parental, *DCAF10* KO, and DCAF10 OE cells. **C**. Structures of ZA1 and ZA7. **D**. Time-dependent degradation of FKBP12-HiBiT induced by ZA1 in parental HEK293T cells. **E**. D_max_ and DC_50_ values for ZA1-induced FKBP12 degradation at the indicated time points. **F**. Effects of the proteasome inhibitor bortezomib (BTZ), neddylation inhibitor MLN4924, and SLF on ZA1-induced FKBP12 degradation. Cells were pretreated with the indicated compounds for 1 h, followed by treatment with ZA1 for 8 h. Data are presented as mean ± SEM (n = 3). **G**. Global proteomic analysis of ZA1-induced protein degradation in parental and *DCAF10* KO HEK293T cells (5 µM, 24 h). *P* values were calculated using a two-sided t-test and adjusted for multiple comparisons by the Benjamini-Hochberg method (n = 3 for DMSO; n = 2 for ZA1). **H**. Boltz-2-predicted binding model of the ZA1 recruiter moiety within the DCAF10 degron-binding pocket. Key residues involved in binding are highlighted. **I**. Concentration-dependent FKBP12-HiBiT degradation induced by ZA1 in *DCAF10* KO cells and *DCAF10* KO cells re-expressing DCAF10 WT or mutants. Cells were treated for 20 h. Data are presented as mean ± SEM (n = 3).

We next characterized the time dependence of ZA1-induced FKBP12 degradation. ZA1 induced detectable FKBP12 degradation within 2 h and achieved maximal degradation at 24 h (**Figure 2D, 2E** and **S2A**). Furthermore, ZA1-induced FKBP12 degradation was blocked by the proteasome inhibitor bortezomib (BTZ)^25^, the neddylation inhibitor MLN4924^26^, and excess SLF (**Figure 2F**), supporting a proteasome- and cullin-RING ligase (CRL)-dependent degradation mechanism that requires engagement of FKBP12 by the SLF moiety. As an orthogonal approach to validate ZA1 activity, we expressed cytosolic or nuclear-localized FLAG-tagged FKBP12 in HEK293T cells. ZA1 induced degradation of both FKBP12 variants, and DCAF10 overexpression enhanced the degradation (**Figure S2B**). We further evaluated the proteome-wide selectivity of ZA1-induced degradation by performing global proteomic analysis of parental and *DCAF10* KO HEK293T cells expressing FKBP12-HiBiT following treatment with ZA1 or DMSO. FKBP12 was selectively degraded upon ZA1 treatment in parental cells, whereas its degradation was substantially attenuated in *DCAF10* KO cells (**Figure 2G** and **Table S1**).

To understand structural features of DCAF10 involved in ZA1 recognition, we performed co-folding of DCAF10 and the recruiter moiety of ZA1 using Boltz-2^27^. The resulting model suggested that the recruiter moiety of ZA1 occupies the previously characterized degron-binding pocket of DCAF10^23^, with key interactions involving residues K256, F172, and I475 (**Figure 2H** and **S2C**). We then expressed DCAF10 variants harboring individual mutations (K256A, F172G, or I475G) or a combined triple mutation (K256A/F172G/I475G) in *DCAF10* KO HEK293T cells. None of these mutants effectively restored ZA1-induced FKBP12 degradation, suggesting the importance of these residues for degrader activity, although residual degradation was observed at higher concentrations (**Figure 2I**).

### DCAF10 and FBXO22 differentially contribute to ZA1-induced FKBP12 degradation

Our data showed residual ZA1-induced degradation activity at higher concentrations in *DCAF10* KO cells (**Figure 2A**), suggesting the potential involvement of additional E3 ligases. To investigate this possibility, we performed AP-MS using FLAG-FKBP12 as the bait. ZA1 treatment induced interactions of FKBP12 with DCAF10 and its CRL4 adaptor DDB1, consistent with the formation of a ZA1-induced ternary complex involving DCAF10 and FKBP12 (**Figure 3A** and **Table S2**). Notably, AP-MS also identified FBXO22 and its associated SKP1-CUL1-F-box (SCF) component SKP1 as additional proteins enriched upon ZA1 treatment (**Figure 3A**), although their substantially lower spectral counts compared with those of DCAF10 and DDB1 suggested lower relative enrichment abundance (**Figure 3B**).

**Figure 3.**
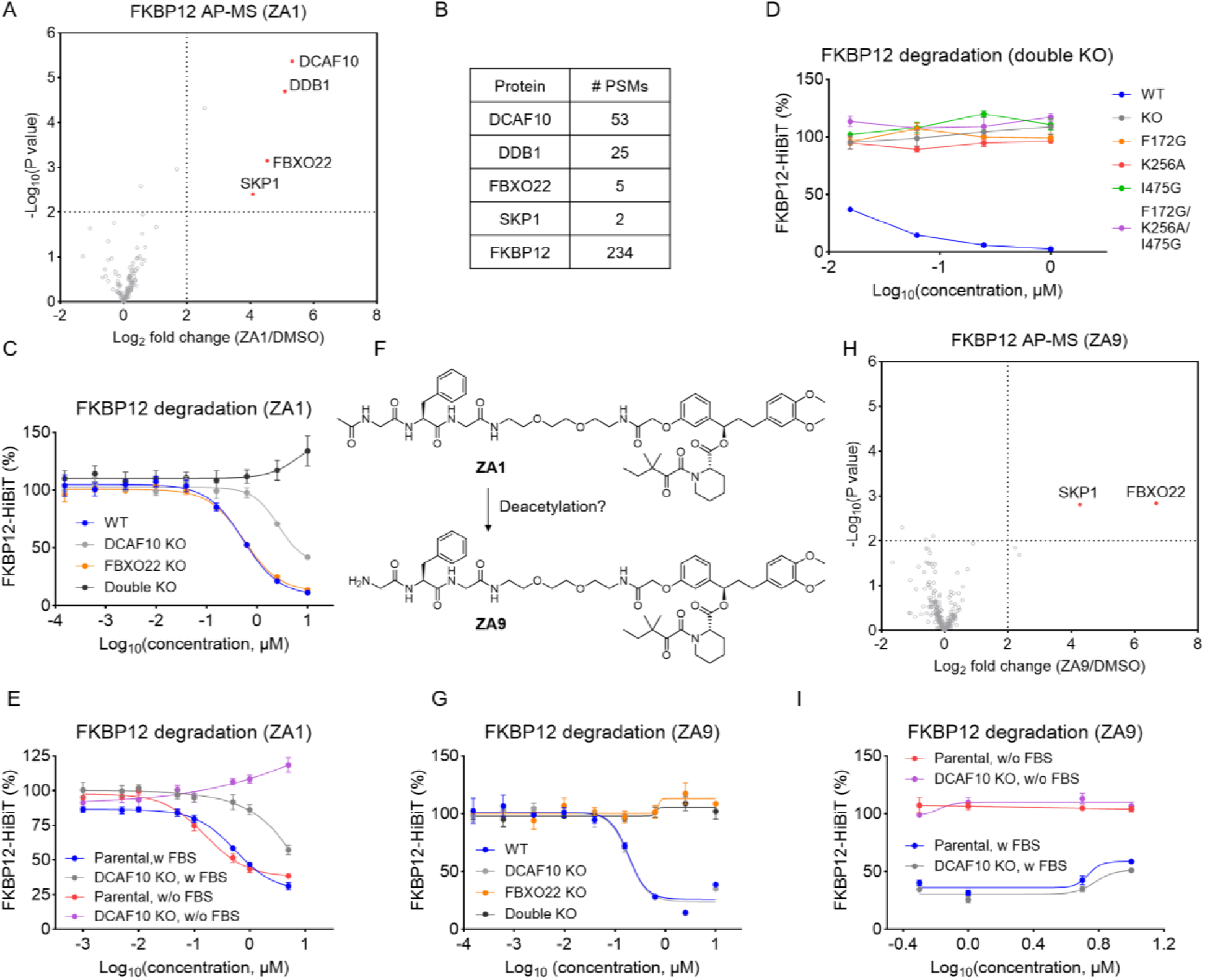
DCAF10 and FBXO22 differentially contribute to ZA1-induced FKBP12 degradation. **A**. AP-MS analysis of ZA1-induced protein interactions with FLAG-FKBP12. Cells were pretreated with 100 nM BTZ for 1 h, followed by treatment with DMSO or 5 µM ZA1 for 3 h. *P* values were calculated using a two-sided t-test and adjusted for multiple comparisons by the Benjamini-Hochberg method (n = 3). **B**. Spectral counts of DCAF10, DDB1, FBXO22, SKP1, and FKBP12 identified by AP-MS. **C**. Concentration-dependent FKBP12-HiBiT degradation induced by ZA1 in parental, *DCAF10* KO, *FBXO22* KO, and *DCAF10*/*FBXO22* double KO HEK293T cells. Cells were treated for 20 h. Data are presented as mean ± SEM (n = 3). **D**. Concentration-dependent FKBP12-HiBiT degradation induced by ZA1 in *DCAF10*/*FBXO22* double KO cells re-expressing DCAF10 WT or mutants. Cells were treated for 20 h. Data are presented as mean ± SEM (n = 3). **E**. Effects of FBS depletion on ZA1-induced FKBP12-HiBiT degradation in parental and *DCAF10* KO cells. Cells were treated for 20 h. Data are presented as mean ± SEM (n = 3). **F**. Proposed conversion of ZA1 to ZA9, the N-terminally deacetylated analog of ZA1. **G**. Concentration-dependent FKBP12-HiBiT degradation induced by ZA9 in parental, *DCAF10* KO, and *FBXO22* KO cells. Cells were treated for 20 h. Data are presented as mean ± SEM (n = 3). **H**. AP-MS analysis of ZA9-induced protein interactions with FLAG-FKBP12. Cells were pretreated with 100 nM BTZ for 1 h, followed by treatment with DMSO or 5 µM ZA1 for 3 h. *P* values were calculated using a two-sided t-test and adjusted for multiple comparisons by the Benjamini-Hochberg method (n = 2). **I**. Effects of FBS depletion on ZA9-induced FKBP12-HiBiT degradation in parental and *DCAF10* KO cells. Cells were treated for 20 h. Data are presented as mean ± SEM (n = 3).

To investigate the dependence on these two E3 ligases, we generated *FBXO22* single KO and *DCAF10*/*FBXO22* double KO FKBP12-HiBiT HEK293T cells. Unlike *DCAF10* single KO, which dramatically reduced ZA1-induced FKBP12 degradation, *FBXO22* single KO had little effect, whereas degradation was completely abolished in *DCAF10*/*FBXO22* double KO cells (**Figure 3C**). These results suggest that DCAF10 is the predominant E3 ligase responsible for ZA1-induced FKBP12 degradation, while FBXO22 contributes to the residual degradation activity observed in the absence of DCAF10. Moreover, re-expression of DCAF10 variants (K256A, F172G, I475G, or K256A/F172G/I475G) in the double KO cells did not restore ZA1-induced FKBP12 degradation (**Figure 3D**). Given that residual degradation activity was observed following re-expression of these variants in *DCAF10* single KO cells (**Figure 2I**), this residual activity is likely mediated by FBXO22.

This dual E3-dependent mechanism, in which DCAF10 plays a predominant role, prompted us to consider whether a potential contaminant in ZA1 could engage FBXO22 to support FKBP12 degradation. However, neither LC-MS nor NMR analysis revealed clear evidence of such contamination (**Figure S3A**). Although we could not completely rule out the possibility that trace amounts of a contaminant might induce robust degradation, we sought to investigate alternative mechanisms that could explain the unexpected involvement of FBXO22 in ZA1-induced FKBP12 degradation. Previous studies have shown that heterobifunctional compounds containing a terminal primary amine can be converted into aldehydes by metabolic enzymes present in culture serum, which can covalently recruit FBXO22 for TPD^28,29^. Guided by this mechanism, we first evaluated ZA1-induced FKBP12 degradation in parental and *DCAF10* KO HEK293T cells cultured in the presence or absence of fetal bovine serum (FBS). Either FBS removal or *FBXO22* knockout completely abolished FKBP12 degradation induced by ZA8 (**Figure S3B**), an analog of the previously reported amine-containing FKBP12-targeting degrader SP3N^28^ (**Figure S3C** and **S3D**). Using this system, we tested ZA1 and found that removing FBS completely abolished the residual FKBP12 degradation observed in *DCAF10* KO cells (**Figure 3E**). This suggests a potentially similar mechanism in which a precursor species (e.g., a compound containing a terminal primary amine) is converted into an active degrader by metabolic enzymes present in serum. Interestingly, ZA1-induced FKBP12 degradation in parental cells was largely unaffected by FBS removal (**Figure 3E**), suggesting that in the presence of endogenous DCAF10, ZA1-induced FKBP12 degradation is predominantly mediated by DCAF10.

Given that ZA1 lacks a terminal primary amine, we hypothesized that FBS-dependent metabolic processes might generate a derivative of ZA1 containing an N-terminal free primary amine. To explore this possibility, we synthesized ZA9, the N-terminally deacetylated analog of ZA1 (**Figure 3F**), which induced dose-dependent FKBP12 degradation (**Figure 3G**). AP-MS using FLAG-FKBP12 as the bait revealed selective enrichment of FBXO22 and SKP1, but not DCAF10 and DDB1, upon ZA9 treatment (**Figure 3H** and **Table S3**). Consistently, ZA9-induced FKBP12 degradation was completely abolished by *FBXO22* KO but was unaffected by *DCAF10* KO (**Figure 3G**). Moreover, ZA9-induced degradation was abolished in the absence of FBS in both parental and *DCAF10* KO cells (**Figure 3I**), suggesting that the primary amine may undergo metabolic conversion to an aldehyde that serves as the active species, although this proposed mechanism requires further validation.

Another potential pathway for generating a species containing a terminal primary amine is cleavage of the amide bond between the second residue (Phe) and the third residue (Gly). We therefore synthesized the corresponding compound, ZA10 (**Figure S4A**), and confirmed that, similar to ZA9, ZA10 induced dose-dependent FKBP12 degradation. This degradation was dependent only on FBXO22 and abolished by FBS removal (**Figure S4B** and **S4C**). Our results suggest a potential mechanism in which ZA1 may undergo metabolic conversion to generate species containing terminal primary amines that subsequently support FBXO22-dependent degradation. Further investigation of this mechanism may provide valuable insights into the metabolic regulation and E3 ligase selectivity of targeted protein degraders.

### Expanding DCAF10-mediated target degradation to protein kinases

To explore the applicability of DCAF10 recruitment for degrading additional proteins, we chose TL13-87, a multikinase binder that has previously been used to develop heterobifunctional kinase degraders^30–32^. We synthesized two kinase degraders incorporating the ZA1-derived DCAF10 recruiter with either a PEG2 (ZA11) or PEG3 (ZA12) linker (**Figure 4A**). Given the essential roles of many protein kinases in cell survival, comparing the cytotoxic effects of kinase degraders in DCAF10-expressing and *DCAF10* KO cells could provide an initial indication of potential target degradation. This strategy was validated in our previous study of FBXO31-recruiting kinase degraders, which showed significantly greater cytotoxicity in FBXO31-expressing cells than in *FBXO31* KO cells^16^. As expected, cytotoxicity induced by both ZA11 and ZA12 was enhanced in *DCAF10* KO HEK293T cells re-expressing DCAF10 compared with *DCAF10* KO cells (**Figure 4B** and **4C**). We also evaluated a CRBN-recruiting degrader SK-3-91, which shares the same kinase-binding scaffold^32^ (**Figure S5A**), and found that it exhibited comparable cytotoxicity in DCAF10-expressing and *DCAF10* KO cells (**Figure 4B** and **4C**).

**Figure 4.**
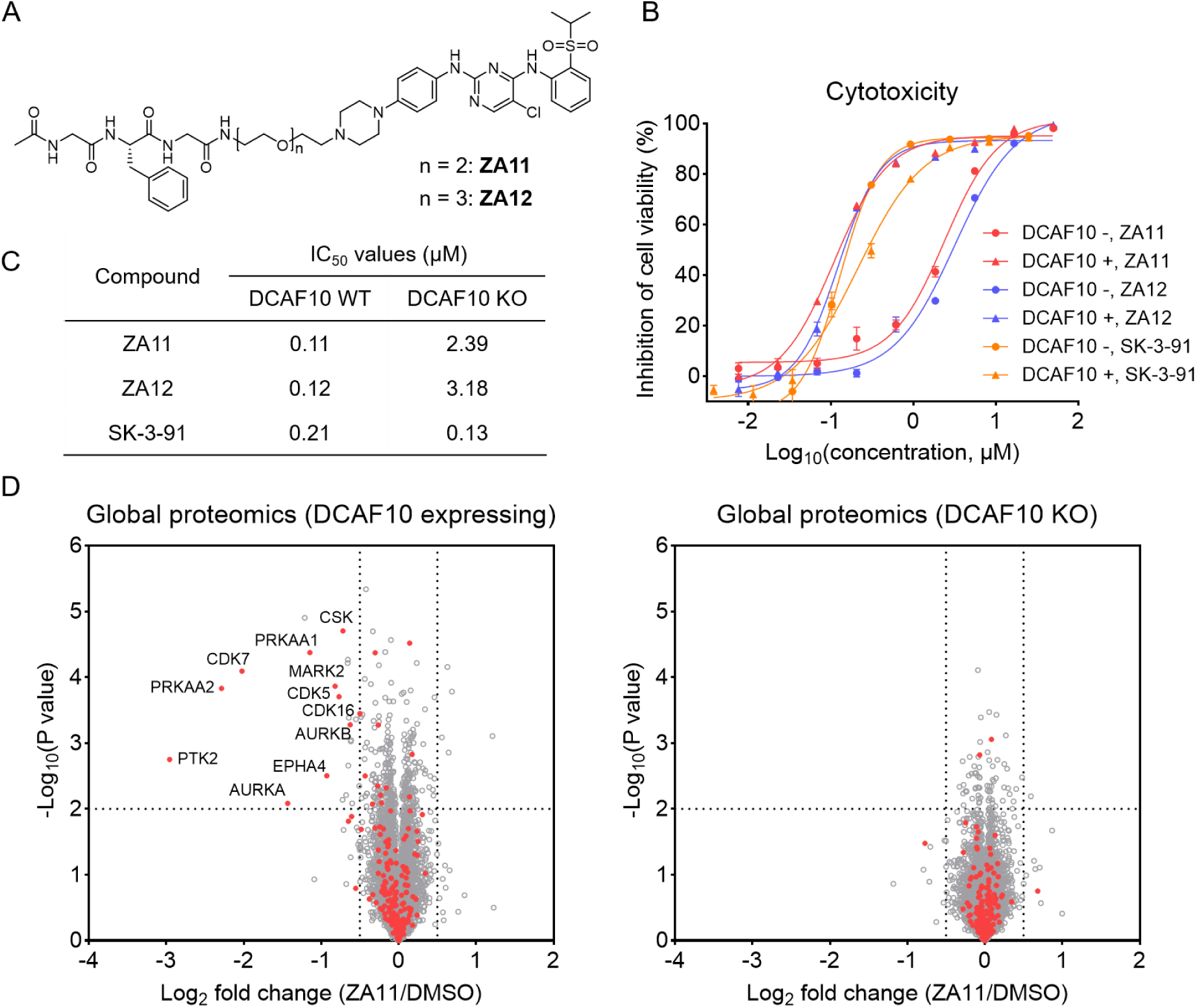
Expanding DCAF10-mediated target degradation to protein kinases. **A**. Structures of the DCAF10-recruiting kinase degraders ZA11 and ZA12. **B**. Concentration-dependent cytotoxicity induced by ZA11, ZA12 and SK-3-91 in *DCAF10* KO HEK293T cells and *DCAF10* KO cells re-expressing DCAF10. Cells were treated for 72 h. Data are presented as mean ± SEM (n = 3). **C**. Half-maximal inhibitory concentration (IC_50_) values of ZA11, ZA12 and SK-3-91 in *DCAF10* KO HEK293T cells and *DCAF10* KO cells re-expressing DCAF10. **D**. Global proteomic analysis of ZA11-induced protein degradation in *DCAF10* KO HEK293T cells and *DCAF10* KO cells re-expressing DCAF10 (100 nM, 24 h). *P* values were calculated using a two-sided t-test and adjusted for multiple comparisons by the Benjamini-Hochberg method (n = 3).

To further assess kinase degradation, we performed global proteomics following ZA11 treatment in *DCAF10* KO HEK293T cells and *DCAF10* KO cells re-expressing DCAF10 (**Figure S5B** and **Table S4**). ZA11 induced significant degradation of multiple kinases in DCAF10-expressing cells, whereas no significant kinase degradation was detected in *DCAF10* KO cells (**Figure 4D** and **Table S4**). In contrast, the CRBN-recruiting degrader SK-3-91 induced degradation of a subset of kinases in both DCAF10-expressing and *DCAF10* KO cells (**Figure S5C** and **Table S4**). Interestingly, SK-3-91 appeared to degrade these kinases more effectively in *DCAF10* KO cells. We found that CRBN expression was upregulated in *DCAF10* KO cells (**Figure S5D**), which may explain the enhanced degradation activity. Nevertheless, how DCAF10 expression regulates CRBN expression remains to be investigated.

### Optimization of the DCAF10 recruiters

We next sought to optimize the structure of ZA1 to improve degrader activity by modifying the N-terminal group and each of the three amino acid residues within the ZA1-derived recruiter (**Figure 5A**). All resulting analogs were conjugated to SLF through a PEG2 linker and evaluated for FKBP12 degradation using the HiBiT assay. We first investigated modifications to the N-terminal acetyl group. Replacing the acetyl group with an ethyl group (ZA13) slightly reduced degradation activity, whereas introducing an N-methyl group on the amide nitrogen (ZA14) or replacing the amide linkage with a ketone (ZA15) or ester (ZA16) markedly impaired degradation activity (**Figure 5B, 5C** and **S6**). These results suggest that the N-terminal amide moiety is important for productive degrader activity. We next examined the effect of N-terminal acyl chain length. Whereas N-octanoylation (ZA17) reduced degradation activity, N-propionylation (ZA18) improved degradation potency by approximately 3-fold (DC_50_ = 98 nM) while maintaining a similar D_max_ (86%) (**Figure 5B, 5C** and **S6**).

**Figure 5.**
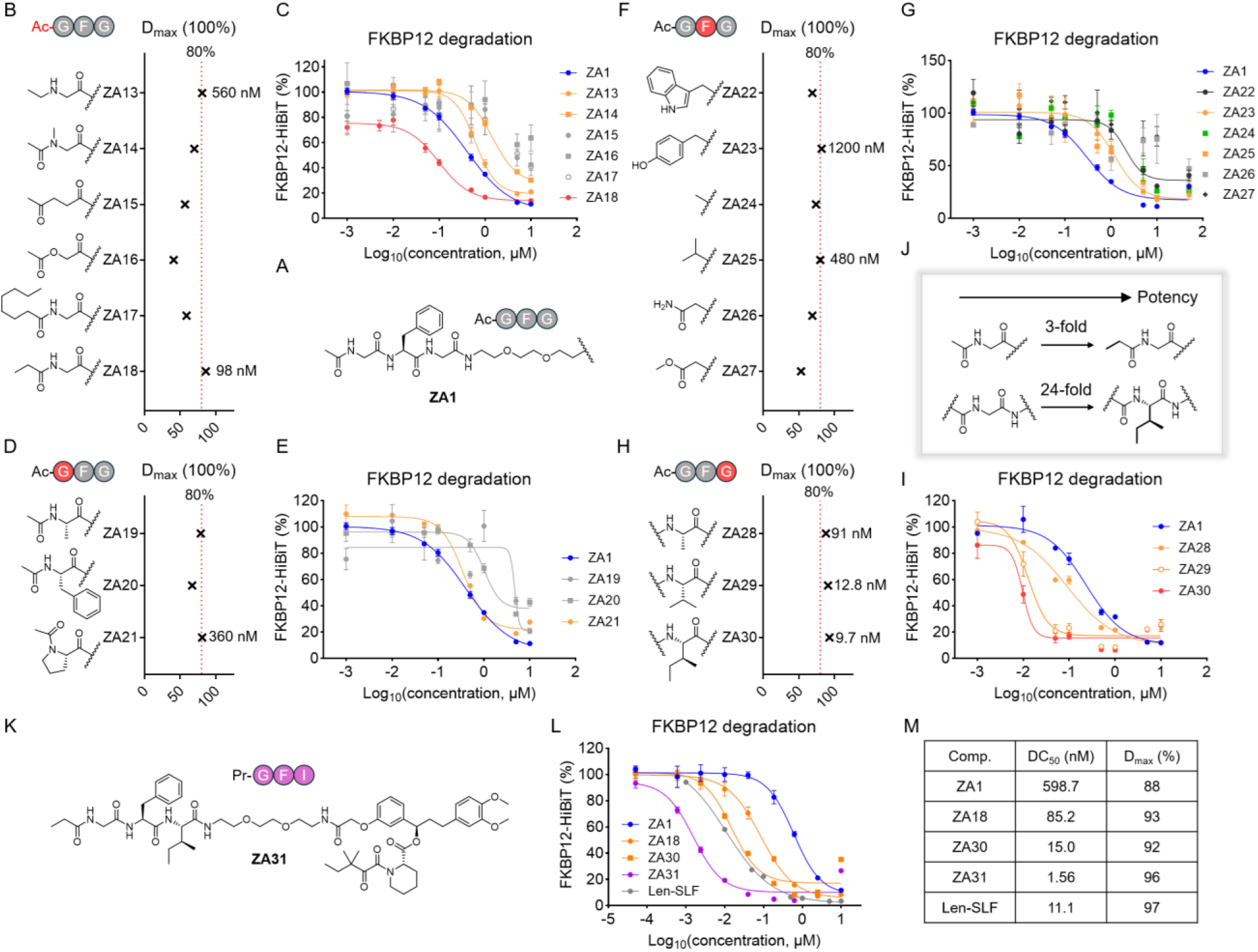
Optimization of the DCAF10 recruiters. **A**. Structure of the ZA1-derived DCAF10 recruiter showing the positions for optimization. **B**. D_max_ values of FKBP12 degradation induced by ZA13-ZA18 in HEK293T cells. **C**. Concentration-dependent FKBP12-HiBiT degradation induced by ZA1 and ZA13-ZA18 in HEK293T cells. Cells were treated for 20 h. Data are presented as mean ± SEM (n = 3). **D**. D_max_ values of FKBP12 degradation induced by ZA19-ZA21 in HEK293T cells. **E**. Concentration-dependent FKBP12-HiBiT degradation induced by ZA1 and ZA19-ZA21 in HEK293T cells. Cells were treated for 20 h. Data are presented as mean ± SEM (n = 3). **F**. D_max_ values of FKBP12 degradation induced by ZA22-ZA27 in HEK293T cells. **G**. Concentration-dependent FKBP12-HiBiT degradation induced by ZA1 and ZA22-ZA27 in HEK293T cells. Cells were treated for 20 h. Data are presented as mean ± SEM (n = 3). **H**. D_max_ values of FKBP12 degradation induced by ZA28-ZA30 in HEK293T cells. **I**. Concentration-dependent FKBP12-HiBiT degradation induced by ZA1 and ZA28-ZA29 in HEK293T cells. Cells were treated for 20 h. Data are presented as mean ± SEM (n = 3). **J**. Comparison of FKBP12 degradation potency following N-terminal propionylation or isoleucine substitution at the third amino acid position. **K**. Structure of ZA31. **L**. Concentration-dependent FKBP12-HiBiT degradation induced by ZA1, ZA18, ZA30, ZA31, and Len-SLF in HEK293T cells. Cells were treated for 20 h. Data are presented as mean ± SEM (n = 3). **M**. D_max_ and DC_50_ values of FKBP12 degradation induced by ZA18, ZA30, ZA31, and Len-SLF in HEK293T cells.

We next examined the contribution of the first amino acid residue. Substitution of glycine with alanine (ZA19) or phenylalanine (ZA20) reduced degradation potency, consistent with previous reports indicating the importance of an N-terminal glycine residue^23^ (**Figure 5D, 5E** and **S6**). Interestingly, replacing glycine with proline (ZA21) preserved robust FKBP12 degradation. AP-MS analysis revealed that ZA21 recruited FBXO22 rather than DCAF10 (**Figure S7A**, **S7B** and **Table S3**). Consistent with this, ZA21-induced FKBP12 degradation was completely abolished by *FBXO22* KO but unaffected by *DCAF10* KO (**Figure S7C**). We then explored substitutions at the second amino acid position, including bulky aromatic residues such as tryptophan (ZA22) and tyrosine (ZA23), hydrophobic residues such as alanine (ZA24) and valine (ZA25), the polar residue asparagine (ZA26), and esterified aspartate (ZA27). However, none of these substitutions improved FKBP12 degradation relative to the phenylalanine-containing ZA1 (**Figure 5F, 5G** and **S6**). We further investigated the third amino acid position and found that increasing side-chain size and hydrophobicity from alanine (ZA28) to valine (ZA29) and isoleucine (ZA30) progressively enhanced degradation potency, with ZA30 achieving the DC_50_ of 9.7 nM and the D_max_ of 93% (**Figure 5H, 5I** and **S6**).

As both N-terminal propionylation and isoleucine substitution at the third amino acid position improved degradation potency (**Figure 5J**), we combined these two modifications to generate ZA31 (**Figure 5K**). Remarkably, ZA31 exhibited the highest degradation potency and efficacy among all compounds tested, achieving the DC_50_ of 1.56 nM and the D_max_ of 96% (**Figure 5L** and **5M**). ZA31 also outperformed the CRBN-recruiting FKBP12 degrader Len-SLF in our system (**Figure 5L** and **5M**). Finally, we explored additional modifications to the N-terminal propionyl group of ZA31, including replacement with butyryl (ZA32), isobutyryl (ZA33), and cyclopropanecarbonyl (ZA34) groups. However, all three modifications reduced degradation potency relative to ZA31 (**Figure S8A** and **S8B**).

### The Pr-GFI motif is a potent and selective DCAF10 recruiter for TPD

The optimized ZA31 degrader containing the N-propionyl-Gly-Phe-Ile (Pr-GFI) motif exhibited DCAF10-dependent FKBP12 degradation (**Figure 6A**). Unlikely ZA1, *DCAF10* KO completely abolished ZA31-induced FKBP12 degradation, whereas *FBXO22* KO or FBS depletion had no effect (**Figure 6A** and **S9A**). These results suggest that the Pr-GFI motif does not undergo the potential metabolic conversion observed with the Ac-GFG motif. Co-folding studies of ZA1 and ZA31 with DCAF10 revealed similar binding modes within the degron-binding pocket of DCAF10 (**Figure 6B**). Moreover, re-expression of the F172G, K256A, or F172G/K256A/I475G DCAF10 variants failed to restore ZA31-induced FKBP12 degradation in *DCAF10* KO cells, with no residual degradation activity observed (**Figure 6C**). Interestingly, the I475G mutant partially restored ZA31-induced degradation at higher concentrations (**Figure 6C**). The Boltz-2-predicted ZA31-DCAF10 co-folded structure revealed that I475 faces the third amino acid residue of the DCAF10-recruiting motif (**Figure S9B**). Our initial degrader ZA1 contains glycine at this position, and the I475G mutation creates a cavity that weakens productive interactions with ZA1 (**Figure S9B**). In contrast, the optimized degrader ZA31 contains a bulkier isoleucine residue at this position, which may occupy the cavity created by the I475G (**Figure S9B**). This potential “bump-and-hole” mechanism may explain the partially retained degradation activity of ZA31.

**Figure 6.**
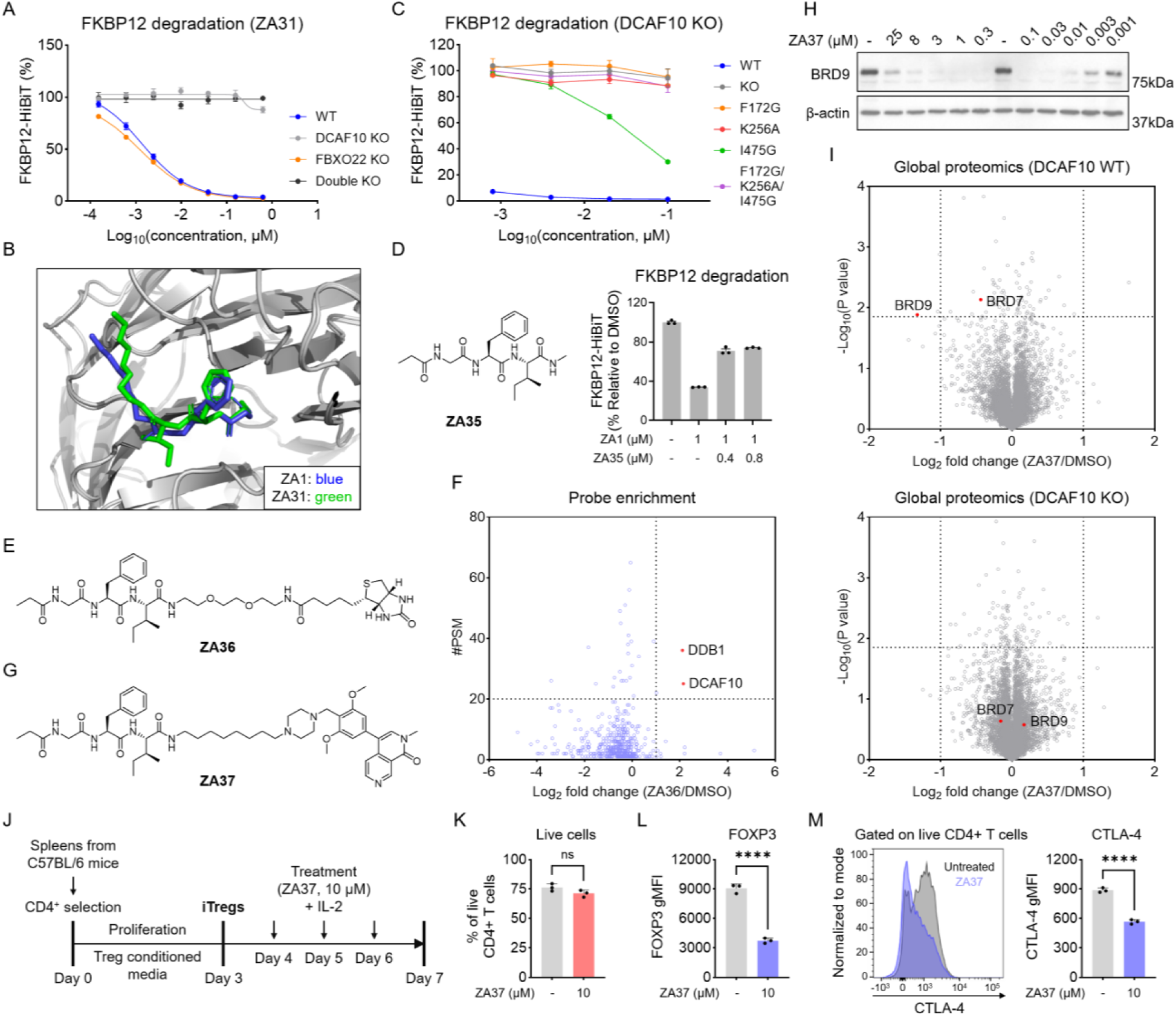
The Pr-GFI motif is a potent and selective DCAF10 recruiter for TPD. **A**. Concentration-dependent FKBP12-HiBiT degradation induced by ZA31 in parental, *DCAF10* KO, and *FBXO22* KO HEK293T cells. Cells were treated for 20 h. Data are presented as mean ± SEM (n = 3). **B**. Boltz-2-predicted binding models of the ZA1 and ZA31 recruiter moieties within the DCAF10 degron-binding pocket. **C**. Concentration-dependent FKBP12-HiBiT degradation induced by ZA31 in *DCAF10* KO cells re-expressing DCAF10 WT or mutants. Cells were treated for 20 h. Data are presented as mean ± SEM (n = 3). **D**. Structure of ZA35 and its effects on ZA1-induced FKBP12-HiBiT degradation. Cells were pretreated with 0.4 or 0.8 µM ZA35 for 1 h, followed by treatment with 1 µM ZA1 for 20 h. Data are presented as mean ± SEM (n = 3). **E**. Structure of the biotinylated Pr-GFI probe ZA36. **F**. Streptavidin pulldown analysis of DCAF10 and DDB1 enrichment by ZA36. HEK293T cell lysates were incubated with 100 nM ZA36 for 2 h at 4 °C, followed by streptavidin enrichment for 2 h. Data are representative of two independent experiments (n = 2). **G**. Structure of the DCAF10-recruiting BRD9 degrader ZA37. **H**. Concentration-dependent BRD9 degradation induced by ZA37 in HEK293T cells. Cells were treated for 24 h. **I**. Global proteomic analysis of ZA37-induced protein degradation in parental and *DCAF10* KO HEK293T cells (100 nM, 24 h). *P* values were calculated using a two-sided t-test and adjusted for multiple comparisons by the Benjamini-Hochberg method (n = 3 for DMSO; n = 2 for ZA37). **J**. Experimental scheme for evaluating the effects of ZA37 on mouse iTregs derived from splenic CD4+ T cells. **K**. Percentage of live CD4^+^ T cells following treatment with ZA37 (10 µM) or untreated for 72 h. The bar graph (n = 3) is representative of two independent experiments. The statistical significance was assessed using unpaired two-tailed Student’s t-tests. *ns*: not significant. **L**. Relative FOXP3 expression in untreated and ZA37-treated iTregs (10 µM, 72 h). The bar graph (n = 3) is representative of two independent experiments. The statistical significance was assessed using unpaired two-tailed Student’s t-tests. \*\*\*\**P* < 0.0001. **M**. Left: Representative flow cytometry histograms showing FOXP3 expression in untreated and ZA37-treated iTregs. Right: Relative CTLA-4 expression in untreated and ZA37-treated iTregs (10 µM, 72 h). The bar graph (n = 3) is representative of two independent experiments. The statistical significance was assessed using unpaired two-tailed Student’s t-tests. \*\*\*\**P* < 0.0001.

To investigate whether the Pr-GFI motif itself can effectively engage DCAF10, we synthesized a C-terminal N-methylamidated Pr-GFI peptide (ZA35, **Figure 6D**). ZA35 blocked ZA1-induced FKBP12 degradation (**Figure 6D**). We then synthesized a biotinylated Pr-GFI probe (ZA36, **Figure 6E**). Incubation of ZA36 with cell lysates, followed by streptavidin enrichment, led to selective enrichment of both DCAF10 and DDB1 (**Figure 6F** and **Table S5**). These results suggest that the Pr-GFI motif directly engages DCAF10 and has the potential to serve as a versatile chemical handle for developing additional bifunctional degraders.

To further explore the applicability of the Pr-GFI motif for degrading additional targets, we chose BRD9, a component of the noncanonical BAF (ncBAF) chromatin-remodeling complex that has emerged as a promising therapeutic target in cancers such as synovial sarcoma^33–35^. Beyond its role in cancer, BRD9 also contributes to the maintenance of regulatory T-cell (Treg) identity and immunosuppressive function, suggesting its potential relevance to cancer immunotherapy^36^. We developed a BRD9-targeting degrader, ZA37, by conjugating the dual BRD7/9 inhibitor BI-7273^39^ to the ZA31-derived recruiter through a linker containing piperazine and an aliphatic carbon chain (**Figure 6G**). ZA37 induced potent, concentration-dependent and DCAF10-dependent degradation of BRD9 in HEK293T cells (**Figure 6H** and **S9C**). BRD9 degradation was observed at concentrations as low as 1 nM (**Figure 6H**). Global proteomic analysis further confirmed DCAF10-dependent BRD9 degradation (**Figure 6I** and **Table S6**). Despite the dual BRD7/9 binding profile of BI-7273, BRD7 was not significantly depleted by ZA37 (**Figure 6I** and **Table S6**), indicating preferential degradation of BRD9 over BRD7.

To further investigate the functional consequences of BRD9 degradation, we evaluated ZA37 in mouse induced regulatory T cells (iTregs), given the role of BRD9 in maintaining Treg identity and immunosuppressive function^36^. Mouse splenic CD4^+^ T cells were isolated and differentiated into iTregs, followed by treatment with ZA37 (**Figure 6J**). We first assessed the effects of ZA37 on cell viability and identified treatment with 10 µM ZA37 for 72 h as a condition with minimal cytotoxicity (**Figure 6K** and **S9D**). Under this condition, ZA37 treatment substantially reduced the expression of FOXP3, a master transcription factor essential for Treg identity and function (**Figure 6L**). We next examined CTLA-4, an inhibitory immune checkpoint that plays a critical role in Treg-mediated immunosuppression. ZA37 treatment also significantly decreased CTLA-4 expression in iTregs (**Figure 6M**). These results suggest that DCAF10-recruiting BRD9 degraders may provide a chemical strategy to modulate Treg identity and immunosuppressive function.

## Discussion

The human proteome encodes more than 600 E3 ligases, yet only a small subset has been exploited for TPD^8^. Expanding the repertoire of TPD-competent E3 ligases remains an important goal for broadening the applicability of this strategy. Two general approaches are used to discover new TPD-competent E3 ligases and their corresponding recruiters. One is the ‘degradation-first’ approach, in which compounds are screened for their ability to induce target protein degradation, followed by deconvolution of the responsible E3 ligases^28,29,37–42^. The second is the ‘binder-first’ approach, in which E3 ligase binders are identified through rational or computational design, screening, or chemical proteomics and subsequently incorporated into bifunctional degraders^16,19,43–46^. As a ‘binder-first’ approach, degron-guided design leverages the endogenous substrate-recognition preferences of E3 ligases to facilitate the rational discovery of E3 recruiters. In this study, we applied this strategy to develop potent and selective recruiters for DCAF10, an E3 ligase not previously exploited for TPD. We further demonstrated the applicability of this recruitment strategy by achieving DCAF10-dependent degradation of multiple protein targets, including FKBP12, protein kinases, and BRD9.

DCAF10 remains a relatively understudied E3 ligase. Previous studies have implicated DCAF10 in protective roles in ovarian cancer and lung adenocarcinoma^47,48^, as well as in adenovirus-mediated suppression of antiviral responses^49^. More recently, DCAF10 was identified as a substrate receptor that recognizes N-terminal Ac-Gly degrons to promote protein degradation^23^. Our structure-activity relationship (SAR) studies of DCAF10 recruiters revealed a strong preference for hydrophobic residues, particularly phenylalanine and isoleucine at the second and third positions, respectively. These findings may help guide the identification of additional endogenous DCAF10 substrates bearing similar sequence motifs. Moreover, we identified N-propionyl-Gly-Phe-Ile as an effective recruiting motif for DCAF10. Although relatively rare, N-terminally propionylated proteins have been identified in both humans and yeast, including proteins bearing N-terminal propionylglycine^50–52^. These observations raise the intriguing possibility that N-terminal propionylation may represent an additional substrate-recognition signal for DCAF10. Identifying such substrates could provide new insights into the physiological substrate-recognition preferences and biological functions of DCAF10.

Moreover, a recent systematic analysis of the DCAF family identified DCAF10 as a high-activity CRL4 substrate receptor based on its robust CRL4 complex formation and autodegradation activity^53^. Notably, DCAF10 ranked among the highest-activity DCAFs, with several established TPD-competent E3 ligases, including CRBN, DCAF11, DCAF15, and DCAF16. These findings suggest that DCAF10 possesses favorable biochemical properties for supporting protein ubiquitination and degradation. Despite these observations, DCAF10 had not previously been exploited for the development of targeted protein degraders. Our work validates its TPD potential and establishes a chemical recruitment strategy for harnessing DCAF10 to degrade diverse protein targets.

Despite these advances, several limitations remain. Although ZA37 achieved low-nanomolar degradation potency, its structural basis of recognition has yet to be directly characterized. Furthermore, while our SAR studies improved degrader potency, this work focused on cellular degradation activity rather than the in vivo efficacy. Given the peptide-derived nature of the recruiter, further evaluation and optimization of metabolic stability and pharmacokinetics will be crucial for assessing its suitability for future in vivo applications. Our data also suggest the potential metabolic conversion of DCAF10 recruiter to terminal amino containing species that may engage FBXO22 to promote target degradation. The origin of such species and the underlying mechanism remain to be addressed. Addressing these questions will help guide the further development of DCAF10 recruiters and understand the drug stability in biological environment.

## Acknowledgements

We gratefully acknowledge support from the International Institute for Nanotechnology Seed Funding Program (X.Z.), the Sherman Fairchild Bridge Fund (X.Z.), and the Chemistry of Life Processes Predoctoral Training Program at Northwestern University (Y.A.).

