## Supplementary Information for "Degron-Informed Design of DCAF10 Recruiter for Targeted Protein Degradation"

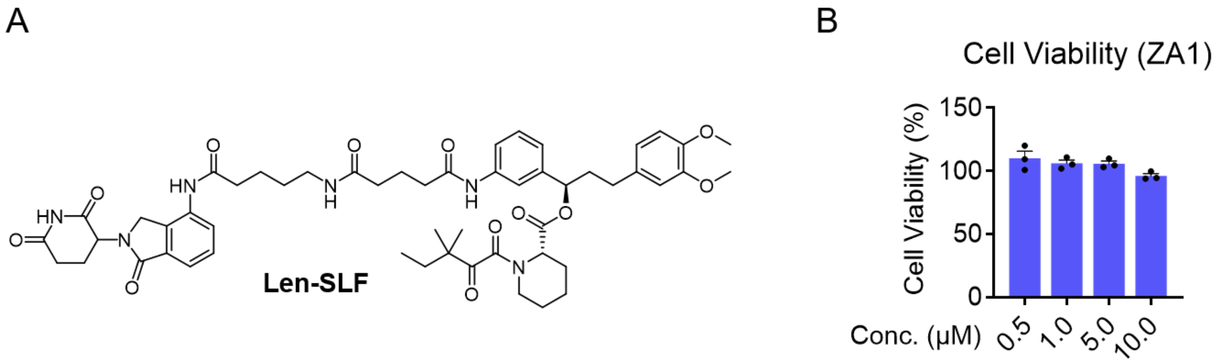


**Supplementary Figure 1. Evaluation of FKBP12 degradation and cytotoxicity of ZA1. A**. Structure of Len-SLF. **B**. Cell viability of HEK293T cells following treatment with increasing concentrations of ZA1 for 20 h. Data are presented as mean ± SEM (n = 3).

A

| Treatment time | $DC_{50}$ (nM) | $D_{max}$ (%) |
|---|---|---|
| 2 h | 1773 | 43 |
| 4 h | 2509 | 52 |
| 8 h | 1335 | 83 |
| 16 h | 703 | 87 |
| 24 h | 586 | 87 |
| 32 h | 736 | 89 |

B

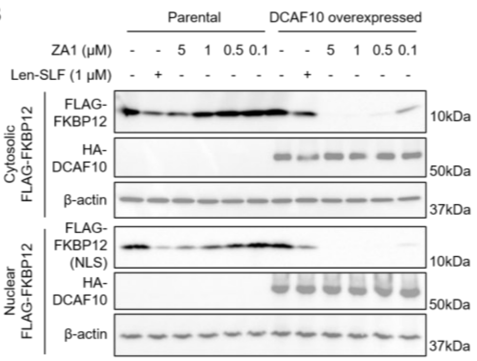


C

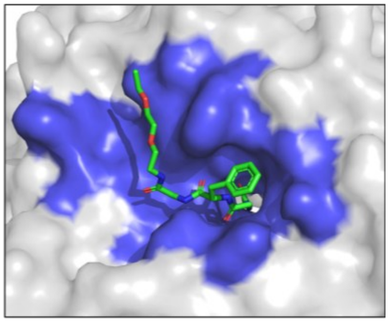

**Supplementary Figure 2. Mechanistic characterization of DCAF10-dependent FKBP12 degradation by ZA1. A**. Summary of $DC_{50}$ and $D_{max}$ values for ZA1-induced FKBP12 degradation at different treatment time points in parental FKBP12-HiBiT HEK293T cells. **B**. Western blot analysis of cytosolic and nuclear-localized FLAG-tagged FKBP12 following ZA1 treatment in HEK293T cells with or without DCAF10 overexpression for 20 h. **C**. Boltz-2-predicted binding model of the ZA1-derived recruiter within the DCAF10 degron-binding pocket. The recruiter is shown as green sticks, and the DCAF10 binding pocket is highlighted in blue.

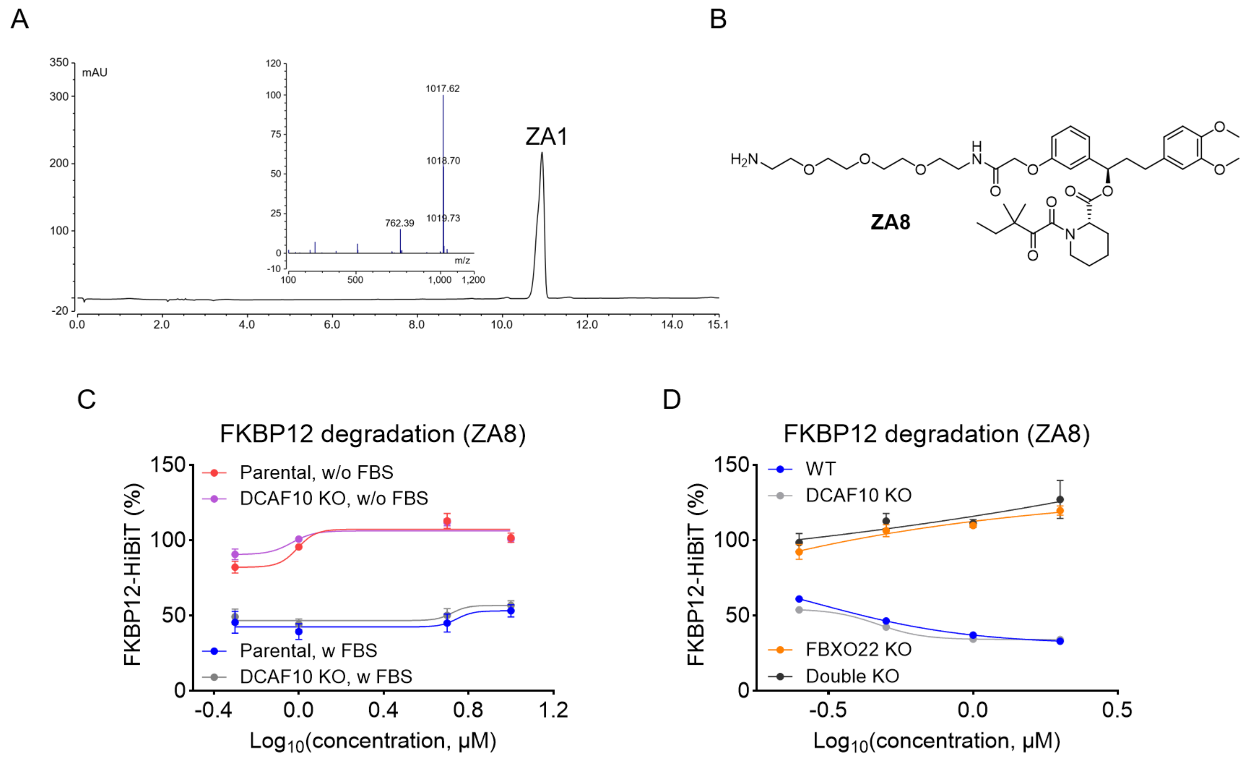


**Supplementary Figure 3. DCAF10 and FBXO22 differentially contribute to ZA1-induced FKBP12 degradation. A**. LC-MS analysis of ZA1 to assess compound purity. **B**. Structure of ZA8, an analog of the previously reported FBXO22-recruiting degrader SP3N. **C**. ZA8-induced FKBP12 degradation in parental and *DCAF10* KO HEK293T cells cultured with or without FBS. Cells were treated for 20 h. Data are presented as mean ± SEM (n = 3). **D**. ZA8-induced FKBP12 degradation in parental, *DCAF10* KO, *FBXO22* KO, and *DCAF10*/*FBXO22* double KO HEK293T cells. Cells were treated for 20 h. Data are presented as mean ± SEM (n = 3).

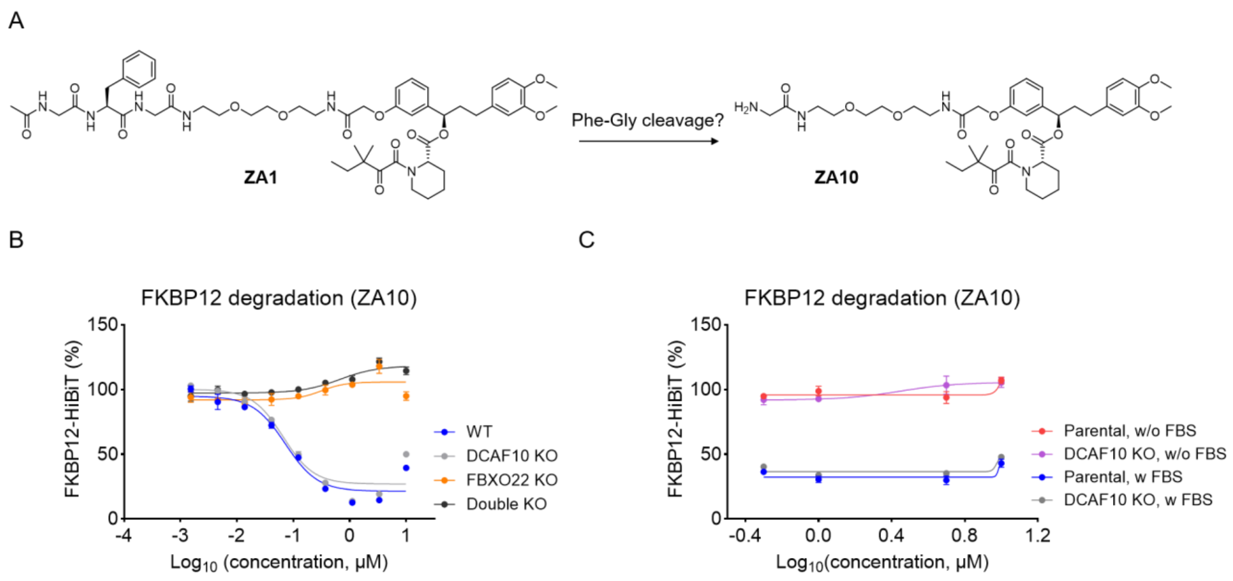


**Supplementary Figure 4. Characterization of FBXO22-dependent FKBP12 degradation induced by ZA10. A**. Structure of ZA10. **B**. Dose-dependent FKBP12 degradation induced by ZA10 in parental, *DCAF10* KO, *FBXO22* KO, and *DCAF10*/*FBXO22* double KO HEK293T cells. Cells were treated for 20 h. Data are presented as mean ± SEM (n = 3). **C**. Dose-dependent FKBP12 degradation induced by ZA10 in parental and *DCAF10* KO HEK293T cells cultured in the presence or absence of FBS. Cells were treated for 20 h. Data are presented as mean ± SEM (n = 3).

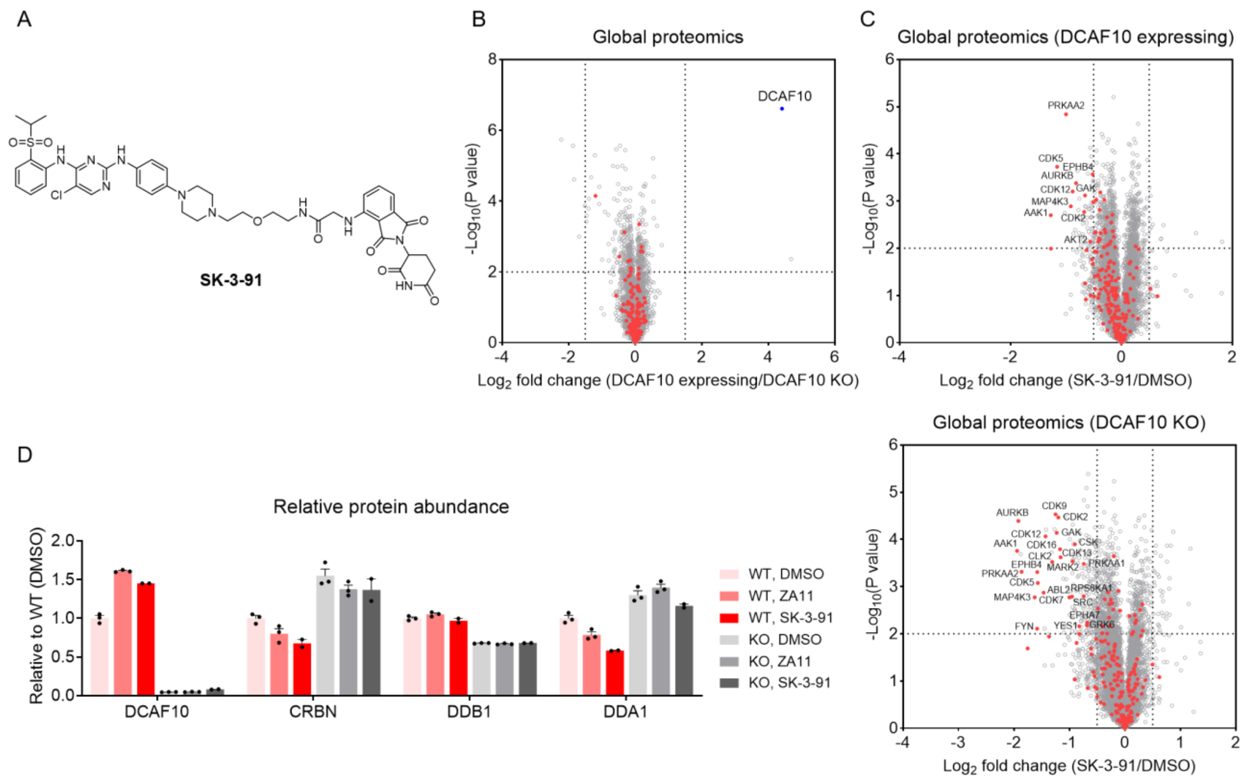


**Supplementary Figure 5. Proteomic characterization of DCAF10- and CRBN-dependent kinase degradation. A**. Structure of the CRBN-recruiting kinase degrader SK-3-91. **B**. Global proteomic comparison of DCAF10-expressing and *DCAF10* KO HEK293T cells. Protein kinases are highlighted in red. *P* values were calculated using a two-sided t-test and adjusted for multiple comparisons by the Benjamini-Hochberg method (n = 3). **C**. Global proteomic analysis of SK-3-91-treated *DCAF10* KO HEK293T cells with or without DCAF10 re-expression (100 nM, 24 h). Protein kinases are highlighted in red. *P* values were calculated using a two-sided t-test and adjusted for multiple comparisons by the Benjamini-Hochberg method (n = 3 for DMSO; n = 2 for SK-3-91). **D**. Relative protein abundance of DCAF10, CRBN, DDB1, and DDA1 in *DCAF10* KO HEK293T cells with or without DCAF10 re-expression, determined by global proteomic analysis. Data are presented as mean ± SEM (n = 3).

Ac-G-F-G

| | $DC_{50}$ (μM) | $D_{max}$ (%) |
|---|---|---|
| ZA1 | 0.30 | 89 |
| ZA13 | 0.56 | 81 |
| ZA14 | 1.0 | 70 |
| ZA15 | - | 57 |
| ZA16 | - | 41 |
| ZA17 | - | 59 |
| ZA18 | 0.098 | 86 |

Ac-G-F-G

| | $DC_{50}$ (μM) | $D_{max}$ (%) |
|---|---|---|
| ZA1 | 0.30 | 89 |
| ZA19 | 4.7 | 79 |
| ZA20 | 1.1 | 67 |
| ZA21 | 0.36 | 81 |

Ac-G-F-G

| | $DC_{50}$ (μM) | $D_{max}$ (%) |
|---|---|---|
| ZA1 | 0.30 | 89 |
| ZA22 | 2.1 | 69 |
| ZA23 | 1.2 | 82 |
| ZA24 | 4.7 | 74 |
| ZA25 | 0.48 | 80 |
| ZA26 | - | 69 |
| ZA27 | - | 53 |

Ac-G-F-G

| | $DC_{50}$ (nM) | $D_{max}$ (%) |
|---|---|---|
| ZA1 | 235 | 88 |
| ZA28 | 91 | 88 |
| ZA29 | 12.8 | 91 |
| ZA30 | 9.7 | 93 |

**Supplementary Figure 6.** Summary of $DC_{50}$ and $D_{max}$ values for ZA1-derived FKBP12 degraders.

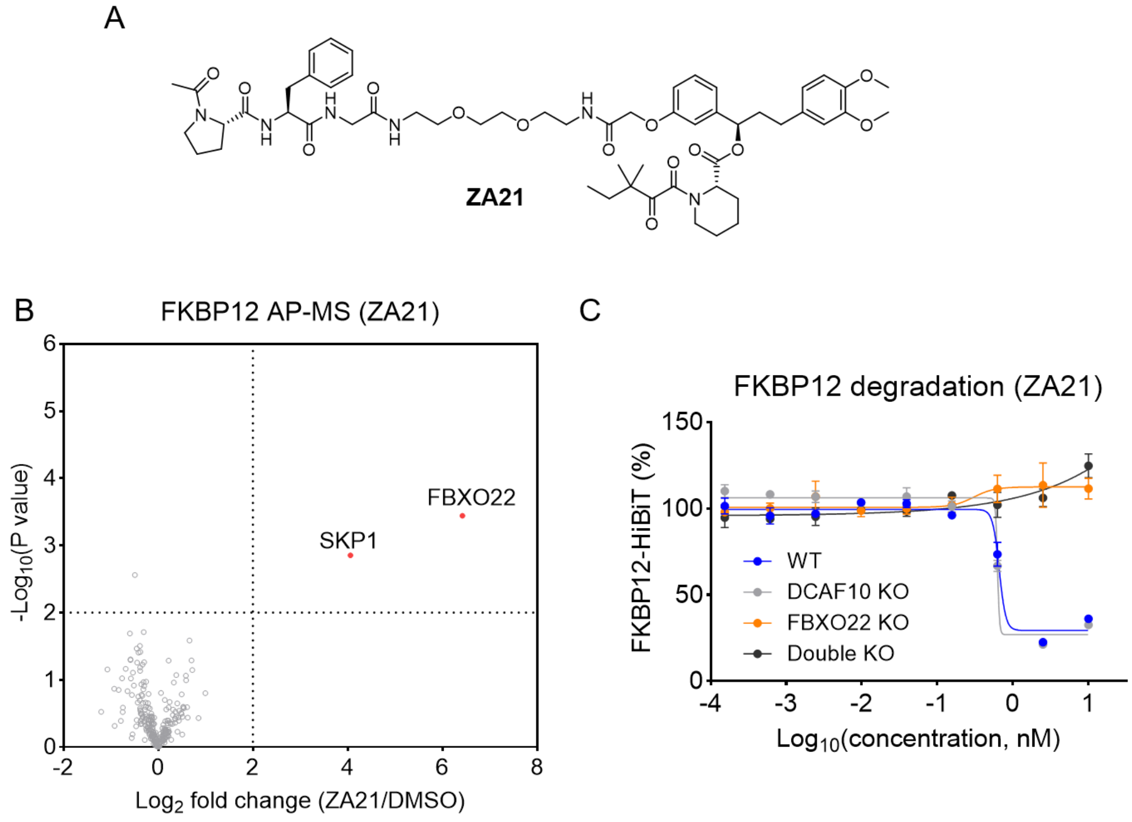


**Supplementary Figure 7. Characterization of FBXO22-dependent FKBP12 degradation induced by ZA21. A**. Structure of ZA21. **B**. AP-MS analysis of ZA21-induced protein interactions with FLAG-FKBP12 in HEK293T cells. Cells were pretreated with 100 nM BTZ for 1 h, followed by treatment with DMSO or 5 μM ZA1 for 3 h. *P* values were calculated using a two-sided t-test and adjusted for multiple comparisons by the Benjamini-Hochberg method (n = 2). **C**. Dose-dependent FKBP12 degradation induced by ZA21 in parental, *DCAF10* KO, *FBXO22* KO, and *DCAF10*/*FBXO22* double KO HEK293T cells. Cells were treated for 20 h. Data are presented as mean ± SEM (n = 3).

A

| | | $DC_{50}$ (nM) | $D_{max}$ (%) |
|---|---|---|---|
| | ZA1 | 488 | 88 |
| 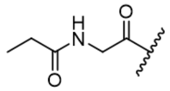 | ZA31 | 0.9 | 96 |
| 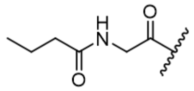 | ZA32 | 7.9 | 94 |
| 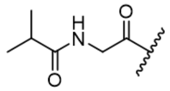 | ZA33 | 51.2 | 86 |
| 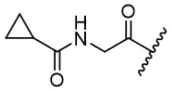 | ZA34 | 37.9 | 89 |

B

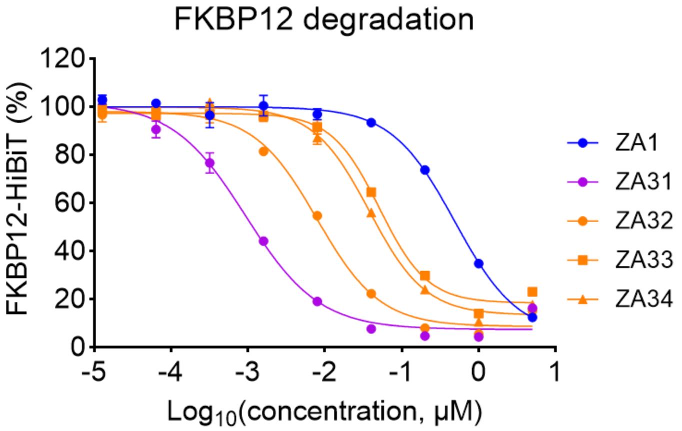


**Supplementary Figure 8. Further optimization of the ZA31-derived DCAF10 recruiter. A**. Summary of $DC_{50}$ and $D_{max}$ values for ZA31-derived FKBP12 degraders. **B**. Dose-dependent FKBP12 degradation induced by ZA31, ZA32, ZA33, and ZA34 in parental FKBP12-HiBiT HEK293T cells. Cells were treated for 20 h. Data are presented as mean ± SEM (n = 3).

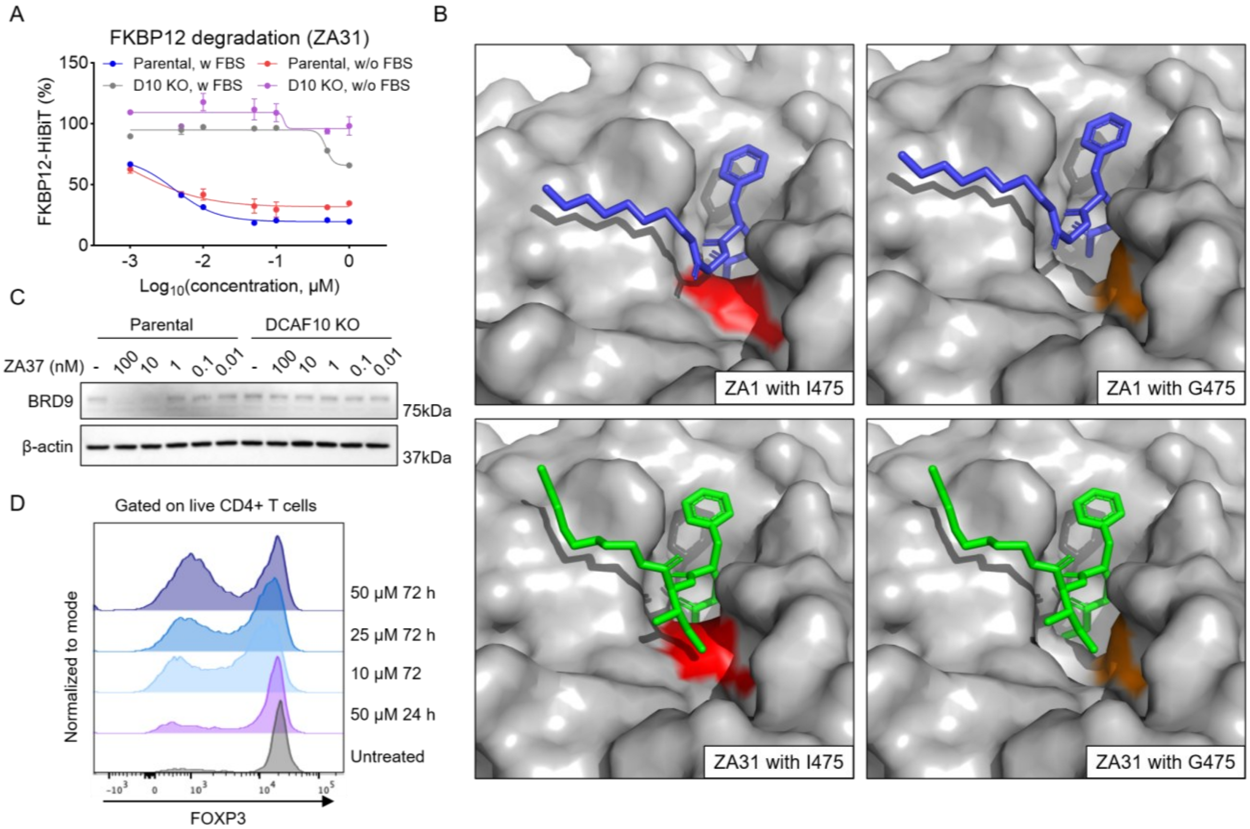


**Supplementary Figure 9. Characterization of DCAF10-dependent degradation by ZA31 and ZA37. A**. Dose-dependent FKBP12 degradation induced by ZA31 in parental and *DCAF10* KO FKBP12-HiBiT HEK293T cells cultured in the presence or absence of FBS. Cells were treated for 20 h. Data are presented as mean ± SEM (n = 3). **B**. Boltz-2-predicted binding models of the ZA1- and ZA31-derived recruiters within the DCAF10 degron-binding pocket containing either wild-type I475 or the I475G mutation. Residue 475 is highlighted in red (I475) or orange (G475). **C**. Western blot analysis of BRD9 degradation induced by ZA37 in parental and *DCAF10* KO HEK293T cells. Cells were treated for 24 h. **D**. Flow cytometry histograms showing the effects of ZA37 treatment on iTreg cell viability.

### Methods

#### Reagents

Anti-HA (clone #C29F4, cat. #3724, dilution 1:5000), HRP-linked rabbit IgG (cat. #7074, dilution 1:5000), and anti-BRD9 (E9R2I) Rabbit Monoclonal Antibody (cat. #58906S, 1:1000) were purchased from Cell Signaling Technology. Anti-FLAG HRP antibody (clone M2, cat. #A8592, dilution 1:5000), anti-FLAG affinity gel (clone M2, cat. #A2220), anti-HA HRP antibody (clone #3F10, cat. #12013819001, dilution 1:5000), and cOmplete protease inhibitor cocktail (cat. #11873580001) were purchased from Sigma-Aldrich. Anti-β-Actin HRP antibody (clone #C4, cat. #sc- 47778, dilution 1:1000) was purchased from Santa Cruz Biotechnology. Cas9 endonuclease was purchased from Integrated DNA Technologies (cat. #1081061). FuGene 6 (cat. #E2692) transfection reagent, Nano-Glo HiBiT Lytic Detection System (cat. #N3040), Cell Titer Glo (cat. #G7573) and sequencing grade trypsin/Lys-C mix (cat. #V5071) were purchased from Promega. Blastcidin (cat. #ant-bl-05) was purchased from InvivoGen. Bortezomib (cat. #HY-10227) and SK-3-91 (cat. #HY-137341) were purchased from MedChemExpress. MLN4924 (cat. #15217) and SLF (cat. #10007974) were purchased from Cayman Chemical. DC protein assay kit (cat. #5000112) was purchased from Bio-Rad. Pierce high pH reversed-phase peptide fractionation kit (cat. #84868), Pierce ECL Western Blotting Substrate (cat. #32209), SuperSignal West Pico PLUS Chemiluminescent Substrate (cat. #34580) and Tandem Mass Tag (TMT) isobaric label reagent (cat. #90110) were purchased from Thermo Scientific. Opti-MEM I Reduced Serum Medium (cat. #31985070) and poly-D-lysine (0.1 mg/mL, cat. #A3890401) were purchased from Gibco.

#### Cell Lines

HEK293T cells were obtained from ATCC. HEK293T were cultured in Dulbecco's Modified Eagle Medium (DMEM, Corning, cat. #15013CV) with 10% (v/v) fetal bovine serum (FBS, Omega Scientific, cat. #FB-01) and L-glutamine (2 mM, Gibco, cat.

#25030081). All the cell lines were tested negative for mycoplasma contamination using Universal Mycoplasma Detection Kit (ATCC, cat. #30-1012K). FKBP12-HiBiT HEK293T was generated as previously described[1].

**Generation of *DCAF10* and *FBXO22* Knockout Cells**

*DCAF10* and *FBXO22* knockout FKBP12-HiBiT HEK293T cells were generated by electroporating Cas9 ribonucleoprotein complexes containing two or three mixed sgRNAs using a 4D-Nucleofector (Lonza Bioscience) with pulse code CM-130. To validate the knockout, genomic DNA was extracted using the PureLink Genomic DNA Mini Kit, and the *DCAF10* and *FBXO22* loci were amplified by polymerase chain reaction (PCR). PCR products were analyzed by agarose gel electrophoresis and Sanger sequencing. The sgRNA sequences and sequencing primers are listed below.

*DCAF10* sgRNA#1: ATGTTGATTTCAACGTCCTC
*DCAF10* sgRNA#2: CTTTACATGGTCACACTAGC
*DCAF10* sgRNA#3: TGACACTACAATAGCACTAT
*FBXO22* sgRNA#1: TCAACACGAAGGTGCTCCGC
*FBXO22* sgRNA#2: GGACCCATCGGAGCGTAACC
*DCAF10* sequencing primers#1: 5'-CTCTCGGCAATCAAGTCTTCA-3'
*DCAF10* sequencing primers#2: 5'-CCTAAAGCCAGTTTCCTTCTTGA-3'
*FBXO22* sequencing primers#1: 5'-TATGCTGGCGTAATCGGGTT-3'
*FBXO22* sequencing primers#2: 5'-CCTGCAAATGACTGTGCATGACA-3'

**Cloning, Mutagenesis, and Lentiviral Transduction**

The pCDH-CMV-MCS-EF1-Blast construct encoding human DCAF10 with an N-terminal HA tag was purchased from GenScript. The FLAG-FKBP12 expression construct was prepared as previously reported[2]. DCAF10 mutants (F172G, K256A, I475G, F172G/K256A/I475G) were generated by Gibson Assembly (New England Biolabs). For lentivirus production, HEK293T cells were transfected with the expression construct together with the packaging plasmid psPAX2 and envelope plasmid pMD2.G using FuGENE 6. Culture supernatants containing lentiviral particles were collected 48 hours later and filtered through 0.45 µm Millex-HV sterile syringe filters (MilliporeSigma). The filtered supernatants were applied to HEK293T cells for transduction, followed by selection with blastcidin at 10 µg/mL for 7 days to generate stable polyclonal cell lines.

**Cell Viability Assay**

FKBP12-HiBiT HEK293T cells were seeded in 100 µL of DMEM in a 96-well clear-bottom white plate (Corning, cat. #3610, 3000 cells per well). Serial compound dilutions were prepared in 100 µL of DMEM and applied for 72 hours, with DMSO maintained at a final concentration of 0.5%. Viability was measured by adding 50 µL of CellTiter-Glo reagent (Promega, cat. #G7573) per well and allowing incubation for 10 minutes at room temperature before recording luminescence on a CLARIOstar Plus microplate reader (BMG LABTECH).

**HiBiT Assay**

FKBP12-HiBiT-expressing HEK293T cells were plated in white 96-well plates (5,000 cells per well). At the end of treatment, HiBiT luminescence was assessed by adding 50 µL of Nano-Glo HiBiT Lytic Detection System reagent (Promega, cat. #N3040) to each well. After 10 minute incubation at room temperature, luminescence was recorded on a CLARIOstar Plus microplate reader.

**Cell Lysis and Western Blot**

Cell lysates were prepared in RIPA buffer (25 mM Tris-HCl at pH 7.6, 150 mM NaCl, 1% NP-40, 1% sodium deoxycholate, and 0.1% SDS), with Universal Nuclease added before use. Protein concentration were quantified using the DC protein assay kit, after which lysates were mixed with Laemmli sample buffer and incubated at 95°C for 10 minutes. Samples were analyzed on 4-20% Novex Tris-Glycine gels (Thermo Scientific, cat. #EC62255BOX) and transferred to PVDF membranes (Bio-Rad, cat. #1620177). 5% non-fat milk in TBST (20 mM Tris-HCl, pH 7.6, 150 mM NaCl, and 0.1% Tween-20) was used for both membrane blocking and primary antibody dilution. Blocking proceeded for 30 minutes at room temperature. Primary antibodies were diluted in the same blocking solution. Incubation with anti-FLAG, anti-HA, and anti-β-actin antibodies was performed for 1 hour at room temperature. Other primary antibodies were incubated overnight at 4°C. Membranes were washed with TBST and incubated with secondary antibodies for 1 hour at room temperature. Membranes were washed three times with TBST before detection with chemiluminescent reagent on a ChemiDoc MP imaging system (Bio-Rad).

**Quantitative Global Proteomics**

Proteins were extracted by sonicating cells in 100 µL PBS containing cOmplete protease inhibitor cocktail. Sonication consisted of three cycles, each comprising five pulses at 40% amplitude. Following protein quantification with the DC assay, an aliquot containing 100 µg of protein in 50 µL of PBS was mixed with 50 µL of 12 M urea in PBS. Reduction was carried out at 65°C for 15 minutes after the addition of 5 µL of 200 mM DTT. Samples were then alkylated by adding 5 µL of 400 mM iodoacetamide in water and incubate at 37°C for 30 minutes in the dark. Following dilution with 300 µL of PBS, samples were digested with 2 µg of trypsin/LysC for 16 hours at 37°C. For each TMT channel, 8.5 µg of peptides was resuspended in 35 µL of PBS and supplemented with 9 µL of acetonitrile. The samples were then reacted with TMT reagents (5 µL, 20 µg/µL in dry acetonitrile) at room temperature for 1 hour. Reactions were quenched by adding 6 µL of 5% hydroxylamine and 2.5 µL of formic acid. Labeled samples were pooled and fractionated into ten fractions by high-pH reversed-phase peptide fractionation kit. Fractions were dried by vacuum centrifugation before LC-MS analysis on an Orbitrap Eclipse Tribrid mass spectrometer coupled to a Vanquish Neo UHPLC system.

Dried fractions were dissolved in 20 µL buffer contain 95% $H_2O$, 5% acetonitrile, 0.1% formic acid and loaded onto an EASY-Spray analytical column (C18, 2 µm particles, 75 μm i.d., 150 mm length) and separated at a flow rate of 250 nL/min using a multistep gradient. Mobile phases A and B consisted of 0.1% formic acid in water and 0.1% formic acid in 80% acetonitrile, respectively. A 180-min gradient consisted of an initial 15 min at 5% B, followed by a linear increase to 45% B over 140 min and a final increase to 100% B over 25 min. The nanoelectrospray voltage was set to 1.5 kV. Full MS scans were collected in the Orbitrap at a resolution of 60,000 over an m/z range of 375-1600, with the RF lens set to 60%, standard automatic gain control (AGC), and automatic maximum injection time. For MS2, quadrupole isolation of precursor ions used a 0.7 m/z window. Precursors underwent higher-energy collisional dissociation (HCD) at 27% collision energy, and the resulting MS2 spectra were recorded in the ion trap using standard AGC and a maximum injection time of 35 ms. Each MS2 scan was followed by synchronous precursor selection (SPS) of up to ten fragment ions for MS3 acquisition. These ions were further fragmented by HCD at 55% collision energy and analyzed in the Orbitrap at 60,000 resolution, with an AGC target of 250% and a maximum injection time of 200 ms. Raw spectra were acquired using Xcalibur software (v4.5.445.18, Thermo Scientific).

Proteome Discoverer (v2.5, Thermo Scientific) was used for peptide identification and TMT-based quantification. Spectra were searched against the UniProt human reference proteome (UP000005640_9606_Human.fasta) using Sequest HT with tryptic specificity. Precursor and fragment mass tolerances were set to 10 ppm and 0.6 Da, respectively. The search included fixed modifications for carbamidomethylated cysteine residues and TMT-labeled peptide N-termini and lysine residues. Methionine oxidation and protein N-terminal acetylation were set as variable modifications. Peptide-spectrum matches were assessed using Percolator, with a 1% false discovery rate applied at both the peptide and protein levels. For quantification, reporter ion signals were integrated within a 20 ppm window in SPS-MS3 spectra. Protein abundances were calculated from unique and razor peptides.

**Affinity Purification Mass Spectrometry (AP-MS)**

Proteins were extracted by sonicating in NP-40 lysis buffer containing 25 mM Tris-HCl at pH 7.6, 150 mM NaCl, 10% glycerol, 1% NP-40, and cOmplete protease inhibitor cocktail. Sonication consisted of three cycles, each comprising five pulses at 25% amplitude. Lysates were clarified at 16,000 × g for 10 minutes at 4 °C. Each supernatant was then combined with 30 µL of FLAG affinity agarose slurry and rotated for 2 hours at 4°C. The agarose was washed four times with immunoprecipitation wash buffer and twice with PBS. Captured proteins were released by incubating the agarose in 8 M urea in PBS at 65°C for 10 minutes. The recovered proteins were subjected to sequential reduction and alkylation using 12.5 mM DTT at 65°C for 15 minutes and 25 mM iodoacetamide at 37°C for 30 minutes, respectively. The urea concentration was subsequently lowered to 2 M by dilution with PBS, and 2 µg of trypsin/LysC was added for digestion at 37°C for 16 hours. TMT labeling was performed for 1 hour at room temperature using 6 µL of reagent dissolved in anhydrous acetonitrile. To quench the reaction, 6 µL of 5% hydroxylamine and 8.5 µL of formic acid were added. Peptides from the individual samples were combined and cleaned up on a Sep-Pak C18 cartridge (Waters, cat. #WAT054955), followed by vacuum drying. The pooled peptides were analyzed using the LC-MS workflow and data analysis procedures described above.

For ZA36 AP-MS, DMSO or 100nM ZA36 was added to the cell lysate and rotate at room temperature for 2 hours. Then streptavidin beads affinity agarose (50 µL slurry per sample) was added to the lysates and incubated with rotation for 2 hours at 4 °C. The agarose was washed four times with immunoprecipitation wash buffer and twice with PBS. Captured proteins were released by incubating the agarose in 8 M urea in PBS at 65°C for 10 minutes. The recovered proteins were subjected to sequential reduction and alkylation using 12.5 mM DTT at 65°C for 15 minutes and 25 mM iodoacetamide at 37°C for 30 minutes, respectively. The urea concentration was subsequently lowered to 2 M by dilution with PBS, and 2 µg of trypsin/LysC was added for digestion at 37 °C for 16 hours. The samples were then added with 8.5 µL of formic acid and desalted on a Sep-Pak C18 cartridge, then analyzed by LC-MS in label free quantification.

**Isolation and Culture of Primary Mouse iTreg Cells**

Spleens from C57BL/6 mice (000664, The Jackson Laboratory) were harvested and dissociated in a solution containing 1x PBS with 2% FBS and 1 mM EDTA. $CD4^+$ T cells were collected and separated using a $CD4^+$ negative selection magnetic bead kit (StemCellTechnologies, 19752). Cells were then plated at 0.5 $\times 10^6$ cells/mL in a 24-well plate precoated with plate-bound αCD3 and αCD28 antibodies at 3 mg/mL and 5 mg/mL, respectively (Invitrogen, 16-0031-86 and 16-0281-86). Cells were plated with Treg cell-conditioned media including complete RPMI with 10% FBS, 1% penicillin/streptomycin, 1% non-essential amino acids, 1% sodium-pyruvate, 1% glutamine, 0.5% HEPES, plus the following cytokines: 5 ng/mL IL-2, 2 μg/mL anti-IL-4, 2 μg/mL anti-IFNγ, and 5 ng/mL TGF-β, and polarized in conditioned media for 72 h at 5% $CO_2$ at 37°C. Cells were then harvested and replated in high-dose IL-2 resting medium (2,000 IU/mL IL-2), with or without treatment with ZA37, for subsequent experimental analysis.

**Flow Cytometry of iTregs**

iTreg suspensions were incubated with Fixable Viability Dye (Thermo Fisher Scientific), followed by staining with surface-directed antibodies, including CD4 (GK1.5), CD44 (IM7), and CD25 (PC61), for 30 min at room temperature. For the detection of intracellular proteins, the FOXP3/Transcription Factor Staining Buffer Set (Thermo Fisher Scientific, 00-5223-56) was used. Intracellular antibodies included FOXP3 (FJK-16s) and CTLA-4 (UC10-4B9). Flow cytometry analyses were conducted with FlowJo v10.10.

**Statistical Analysis**

Quantitative data are displayed as scatter plots with the mean ± standard error of the mean (SEM). Statistical significance in proteomics experiments was assessed using two-sided Student's t-tests, and *P* values were adjusted for multiple comparisons using the Benjamini-Hochberg procedure. Statistical significance was defined as $P < 0.05$.

Significance levels were indicated as follows: *$P < 0.05$, **$P < 0.01$, ***$P < 0.001$, ****$P < 0.0001$, and *ns* (not significant).

### Data Availability Statement

The mass spectrometry proteomics data were deposited to the ProteomeXchange Consortium through the PRIDE partner repository with the dataset identifier PXD086915. The data supporting the findings of this study are available within the article and Supplementary Information.

### Synthesis of ZA1, ZA6, ZA8, and ZA10

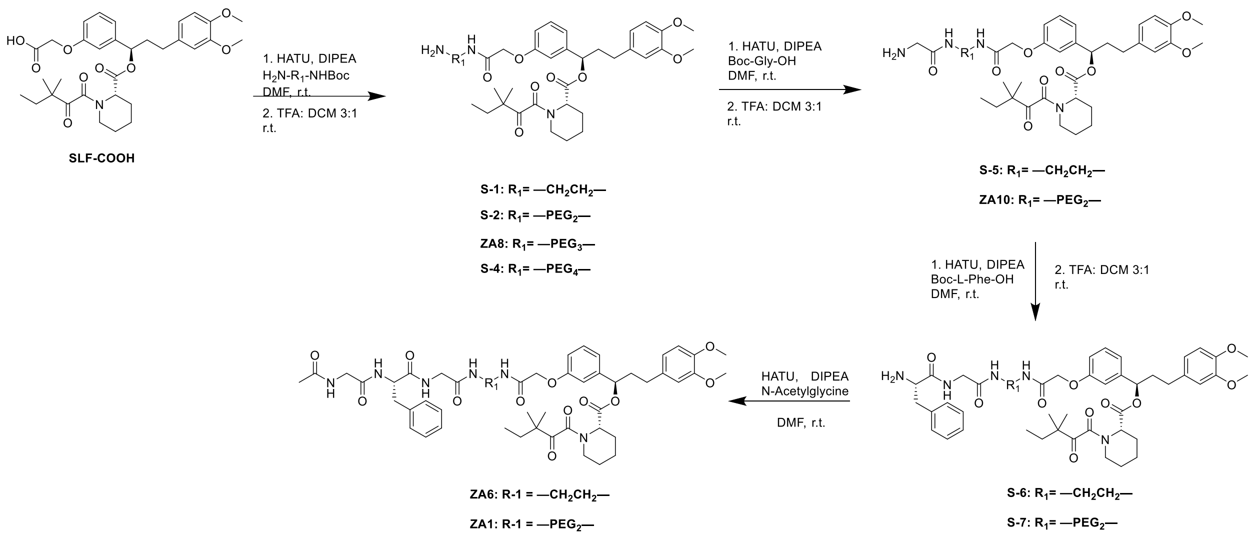


**ZA1:** *(R)-1-(3-(((S)-7-benzyl-2,5,8,11,22-pentaoxo-15,18-dioxa-3,6,9,12,21-pentaazatricosan-23-yl)oxy)phenyl)-3-(3,4-dimethoxyphenyl)propyl (S)-1-(3,3-dimethyl-2-oxopentanoyl)piperidine-2-carboxylate*

**ZA10:** *(R)-1-(3-((14-amino-2,13-dioxo-6,9-dioxa-3,12-diazatetradecyl)oxy)phenyl)-3-(3,4-dimethoxyphenyl)propyl (S)-1-(3,3-dimethyl-2-oxopentanoyl)piperidine-2-carboxylate*

SLF-COOH was synthesized according to the previously reported method[3].

Step 1: SLF-COOH (70mg, 0.120 mmol, 1.0 eq), HATU (59.3mg, 0.156 mmol, 1.3 eq), and $H_2N$-PEG2-NHBoc (59.6mg, 0.240 mmol, 2.0 eq) were dissolved in dry DMF (3.0 mL) at room temperature. DIPEA (105 µL, 0.600 mmol, 5.0 eq) was added to the reaction and stirred for 1 h. Upon completion, the reaction was concentrated, extracted with DCM (20 mL) and washed with brine (1 x 20 mL). The organic layer was dried by $Na_2SO_4$, concentrated, and the resulting residue was used without further purification. The crude product was dissolved in DCM (3.0 mL), TFA (1.0 mL) was added to the reaction and stirred for 0.5 h. The reaction mixture was then concentrated and purified by flash chromatography to provide S-2 as a colorless oil (77.3 mg, 0.095 mmol, 79% over 2 steps).

$^1$H NMR (500 MHz, $CDCl_3$) δ 7.93 (s, 2H), 7.33 – 7.27 (m, 1H), 7.01 – 6.92 (m, 2H), 6.88 – 6.76 (m, 2H), 6.71 – 6.65 (m, 2H), 5.82 – 5.71 (m, 1H), 5.29 (s, 1H), 4.57 – 4.51 (m, 2H), 3.88 – 3.82 (m, 6H), 3.79 – 3.70 (m, 2H), 3.66 – 3.54 (m, 9H), 3.40 – 3.30 (m, 1H), 3.24 – 3.15 (m, 1H), 2.67 – 2.49 (m, 2H), 2.39 (d, *J* = 13.9 Hz, 1H), 2.29 – 2.18 (m, 1H), 2.11 – 2.01 (m, 1H), 1.86 – 1.59 (m, 5H), 1.53 – 1.46 (m, 2H), 1.44 – 1.30 (m, 2H), 1.20 – 1.10 (m, 6H), 0.92 – 0.77 (m, 3H).

Step 2 (ZA10): S-2 (77.3 mg, 0.095 mmol, 1.0 eq), HATU (65.88 mg, 0.173 mmol, 1.6 eq), and Boc-Gly-OH (22.76 mg, 0.130 mmol, 1.2 eq) were dissolved in dry DMF (2.0 mL) at room temperature. DIPEA (59 µL, 0.323 mmol, 3.0 eq) was added to the reaction and stirred for 0.5 h. Upon completion, the reaction was concentrated, extracted with DCM (20 mL) and washed with brine (1 x 20 mL). The organic layer was dried by $Na_2SO_4$, concentrated, and the resulting residue was used without further purification. The crude product was dissolved in DCM (3.0 mL), TFA (1.0 mL) was added to the reaction and stirred for 0.5 h. The reaction mixture was then concentrated and purified by flash chromatography to provide ZA10 as a colorless oil (63 mg, 0.082 mmol, 75% over 2 steps).

$^1$H NMR (500 MHz, $CDCl_3$) δ 8.04 (s, 1H), 7.32 – 7.26 (m, 1H), 7.24 – 7.15 (m, 1H), 7.01 – 6.88 (m, 2H), 6.87 – 6.81 (m, 1H), 6.80 – 6.75 (m, 1H), 6.71 – 6.64 (m, 2H), 5.79 – 5.71 (m, 1H), 5.28 (s, 1H), 4.49 (s, 2H), 3.91 – 3.71 (m, 8H), 3.60 – 3.48 (m, 10H), 3.46 – 3.31 (m, 3H), 3.22 – 3.13 (m, 1H), 2.65 – 2.48 (m, 2H), 2.36 (d, J = 13.6 Hz, 1H), 2.29 – 2.18 (m, 1H), 2.10 – 1.97 (m, 1H), 1.80 – 1.58 (m, 5H), 1.54 – 1.42 (m, 1H), 1.40 – 1.30 (m, 1H), 1.22 – 1.10 (m, 6H), 0.83 (dt, J = 34.3, 7.4 Hz, 3H).

$^{13}$C NMR (126 MHz, $CDCl_3$) δ 208.13, 169.87, 167.49, 157.51, 149.02, 147.49, 141.99, 133.53, 130.19, 120.32, 120.11, 114.10, 113.66, 111.90, 111.50, 76.72, 70.23, 70.18, 69.86, 69.61, 67.31, 56.06, 55.97, 53.57, 51.46, 46.84, 44.32, 41.26, 39.59, 38.99, 38.31, 32.58, 31.39, 29.84, 26.54, 25.04, 23.54, 23.31, 21.29, 8.92.

HRMS (ESI) m/z calcd for $C_{40}H_{59}N_4O_{11}^+$ for $[M+H]^+$: 771.4175, found 771.4175.

Step 3: ZA10 (45mg, 0.058 mmol, 1.0 eq), Boc-L-phenylalanine (18.6 mg, 0.075 mmol, 1.2 eq), and HATU (35.51 mg, 0.093 mmol, 1.6 eq) were dissolved in dry DMF (2.0 mL) at room temperature. DIPEA (30.5 µL, 0.175 mmol, 3.0 eq) was added to the reaction and stirred for 0.5 h. Upon completion, the reaction was concentrated, extracted with DCM (20 mL) and washed with brine (1 x 20 mL). The organic layer was dried by $Na_2SO_4$, concentrated, and the resulting residue was used without further purification. The crude product was dissolved in DCM (3.0 mL), TFA (1.0 mL) was added to the reaction and stirred for 0.5 h. The reaction mixture was then concentrated and purified by flash chromatography to provide S-7 as a colorless oil (38 mg, 0.041 mmol, 71% over 2 steps).

$^1$H NMR (500 MHz, $CDCl_3$) δ 8.61 (s, 1H), 7.42 (s, 1H), 7.32 – 7.14 (m, 8H), 7.01 – 6.87 (m, 2H), 6.85 – 6.80 (m, 1H), 6.80 – 6.74 (m, 1H), 6.71 – 6.63 (m, 2H), 5.79 – 5.71 (m, 1H), 5.30 – 5.27 (m, 1H), 4.50 – 4.36 (m, 3H), 4.05 – 3.93 (m, 1H), 3.83 (d, *J* = 5.6 Hz, 6H), 3.73 – 3.39 (m, 12H), 3.38 – 3.24 (m, 3H), 3.23 – 3.11 (m, 3H), 2.66 – 2.47 (m, 2H), 2.35 (d, *J* = 13.1 Hz, 1H), 2.29 – 2.16 (m, 1H), 2.09 – 1.98 (m, 1H), 1.80 – 1.57 (m, 5H), 1.43 – 1.39 (m, 1H), 1.38 – 1.35 (m, 1H), 1.21 – 1.09 (m, 6H), 0.91 – 0.75 (m, 3H).

ZA1: S-7 (38 mg, 0.041 mmol, 1 eq), N-Acetylglycine (5.8 mg, 0.050 mmol, 1.2 eq), and HATU (25.2 mg, 0.066 mmol, 1.6 eq) were dissolved in dry DMF (1.5 mL) at room temperature. DIPEA (22 μL, 0.124 mmol, 3.0 eq) was added to the reaction and stirred for 0.5 h. Upon completion, the reaction was concentrated, extracted with DCM (20 mL) and washed with brine (1 x 20 mL). The organic layer was dried by $Na_2SO_4$, concentrated, and the resulting residue purified by flash chromatography to provide ZA1 as colorless oil (18.3 mg, 0.018 mmol, 44%).

$^1$H NMR (500 MHz, $CDCl_3$) δ 7.36 – 7.31 (m, 1H), 7.31 – 7.26 (m, 2H), 7.25 (d, J = 8.0 Hz, 2H), 7.20 (d, J = 7.2 Hz, 1H), 7.18 – 7.13 (m, 3H), 7.07 (m, 1H), 6.98 (d, J = 7.7 Hz, 1H), 6.94 – 6.87 (m, 2H), 6.84 (dd, J = 8.1, 2.6 Hz, 1H), 6.78 (dd, J = 8.4, 2.8 Hz, 1H), 6.71 – 6.64 (m, 2H), 5.78 – 5.71 (m, 1H), 5.28 (s, 1H), 4.7-4.6 (m, 1H), 4.49 (s, 2H), 3.97 – 3.74 (m, 10H), 3.61-3.48 (m, 10H), 3.43 – 3.32 (m, 3H), 3.21 – 3.07 (m, 2H), 3.03-2.99 (m, 1H), 2.66 – 2.47 (m, 2H), 2.42 (s, 1H), 2.39-2.33 (m, 1H), 2.09-2.0 (m, 1H), 1.97-1.91 (m, 3H), 1.83 – 1.57 (m, 5H), 1.53 – 1.30 (m, 2H), 1.22 – 1.09 (m, 6H), 0.89-0.78 (m, 3H).

$^{13}$C NMR (126 MHz, $CDCl_3$) δ 208.14, 171.44, 171.25, 169.88, 169.84, 169.01, 168.67, 167.51, 157.56, 149.06, 147.55, 141.99, 136.53, 133.45, 130.24, 129.35, 128.85, 127.25, 120.31, 120.15, 114.11, 113.77, 111.90, 111.50, 76.74, 70.38, 69.79, 69.67, 67.55, 56.08, 56.01, 55.18, 53.57, 51.46, 46.85, 44.31, 43.71, 43.20, 39.48, 39.07, 38.31, 37.79, 32.69, 32.60, 31.40, 29.84, 26.58, 25.06, 23.54, 23.36, 22.88, 21.31, 8.89.

HRMS (ESI) m/z calcd for $C_{53}H_{72}N_6NaO_{14}$ for $[M+Na]^+$: 1039.5004, found 1039.5003.

**ZA6**: *(R)-1-(3-(((S)-11-benzyl-2,7,10,13,16-pentaoxo-3,6,9,12,15-pentaazaheptadecyl)oxy)phenyl)-3-(3,4-dimethoxyphenyl)propyl (S)-1-(3,3-dimethyl-2-oxopentanoyl)piperidine-2-carboxylate*

The synthetic route of ZA6 was similar to ZA1.

$^1$H NMR (500 MHz, $CDCl_3$) δ 7.65 – 7.57 (m, 1H), 7.32 – 7.23 (m, 3H), 7.24 – 7.19 (m, 1H), 7.18 – 7.12 (m, 3H), 7.10 – 7.02 (m, 2H), 6.98 – 6.74 (m, 5H), 6.70 – 6.63 (m, 2H), 5.72 – 5.64 (m, 1H), 5.20 (d, J = 5.7 Hz, 1H), 4.55 – 4.37 (m, 3H), 3.86 – 3.78 (m, 8H), 3.67 – 3.57 (m, 2H), 3.57 – 3.46 (m, 1H), 3.46 – 3.35 (m, 3H), 3.34 – 3.27 (m, 1H), 3.20 – 3.08 (m, 2H), 3.09 – 2.99 (m, 1H), 2.64 – 2.49 (m, 2H), 2.38 (d, J = 13.8 Hz, 1H), 2.31 – 2.18 (m, 1H), 2.11 – 2.01 (m, 1H), 1.87 – 1.77 (m, 4H), 1.75 – 1.63 (m, 4H), 1.53 – 1.29 (m, 2H), 1.23 – 1.11 (m, 6H), 0.90 – 0.78 (m, 3H).

$^{13}$C NMR (126 MHz, $CDCl_3$) δ 208.44, 172.01, 171.32, 170.70, 169.82, 169.74, 169.24, 167.69, 157.76, 149.06, 147.57, 141.78, 136.49, 133.34, 130.23, 129.24, 128.83, 127.20, 120.28, 119.70, 114.09, 111.87, 111.52, 77.00, 67.66, 56.88, 56.06, 56.00, 55.36, 53.55, 51.56, 46.82, 44.30, 44.06, 43.15, 39.72, 39.09, 38.73, 38.16, 36.85, 32.47, 31.42, 26.55, 24.96, 23.47, 23.34, 22.66, 21.27, 8.83.

HRMS (ESI) m/z calcd for $C_{49}H_{64}N_6NaO_{12}$ for $[M+Na]^+$: 951.4480, found 951.4474.

**ZA8**: *(R)-1-(3-((14-amino-2-oxo-6,9,12-trioxa-3-azatetradecyl)oxy)phenyl)-3-(3,4-dimethoxyphenyl)propyl (S)-1-(3,3-dimethyl-2-oxopentanoyl)piperidine-2-carboxylate*

ZA8: SLF-COOH (104.6 mg, 0.179 mmol, 1.0 eq), HATU (115.1 mg, 0.269 mmol, 1.5 eq), and tert-butyl (2-(2-(2-(2-aminoethoxy)ethoxy)ethoxy)ethyl)carbamate (78.6 mg, 0.269 mmol, 1.5 eq) were dissolved in anhydrous DMF (1.8 mL) at room temperature. DIPEA (93.2 µL, 0.538 mmol, 3.0 eq) was added to the reaction and stirred for three hours at room temperature. Upon completion, the reaction was concentrated, extracted with EtOAc, washed with brine, and then washed with water. The organic layer was dried over $Na_2SO_4$ and concentrated. The resulting residue was used without further purification. TFA (0.5 mL) was added to a stirred solution of the above residue in DCM (1 mL). The reaction was stirred for two hours at room temperature. Upon completion, the reaction was concentrated. The resulting residue was purified by flash chromatography (7N $NH_3$ in MeOH / DCM) to provide ZA8 as a white solid (114.8 mg, 0.151 mmol, 84%).

$^1$H NMR (500 MHz, CDCl3) δ 7.32 – 7.26 (m, 1H), 7.21 – 7.13 (m, 1H), 7.01 – 6.82 (m, 3H), 6.80 – 6.74 (m, 1H), 6.71 – 6.64 (m, 2H), 5.80 – 5.70 (m, 1H), 5.30 (d, J = 3.9 Hz, 1H), 4.49 (s, 2H), 3.84 (d, J = 5.8 Hz, 6H), 3.65 – 3.57 (m, 10H), 3.57 – 3.52 (m, 2H), 3.49 (t, J = 5.2 Hz, 2H), 3.36 (d, J = 11.2 Hz, 1H), 3.16 (td, J = 13.1, 3.1 Hz, 1H), 2.89 – 2.79 (m, 2H), 2.64 – 2.47 (m, 2H), 2.36 (d, J = 12.2 Hz, 1H), 2.30 – 2.17 (m, 1H), 2.10 – 1.98 (m, 3H), 1.80 – 1.59 (m, 5H), 1.54 – 1.41 (m, 1H), 1.39 – 1.27 (m, 1H), 1.20 (d, J = 9.4 Hz, 5H), 1.11 (d, J = 4.9 Hz, 1H), 0.91 – 0.76 (m, 3H).

$^{13}$C NMR (126 MHz, CDCl3) δ 207.93, 169.77, 168.24, 167.38, 157.59, 149.01, 147.48, 141.93, 133.45, 130.10, 120.25, 120.16, 114.40, 113.38, 111.83, 111.42, 77.41, 77.16, 76.91, 76.61, 70.61, 70.39, 70.31, 69.88, 67.54, 56.03, 55.94, 51.36, 46.80, 44.26, 41.73, 38.97, 38.28, 32.58, 31.35, 26.53, 25.04, 23.58, 23.24, 21.31, 8.86.

HRMS (ESI) m/z calcd for $C_{40}H_{60}N_3O_{11}$ for $[M+H]^+$: 758.4228, found 758.4257.

**Synthesis of ZA3 and ZA5**

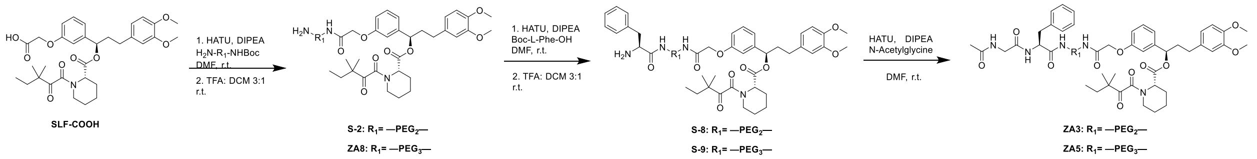

**ZA3:** *(R)-1-(3-(((S)-7-benzyl-2,5,8,19-tetraoxo-12,15-dioxa-3,6,9,18-tetraazaicosan-20-yl)oxy)phenyl)-3-(3,4-dimethoxyphenyl)propyl (S)-1-(3,3-dimethyl-2-oxopentanoyl)piperidine-2-carboxylate*

Step 1: S-2 (50 mg, 0.070 mmol, 1.0 eq), Boc-L-phenylalanine (22.3 mg, 0.084 mmol, 1.2 eq), and HATU (42.6 mg, 0.112 mmol, 1.6 eq) were dissolved in dry DMF (2.0 mL) at room temperature. DIPEA (37 µL, 0.210 mmol, 3.0 eq) was added to the reaction and stirred for 0.5 h. Upon completion, the reaction was concentrated, extracted with DCM (20 mL) and washed with brine (1 x 20 mL). The organic layer was dried by $Na_2SO_4$, concentrated, and the resulting residue was used without further purification. The crude product was dissolved in DCM (3.0 mL), TFA (1.0 mL) was added to the reaction and stirred for 0.5 h. The reaction mixture was then concentrated and purified by flash chromatography to provide S-8 as a colorless oil (51.7 mg, 0.060 mmol, 86% over 2 steps).

$^1$H NMR (500 MHz, $CDCl_3$) δ 7.96 (s, 1H), 7.29 – 7.15 (m, 8H), 6.98 – 6.91 (m, 2H), 6.84 – 6.73 (m, 2H), 6.72 – 6.63 (m, 2H), 5.78 – 5.71 (m, 1H), 5.29 (d, *J* = 4.1 Hz, 1H), 4.55 – 4.26 (m, 3H), 3.83 (d, *J* = 5.4 Hz, 6H), 3.61 – 3.09 (m, 17H), 2.63 – 2.47 (m, 2H), 2.35 (d, *J* = 13.3 Hz, 1H), 2.31 – 2.17 (m, 1H), 2.09 – 1.98 (m, 1H), 1.80 – 1.56 (m, 5H), 1.51 – 1.29 (m, 2H), 1.23 – 1.08 (m, 6H), 0.91 – 0.75 (m, 3H).

ZA3**:** S-8 (51.7 mg, 0.060 mmol, 1 eq), N-Acetylglycine (8.44 mg, 0.072 mmol, 1.2 eq), and HATU (36.5 mg, 0.096 mmol, 1.6 eq) were dissolved in dry DMF (2 mL) at room temperature. DIPEA (31 µL, 0.180 mmol, 3.0 eq) was added to the reaction and stirred for 0.5 h. Upon completion, the reaction was concentrated, extracted with DCM (20 mL) and washed with brine (1 x 20 mL). The organic layer was dried by $Na_2SO_4$, concentrated, and the resulting residue purified by flash chromatography to provide ZA3 as colorless oil (34.6 mg, 0.038 mmol, 65%).

$^1$H NMR (500 MHz, $CDCl_3$) δ 7.32 – 7.23 (m, 3H), 7.23 – 7.12 (m, 4H), 7.01 – 6.87 (m, 4H), 6.86 – 6.82 (m, 1H), 6.80 – 6.75 (m, 1H), 6.72 – 6.48 (m, 4H), 5.79 – 5.72 (m, 1H), 5.30 (s, 1H), 4.68 – 4.59 (m, 1H), 4.48 (s, 2H), 3.94 – 3.74 (m, 8H), 3.62 – 3.43 (m, 9H), 3.41 – 3.32 (m, 4H), 3.22 – 3.13 (m, 1H), 3.09 – 3.02 (m, 2H), 2.66 – 2.48 (m, 2H), 2.37 (d, J = 13.8 Hz, 1H), 2.28 – 2.18 (m, 1H), 2.10 – 1.99 (m, 0H), 1.99 – 1.95 (m, 3H), 1.81 – 1.60 (m, 5H), 1.51 – 1.44 (m, 1H), 1.37 – 1.32 (m, 1H), 1.23 – 1.10 (m, 6H), 0.91 – 0.77 (m, 3H).

$^{13}C$ NMR (126 MHz, $CDCl_3$) δ 208.04, 170.96, 170.72, 169.84, 168.99, 168.58, 167.47, 157.56, 149.03, 147.52, 141.99, 136.74, 133.46, 130.18, 129.46, 128.70, 127.08, 120.29, 120.17, 114.20, 113.64, 111.88, 111.48, 76.65, 70.42, 70.29, 69.76, 69.63, 67.56, 56.06, 55.98, 54.71, 51.43, 46.83, 44.29, 43.37, 39.44, 39.02, 38.44, 38.31, 32.59, 31.38, 26.56, 25.05, 23.56, 23.31, 22.96, 21.31, 18.70, 17.39, 8.88.

HRMS (ESI) m/z calcd for $C_{51}H_{69}N_5NaO_{13}$ for $[M+Na]^+$: 982.4790, found 982.4782.

**ZA5:** *(R)-1-(3-(((S)-7-benzyl-2,5,8,22-tetraoxo-12,15,18-trioxa-3,6,9,21-tetraazatricosan-23-yl)oxy)phenyl)-3-(3,4-dimethoxyphenyl)propyl (S)-1-(3,3-dimethyl-2-oxopentanoyl)piperidine-2-carboxylate*

The synthetic route of ZA5 was similar to ZA3.

$^1H$ NMR (500 MHz, $CDCl_3$) δ 7.32 – 7.24 (m, 3H), 7.24 – 7.15 (m, 4H), 7.00 – 6.97 (m, 1H), 6.95 – 6.87 (m, 2H), 6.87 – 6.82 (m, 1H), 6.81-6.74 (m, 2H), 6.72 – 6.64 (m, 2H), 6.58-6.51 (m, 1H), 5.79-5.72 (m, 1H), 5.31 (s, 1H), 4.69-4.61 (m, 1H), 4.49 (s, 2H), 3.94 – 3.75 (m, 8H), 3.63 – 3.43 (m, 13H), 3.40-3.30 (m, 4H), 3.22-3.14 (m, 1H), 3.12 – 3.00 (m, 2H), 2.67-2.48 (m, 2H), 2.40-2.33 (m, 1H), 2.30 – 2.18 (m, 1H), 2.11 – 2.01 (m, 1H),1.97 (m, 3H), 1.81 – 1.60 (m, 5H), 1.55 – 1.46 (m, 1H), 1.40-1.32 (m, 1H), 1.23 – 1.10 (m, 6H), 0.90-0.77 (m, 6H).

$^{13}C$ NMR (126 MHz, $CDCl_3$) δ 208.03, 171.18, 170.89, 169.85, 168.96, 168.62, 167.48, 157.59, 149.05, 147.53, 142.00, 136.70, 133.48, 130.17, 129.49, 128.70, 127.08, 120.30, 120.17, 114.29, 113.59, 111.89, 111.49, 76.67, 70.61, 70.57, 70.34, 70.27, 69.77, 69.67, 67.53, 56.07, 55.99, 54.65, 53.56, 51.44, 46.84, 44.30, 43.37, 39.52, 38.98, 38.50, 38.33, 36.69, 32.60, 31.40, 26.57, 25.06, 23.58, 23.31, 22.92, 21.33, 8.89.

HRMS (ESI) m/z calcd for $C_{53}H_{73}N_5NaO_{14}$ for $[M+Na]^+$: 1026.5052, found 1026.5048.

**Synthesis of ZA4**

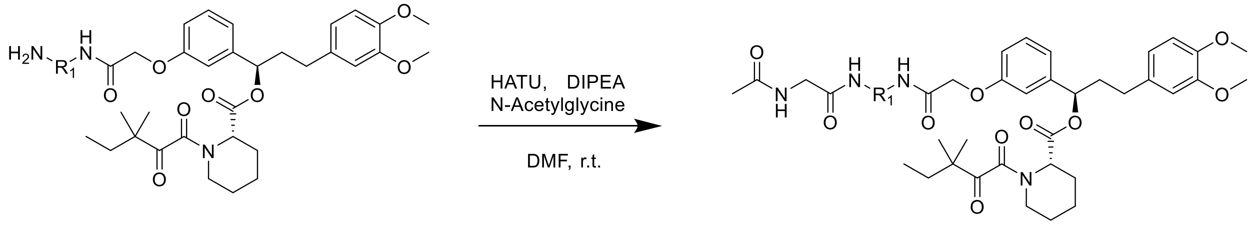


S-4: $R_1$= —$PEG_4$— ZA4: $R_1$= —$PEG_4$—

**ZA4**: *(R)-3-(3,4-dimethoxyphenyl)-1-(3-((2,5,22-trioxo-9,12,15,18-tetraoxa-3,6,21-triazatricosan-23-yl)oxy)phenyl)propyl (S)-1-(3,3-dimethyl-2-oxopentanoyl)piperidine-2-carboxylate*

ZA4: S-4 (60 mg, 0.075 mmol, 1 eq), N-Acetylglycine (10.51 mg, 0.080 mmol, 1.2 eq), and HATU (45.52 mg, 0.120 mmol, 1.6 eq) were dissolved in dry DMF (2 mL) at room temperature. DIPEA (39 μL, 0.224mmol, 3.0 eq) was added to the reaction and stirred for 0.5 h. Upon completion, the reaction was concentrated, extracted with DCM (20 mL) and washed with brine (1 x 20 mL). The organic layer was dried by $Na_2SO_4$, concentrated, and the resulting residue purified by flash chromatography to provide ZA4 as colorless oil (13.3 mg, 0.015 mmol, 20%).

$^1$H NMR (500 MHz, $CDCl_3$) δ 7.32 – 7.26 (m, 2H), 7.10 (t, J = 5.3 Hz, 1H), 7.01 – 6.83 (m, 3H), 6.81 – 6.75 (m, 1H), 6.72 – 6.64 (m, 2H), 6.56 – 6.50 (m, 1H), 5.81 – 5.68 (m, 1H), 5.30 (d, J = 5.6 Hz, 1H), 4.50 (s, 2H), 3.88 (dd, J = 26.6, 5.2 Hz, 8H), 3.65 – 3.50 (m, 18H), 3.44 (q, J = 5.2 Hz, 2H), 3.40 – 3.32 (m, 1H), 3.22 – 3.12 (m, 1H), 2.65 – 2.47 (m, 2H), 2.40 – 2.32 (m, 1H), 2.30 – 2.19 (m, 1H), 2.09 – 1.96 (m, 4H), 1.81 – 1.59 (m, 5H), 1.54 – 1.45 (m, 1H), 1.42 – 1.29 (m, 1H), 1.23 – 1.10 (m, 6H), 0.92 – 0.76 (m, 3H).

$^{13}$C NMR (126 MHz, $CDCl_3$) δ 207.98, 170.51, 169.81, 168.96, 168.47, 167.43, 157.67, 149.04, 147.51, 141.92, 133.50, 130.13, 120.28, 120.12, 114.44, 113.48, 111.87, 111.46, 76.66, 70.67, 70.61, 70.58, 70.56, 70.30, 70.22, 69.84, 69.82, 67.59, 56.06, 55.98, 51.41, 46.83, 44.29, 43.02, 39.46, 38.80, 38.31, 32.60, 31.39, 26.57, 25.07, 23.60, 23.27, 23.10, 21.34, 8.89.

HRMS (ESI) m/z calcd for $C_{46}H_{68}N_4NaO_{14}$ for $[M+Na]^+$: 923.4630, found 923.4626.

**Synthesis of ZA7**

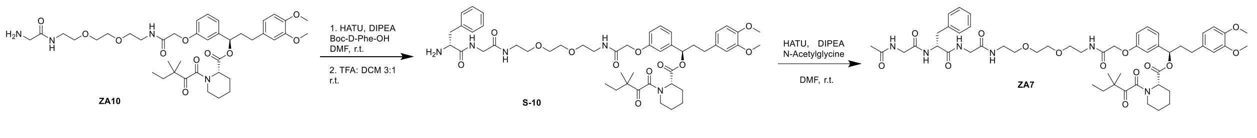


ZA7**:** *(R)-1-(3-(((R)-7-benzyl-2,5,8,11,22-pentaoxo-15,18-dioxa-3,6,9,12,21-pentaazatricosan-23-yl)oxy)phenyl)-3-(3,4-dimethoxyphenyl)propyl (S)-1-(3,3-dimethyl-2-oxopentanoyl)piperidine-2-carboxylate*

Step1: ZA10 (45 mg, 0.058 mmol, 1.0 eq), Boc-D-phenylalanine (18.6 mg, 0.075 mmol, 1.2 eq), and HATU (35.51 mg, 0.093 mmol, 1.6 eq) were dissolved in dry DMF (2.0 mL) at room temperature. DIPEA (30.5 µL, 0.175 mmol, 3.0 eq) was added to the reaction and stirred for 0.5 h. Upon completion, the reaction was concentrated, extracted with DCM (20 mL) and washed with brine (1 x 20 mL). The organic layer was dried by $Na_2SO_4$, concentrated, and the resulting residue was used without further purification. The crude product was dissolved in DCM (3.0 mL), TFA (1.0 mL) was added to the reaction and stirred for 0.5 h. The reaction mixture was then concentrated and purified by flash chromatography to provide S-10 as a colorless oil (44.1 mg, 0.048 mmol, 82% over 2 steps).

$^1$H NMR (500 MHz, CDCl3) δ 8.68 (s, 1H), 7.54 (s, 1H), 7.32 – 7.14 (m, 7H), 7.00 – 6.88 (m, 2H), 6.86 – 6.80 (m, 1H), 6.80 – 6.74 (m, 1H), 6.71 – 6.64 (m, 2H), 5.79 – 5.72 (m, 1H), 5.28 (s, 1H), 4.56 – 4.31 (m, 3H), 4.09 (s, 1H), 3.88 – 3.80 (m, 6H), 3.75 – 3.06 (m, 19H), 2.65 – 2.48 (m, 2H), 2.36 (d, J = 13.9 Hz, 1H), 2.30 – 2.15 (m, 1H), 2.11 – 1.97 (m, 1H), 1.83 – 1.58 (m, 5H), 1.47 – 1.34 (m, 2H), 1.23 – 1.09 (m, 6H), 0.93 – 0.75 (m, 3H).

ZA7: S-10 (44.1 mg, 0.048 mmol, 1 eq), N-Acetylglycine (6.75 mg, 0.080 mmol, 1.2 eq), and HATU (29.2 mg, 0.120 mmol, 1.6 eq) were dissolved in dry DMF (2 mL) at room temperature. DIPEA (25 µL, 0.144mmol, 3.0 eq) was added to the reaction and stirred for 0.5 h. Upon completion, the reaction was concentrated, extracted with DCM (20 mL) and washed with brine (1 x 20 mL). The organic layer was dried by $Na_2SO_4$, concentrated, and the resulting residue purified by flash chromatography to provide ZA7 as colorless oil (20.9 mg, 0.021 mmol, 42.8%).

$^1$H NMR (500 MHz, $CDCl_3$) δ 7.31 – 7.27 (m, 2H), 7.26 – 7.18 (m, 4H), 7.18 – 7.14 (m, 3H), 7.02-6.95 m, 2H), 6.94-6.91 (m, 1H), 6.86-6.80 (m Hz, 2H), 6.79-6.75 (m, 1H), 6.72 – 6.64 (m, 2H), 5.79-5.71 (m, 1H), 5.27 (s, 1H), 4.66 (q, J = 7.1 Hz, 1H), 4.49 (s, 2H), 3.95 – 3.74 (m, 10H), 3.61-3.49 (m, 10H), 3.43 – 3.31 (m, 3H), 3.21 – 3.07 (m, 2H), 3.06 – 2.97 (m, 1H), 2.67 – 2.47 (m, 2H), 2.29 – 2.18 (m, 1H), 2.11-1.99 (m, 2H), 1.94 (s, 3H), 1.82 – 1.60 (m, 5H), 1.55 – 1.30 (m, 2H), 1.23 – 1.10 (m, 6H), 0.90-0.77 (m, 3H).

$^{13}$C NMR (126 MHz, $CDCl_3$) δ 208.11, 171.40, 171.37, 169.85, 169.82, 169.06, 168.64, 167.49, 157.54, 149.03, 147.52, 141.96, 136.51, 133.45, 130.20, 129.31, 128.79, 127.20, 120.29, 120.13, 114.17, 113.69, 111.88, 111.48, 76.72, 70.37, 70.34, 69.78, 69.66, 67.51, 56.06, 55.99, 55.06, 53.56, 51.44, 46.83, 44.29, 43.60, 43.18, 39.45, 39.04, 38.29, 37.92, 32.58, 31.38, 31.05, 26.55, 25.04, 23.53, 23.33, 22.85, 21.30, 8.88.

HRMS (ESI) m/z calcd for $C_{53}H_{72}N_6NaO_{14}$ for $[M+Na]^+$: 1039.5004, found 1039.4999.

**Synthesis of ZA9 and ZA13-21**

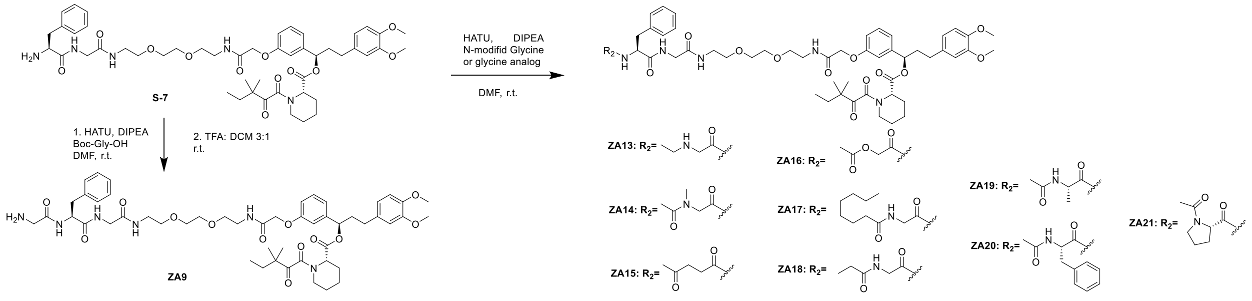

**ZA9:** *(R)-1-(3-(((S)-1-amino-4-benzyl-2,5,8,19-tetraoxo-12,15-dioxa-3,6,9,18-tetraazaicosan-20-yl)oxy)phenyl)-3-(3,4-dimethoxyphenyl)propyl (S)-1-(3,3-dimethyl-2-oxopentanoyl)piperidine-2-carboxylate*

ZA9: S-7 (65 mg, 0.071 mmol, 1.0 eq), Boc-Gly-OH (14.9 mg, 0.085 mmol, 1.2 eq), and HATU (43.07 mg, 0.113 mmol, 1.6 eq) were dissolved in dry DMF (2.0 mL) at room temperature. DIPEA (37 µL, 0.212 mmol, 3.0 eq) was added to the reaction and stirred for 0.5 h. Upon completion, the reaction was concentrated, extracted with DCM (20 mL) and washed with brine (1 x 20 mL). The organic layer was dried by $Na_2SO_4$, concentrated, and the resulting residue was used without further purification. The crude product was dissolved in DCM (3.0 mL), TFA (1.0 mL) was added to the reaction and stirred for 0.5 h. The reaction mixture was then concentrated and purified by flash chromatography to provide ZA9 as a colorless oil (32.6 mg, 0.033 mmol, 47% over 2 steps).

$^1$H NMR (500 MHz, $CDCl_3$) δ 7.83 (s, 1H), 7.32 – 7.26 (m, 3H), 7.24 – 7.18 (m, 4H), 7.15 – 7.10 (m, 1H), 7.00 – 6.88 (m, 2H), 6.88 – 6.75 (m, 3H), 6.72 – 6.65 (m, 2H), 5.79 – 5.72 (m, 1H), 5.28 (s, 1H), 4.61 – 4.47 (m, 3H), 3.96 – 3.75 (m, 8H), 3.63 – 3.49 (m, 10H), 3.43 – 3.25 (m, 5H), 3.23 – 3.13 (m, 2H), 3.08 – 2.99 (m, 1H), 2.65 – 2.48 (m, 2H), 2.37 (d, *J* = 13.7 Hz, 1H), 2.29 – 2.18 (m, 1H), 2.11 – 2.00 (m, 1H), 1.81 – 1.60 (m, 5H), 1.55 – 1.42 (m, 1H), 1.41 – 1.32 (m, 1H), 1.23 – 1.10 (m, 6H), 0.91 – 0.77 (m, 3H).

$^{13}$C NMR (126 MHz, $CDCl_3$) δ 208.08, 171.55, 169.85, 169.07, 168.67, 167.49, 157.60, 149.05, 147.54, 141.99, 136.72, 133.48, 130.19, 129.32, 128.84, 127.20, 120.31, 120.20, 114.17, 113.66, 111.89, 111.49, 76.70, 70.39, 70.33, 69.88, 69.64, 67.55, 56.84, 56.08, 56.01, 55.15, 51.44, 46.85, 44.31, 43.15, 39.38, 39.09, 38.32, 37.67, 32.69, 32.61, 31.40, 27.82, 26.58, 25.07, 23.72, 23.58, 23.33, 21.32, 8.90.

HRMS (ESI) m/z calcd for $C_{51}H_{71}N_6O_{13}$ for $[M+H]^+$: 975.5079, found 975.5079.

**ZA13:** *(R)-1-(3-(((S)-7-benzyl-5,8,11,22-tetraoxo-15,18-dioxa-3,6,9,12,21-pentaazatricosan-23-yl)oxy)phenyl)-3-(3,4-dimethoxyphenyl)propyl (S)-1-(3,3-dimethyl-2-oxopentanoyl)piperidine-2-carboxylate*

ZA13: S-7 (34.8 mg, 0.038 mmol, 1.0 eq), HATU (23.1 mg, 0.061 mmol, 1.6 eq) and N-(tert-butoxycarbonyl)-N-ethylglycine (9.24 mg, 0.045 mmol, 1.2 eq) were dissolved in anhydrous DMF 0.4 mL at room temperature. DIPEA (19.9 µL, 0.11 mmol, 3.0 eq) was added to the reaction and stirred for one hour at room temperature. Upon completion, the reaction was concentrated, extracted with DCM, washed with brine, and then washed with water. The organic layer was dried over $Na_2SO_4$ and concentrated. The crude residue was dissolved in 30% TFA in DCM (1 mL) and stirred at room temperature for two hours at room temperature. The reaction was concentrated, and the resulting residue was purified by flash chromatography (MeOH/DCM) to provide ZA13 as a white solid (27.6 mg, 0.28 mmol, 73%).

$^1$H NMR (500 MHz, $CDCl_3$) δ 8.97 (s, 1H), 8.37 (s, 1H), 7.44 (s, 1H), 7.32 – 7.26 (m, 1H), 7.24 – 7.07 (m, 5H), 7.00 – 6.87 (m, 2H), 6.84 (dd, J = 8.1, 2.5 Hz, 1H), 6.80 – 6.74 (m, 1H), 6.71 – 6.64 (m, 2H), 5.81 – 5.69 (m, 1H), 5.29 (d, J = 5.4 Hz, 1H), 4.76 (d, J = 7.7 Hz, 1H), 4.49 (s, 2H), 3.93 (d, J = 15.2 Hz, 2H), 3.84 (d, J = 5.3 Hz, 6H), 3.79 – 3.65 (m, 3H), 3.60 – 3.45 (m, 9H), 3.41 – 3.26 (m, 3H), 3.23 – 3.05 (m, 3H), 3.01 – 2.86 (m, 3H), 2.64 – 2.49 (m, 2H), 2.36 (d, J = 13.5 Hz, 1H), 2.27 – 2.18 (m, 1H), 2.09 – 1.99 (m, 1H), 1.83 – 1.57 (m, 5H), 1.55 – 1.42 (m, 1H), 1.38 – 1.31 (m, 2H), 1.26 – 1.17 (m, 8H), 1.12 (d, J = 2.4 Hz, 1H), 0.91 – 0.76 (m, 3H).

$^{13}$C NMR (126 MHz, $CDCl_3$) δ 207.96, 171.87, 169.70, 168.82, 167.33, 167.13, 157.36, 148.91, 147.41, 141.89, 136.48, 133.37, 130.06, 129.22, 128.52, 126.94, 120.18, 120.07, 114.06, 113.44, 111.76, 111.36, 76.54, 70.15, 70.10, 69.64, 69.27, 67.24, 55.93, 55.85, 55.58, 52.93, 51.30, 46.71, 44.18, 43.13, 43.07, 42.76, 39.25, 38.89, 38.18, 32.46, 31.25, 26.42, 24.93, 23.43, 23.19, 21.17, 11.12, 10.98, 8.75.

HRMS (ESI) m/z calcd for $C_{53}H_{75}N_6O_{13}$ for $[M+H]^+$: 1003.5392, found 1003.5390.

**ZA14:** *(S)-1-(3-(((S)-7-benzyl-3-methyl-2,5,8,11,22-pentaoxo-15,18-dioxa-3,6,9,12,21-pentaazatricosan-23-yl)oxy)phenyl)-3-(3,4-dimethoxyphenyl)propyl (R)-1-(3,3-dimethyl-2-oxopentanoyl)piperidine-2-carboxylate*

ZA14: S-7 (37.5 mg, 0.041 mmol, 1.0 eq), HATU (24.9 mg, 0.065 mmol, 1.6 eq) and *N*-acetyl-*N*-methylglycine (6.4 mg, 0.049 mmol, 1.2 eq) were dissolved in anhydrous DMF (0.4 mL) at room temperature. DIPEA (21.4 μL, 0.12 mmol, 3.0 eq) was added to the reaction and stirred overnight at room temperature. Upon completion, the reaction was concentrated, extracted with DCM, washed with brine, and then washed with water. The organic layer was dried over $Na_2SO_4$ and concentrated. The resulting residue was purified by flash chromatography (MeOH/DCM) to provide ZA14 as a white solid (16.8 mg, 0.016 mmol, 40%).

$^1$H NMR (500 MHz, $CDCl_3$) δ 7.32 – 7.26 (m, 3H), 7.25 – 7.15 (m, 4H), 7.14 – 7.06 (m, 1H), 7.00 – 6.94 (m, 1H), 6.94 – 6.88 (m, 1H), 6.85 (dd, *J* = 8.2, 2.6 Hz, 1H), 6.78 (dd, *J* = 8.3, 3.2 Hz, 2H), 6.74 – 6.65 (m, 3H), 5.76 (dd, *J* = 8.1, 5.6 Hz, 1H), 5.29 (d, *J* = 4.5 Hz, 1H), 4.61 (td, *J* = 8.1, 5.7 Hz, 1H), 4.49 (s, 2H), 3.92 – 3.77 (m, 10H), 3.63 – 3.48 (m, 10H), 3.47 – 3.33 (m, 3H), 3.23 – 3.12 (m, 2H), 3.06 – 2.98 (m, 1H), 2.95 (s, 2H), 2.64 – 2.48 (m, 2H), 2.36 (d, *J* = 13.7 Hz, 1H), 2.28 – 2.19 (m, 1H), 2.09 – 1.98 (m, 5H), 1.81 – 1.61 (m, 5H), 1.55 – 1.42 (m, 1H), 1.41 – 1.31 (m, 1H), 1.21 (d, *J* = 7.7 Hz, 5H), 1.12 (d, *J* = 3.2 Hz, 1H), 0.91 – 0.76 (m, 3H).

$^{13}$C NMR (126 MHz, $CDCl_3$) δ 208.02, 172.22, 171.29, 169.81, 169.72, 168.99, 168.49, 167.44, 157.56, 149.02, 147.50, 141.96, 136.70, 133.47, 130.18, 129.27, 128.84, 127.18, 120.29, 120.17, 114.25, 113.60, 111.87, 111.47,76.66, 70.50, 70.33, 69.79, 69.66, 67.54, 56.05, 55.98, 54.73, 52.68, 51.41, 46.83, 44.29, 43.25, 39.40, 39.04, 38.29, 38.00, 37.35, 32.59, 31.37, 26.55, 25.05, 23.57, 23.29, 21.47, 21.31, 8.88.

HRMS (ESI) m/z calcd for $C_{54}H_{74}N_6NaO_{14}$ for $[M+Na]^+$: 1053.5161, found 1053.5160.

**ZA15**: *(S)-1-(3-(((S)-17-benzyl-2,13,16,19,22-pentaoxo-6,9-dioxa-3,12,15,18-tetraazatricosyl)oxy)phenyl)-3-(3,4-dimethoxyphenyl)propyl (R)-1-(3,3-dimethyl-2-oxopentanoyl)piperidine-2-carboxylate*

ZA15: S-7 (36.5 mg, 0.040 mmol, 1.0 eq), HATU (24.2 mg, 0.064 mmol, 1.6 eq) and 4-oxopentanoic acid (5.5 mg, 0.048 mmol, 1.2 eq) were dissolved in anhydrous DMF (0.4 mL) at room temperature. DIPEA (20.8 µL, 0.12 mmol, 3.0 eq) was added to the reaction and stirred overnight at room temperature. Upon completion, the reaction was concentrated, extracted with DCM, washed with brine, and then washed with water. The organic layer was dried over $Na_2SO_4$ and concentrated. The resulting residue was purified by flash chromatography (MeOH/DCM) to provide ZA15 as a white solid (30.5 mg, 0.030 mmol, 75%).

$^1$H NMR (500 MHz, $CDCl_3$) δ 7.31 – 7.26 (m, 2H), 7.24 – 7.15 (m, 4H), 7.15 – 7.07 (m, 1H), 7.01 – 6.93 (m, 1H), 6.93 – 6.86 (m, 1H), 6.84 (dd, *J* = 8.1, 2.6 Hz, 1H), 6.80 – 6.70 (m, 2H), 6.70 – 6.63 (m, 2H), 6.55 (d, *J* = 7.1 Hz, 1H), 5.79 – 5.71 (m, 1H), 5.29 (d, *J* = 4.3 Hz, 1H), 4.64 – 4.56 (m, 1H), 4.48 (s, 2H), 3.95 – 3.78 (m, 8H), 3.62 – 3.47 (m, 10H), 3.46 – 3.31 (m, 3H), 3.21 – 3.07 (m, 2H), 3.07 – 2.98 (m, 1H), 2.89 – 2.48 (m, 5H), 2.39 – 2.12 (m, 4H), 2.10 (s, 3H), 2.08 – 2.00 (m, 1H), 1.80 – 1.60 (m, 5H), 1.54 – 1.41 (m, 1H), 1.40 – 1.30 (m, 1H), 1.20 (d, *J* = 7.6 Hz, 5H), 1.11 (d, *J* = 2.9 Hz, 1H), 0.90 – 0.76 (m, 3H).

$^{13}$C NMR (126 MHz, $CDCl_3$) δ 208.54, 208.00, 172.74, 171.59, 169.78, 169.07, 168.47, 167.41, 157.53, 148.99, 147.47, 141.93, 136.71, 133.45, 130.15, 129.23, 128.81, 127.13, 120.26, 120.14, 114.23, 113.58, 111.85, 111.45, 76.63, 70.46, 70.31, 69.76, 69.64, 67.50, 56.02, 55.95, 54.99, 51.38, 46.79, 44.26, 43.26, 39.37, 39.01, 38.59, 38.26, 37.42, 32.56, 31.34, 29.85, 29.83, 26.53, 25.02, 23.54, 23.26, 21.28, 8.85.

HRMS (ESI) m/z calcd for $C_{54}H_{73}N_5NaO_{14}$ for $[M+Na]^+$: 1038.5052, found 1038.5052.

**ZA16:** *(S)-1-(3-(((S)-17-benzyl-2,13,16,19,22-pentaoxo-6,9,21-trioxa-3,12,15,18-tetraazatricosyl)oxy)phenyl)-3-(3,4-dimethoxyphenyl)propyl (R)-1-(3,3-dimethyl-2-oxopentanoyl)piperidine-2-carboxylate*

The synthetic route of ZA16 was similar to ZA15.

$^1$H NMR (500 MHz, $CDCl_3$) δ 7.32 – 7.26 (m, 3H), 7.25 – 7.20 (m, 1H), 7.20 – 7.15 (m, 2H), 7.11 (t, *J* = 5.8 Hz, 1H), 7.06 – 6.95 (m, 2H), 6.94 – 6.87 (m, 1H), 6.87 – 6.81 (m, 2H), 6.80 – 6.74 (m, 1H), 6.71 – 6.64 (m, 3H), 5.80 – 5.72 (m, 1H), 5.29 (d, *J* = 4.8 Hz, 1H), 4.73 – 4.64 (m, 1H), 4.54 – 4.44 (m, 4H), 3.98 – 3.88 (m, 1H), 3.84 (d, *J* = 5.5 Hz, 6H), 3.81 – 3.74 (m, 1H), 3.62 – 3.57 (m, 4H), 3.56 – 3.52 (m, 3H), 3.50 (t, *J* = 5.3 Hz, 2H), 3.45 – 3.33 (m, 3H), 3.22 – 3.03 (m, 3H), 2.66 – 2.47 (m, 2H), 2.36 (d, *J* = 13.8 Hz, 1H), 2.29 – 2.17 (m, 1H), 2.10 – 2.00 (m, 5H), 1.81 – 1.60 (m, 5H), 1.55 – 1.42 (m, 1H), 1.41 – 1.30 (m, 1H), 1.20 (d, *J* = 7.0 Hz, 5H), 1.12 (d, *J* = 2.6 Hz, 1H), 0.83 (dt, *J* = 37.0, 7.4 Hz, 3H).

$^{13}$C NMR (126 MHz, $CDCl_3$) δ 208.01, 170.96, 169.79, 169.69, 168.63, 168.56, 167.46, 167.44, 157.51, 149.01, 147.49, 141.97, 136.18, 133.45, 130.17, 129.33, 128.86, 127.34, 120.27, 120.18, 114.08, 113.70, 111.86, 111.46,76.62, 70.54, 70.29, 69.76, 69.58, 67.50, 62.94, 56.04, 55.96, 54.35, 51.40, 46.81, 44.27, 43.11, 39.44, 39.05, 38.28, 38.05, 32.57, 31.35, 26.53, 25.03, 23.54, 23.28, 21.28, 20.69, 8.86.

HRMS (ESI) m/z calcd for $C_{53}H_{71}N_5NaO_{15}$ for $[M+Na]^+$: 1040.4844, found 1040.4844.

**ZA17:** *(S)-1-(3-(((S)-17-benzyl-2,13,16,19,22-pentaoxo-6,9-dioxa-3,12,15,18,21-pentaazanonacosyl)oxy)phenyl)-3-(3,4-dimethoxyphenyl)propyl (R)-1-(3,3-dimethyl-2-oxopentanoyl)piperidine-2-carboxylate*

The synthetic route of ZA17 was similar to ZA14.

$^{1}$H NMR (500 MHz, $CDCl_3$) δ 7.40 – 7.23 (m, 5H), 7.22 – 7.10 (m, 4H), 7.09 – 7.04 (m, 1H), 7.00 – 6.88 (m, 2H), 6.84 (dd, *J* = 8.3, 2.3 Hz, 1H), 6.79 – 6.71 (m, 2H), 6.70 – 6.64 (m, 2H), 5.75 (dt, *J* = 8.1, 5.3 Hz, 1H), 5.29 (q, *J* = 2.2 Hz, 1H), 4.69 (q, *J* = 7.1 Hz, 1H), 4.48 (s, 2H), 3.99 – 3.85 (m, 3H), 3.85 – 3.78 (m, 7H), 3.61 – 3.49 (m, 9H), 3.44 – 3.32 (m, 3H), 3.16 (td, *J* = 13.2, 3.1 Hz, 1H), 3.13 – 2.95 (m, 2H), 2.66 – 2.47 (m, 2H), 2.36 (d, *J* = 13.9 Hz, 1H), 2.29 – 2.13 (m, 4H), 2.11 – 1.99 (m, 1H), 1.83 – 1.59 (m, 5H), 1.59 – 1.31 (m, 4H), 1.31 – 1.22 (m, 8H), 1.20 (d, *J* = 7.0 Hz, 5H), 1.12 (d, *J* = 2.5 Hz, 1H), 0.90 – 0.77 (m, 6H).

$^{13}$C NMR (126 MHz, $CDCl_3$) δ 207.93, 174.18, 171.18, 169.71, 169.48, 168.81, 168.42, 167.34, 157.43, 148.91, 147.40, 141.85, 136.42, 133.34, 130.07, 129.25, 128.63, 127.06, 120.17, 120.04, 114.13, 113.48, 111.76, 111.37, 76.56, 70.29, 70.23, 69.67, 69.54, 67.40, 55.94, 55.86, 54.84, 51.31, 46.71, 44.18, 43.33, 43.10, 39.31, 38.91, 38.18, 36.15, 32.47, 31.68, 31.26, 29.29, 29.02, 26.43, 25.55, 24.93, 23.44, 23.19, 22.61, 21.19, 14.08, 8.76.

HRMS (ESI) m/z calcd for $C_{59}H_84N_6NaO_{14}$ for $[M+Na]^+$: 1123.5943, found 1123.5943.

**ZA18**: *(R)-1-(3-(((S)-17-benzyl-2,13,16,19,22-pentaoxo-6,9-dioxa-3,12,15,18,21-pentaazatetracosyl)oxy)phenyl)-3-(3,4-dimethoxyphenyl)propyl (S)-1-(3,3-dimethyl-2-oxopentanoyl)piperidine-2-carboxylate*

The synthetic route of ZA18 was similar to ZA14.

$^{1}$H NMR (500 MHz, $CDCl_3$) δ 7.37-7.22 (m, 5H), 7.21-7.18 (m, 1H), 7.17 – 7.03 (m, 4H), 6.98 (d, J = 7.7 Hz, 1H), 6.92 (t, J = 2.1 Hz, 1H), 6.85-6.82 (m, 1H), 6.80-6.74 (m, 2H), 6.71 – 6.64 (m, 2H), 5.78-5.71 (m, 1H), 5.27 (s, 1H), 4.70-4.62 (m, 2H), 4.48 (s, 1H), 3.96 – 3.78 (m, 10H), 3.60 – 3.49 (m, 10H), 3.41-3.35 (m, 2H), 3.21 – 3.05 (m, 2H), 3.04-2.97 (m, 1H), 2.63 – 2.48 (m, 2H), 2.30 – 2.14 (m, 4H), 2.10 – 1.98 (m, 1H), 1.83 – 1.59 (m, 5H), 1.53-1.42 (m, 1H), 1.41-1.33 (m, 2H), 1.20 (d, J = 7.0 Hz, 5H), 1.12 – 1.05 (m, 4H), 0.89-0.77 (m, 3H).

$^{13}C$ NMR (126 MHz, $CDCl_3$) δ 208.04, 174.98, 171.33, 169.80, 169.77, 169.03, 168.52, 167.44, 157.52, 149.00, 147.49, 141.95, 136.49, 133.44, 130.16, 129.32, 128.74, 127.17, 120.27, 120.13, 114.25, 113.52, 111.86, 111.46, 76.66, 70.37, 70.33, 69.74, 69.61, 67.48, 56.03, 55.96, 55.01, 53.55, 51.40, 46.80, 44.27, 43.49, 43.20, 39.43, 39.00, 38.27, 38.08, 32.55, 31.35, 29.16, 26.53, 25.02, 23.53, 23.28, 21.28, 12.54, 9.69, 8.85.

HRMS (ESI) m/z calcd for $C_{54}H_{74}N_6NaO_{14}$ for $[M+Na]^+$: 1053.5161, found 1053.5157.

**ZA19**: *(S)-1-(3-(((4S,7S)-7-benzyl-4-methyl-2,5,8,11,22-pentaoxo-15,18-dioxa-3,6,9,12,21-pentaazatricosan-23-yl)oxy)phenyl)-3-(3,4-dimethoxyphenyl)propyl (R)-1-(3,3-dimethyl-2-oxopentanoyl)piperidine-2-carboxylate*

The synthetic route of ZA19 was similar to ZA14.

$^1H$ NMR (500 MHz, $CDCl_3$) δ 7.41 – 7.34 (m, 1H), 7.29 (h, 2H), 7.25 – 7.20 (m, 2H), 7.20 – 7.10 (m, 4H), 7.09 – 7.02 (m, 1H), 7.00 – 6.94 (m, 1H), 6.93 – 6.88 (m, 1H), 6.84 (dd, *J* = 8.3, 2.6 Hz, 1H), 6.80 – 6.74 (m, 1H), 6.73 – 6.62 (m, 3H), 5.78 – 5.71 (m, 1H), 5.28 (d, *J* = 5.1 Hz, 1H), 4.73 – 4.61 (m, 1H), 4.48 (s, 2H), 4.47 – 4.40 (m, 1H), 3.99 – 3.90 (m, 1H), 3.89 – 3.80 (m, 7H), 3.63 – 3.56 (m, 6H), 3.56 – 3.49 (m, 3H), 3.44 – 3.31 (m, 3H), 3.21 – 3.08 (m, 2H), 3.03 (dd, *J* = 13.9, 7.7 Hz, 1H), 2.66 – 2.47 (m, 2H), 2.36 (d, *J* = 13.8 Hz, 1H), 2.28 – 2.17 (m, 3H), 2.09 – 1.99 (m, 1H), 1.89 (s, 2H), 1.81 – 1.59 (m, 5H), 1.54 – 1.41 (m, 1H), 1.40 – 1.29 (m, 1H), 1.28 – 1.22 (m, 3H), 1.20 (d, *J* = 7.0 Hz, 5H), 1.11 (d, *J* = 2.4 Hz, 1H), 0.91 – 0.77 (m, 3H).

$^{13}C$ NMR (126 MHz, $CDCl_3$) δ 208.03, 172.89, 171.23, 170.82, 169.80, 169.00, 168.46, 167.44, 157.52, 149.00, 147.49, 141.93, 136.75, 133.42, 130.17, 129.37, 128.64, 127.02, 120.27, 120.12, 114.25, 113.55, 111.85, 111.46, 76.67, 70.40, 70.36, 69.79, 69.67, 67.50, 56.03, 55.95, 54.88, 51.40, 49.44, 46.80, 44.26, 43.09, 39.43, 38.98, 38.26, 38.03, 32.56, 31.35, 26.53, 25.02, 23.53, 23.28, 23.01, 21.28, 18.14, 8.86.

HRMS (ESI) m/z calcd for $C_{54}H_{74}N_6NaO_{14}$ for $[M+Na]^+$: 1053.5161, found 1053.5159.

**ZA20:** *(S)-1-(3-(((4S,7S)-4,7-dibenzyl-2,5,8,11,22-pentaoxo-15,18-dioxa-3,6,9,12,21-pentaazatricosan-23-yl)oxy)phenyl)-3-(3,4-dimethoxyphenyl)propyl (R)-1-(3,3-dimethyl-2-oxopentanoyl)piperidine-2-carboxylate*

The synthetic route of ZA20 was similar to ZA14.

$^1$H NMR (500 MHz, $CDCl_3$) δ 7.29 (t, *J* = 8.0 Hz, 1H), 7.24 – 7.14 (m, 7H), 7.13 – 7.09 (m, 3H), 7.09 – 7.03 (m, 2H), 7.01 – 6.87 (m, 4H), 6.83 (dd, *J* = 7.8, 2.2 Hz, 1H), 6.79 – 6.74 (m, 1H), 6.71 – 6.64 (m, 2H), 6.55 (d, *J* = 7.3 Hz, 1H), 5.75 (dd, *J* = 8.1, 5.5 Hz, 1H), 5.29 (d, *J* = 5.1 Hz, 1H), 4.72 – 4.58 (m, 2H), 4.48 (s, 2H), 3.93 – 3.75 (m, 7H), 3.63 – 3.47 (m, 9H), 3.46 – 3.30 (m, 3H), 3.22 – 3.04 (m, 2H), 3.04 – 2.78 (m, 3H), 2.69 – 2.47 (m, 2H), 2.41 – 2.30 (m, 1H), 2.28 – 2.18 (m, 1H), 2.15 (s, 1H), 2.11 – 1.99 (m, 1H), 1.91 – 1.83 (m, 3H), 1.83 – 1.59 (m, 5H), 1.54 – 1.28 (m, 2H), 1.20 (d, *J* = 7.2 Hz, 5H), 1.12 (d, *J* = 2.9 Hz, 1H), 0.91 – 0.74 (m, 3H).

$^{13}$C NMR (126 MHz, $CDCl_3$) δ 208.04, 171.47, 171.00, 170.73, 169.83, 168.95, 168.49, 167.42, 157.51, 149.01, 147.50, 141.96, 136.67, 136.51, 133.43, 130.18, 129.38, 129.31, 128.71, 128.65, 127.11, 127.01, 120.27, 120.13, 114.18, 113.65, 111.85, 111.46, 76.68, 70.46, 70.34, 69.80, 69.70, 67.50, 56.04, 55.96, 54.87, 54.76, 51.39, 46.81, 44.26, 43.16, 39.46, 38.98, 38.28, 38.06, 37.88, 32.57, 31.36, 26.53, 25.03, 23.54, 23.29, 23.02, 21.28, 8.87.

HRMS (ESI) m/z calcd for $C_{60}H_{78}N_6NaO_{14}$ for $[M+Na]^+$: 1129.5474, found 1129.5474.

**ZA21**: *(S)-1-(3-(((S)-1-((S)-1-acetylpyrrolidin-2-yl)-3-benzyl-1,4,7,18-tetraoxo-11,14-dioxa-2,5,8,17-tetraazanonadecan-19-yl)oxy)phenyl)-3-(3,4-dimethoxyphenyl)propyl (R)-1-(3,3-dimethyl-2-oxopentanoyl)piperidine-2-carboxylate*

The synthetic route of ZA21 was similar to ZA14.

$^{1}H$ NMR (500 MHz, $CDCl_3$) δ 7.31 – 7.24 (m, 4H), 7.23 – 7.07 (m, 5H), 6.99 – 6.89 (m, 2H), 6.89 – 6.75 (m, 3H), 6.69 – 6.64 (m, 2H), 5.75 (dd, *J* = 8.0, 5.6 Hz, 1H), 5.28 (d, *J* = 4.5 Hz, 1H), 4.56 – 4.45 (m, 3H), 4.34 (dd, *J* = 7.8, 3.6 Hz, 1H), 3.96 – 3.85 (m, 2H), 3.83 (d, *J* = 6.3 Hz, 6H), 3.62 – 3.49 (m, 10H), 3.46 – 3.31 (m, 5H), 3.28 – 3.01 (m, 3H), 2.64 – 2.47 (m, 2H), 2.35 (d, *J* = 13.9 Hz, 1H), 2.29 – 2.17 (m, 2H), 2.14 – 1.98 (m, 2H), 1.92 – 1.59 (m, 10H), 1.54 – 1.41 (m, 1H), 1.39 – 1.30 (m, 1H), 1.20 (d, *J* = 8.8 Hz, 5H), 1.10 (d, *J* = 4.1 Hz, 1H), 0.89 – 0.76 (m, 3H).

$^{13}C$ NMR (126 MHz, $CDCl_3$) δ 207.96, 172.22, 171.35, 171.31, 169.76, 169.20, 168.35, 167.40, 157.56, 148.98, 147.45, 141.88, 137.05, 133.45, 130.12, 129.25, 128.65, 127.00, 120.25, 120.15, 114.34, 113.45, 111.83, 111.43, 76.63, 70.42, 70.34, 69.77, 69.62, 67.50, 60.39, 56.02, 55.93, 54.78, 51.37, 48.34, 46.78, 44.25, 43.24, 39.35, 38.99, 38.24, 36.73, 32.56, 31.33, 28.19, 26.52, 25.02, 24.98, 23.56, 23.23, 22.43, 21.28, 8.85.

HRMS (ESI) m/z calcd for $C_{56}H_{76}N_6NaO_{14}$ for $[M+Na]^+$: 1079.5317, found 1079.5314.

**Synthesis of ZA22-27**

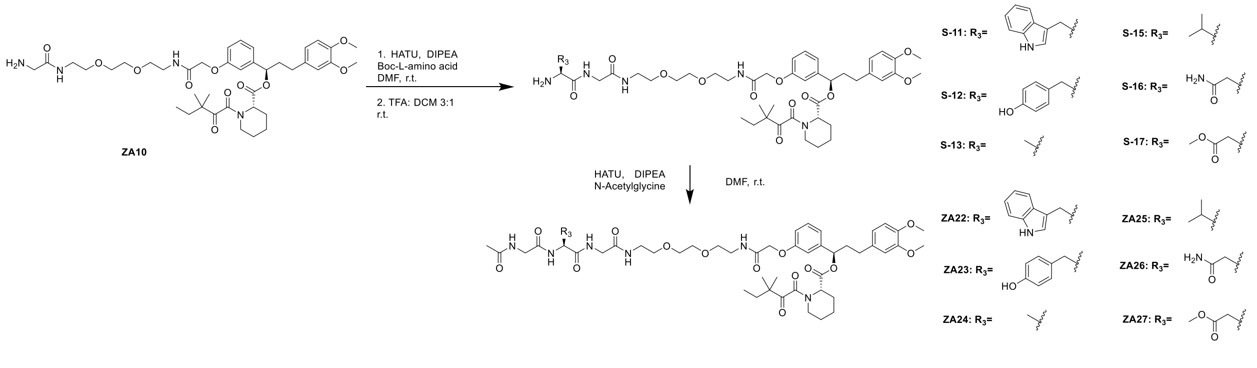


**ZA22**: *(R)-1-(3-(((S)-7-((1H-indol-3-yl)methyl)-2,5,8,11,22-pentaoxo-15,18-dioxa-3,6,9,12,21-pentaazatricosan-23-yl)oxy)phenyl)-3-(3,4-dimethoxyphenyl)propyl (S)-1-(3,3-dimethyl-2-oxopentanoyl)piperidine-2-carboxylate*

Step 1: ZA10 (62.3 mg, 0.081mmol, 1 eq), Boc-Trp-OH (29.5 mg, 0.097 mmol, 1.2 eq), and HATU (49.16 mg, 0.129 mmol, 1.6 eq) were dissolved in dry DMF (2.0 mL) at room temperature. DIPEA (42 μL, 0.242 mmol, 3.0 eq) was added to the reaction and stirred for 0.5 h. Upon completion, the reaction was concentrated, extracted with DCM (20 mL) and washed with brine (1 x 20 mL). The organic layer was dried by $Na_2SO_4$, concentrated, and the resulting residue was used without further purification. The crude product was dissolved in DCM (3.0 mL), TFA (1.0 mL) was added to the reaction and stirred for 0.5 h. The reaction mixture was then concentrated and purified by flash chromatography to provide S-11 as a colorless oil (52 mg, 0.051 mmol, 64% over 2 steps).

ZA22: S-11 (52 mg, 0.051 mmol, 1 eq), N-Acetylglycine (7.63 mg, 0.065 mmol, 1.2 eq), and HATU (33.05 mg, 0.87 mmol, 1.6 eq) were dissolved in dry DMF (1.5 mL) at room temperature. DIPEA (28 μL, 0.162 mmol, 3.0 eq) was added to the reaction and stirred for 0.5 h. Upon completion, the reaction was concentrated, extracted with DCM (20 mL) and washed with brine (1 x 20 mL). The organic layer was dried by $Na_2SO_4$, concentrated, and the resulting residue purified by flash chromatography to provide ZA22 as colorless oil (22.1 mg, 0.021 mmol, 38.5%).

$^1$H NMR (500 MHz, $CDCl_3$) δ 9.02 – 8.97 (m, 1H), 7.58 – 7.51 (m, 1H), 7.34 – 7.24 (m, 2H), 7.20 – 7.09 (m, 3H), 7.08 – 7.01 (m, 3H), 7.00 – 6.92 (m, 1H), 6.87 – 6.84 (m, 1H), 6.82 – 6.64 (m, 6H), 5.78 – 5.71 (m, 1H), 5.29 (s, 1H), 4.75 – 4.67 (m, 1H), 4.47 – 4.33 (m, 2H), 3.84 (d, J = 4.5 Hz, 6H), 3.81 – 3.75 (m, 4H), 3.63 – 3.45 (m, 10H), 3.39 – 3.12 (m, 6H), 2.64 – 2.47 (m, 2H), 2.37 (d, J = 13.7 Hz, 1H), 2.30 – 2.18 (m, 1H), 2.11 – 1.98 (m, 1H), 1.96 (s, 2H), 1.78 (s, 3H), 1.73 – 1.64 (m, 3H), 1.54 – 1.42 (m, 1H), 1.38 – 1.30 (m, 1H), 1.23 – 1.10 (m, 6H), 0.91 – 0.77 (m, 3H).

$^{13}$C NMR (126 MHz, $CDCl_3$) δ 174.85, 171.02, 170.93, 170.85, 169.32, 169.24, 161.62, 159.73, 151.55, 150.76, 142.01, 136.31, 136.14, 135.69, 129.44, 128.57, 127.02, 120.59, 118.20, 117.72, 105.37, 77.36, 58.68, 57.95, 57.91, 55.96, 54.55, 54.46, 52.82,

52.05, 48.71, 43.07, 39.59, 39.46, 38.54, 37.26, 37.15, 29.45, 29.32, 29.31, 29.22, 29.18, 29.14, 27.53, 26.93, 26.49, 24.90, 15.34, 11.19, 9.73.

HRMS (ESI) m/z calcd for $C_{55}H_{73}N_7NaO_{14}$ for $[M+Na]^+$: 1078.5113, found 1078.5106.

**ZA23**: *(R)-3-(3,4-dimethoxyphenyl)-1-(3-(((S)-7-(4-hydroxybenzyl)-2,5,8,11,22-pentaoxo-15,18-dioxa-3,6,9,12,21-pentaazatricosan-23-yl)oxy)phenyl)propyl (S)-1-(3,3-dimethyl-2-oxopentanoyl)piperidine-2-carboxylate*

The synthetic route of ZA23 was similar to ZA22.

$^1$H NMR (500 MHz, $CDCl_3$) δ 7.91 (s, 1H), 7.40 (s, 1H), 7.34 – 7.27 (m, 2H), 7.20 (t, J = 5.8 Hz, 1H), 7.09 – 7.01 (m, 1H), 7.00 – 6.89 (m, 4H), 6.89 – 6.80 (m, 2H), 6.80 – 6.75 (m, 1H), 6.73 – 6.64 (m, 4H), 5.75 (dd, J = 8.1, 5.3 Hz, 1H), 5.28 (s, 1H), 4.61 (q, J = 6.9 Hz, 1H), 4.47 (s, 2H), 3.88 – 3.77 (m, 10H), 3.61 – 3.46 (m, 10H), 3.41 – 3.31 (m, 3H), 3.22 – 3.13 (m, 1H), 2.64 – 2.48 (m, 2H), 2.36 (d, J = 13.6 Hz, 1H), 2.29 – 2.17 (m, 1H), 2.09 – 1.98 (m, 3H), 1.95 (s, 3H), 1.82 – 1.58 (m, 5H), 1.55 – 1.41 (m, 1H), 1.40 – 1.28 (m, 1H), 1.22 – 1.09 (m, 6H), 0.91 – 0.76 (m, 3H).

$^{13}$C NMR (126 MHz, $CDCl_3$) δ 208.10, 171.71, 171.64, 169.86, 169.70, 169.31, 168.80, 167.52, 157.51, 155.92, 149.02, 147.51, 141.93, 133.48, 130.45, 130.19, 127.38, 120.32, 120.12, 115.83, 114.26, 113.59, 111.91, 111.52, 76.75, 70.30, 70.28, 69.80, 69.66, 67.44, 56.85, 56.07, 56.00, 55.09, 51.47, 46.84, 44.32, 43.45, 43.18, 39.51, 39.06, 38.74, 38.29, 37.25, 32.57, 31.37, 26.56, 25.03, 23.54, 23.32, 22.85, 21.29, 8.88.

HRMS (ESI) m/z calcd for $C_{53}H_{72}N_6NaO_{15}$ for $[M+Na]^+$: 1055.4953, found 1055.4948.

**ZA24:** *(R)-3-(3,4-dimethoxyphenyl)-1-(3-(((S)-7-methyl-2,5,8,11,22-pentaoxo-15,18-dioxa-3,6,9,12,21-pentaazatricosan-23-yl)oxy)phenyl)propyl (S)-1-(3,3-dimethyl-2-oxopentanoyl)piperidine-2-carboxylate*

The synthetic route of ZA24 was similar to ZA22.

$^1$H NMR (500 MHz, $CDCl_3$) δ 7.46 – 7.38 (m, 2H), 7.33 – 7.27 (m, 1H), 7.17 – 7.10 (m, 2H), 7.02 – 6.96 (m, 2H), 6.94 – 6.91 (m, 1H), 6.87 – 6.82 (m, 1H), 6.81 – 6.75 (m, 1H), 6.72 – 6.64 (m, 2H), 5.79 – 5.72 (m, 1H), 5.29 (s, 1H), 4.57 – 4.47 (m, 3H), 3.98 – 3.92 (m, 4H), 3.87 – 3.82 (m, 6H), 3.62 – 3.50 (m, 10H), 3.47 – 3.32 (m, 3H), 3.23 – 3.13 (m, 1H), 2.66 – 2.48 (m, 2H), 2.29 – 2.18 (m, 1H), 2.11 – 1.98 (m, 4H), 1.96 (s, 1H), 1.83 – 1.59 (m, 5H), 1.55 – 1.42 (m, 1H), 1.42 – 1.30 (m, 4H), 1.23 – 1.10 (m, 6H), 0.92 – 0.76 (m, 3H).

$^{13}$C NMR (126 MHz, $CDCl_3$) δ 208.11, 172.67, 171.41, 169.86, 169.56, 169.15, 168.61, 167.51, 157.56, 149.04, 147.54, 141.99, 133.45, 130.22, 120.30, 120.16, 114.16, 113.68, 111.89, 111.49, 76.71, 70.37, 69.79, 69.68, 67.55, 56.83, 56.07, 56.00, 51.46, 49.63, 46.84, 44.31, 43.61, 43.21, 39.48, 39.04, 38.30, 32.59, 31.39, 26.57, 25.05, 23.54, 23.34, 22.94, 21.31, 18.25, 8.88.

HRMS (ESI) m/z calcd for $C_{47}H_{68}N_6NaO_{14}$ for $[M+Na]^+$: 963.4691, found 963.4689.

**ZA25**: *(R)-3-(3,4-dimethoxyphenyl)-1-(3-(((S)-7-isopropyl-2,5,8,11,22-pentaoxo-15,18-dioxa-3,6,9,12,21-pentaazatricosan-23-yl)oxy)phenyl)propyl (S)-1-(3,3-dimethyl-2-oxopentanoyl)piperidine-2-carboxylate*

The synthetic route of ZA25 was similar to ZA22.

$^1$H NMR (500 MHz, $CDCl_3$) δ 7.40 (s, 1H), 7.33 – 7.26 (m, 2H), 7.17 – 7.09 (m, 2H), 7.01 – 6.88 (m, 3H), 6.85 (dd, $J$ = 8.2, 2.5 Hz, 1H), 6.78 (dd, $J$ = 8.5, 3.8 Hz, 1H), 6.72 – 6.64 (m, 2H), 5.79 – 5.72 (m, 1H), 5.32 – 5.27 (m, 1H), 4.50 (s, 2H), 4.35 – 4.28 (m, 1H), 4.03 – 3.93 (m, 4H), 3.84 (d, $J$ = 5.0 Hz, 6H), 3.63 – 3.49 (m, 10H), 3.46 – 3.32 (m, 3H), 3.22 – 3.13 (m, 1H), 2.66 – 2.48 (m, 2H), 2.30 – 2.18 (m, 1H), 2.17 – 2.09 (m, 1H), 2.09 – 1.98 (m, 4H), 1.82 – 1.60 (m, 5H), 1.55 – 1.28 (m, 2H), 1.23 – 1.10 (m, 6H), 0.94 (t, $J$ = 7.2 Hz, 6H), 0.90 – 0.76 (m, 3H).

$^{13}C$ NMR (126 MHz, $CDCl_3$) δ 208.04, 171.53, 171.21, 169.83, 169.02, 168.57, 167.46, 157.54, 149.03, 147.52, 141.99, 133.46, 130.19, 120.29, 120.19, 114.23, 113.59, 111.88, 111.48, 76.66, 70.38, 70.34, 69.80, 69.69, 67.53, 59.14, 56.06, 55.99, 51.43, 46.83, 44.30, 43.63, 43.11, 39.44, 39.02, 38.30, 32.59, 31.37, 26.55, 25.05, 23.56, 23.31, 22.95, 21.31, 19.35, 18.25, 8.88.

HRMS (ESI) m/z calcd for $C_{49}H_{72}N_6NaO_{14}$ for $[M+Na]^+$: 991.5004, found 991.5003.

**ZA26**: *(R)-1-(3-(((S)-7-(2-amino-2-oxoethyl)-2,5,8,11,22-pentaoxo-15,18-dioxa-3,6,9,12,21-pentaazatricosan-23-yl)oxy)phenyl)-3-(3,4-dimethoxyphenyl)propyl (S)-1-(3,3-dimethyl-2-oxopentanoyl)piperidine-2-carboxylate*

The synthetic route of ZA26 was similar to ZA22.

$^{1}H$ NMR (500 MHz, $CDCl_3$) δ 8.06 (d, *J* = 63.4 Hz, 2H), 7.42 – 7.27 (m, 3H), 7.26 – 7.18 (m, 1H), 7.03 – 6.88 (m, 3H), 6.87 – 6.82 (m, 1H), 6.81 – 6.74 (m, 1H), 6.72 – 6.64 (m, 2H), 6.40 (s, 1H), 5.78 – 5.71 (m, 1H), 5.29 – 5.26 (m, 1H), 4.76 (s, 1H), 4.50 (s, 2H), 3.98 (d, *J* = 13.0 Hz, 1H), 3.88 – 3.77 (m, 9H), 3.60 – 3.49 (m, 10H), 3.38 – 3.33 (m, 3H), 3.23 – 3.14 (m, 1H), 2.95 – 2.89 (m, 1H), 2.67 (d, *J* = 13.9 Hz, 1H), 2.63 – 2.48 (m, 2H), 2.36 (d, *J* = 13.6 Hz, 1H), 2.28 – 2.18 (m, 1H), 2.10 – 1.99 (m, 1H), 1.98 (s, 3H), 1.82 – 1.60 (m, 5H), 1.45 – 1.39 (m, 1H), 1.38 – 1.31 (m, 1H), 1.22 – 1.10 (m, 6H), 0.91 – 0.76 (m, 3H).

$^{13}C$ NMR (126 MHz, $CDCl_3$) δ 208.13, 173.96, 172.30, 171.79, 170.20, 169.88, 168.84, 167.52, 157.57, 149.04, 147.52, 141.98, 133.48, 130.23, 120.32, 120.09, 114.28, 113.60, 111.91, 111.51, 76.76, 70.30, 70.21, 69.73, 69.63, 67.50, 56.08, 56.01, 53.71, 53.56, 51.47, 50.45, 46.84, 44.32, 43.82, 43.32, 39.48, 39.05, 38.30, 32.58, 31.39, 26.58, 25.04, 23.54, 23.34, 22.83, 21.31, 8.89.

HRMS (ESI) m/z calcd for $C_{48}H_{69}N_7NaO_{15}$ for $[M+Na]^+$: 1006.4749, found 1006.4743.

**ZA27**: *(R)-3-(3,4-dimethoxyphenyl)-1-(3-(((S)-7-(2-methoxy-2-oxoethyl)-2,5,8,11,22-pentaoxo-15,18-dioxa-3,6,9,12,21-pentaazatricosan-23-yl)oxy)phenyl)propyl (S)-1-(3,3-dimethyl-2-oxopentanoyl)piperidine-2-carboxylate*

The synthetic route of ZA27 was similar to ZA22.

$^{1}$H NMR (500 MHz, $CDCl_3$) δ 7.62 (t, *J* = 5.9 Hz, 1H), 7.55 (t, *J* = 8.3 Hz, 1H), 7.30 (dd, *J* = 9.2, 6.7 Hz, 1H), 7.20 (t, *J* = 5.9 Hz, 1H), 7.01 – 6.82 (m, 5H), 6.78 (dd, *J* = 8.4, 3.6 Hz, 1H), 6.72 – 6.65 (m, 2H), 5.75 (dd, *J* = 8.1, 5.5 Hz, 1H), 5.28 (d, *J* = 2.0 Hz, 1H), 4.79 (ddt, *J* = 13.8, 8.4, 5.4 Hz, 1H), 4.50 (s, 2H), 4.05 – 3.96 (m, 1H), 3.95 – 3.75 (m, 9H), 3.67 (s, 3H), 3.63 – 3.49 (m, 10H), 3.49 – 3.31 (m, 3H), 3.19 (td, *J* = 13.1, 3.1 Hz, 1H), 3.04 (ddd, *J* = 17.2, 5.2, 2.1 Hz, 1H), 2.78 (ddd, *J* = 17.1, 9.0, 5.6 Hz, 1H), 2.65 – 2.48 (m, 2H), 2.41 – 2.34 (m, 1H), 2.29 – 2.19 (m, 1H), 2.09 – 1.93 (m, 4H), 1.83 – 1.59 (m, 4H), 1.48 (qt, *J* = 12.8, 4.0 Hz, 1H), 1.37 (tt, *J* = 13.2, 3.7 Hz, 1H), 1.22 – 1.10 (m, 6H), 0.84 (dt, *J* = 35.0, 7.4 Hz, 3H).

$^{13}$C NMR (126 MHz, $CDCl_3$) δ 208.19, 172.48, 171.87, 170.73, 169.88, 169.20, 168.66, 167.50, 157.56, 149.03, 147.52, 141.96, 133.43, 130.23, 120.28, 120.07, 114.09, 113.76, 111.87, 111.48, 76.76, 70.38, 70.35, 69.76, 69.64, 67.53, 56.06, 55.98, 53.55, 52.27, 51.45, 49.78, 46.83, 44.29, 43.95, 43.39, 39.45, 39.04, 38.27, 35.42, 32.57, 31.38, 26.56, 25.04, 23.50, 23.35, 22.80, 21.28, 8.86.

HRMS (ESI) m/z calcd for $C_{49}H_{70}N_6NaO_{16}$ for $[M+Na]^+$: 1021.4746, found. 1021.4746

**Synthesis of ZA28-31**

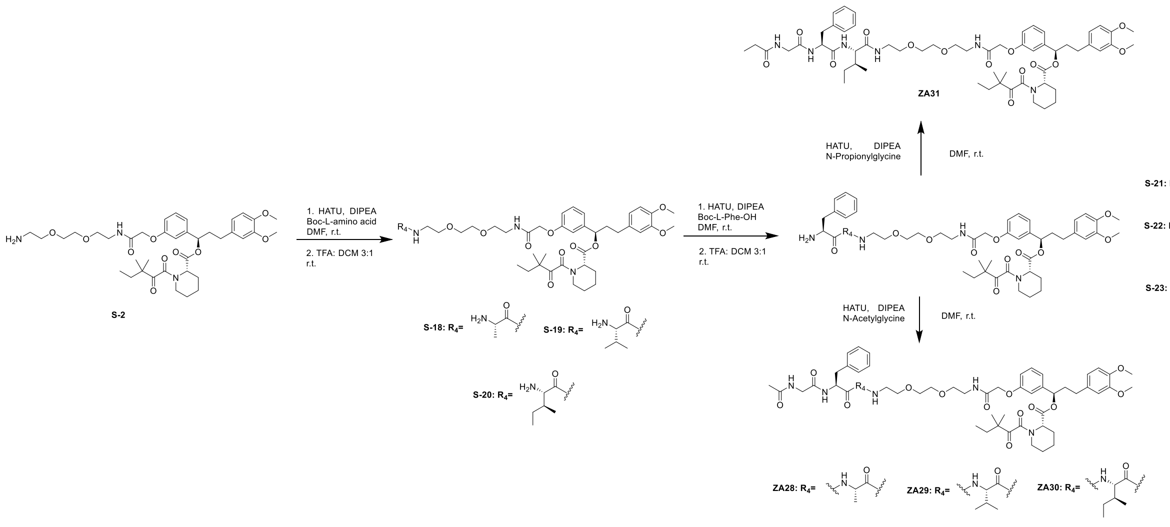

**ZA28**: *(R)-1-(3-(((7S,10S)-7-benzyl-10-methyl-2,5,8,11,22-pentaoxo-15,18-dioxa-3,6,9,12,21-pentaazatricosan-23-yl)oxy)phenyl)-3-(3,4-dimethoxyphenyl)propyl (S)-1-(3,3-dimethyl-2-oxopentanoyl)piperidine-2-carboxylate*

Step 1: S-2 (56.6 mg, 0.079mmol, 1 eq), Boc-Ala-OH (18 mg, 0.095 mmol, 1.2 eq), and HATU (48.2 mg, 0.126 mmol, 1.6 eq) were dissolved in dry DMF (2.0 mL) at room temperature. DIPEA (41 μL, 0.228 mmol, 3.0 eq) was added to the reaction and stirred for 0.5 h. Upon completion, the reaction was concentrated, extracted with DCM (20 mL) and washed with brine (1 x 20 mL). The organic layer was dried by $Na_2SO_4$, concentrated, and the resulting residue was used without further purification. The crude product was dissolved in DCM (3.0 mL), TFA (1.0 mL) was added to the reaction and stirred for 0.5 h. The reaction mixture was then concentrated and purified by flash chromatography to provide S-18 as a colorless oil (60 mg, 0.076 mmol, 96% over 2 steps).

$^1$H NMR (500 MHz, $CDCl_3$) δ 8.52 – 7.97 (m, 3H), 7.31 – 7.26 (m, 1H), 7.23 – 7.13 (m, 1H), 6.99 – 6.91 (m, 2H), 6.87 – 6.81 (m, 1H), 6.80 – 6.74 (m, 1H), 6.71 – 6.63 (m, 2H), 5.75 (dd, *J* = 8.1, 5.4 Hz, 1H), 5.29 (s, 1H), 4.49 (s, 2H), 4.15 (s, 1H), 3.84 (d, *J* = 5.9 Hz, 6H), 3.71 – 3.40 (m, 12H), 3.38 – 3.31 (m, 2H), 3.22 – 3.12 (m, 1H), 3.12 – 3.03 (m, 1H), 2.65 – 2.47 (m, 2H), 2.39 – 2.33 (m, 1H), 2.29 – 2.18 (m, 1H), 2.10 – 1.98 (m, 1H),

1.81 – 1.58 (m, 5H), 1.54 – 1.46 (m, 3H), 1.43 – 1.34 (m, 2H), 1.22 – 1.08 (m, 6H), 0.91 – 0.76 (m, 3H).

Step 2: S-18 (60 mg, 0.076 mmol, 1 eq), Boc-Phe-OH (24.3 mg, 0.092 mmol, 1.2 eq), and HATU (46.5 mg, 0.122 mmol, 1.6 eq) were dissolved in dry DMF (2.0 mL) at room temperature. DIPEA (40 µL, 0.228 mmol, 3.0 eq) was added to the reaction and stirred for 0.5 h. Upon completion, the reaction was concentrated, extracted with DCM (20 mL) and washed with brine (1 x 20 mL). The organic layer was dried by $Na_2SO_4$, concentrated, and the resulting residue was used without further purification. The crude product was dissolved in DCM (3.0 mL), TFA (1.0 mL) was added to the reaction and stirred for 0.5 h. The reaction mixture was then concentrated and purified by flash chromatography to provide S-21 as a colorless oil (50 mg, 0.054 mmol, 70% over 2 steps).

1H NMR (500 MHz, CDCl3) δ 7.47 (s, 1H), 7.32 – 7.13 (m, 9H), 6.99 – 6.94 (m, 1H), 6.94 – 6.86 (m, 1H), 6.85 – 6.74 (m, 2H), 6.71 – 6.64 (m, 2H), 5.78 – 5.72 (m, 1H), 5.31 (s, 1H), 4.55 – 4.26 (m, 4H), 3.83 (d, J = 4.5 Hz, 6H), 3.60 – 3.38 (m, 16H), 3.38 – 3.31 (m, 1H), 3.30 – 3.05 (m, 4H), 2.63 – 2.49 (m, 2H), 2.36 (d, J = 13.4 Hz, 1H), 2.29 – 2.18 (m, 1H), 2.12 – 1.98 (m, 1H), 1.81 – 1.58 (m, 5H), 1.53 – 1.41 (m, 1H), 1.40 – 1.29 (m, 1H), 1.28 – 1.09 (m, 9H), 0.90 – 0.76 (m, 3H).

ZA28: S-21 (50 mg, 0.054 mmol, 1 eq), N-Acetylglycine (7.57 mg, 0.065 mmol, 1.2 eq), and HATU (32.77 mg, 0.086 mmol, 1.6 eq) were dissolved in dry DMF (1.5 mL) at room temperature. DIPEA (28 µL, 0.161 mmol, 3.0 eq) was added to the reaction and stirred for 0.5 h. Upon completion, the reaction was concentrated, extracted with DCM (20 mL) and washed with brine (1 x 20 mL). The organic layer was dried by $Na_2SO_4$, concentrated, and the resulting residue purified by flash chromatography to provide ZA28 as colorless oil (22.4 mg, 0.022 mmol, 40.3%).

$^1$H NMR (500 MHz, $CDCl_3$) δ 7.29 (t, *J* = 7.9 Hz, 3H), 7.25-7.22 (m, 2H), 7.21-7.16 (m, 2H), 7.16 – 7.05 (m, 4H), 7.00 – 6.96 (m, 1H), 6.92 (t, *J* = 2.0 Hz, 1H), 6.87 – 6.82 (m,

1H), 6.79-6.75 (m, 1H), 6.71 – 6.64 (m, 2H), 5.78-5.72 (m, 1H), 5.30 (s, 1H), 4.73 (s, 1H), 4.58-4.39 (m, 3H), 3.96-3.78 (m, 8H), 3.63 – 3.49 (m, 10H), 3.43 – 3.32 (m, 3H), 3.17 (td, *J* = 13.4, 3.3 Hz, 1H), 3.02 (d, *J* = 6.5 Hz, 2H), 2.65-2.47 (m, 2H), 2.29 – 2.17 (m, 1H), 1.99-1.92 (m, 4H), 1.81 – 1.59 (m, 5H), 1.54-1.45 (m, 1H), 1.36-1.31 (m, 1H), 1.29 (d, *J* = 6.9 Hz, 3H), 1.23 – 1.07 (m, 6H), 0.89-0.77 (m, 3H).

$^{13}$C NMR (126 MHz, $CDCl_3$) δ 208.04, 172.42, 171.44, 170.49, 169.82, 168.49, 167.46, 157.55, 149.03, 147.51, 141.98, 136.28, 133.45, 130.18, 129.43, 128.73, 127.20, 120.28, 120.16, 114.37, 113.42, 111.87, 111.48, 76.68, 70.41, 70.38, 69.82, 69.71, 67.51, 56.05, 55.97, 54.77, 53.56, 51.43, 49.10, 46.82, 44.29, 43.71, 39.47, 38.99, 38.29, 32.57, 31.37, 26.56, 25.04, 23.53, 23.30, 22.82, 21.30, 18.73, 18.55, 17.37, 12.58, 8.88.

HRMS (ESI) m/z calcd for $C_{54}H_{74}N_6NaO_{14}$ for $[M+Na]^+$: 1053.5161, found: 1053.5159

**ZA29**: *(R)-1-(3-(((7S,10S)-7-benzyl-10-isopropyl-2,5,8,11,22-pentaoxo-15,18-dioxa-3,6,9,12,21-pentaazatricosan-23-yl)oxy)phenyl)-3-(3,4-dimethoxyphenyl)propyl (S)-1-(3,3-dimethyl-2-oxopentanoyl)piperidine-2-carboxylate*

The synthetic route of ZA29 was similar to ZA28.

$^{1}$H NMR (500 MHz, $CDCl_3$) δ 7.57 (s, 1H), 7.37 – 7.26 (m, 3H), 7.23 – 7.13 (m, 4H), 7.12 – 7.04 (m, 3H), 6.99 (d, *J* = 7.7 Hz, 1H), 6.92 (t, *J* = 2.1 Hz, 1H), 6.87-6.82 (m, 1H), 6.79-6.75 (m, 1H), 6.72 – 6.63 (m, 2H), 5.80-5.72 (m, 1H), 5.31 (s, 1H), 4.94-4.84 (m, 1H), 4.50 (s, 2H), 4.34 – 4.27 (m, 1H), 4.03 – 3.93 (m, 2H), 3.84 (d, *J* = 5.4 Hz, 6H), 3.64 – 3.51 (m, 10H), 3.40-3.33 (m, 2H), 3.22 – 3.11 (m, 1H), 2.98 (d, *J* = 6.4 Hz, 2H), 2.66 – 2.48 (m, 2H), 2.30 – 2.18 (m, 1H), 2.10 – 1.94 (m, 5H), 1.80-1.59 (m, 5H), 1.54 – 1.30 (m, 4H), 1.23 – 1.10 (m, 6H), 0.92 – 0.77 (m, 9H).

$^{13}$C NMR (126 MHz, $CDCl_3$) δ 208.01, 171.07, 170.82, 169.83, 169.10, 168.41, 167.44, 157.55, 149.04, 147.52, 142.00, 136.39, 133.47, 130.18, 129.51, 128.63, 127.06,

120.29, 120.19, 114.44, 113.39, 111.88, 111.48, 76.66, 70.38, 69.85, 69.74, 67.53, 58.70, 56.82, 56.06, 55.98, 55.62, 54.66, 53.56, 51.42, 46.83, 44.30, 43.48, 39.32, 38.97, 38.31, 32.60, 31.38, 26.56, 25.06, 23.58, 23.30, 22.88, 21.33, 19.19, 18.76, 18.42, 17.40, 12.58, 8.89.

HRMS (ESI) m/z calcd for $C_{56}H_{78}N_6NaO_{14}$ for $[M+Na]^+$: 1081.5474, found: 1081.5471

**ZA30**: *(R)-1-(3-(((7S,10S)-7-benzyl-10-((S)-sec-butyl)-2,5,8,11,22-pentaoxo-15,18-dioxa-3,6,9,12,21-pentaazatricosan-23-yl)oxy)phenyl)-3-(3,4-dimethoxyphenyl)propyl (S)-1-(3,3-dimethyl-2-oxopentanoyl)piperidine-2-carboxylate*

The synthetic route of ZA30 was similar to ZA28.

$^1H$ NMR (500 MHz, $CDCl_3$) δ 7.55 (s, 1H), 7.35 – 7.26 (m, 3H), 7.22-7.13 (m, 4H), 7.11-7.05 (m, 3H), 7.00 – 6.97 (m, 1H), 6.92 (t, J = 2.0 Hz, 1H), 6.87 – 6.83 (m, 1H), 6.79-6.75 (m, 1H), 6.72 – 6.62 (m, 2H), 5.80-5.73 (m, 1H), 5.31 (s, 1H), 4.88 (s, 1H), 4.50 (s, 2H), 4.33 (t, J = 8.2 Hz, 1H), 4.02-3.91 (m, 2H), 3.86-3.83 (m, 6H), 3.62 – 3.52 (m, 10H), 3.52-3.43 (m, 1H), 3.40-3.32 (m, 2H), 3.22 – 3.10 (m, 1H), 2.98 (d, J = 6.7 Hz, 2H), 2.66 – 2.48 (m, 2H), 2.31 – 2.18 (m, 1H), 2.10 – 2.01 (m, 1H), 2.00 (s, 3H), 1.80 – 1.60 (m, 6H), 1.46 – 1.37 (m, 4H), 1.23 – 1.09 (m, 6H), 1.06-0.95 (m, 1H), 0.92 – 0.76 (m, 9H).

$^{13}C$ NMR (126 MHz, $CDCl_3$) δ 208.01, 171.07, 170.72, 169.83, 169.12, 168.41, 167.44, 157.55, 149.04, 147.52, 142.00, 136.37, 133.47, 130.17, 129.52, 128.63, 127.06, 120.28, 120.19, 114.45, 113.37, 111.87, 111.47, 76.66, 70.38, 69.85, 69.72, 67.52, 57.91, 56.06, 55.98, 55.61, 54.64, 53.56, 51.41, 46.83, 44.29, 43.47, 39.31, 38.97, 38.31, 37.60, 32.59, 31.37, 31.06, 26.55, 25.06, 25.02, 23.58, 23.30, 22.88, 21.32, 18.75, 17.39, 15.39, 11.41, 8.89.

HRMS (ESI) m/z calcd for $C_{57}H_{80}N_6NaO_{14}$ for $[M+Na]^+$: 1095.5630, found: 1095.5630

**ZA31**: *(R)-1-(3-(((14S,17S)-17-benzyl-14-((S)-sec-butyl)-2,13,16,19,22-pentaoxo-6,9-dioxa-3,12,15,18,21-pentaazatetracosyl)oxy)phenyl)-3-(3,4-dimethoxyphenyl)propyl (S)-1-(3,3-dimethyl-2-oxopentanoyl)piperidine-2-carboxylate*

ZA31: S-23 (51.3 mg, 0.053 mmol, 1 eq), N-Propionylglycine (8.29 mg, 0.063 mmol, 1.2 eq), and HATU (32.04 mg, 0.084 mmol, 1.6 eq) were dissolved in dry DMF (1.5 mL) at room temperature. DIPEA (27 μL, 0.158 mmol, 3.0 eq) was added to the reaction and stirred for 0.5 h. Upon completion, the reaction was concentrated, extracted with DCM (20 mL) and washed with brine (1 x 20 mL). The organic layer was dried by $Na_2SO_4$, concentrated, and the resulting residue purified by flash chromatography to provide ZA31 as colorless oil (27.5 mg, 0.025 mmol, 48%).

$^1$H NMR (500 MHz, $CDCl_3$) δ 7.38 (s, 1H), 7.31-7.26 (m, 1H), 7.25 – 7.14 (m, 4H), 7.13-7.03 (m, 4H), 7.01 – 6.94 (m, 1H), 6.92 (t, J = 2.0 Hz, 1H), 6.90 – 6.81 (m, 2H), 6.80-6.75 (m, 1H), 6.72 – 6.64 (m, 2H), 5.80-5.72 (m, 1H), 5.31 (s, 1H), 4.79 (s, 1H), 4.50 (s, 2H), 4.33 (t, J = 8.0 Hz, 1H), 4.03 – 3.87 (m, 2H), 3.86-3.83 (m, 6H), 3.63 – 3.51 (m, 10H), 3.41-3.32 (m, 2H), 3.17 (td, J = 13.2, 3.1 Hz, 1H), 3.00 (d, J = 6.4 Hz, 2H), 2.66-2.47 (m, 2H), 2.29 – 2.18 (m, 3H), 2.10-1.99 (m, 1H), 1.81 – 1.60 (m, 7H), 1.56 – 1.28 (m, 4H), 1.21 (d, J = 7.8 Hz, 5H), 1.14 – 1.08 (m, 4H), 1.08 – 0.96 (m, 1H), 0.92 – 0.77 (m, 9H).

$^{13}$C NMR (126 MHz, $CDCl_3$) δ 207.87, 174.72, 171.02, 170.66, 169.84, 169.23, 168.43, 167.45, 157.56, 149.04, 147.53, 141.99, 136.33, 133.47, 130.17, 129.49, 128.70, 127.12, 120.29, 120.19, 114.42, 113.42, 111.88, 111.48, 76.67, 70.40, 70.38, 69.85, 69.75, 67.54, 57.94, 56.06, 55.98, 54.70, 53.56, 51.42, 46.84, 44.30, 43.43, 39.33, 38.98, 38.79, 38.31, 37.51, 32.60, 31.38, 29.19, 26.56, 25.07, 24.98, 23.69, 23.58, 23.30, 21.33, 15.40, 11.43, 9.81, 8.89.

HRMS (ESI) m/z calcd for $C_{58}H_{82}N_6NaO_{14}$ for $[M+Na]^+$: 1109.5787, found: 1109.5781

**Synthesis of ZA2**

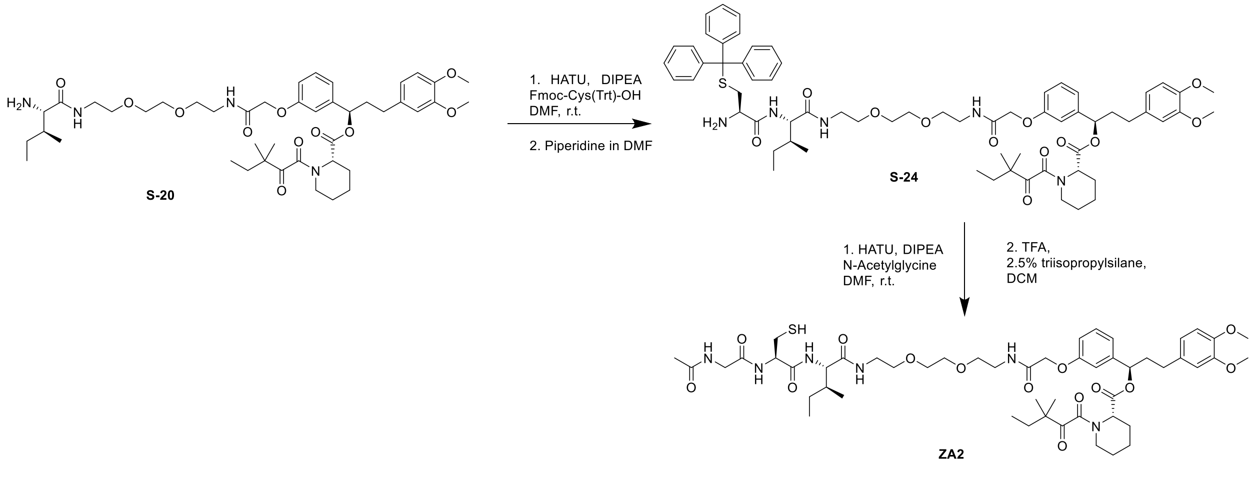


**ZA2**: *(R)-1-(3-(((7R,10S)-10-((S)-sec-butyl)-7-(mercaptomethyl)-2,5,8,11,22-pentaoxo-15,18-dioxa-3,6,9,12,21-pentaazatricosan-23-yl)oxy)phenyl)-3-(3,4-dimethoxyphenyl)propyl (S)-1-(3,3-dimethyl-2-oxopentanoyl)piperidine-2-carboxylate*

Step1: S-20 (115 mg, 0.139mmol, 1 eq), Fmoc-Cys(Trt)-OH (97.7 mg, 0.167 mmol, 1.2 eq), and HATU (84.6 mg, 0.222 mmol, 1.6 eq) were dissolved in dry DMF (3.0 mL) at room temperature. DIPEA (73 µL, 0.417 mmol, 3.0 eq) was added to the reaction and stirred for 0.5 h. Upon completion, the reaction was concentrated, extracted with DCM (20 mL) and washed with brine (1 x 20 mL). The organic layer was dried by $Na_2SO_4$, concentrated, and the resulting residue was used without further purification. The crude product was dissolved in DMF (3.0 mL), piperidine (34 µL, 0.344 mmol, 5 eq) was added to the reaction and stirred for 1 h. The reaction mixture was then concentrated and purified by flash chromatography to provide S-24 (80 mg, 0.068 mmol, 58% over 2 steps).

ZA2: S-24 (80 mg, 0.068 mmol, 1 eq), N-Acetylglycine (9.59 mg, 0.082 mmol, 1.2 eq), and HATU (41.5 mg, 0.109 mmol, 1.6 eq) were dissolved in dry DMF (2.0 mL) at room temperature. DIPEA (35 µL, 0.204 mmol, 3.0 eq) was added to the reaction and stirred for 0.5 h. Upon completion, the reaction was concentrated, extracted with DCM (20 mL) and washed with brine (1 x 20 mL). The organic layer was dried by $Na_2SO_4$, concentrated, and the resulting residue was used without further purification. The crude product was dissolved in DCM (3.0 mL), TFA (1.0 mL), triisopropylsilane (2.5%), was added to the reaction and stirred for 1 h. The reaction mixture was then concentrated and purified by flash chromatography to provide ZA2 (32 mg, 0.031 mmol, 46% over 2 steps).

$^{1}$H NMR (500 MHz, DMSO) δ 8.14 (t, J = 5.8 Hz, 1H), 8.10 – 8.04 (m, 2H), 7.94 (t, J = 5.6 Hz, 1H), 7.84 (d, J = 8.7 Hz, 1H), 7.30 (q, J = 7.6 Hz, 1H), 6.99 – 6.93 (m, 2H), 6.90 (dd, J = 8.1, 2.5 Hz, 1H), 6.84 (d, J = 8.2 Hz, 1H), 6.79 – 6.74 (m, 1H), 6.67 (dd, J = 8.1, 2.0 Hz, 1H), 5.73 – 5.66 (m, 1H), 5.15 (d, J = 5.7 Hz, 1H), 4.50 – 4.42 (m, 3H), 4.14 – 4.07 (m, 1H), 3.77 – 3.69 (m, 8H), 3.47 (s, 4H), 3.45 – 3.35 (m, 4H), 3.32 – 3.20 (m, 5H), 3.19 – 3.07 (m, 2H), 2.81 – 2.65 (m, 2H), 2.61 – 2.51 (m, 1H), 2.30 – 2.21 (m, 2H), 2.19 – 2.08 (m, 1H), 2.07 – 2.00 (m, 1H), 1.85 (s, 3H), 1.76 – 1.51 (m, 6H), 1.45 – 1.30 (m, 2H), 1.28 – 1.19 (m, 1H), 1.17 – 0.98 (m, 7H), 0.87 – 0.68 (m, 9H).

$^{13}$C NMR (126 MHz, MeOD) δ 209.01, 173.95, 173.50, 172.05, 171.79, 171.12, 170.98, 169.10, 159.34, 150.46, 148.87, 143.37, 135.23, 131.02, 121.71, 121.04, 115.70, 114.29, 113.62, 113.28, 78.20, 77.97, 71.33, 70.48, 68.37, 59.57, 56.81, 56.57, 56.47, 52.83, 47.70, 45.71, 43.66, 40.33, 39.96, 39.24, 37.97, 33.59, 32.26, 27.35, 26.86, 25.96, 25.91, 24.21, 23.84, 23.65, 22.42, 22.15, 15.95, 11.50, 9.11.

HRMS (ESI) m/z calcd for $C_{51}H_{76}N_6NaO_{14}S$ for $[M+Na]^+$: 1051.5038, found: 1051.5030

**Synthesis of ZA32-34**

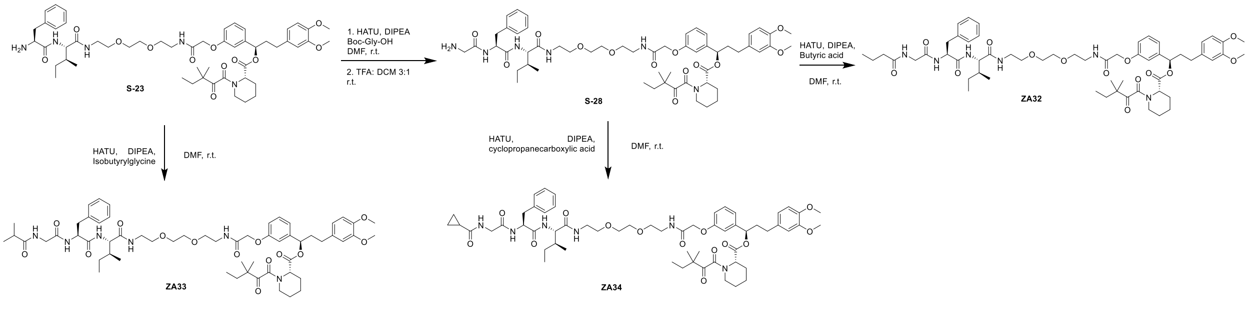

**ZA32**: *(R)-1-(3-(((14S,17S)-17-benzyl-14-((S)-sec-butyl)-2,13,16,19,22-pentaoxo-6,9-dioxa-3,12,15,18,21-pentaazapentacosyl)oxy)phenyl)-3-(3,4-dimethoxyphenyl)propyl (S)-1-(3,3-dimethyl-2-oxopentanoyl)piperidine-2-carboxylate*

Step1: S-23 (90 mg, 0.092 mmol, 1 eq), Boc-Gly-OH (19.4 mg, 0.111 mmol, 1.2 eq), and HATU (56.2 mg, 0.147 mmol, 1.6 eq) were dissolved in dry DMF (3.0 mL) at room temperature. DIPEA (48 μL, 0.277 mmol, 3.0 eq) was added to the reaction and stirred for 0.5 h. Upon completion, the reaction was concentrated, extracted with DCM (20 mL) and washed with brine (1 x 20 mL). The organic layer was dried by $Na_2SO_4$, concentrated, and the resulting residue was used without further purification. The crude product was dissolved in DCM (3.0 mL), TFA (1.0 mL) was added to the reaction and stirred for 0.5 h. The reaction mixture was then concentrated and purified by flash chromatography to provide S-28 as a colorless oil (75.5 mg, 0.073 mmol, 79% over 2 steps).

ZA32: S-23 (41.5 mg, 0.040 mmol, 1 eq), butyric acid (4.25 mg, 0.048 mmol, 1.2 eq), and HATU (24.5 mg, 0.064 mmol, 1.6 eq) were dissolved in dry DMF (1.5 mL) at room temperature. DIPEA (21 μL, 0.120 mmol, 3.0 eq) was added to the reaction and stirred for 0.5 h. Upon completion, the reaction was concentrated, extracted with DCM (20 mL) and washed with brine (1 x 20 mL). The organic layer was dried by $Na_2SO_4$, concentrated, and the resulting residue purified by flash chromatography to provide ZA32 as colorless oil (13.4 mg, 0.012 mmol, 30%).

$^1$H NMR (500 MHz, $CDCl_3$) δ 7.32 – 7.27 (m, 1H), 7.26 – 7.17 (m, 3H), 7.16 – 7.02 (m, 4H), 7.01 – 6.88 (m, 3H), 6.87 – 6.73 (m, 3H), 6.72 – 6.57 (m, 3H), 5.80 – 5.72 (m, 1H), 5.33 – 5.28 (m, 1H), 4.76 – 4.64 (m, 1H), 4.50 (s, 2H), 4.28 (t, *J* = 8.0 Hz, 1H), 3.97 – 3.79 (m, 8H), 3.63 – 3.50 (m, 10H), 3.49 – 3.33 (m, 3H), 3.22 – 3.12 (m, 1H), 3.03 (d, *J* = 6.5 Hz, 2H), 2.68 – 2.47 (m, 2H), 2.37 (d, *J* = 13.7 Hz, 1H), 2.31 – 2.13 (m, 3H), 2.11 – 1.99 (m, 1H), 1.88 – 1.76 (m, 2H), 1.70 – 1.56 (m, 6H), 1.54 – 1.32 (m, 3H), 1.23 – 1.10 (m, 6H), 1.08 – 0.96 (m, 1H), 0.95 – 0.77 (m, 12H).

$^{13}C$ NMR (126 MHz, $CDCl_3$) δ 208.04, 174.05, 170.98, 170.67, 169.85, 169.41, 168.53, 167.47, 157.57, 149.05, 147.54, 142.00, 136.26, 133.48, 130.19, 129.45, 128.80, 127.21, 120.30, 120.19, 114.36, 113.49, 111.89, 111.49, 76.69, 70.40, 70.37, 69.89, 69.81, 67.55, 58.11, 56.84, 56.08, 56.00, 54.83, 51.44, 46.85, 44.31, 43.51, 39.34, 39.00, 38.42, 38.33, 38.10, 37.31, 32.61, 31.39, 26.58, 25.08, 24.94, 23.71, 23.59, 23.32, 21.33, 19.05, 15.47, 13.92, 11.47, 8.90.

HRMS (ESI) m/z calcd for $C_{59}H_{84}N_6NaO_{14}$ for $[M+Na]^+$: 1123.5943, found: 1123.5938

**ZA33**: *(R)-1-(3-(((14S,17S)-17-benzyl-14-((S)-sec-butyl)-23-methyl-2,13,16,19,22-pentaoxo-6,9-dioxa-3,12,15,18,21-pentaazatetracosyl)oxy)phenyl)-3-(3,4-dimethoxyphenyl)propyl (S)-1-(3,3-dimethyl-2-oxopentanoyl)piperidine-2-carboxylate*

The synthetic route of ZA33 was similar to ZA32.

$^1H$ NMR (500 MHz, $CDCl_3$) δ 7.33 – 7.27 (m, 1H), 7.26 – 7.21 (m, 2H), 7.21 – 7.15 (m, 2H), 7.15 – 7.11 (m, 2H), 7.11 – 7.03 (m, 2H), 7.01 – 6.97 (m, 1H), 6.94 – 6.91 (m, 1H), 6.87 – 6.83 (m, 1H), 6.82 – 6.74 (m, 2H), 6.72 – 6.62 (m, 3H), 5.80 – 5.73 (m, 1H), 5.32 – 5.30 (m, 1H), 4.73 (q, *J* = 6.8 Hz, 1H), 4.50 (s, 2H), 4.35 – 4.28 (m, 1H), 4.00 – 3.80 (m, 8H), 3.64 – 3.44 (m, 11H), 3.41 – 3.32 (m, 2H), 3.22 – 3.13 (m, 1H), 3.08 – 2.96 (m, 2H), 2.66 – 2.48 (m, 2H), 2.47 – 2.32 (m, 2H), 2.30 – 2.18 (m, 1H), 2.11 – 1.99 (m, 1H), 1.84 – 1.77 (m, 2H), 1.73 – 1.60 (m, 4H), 1.53 – 1.45 (m, 1H), 1.43 – 1.30 (m, 2H), 1.23 – 1.08 (m, 12H), 1.08 – 0.96 (m, 1H), 0.92 – 0.77 (m, 9H).

$^{13}C$ NMR (126 MHz, $CDCl_3$) δ 208.02, 177.88, 170.92, 170.55, 169.84, 169.28, 168.44, 167.46, 157.53, 149.05, 147.54, 141.99, 136.26, 133.48, 130.18, 129.47, 128.79, 127.21, 120.30, 120.20, 114.40, 113.46, 111.89, 111.48, 76.68, 70.42, 69.86, 69.78, 67.55, 58.02, 56.84, 56.07, 55.99, 54.74, 51.43, 46.85, 44.31, 43.46, 39.33, 38.99, 38.59, 38.32, 37.44, 35.18, 32.61, 31.39, 26.57, 25.08, 24.97, 23.70, 23.60, 23.31, 21.34, 21.15, 19.62, 15.44, 11.47, 8.90.

HRMS (ESI) m/z calcd for $C_{59}H_{84}N_6NaO_{14}$ for $[M+Na]^+$: 1123.5943, found: 1123.5942

**ZA34**: *(R)-1-(3-(((6S,9S)-6-benzyl-9-((S)-sec-butyl)-1-cyclopropyl-1,4,7,10,21-pentaoxo-14,17-dioxa-2,5,8,11,20-pentaazadocosan-22-yl)oxy)phenyl)-3-(3,4-dimethoxyphenyl)propyl (S)-1-(3,3-dimethyl-2-oxopentanoyl)piperidine-2-carboxylate*

The synthetic route of ZA34 was similar to ZA33.

$^1$H NMR (500 MHz, $CDCl_3$) δ 7.32 – 7.27 (m, 1H), 7.25 – 7.16 (m, 3H), 7.15 – 7.06 (m, 4H), 7.02 – 6.94 (m, 2H), 6.94 – 6.88 (m, 2H), 6.88 – 6.81 (m, 2H), 6.81 – 6.75 (m, 1H), 6.72 – 6.64 (m, 2H), 5.80 – 5.72 (m, 1H), 5.31 (d, *J* = 2.5 Hz, 1H), 4.73 (q, *J* = 6.8 Hz, 1H), 4.50 (s, 2H), 4.33 – 4.27 (m, 1H), 4.02 – 3.80 (m, 8H), 3.63 – 3.50 (m, 10H), 3.49 – 3.40 (m, 1H), 3.40 – 3.32 (m, 2H), 3.22 – 3.13 (m, 1H), 3.09 – 2.97 (m, 2H), 2.67 – 2.48 (m, 2H), 2.37 (d, *J* = 13.7 Hz, 1H), 2.30 – 2.18 (m, 1H), 2.11 – 1.99 (m, 1H), 1.87 – 1.73 (m, 3H), 1.69 – 1.59 (m, 3H), 1.53 – 1.43 (m, 2H), 1.43 – 1.30 (m, 2H), 1.23 – 1.11 (m, 6H), 1.06 – 0.95 (m, 1H), 0.95 – 0.69 (m, 13H).

$^{13}$C NMR (126 MHz, $CDCl_3$) δ 208.04, 174.69, 170.98, 170.64, 169.85, 169.48, 168.46, 167.47, 157.54, 149.06, 147.54, 141.99, 136.26, 133.48, 130.19, 129.46, 128.81, 127.21, 120.30, 120.19, 114.40, 113.47, 111.89, 111.49, 76.70, 70.41, 69.85, 69.79, 67.57, 58.05, 56.84, 56.08, 56.00, 54.77, 53.57, 51.44, 46.85, 44.31, 43.85, 39.34, 39.01, 38.38, 38.33, 37.25, 32.61, 31.40, 26.58, 25.08, 24.91, 23.71, 23.59, 23.32, 21.34, 15.49, 14.34, 11.50, 8.90, 7.71, 7.65.

HRMS (ESI) m/z calcd for $C_{59}H_{82}N_6NaO_{14}$ for $[M+Na]^+$: 1121.5787, found 1121.5784.

**Synthesis of ZA35 and ZA36**

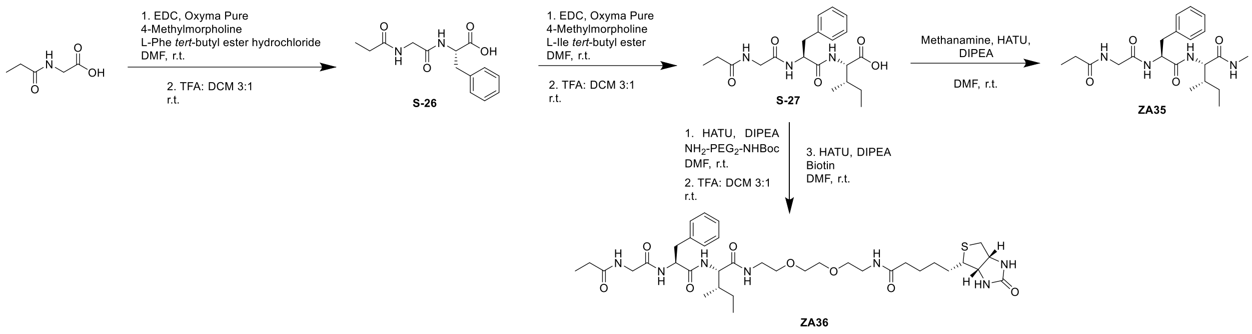


**ZA35**: *(2S,3S)-N,3-dimethyl-2-((S)-3-phenyl-2-(2-propionamidoacetamido)propanamido)pentanamide*

Step1: N-Propionylglycine (200 mg, 1.53 mmol, 1 eq), L-Phe *tert*-butyl ester hydrochloride (412.79 mg, 1.6 mmol, 1.05 eq), Oxyma Pure (433.5 mg, 3.05 mmol, 2 eq) were dissolved in dry DMF (30 mL) on ice. 1-(3-Dimethylaminopropyl)-3-ethylcarbodiimide (EDC) hydrochloride hydrochloride (438.6 mg, 2.29 mmol, 1.5 eq) and 4-Methylmorpholine (838 µL, 7.63 mmol, 5 eq) were then added to the reaction and stirred at room temperature for 2 h. Upon completion, the reaction was concentrated. The crude product was first extracted with EtOAC (40 mL) and 1N HCl (10 mL); then extracted with saturated $NaHCO_3$ solution (10 mL) The organic layer was dried by $Na_2SO_4$, concentrated, and the resulting residue was used without further purification. The crude product was dissolved in DCM (20 mL), TFA (10 mL) was added to the reaction and stirred for 1 h. The reaction mixture was then concentrated to give crude S-26 without further purification.

Step2: Crude S-26 (420 mg, 1 eq), L-Ile *tert*-butyl ester hydrochloride (337.65 mg, 1.51 mmol, 1 eq), Oxyma Pure (346.6 mg, 2.57 mmol, 1.7 eq) were dissolved in dry DMF (20 mL) on ice. 1-(3-Dimethylaminopropyl)-3-ethylcarbodiimide (EDC) hydrochloride hydrochloride (376.1 mg, 1.96 mmol, 1.3 eq) and 4-Methylmorpholine (746 µL, 6.79 mmol, 4.5 eq) were then added to the reaction and stirred at room temperature for 2 hs. Upon completion, the reaction was concentrated. The crude product was first extracted with EtOAC (40 mL) and 1N HCl (10 mL); then extracted with saturated $NaHCO_3$ solution (10 mL) The organic layer was dried by $Na_2SO_4$, concentrated, and the resulting residue was used without further purification. The crude product was dissolved in DCM (20 mL), TFA (10 mL) was added to the reaction and stirred for 1 h. The reaction mixture was then concentrated and purified by flash chromatography to provide S-27 (266 mg, 0.679 mmol, 45% over 4 steps).

$^{1}$H NMR (500 MHz, MeOD) δ 8.18 (d, *J* = 8.4 Hz, 1H), 7.92 (d, *J* = 7.9 Hz, 1H), 7.31 – 7.17 (m, 5H), 4.77 – 4.69 (m, 1H), 4.40 – 4.33 (m, 1H), 3.87 – 3.74 (m, 2H), 3.19 – 3.13 (m, 1H), 2.98 – 2.90 (m, 1H), 2.30 – 2.22 (m, 2H), 1.93 – 1.89 (m, 1H), 1.60 – 1.48 (m, 1H), 1.33 – 1.20 (m, 1H), 1.13 (t, *J* = 7.6 Hz, 3H), 0.98 – 0.91 (m, 6H).

ZA35: S-27 (40 mg, 0.102 mmol, 1 eq), methenamine hydrochloride (7.6 mg, 0.112mmol, 1.1 eq), HATU (50.51 mg, 0.133 mmol, 1.3 eq) were dissolved in dry DMF (3 mL) at room temperature. DIPEA (89 µL, 0.511 mmol, 5 eq) was added to the reaction and stirred for 0.5 h. Upon completion, the reaction was concentrated, extracted with DCM (20 mL) and washed with brine (1 x 20 mL). The organic layer was dried by $Na_2SO_4$, concentrated, and the resulting residue purified by flash chromatography to provide ZA35 as white powder (13 mg, 0.032 mmol, 32%).

$^{1}$H NMR (500 MHz, MeOD) δ 8.00 – 7.75 (m, 1H), 7.34 – 7.21 (m, 6H), 4.71 – 4.60 (m, 1H), 4.16 (d, J = 7.7 Hz, 1H), 3.93 – 3.76 (m, 2H), 3.19 – 3.11 (m, 1H), 3.02 – 2.93 (m, 1H), 2.73 (s, 3H), 2.34 – 2.24 (m, 2H), 1.90 – 1.80 (m, 1H), 1.59 – 1.44 (m, 1H), 1.15 (dt, J = 9.6, 7.6 Hz, 4H), 0.96 – 0.87 (m, 6H).

$^{13}$C NMR (126 MHz, MeOD) δ 177.33, 173.75, 173.04, 171.60, 137.93, 130.29, 129.41, 127.77, 79.31, 59.29, 55.80, 54.66, 43.51, 38.54, 37.85, 29.75, 26.22, 25.84, 15.81, 11.40, 10.07.

HRMS (ESI) m/z calcd for $C_{21}H_{32}N_4NaO_4$ for $[M+Na]^+$: 427.2321, found: 427.2319

**ZA36**: *(2S,3S)-3-methyl-N-(2-(2-(2-(5-((3aS,4S,6aR)-2-oxohexahydro-1H-thieno[3,4-d]imidazol-4-yl)pentanamido)ethoxy)ethoxy)ethyl)-2-((S)-3-phenyl-2-(2-propionamidoacetamido)propanamido)pentanamide*

ZA36: S-27 (50 mg, 0.127 mmol, 1 eq), $NH_2$-$PEG_2$-NHBoc (34.89 mg, 0.140mmol, 1.1 eq), HATU (63.13 mg, 0.166 mmol, 1.3 eq) were dissolved in dry DMF (3 mL) at room temperature. DIPEA (67 μL, 0.383 mmol, 3 eq) was added to the reaction and stirred for 0.5 h. The organic layer was dried by $Na_2SO_4$, concentrated, and the resulting residue was used without further purification. The crude product was dissolved in DCM (3.0 mL), TFA (1.0 mL) was added to the reaction and stirred for 0.5 h. The reaction mixture was then concentrated and dissolved in dry DMF with HATU (61.59 mg, 0.161 mmol, 1.3 eq) and biotin (33.49 mg, 0.137 mmol, 1.1 eq) at room temperature. DIPEA (65 μL, 0.373 mmol, 3 eq) was added to the reaction and stirred for 0.5 h. Upon completion, the reaction was concentrated, extracted with DCM (20 mL) and washed with brine (1 x 20 mL). The organic layer was dried by $Na_2SO_4$, concentrated, and the resulting residue purified by flash chromatography to provide ZA36 (34.2 mg, 0.046 mmol, 37% over 3 steps).

$^1$H NMR (500 MHz, DMSO) δ 8.00 – 7.90 (m, 3H), 7.81 (t, J = 5.7 Hz, 1H), 7.26 – 7.13 (m, 5H), 6.47 – 6.32 (m, 1H), 4.62 – 4.54 (m, 1H), 4.33 – 4.27 (m, 1H), 4.16 – 4.09 (m, 2H), 3.73 – 3.65 (m, 1H), 3.59 – 3.45 (m, 5H), 3.43 – 3.35 (m, 4H), 3.30 (d, J = 11.7 Hz, 3H), 3.21 – 3.13 (m, 3H), 3.12 – 3.05 (m, 1H), 3.02 – 2.94 (m, 1H), 2.85 – 2.73 (m, 2H), 2.57 (d, J = 12.4 Hz, 1H), 2.13 – 2.03 (m, 4H), 1.74 – 1.54 (m, 2H), 1.54 – 1.38 (m, 4H), 1.35 – 1.26 (m, 2H), 1.11 – 1.00 (m, 1H), 0.96 (t, J = 7.6 Hz, 3H), 0.85 – 0.77 (m, 6H).

$^{13}$C NMR (126 MHz, DMSO) δ 173.19, 172.10, 170.83, 170.54, 168.84, 162.71, 162.67, 137.58, 129.25, 127.95, 126.20, 69.52, 69.50, 69.15, 68.90, 61.02, 60.92, 59.18, 59.07, 56.93, 55.40, 53.49, 41.85, 38.44, 38.41, 37.43, 36.66, 35.09, 28.20, 28.15, 28.02, 25.25, 24.37, 15.25, 11.06, 9.72.

HRMS (ESI) m/z calcd for $C_{36}H_{58}N_7O_8S$ for $[M+H]^+$: 748.4068, found: 748.4059

### Synthesis of ZA11 and ZA12

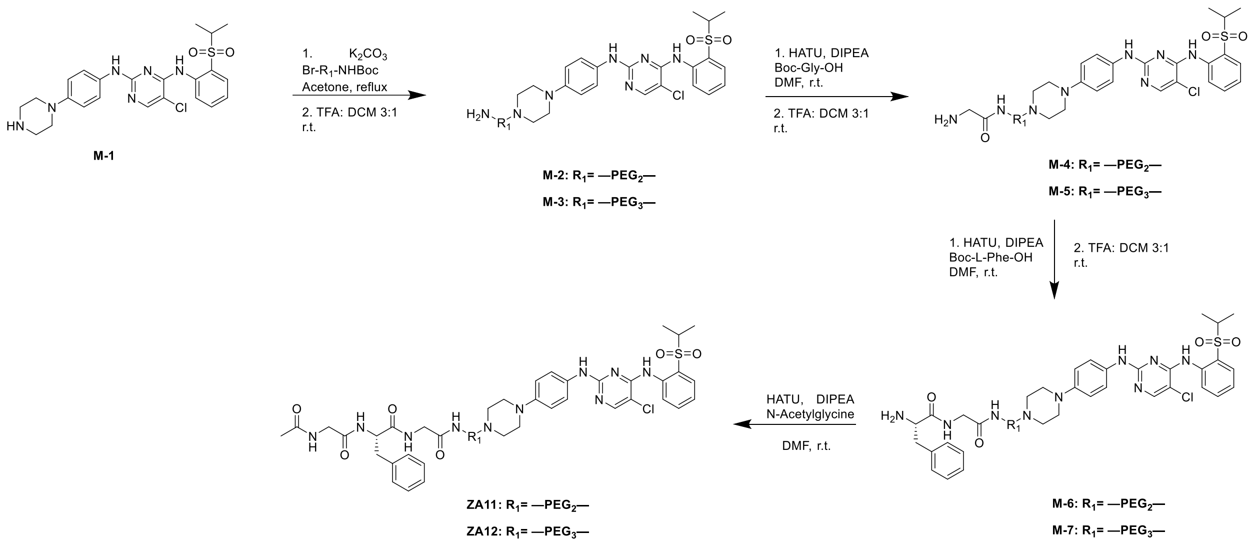


**ZA11**: *(S)-2-(2-acetamidoacetamido)-N-(2-((2-(2-(2-(4-(4-((5-chloro-4-((2-(isopropylsulfonyl)phenyl)amino)pyrimidin-2-yl)amino)phenyl)piperazin-1-yl)ethoxy)ethoxy)ethyl)amino)-2-oxoethyl)-3-phenylpropanamide*

Step1: M-1 (120 mg, 0.246 mmol, 1 eq), $K_2CO_3$ (68.11 mg, 0.492 mmol, 2.0 eq) and Br-PEG2-NHBoc (92.3 mg, 0.296 mmol, 1.2 eq) were dissolved in acetone and refluxed overnight. Upon completion, the reaction was concentrated, extracted with DCM (30 mL) and washed with brine (1 x 20 mL). The organic layer was dried by $Na_2SO_4$, concentrated, and the resulting residue was used without further purification. The crude product was dissolved in DCM (2.0 mL), TFA (0.5 mL) was added to the reaction and stirred for 0.5 h. The reaction mixture was then concentrated and purified by flash chromatography to provide M-2 (88 mg, 0.142 mmol, 58% over 2 steps).

$^1$H NMR (500 MHz, $CDCl_3$) δ 9.65 (s, 1H), 8.55 (d, *J* = 8.4 Hz, 1H), 8.07 (s, 1H), 7.91 – 7.84 (m, 1H), 7.59 – 7.51 (m, 1H), 7.43 – 7.37 (m, 3H), 7.26 – 7.21 (m, 1H), 6.88 – 6.82 (m, 2H), 3.89 – 3.83 (m, 2H), 3.72 – 3.67 (m, 2H), 3.65 – 3.59 (m, 5H), 3.38 (s, 5H), 3.28 – 3.18 (m, 4H), 3.14 – 3.10 (m, 2H), 1.30 (dd, *J* = 6.9, 1.4 Hz, 6H).

Step2: M-2 (88 mg, 0.142 mmol, 1.0 eq), HATU (86.6 mg, 0.228 mmol, 1.6 eq) and Boc-Gly-OH (29.9 mg, 0.171 mmol, 1.2 eq) were dissolved in dry DMF (3.0 mL) at room temperature. DIPEA (74 µL, 0.427 mmol, 5.0 eq) was added to the reaction and stirred for 1 h. Upon completion, the reaction was concentrated, extracted with DCM (20 mL) and washed with brine (1 x 20 mL). The organic layer was dried by $Na_2SO_4$, concentrated, and the resulting residue was used without further purification. The crude product was dissolved in DCM (3.0 mL), TFA (1.0 mL) was added to the reaction and stirred for 0.5 h. The reaction mixture was then concentrated and purified by flash chromatography to provide M-4 (53 mg, 0.078 mmol, 55% over 2 steps).

$^1$H NMR (500 MHz, MeOD) δ 8.66 (s, 1H), 8.16 – 8.12 (m, 1H), 7.95 – 7.89 (m, 1H), 7.73 – 7.66 (m, 1H), 7.53 – 7.46 (m, 2H), 7.39 – 7.32 (m, 1H), 7.05 – 6.99 (m, 2H), 3.97 – 3.89 (m, 2H), 3.78 – 3.68 (m, 8H), 3.62 (t, *J* = 5.7 Hz, 3H), 3.51 – 3.45 (m, 5H), 3.39 – 3.35 (m, 1H), 1.30 – 1.25 (m, 6H).

Step3: M-4 (53 mg, 0.078 mmol, 1.0 eq), HATU (47.75 mg, 0.126 mmol, 1.6 eq) and Boc-L-Phe-OH (24.99 mg, 0.094 mmol, 1.2 eq) were dissolved in dry DMF (3.0 mL) at room temperature. DIPEA (41 µL, 0.235 mmol, 5.0 eq) was added to the reaction and stirred for 1 h. Upon completion, the reaction was concentrated, extracted with DCM (20 mL) and washed with brine (1 x 20 mL). The organic layer was dried by $Na_2SO_4$, concentrated, and the resulting residue was used without further purification. The crude product was dissolved in DCM (3.0 mL), TFA (1.0 mL) was added to the reaction and stirred for 0.5 h. The reaction mixture was then concentrated and purified by flash chromatography to provide M-6 (40 mg, 0.049 mmol, 62% over 2 steps).

$^1$H NMR (500 MHz, MeOD) δ 8.65 (d, *J* = 8.4 Hz, 1H), 8.11 (s, 1H), 7.92 – 7.87 (m, 1H), 7.71 – 7.64 (m, 1H), 7.48 (d, *J* = 8.8 Hz, 2H), 7.40 – 7.26 (m, 8H), 7.03 – 6.96 (m, 2H), 4.13 – 4.07 (m, 1H), 3.92 – 3.83 (m, 3H), 3.74 – 3.67 (m, 4H), 3.59 (t, *J* = 5.7 Hz, 2H), 3.50 (s, 4H), 3.46 – 3.40 (m, 6H), 3.36 (s, 2H), 3.28 – 3.21 (m, 1H), 3.09 – 2.99 (m, 1H), 1.26 (d, *J* = 6.8 Hz, 6H).

ZA11: M-6 (40 mg, 0.049 mmol, 1 eq), N-Acetylglycine (6.83 mg, 0.058 mmol, 1.2 eq), and HATU (29.59 mg, 0.078 mmol, 1.6 eq) were dissolved in dry DMF (1.5 mL) at room temperature. DIPEA (25 μL, 0.146 mmol, 3.0 eq) was added to the reaction and stirred for 0.5 h. Upon completion, the reaction was concentrated, extracted with DCM (20 mL) and washed with brine (1 x 20 mL). The organic layer was dried by $Na_2SO_4$, concentrated, and the resulting residue purified by flash chromatography to provide ZA11 (18 mg, 0.020 mmol, 40.2%).

$^1$H NMR (500 MHz, $CDCl_3$) δ 9.61 (s, 1H), 8.60 (dd, J = 8.5, 1.1 Hz, 1H), 8.09 (s, 1H), 7.91 – 7.86 (m, 1H), 7.59 – 7.52 (m, 1H), 7.41 – 7.34 (m, 2H), 7.31 – 7.25 (m, 2H), 7.24 – 7.19 (m, 3H), 7.18 – 7.13 (m, 2H), 7.09 – 7.03 (m, 1H), 7.00 – 6.95 (m, 2H), 6.92 – 6.85 (m, 2H), 6.80 (s, 1H), 4.64 (q, J = 7.1 Hz, 1H), 3.96 – 3.83 (m, 3H), 3.82 – 3.74 (m, 1H), 3.69 – 3.59 (m, 6H), 3.57 (t, J = 5.2 Hz, 2H), 3.50 – 3.36 (m, 3H), 3.29 – 3.08 (m, 6H), 3.05 – 2.97 (m, 1H), 2.83 – 2.59 (m, 5H), 1.97 (s, 3H), 1.31 (d, J = 6.8 Hz, 6H).

$^{13}$C NMR (126 MHz, $CDCl_3$) δ 171.41, 171.13, 169.74, 168.93, 158.28, 155.50, 155.36, 147.63, 138.64, 136.43, 134.62, 131.94, 131.36, 129.33, 128.86, 127.29, 124.47, 123.53, 123.16, 122.30, 116.91, 105.98, 70.51, 70.30, 69.75, 68.35, 57.95, 55.73, 55.08, 53.64, 53.57, 50.99, 49.61, 43.74, 43.10, 39.52, 37.97, 22.98, 15.50.

HRMS (ESI) m/z calcd for $C_{44}H_{58}ClN_{10}O_8S$ for $[M+H]^+$: 921.3848, found: 921.3853

**ZA12**: *(S)-2-(2-acetamidoacetamido)-N-(14-(4-(4-((5-chloro-4-((2-(isopropylsulfonyl)phenyl)amino)pyrimidin-2-yl)amino)phenyl)piperazin-1-yl)-2-oxo-6,9,12-trioxa-3-azatetradecyl)-3-phenylpropanamide*

The synthetic route of ZA12 was similar to ZA11.

$^1$H NMR (500 MHz, $CDCl_3$) δ 9.60 (s, 1H), 8.63 – 8.57 (m, 1H), 8.07 (s, 1H), 7.88 (dd, J = 7.9, 1.7 Hz, 1H), 7.58 – 7.51 (m, 1H), 7.45 – 7.34 (m, 3H), 7.29 – 7.11 (m, 9H), 6.95 (t,

J = 5.3 Hz, 1H), 6.92 – 6.84 (m, 2H), 4.72 – 4.64 (m, 1H), 3.94 – 3.86 (m, 2H), 3.85 – 3.75 (m, 2H), 3.69 – 3.58 (m, 10H), 3.53 (t, J = 5.1 Hz, 2H), 3.44 – 3.36 (m, 2H), 3.28 – 3.08 (m, 6H), 3.03 – 2.95 (m, 1H), 2.74 – 2.64 (m, 6H), 1.96 (s, 3H), 1.30 (d, J = 6.8 Hz, 6H).

$^{13}$C NMR (126 MHz, $CDCl_3$) δ 171.35, 171.09, 169.75, 168.87, 158.29, 155.47, 155.31, 147.61, 138.63, 136.57, 134.63, 131.91, 131.31, 129.35, 128.76, 127.17, 124.40, 123.54, 123.11, 122.30, 116.83, 105.82, 77.36, 70.72, 70.62, 70.33, 70.09, 69.81, 68.21, 57.96, 55.71, 54.88, 53.62, 49.52.

HRMS (ESI) m/z calcd for $C_{46}H_{62}ClN_{10}O_9S$ for $[M+H]^+$: 965.4105, found: 965.4102

**Synthesis of ZA37**

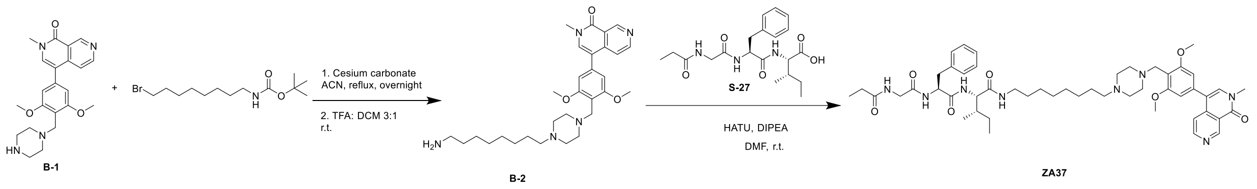


B-1 was synthesized as previously described[4].

Step1: B-1 (65 mg, 0.165 mmol, 1 eq), caesium carbonate (107.4 mg, 0.330 mmol, 2.0 eq) and tert-butyl (8-bromooctyl)carbamate (60.95 mg, 0.198 mmol, 1.2 eq) were dissolved in ACN and refluxed overnight. Upon completion, the reaction was concentrated, extracted with DCM (30 mL) and washed with brine (1 x 20 mL). The organic layer was dried by $Na_2SO_4$, concentrated, and the resulting residue was used without further purification. The crude product was dissolved in DCM (3.0 mL), TFA (1 mL) was added to the reaction and stirred for 0.5 h. The reaction mixture was then concentrated and purified by flash chromatography to provide B-2 (58 mg, 0.111 mmol, 68% over 2 steps).

$^{1}H$ NMR (500 MHz, MeOD) δ 9.50 (s, 1H), 8.66 (s, 1H), 7.78 (s, 1H), 7.66 (d, 1H), 6.83 (s, 2H), 4.37 (s, 2H), 3.93 (s, 6H), 3.69 (s, 3H), 3.54 – 3.33 (m, 10H), 3.06 – 2.99 (m, 2H), 2.95 – 2.88 (m, 2H), 1.76 – 1.61 (m, 4H), 1.45 – 1.35 (m, 8H).

ZA37: B-2 (58 mg, 0.111 mmol, 1 eq), S-27 (37 mg, 0.095 mmol, 0.85 eq), and HATU (46.5 mg, 0.122 mmol, 1.1 eq) were dissolved in dry DMF (1.5 mL) at room temperature. DIPEA (58 µL, 0.333 mmol, 3.0 eq) was added to the reaction and stirred for 1 h. Upon completion, the reaction was concentrated, and the resulting residue purified by flash chromatography to provide ZA37 (22 mg, 0.025 mmol, 22%).

$^{1}H$ NMR (500 MHz, $CDCl_3$) δ 9.66 – 9.62 (m, 1H), 8.65 (d, J = 5.6 Hz, 1H), 7.42 (dd, J = 5.6, 0.9 Hz, 1H), 7.28 (s, 1H), 7.24 – 7.02 (m, 6H), 6.51 (s, 2H), 4.77 – 4.71 (m, 1H), 4.23 – 4.15 (m, 1H), 4.00 – 3.73 (m, 10H), 3.65 (s, 3H), 3.28 – 3.17 (m, 1H), 3.13 – 2.90 (m, 3H), 2.88 – 2.41 (m, 7H), 2.34 (t, J = 8.2 Hz, 2H), 2.27 – 2.18 (m, 3H), 1.85 – 1.64 (m, 1H), 1.52 – 1.35 (m, 5H), 1.32 – 1.19 (m, 8H), 1.15 – 1.05 (m, 3H), 1.04 – 0.95 (m, 1H), 0.85 – 0.73 (m, 6H).

$^{13}C$ NMR (126 MHz, $CDCl_3$) δ 174.85, 171.02, 170.93, 170.85, 169.32, 169.24, 161.62, 159.73, 151.55, 150.76, 142.01, 136.31, 136.14, 135.69, 129.44, 128.57, 127.02, 120.59, 118.20, 117.72, 105.37, 77.36, 58.68, 57.95, 57.91, 55.96, 54.55, 54.46, 52.82, 52.05, 48.71, 43.07, 39.59, 39.46, 38.54, 37.26, 37.15, 29.45, 29.32, 29.31, 29.22, 29.18, 29.14, 27.53, 26.93, 26.49, 24.90, 15.34, 11.19, 9.73.

HRMS (ESI) m/z calcd for $C_{50}H_{71}N_8O_7$ for $[M+H]^+$: 895.5446, found: 895.5447

### $^{1}$H NMR spectrum of ZA1

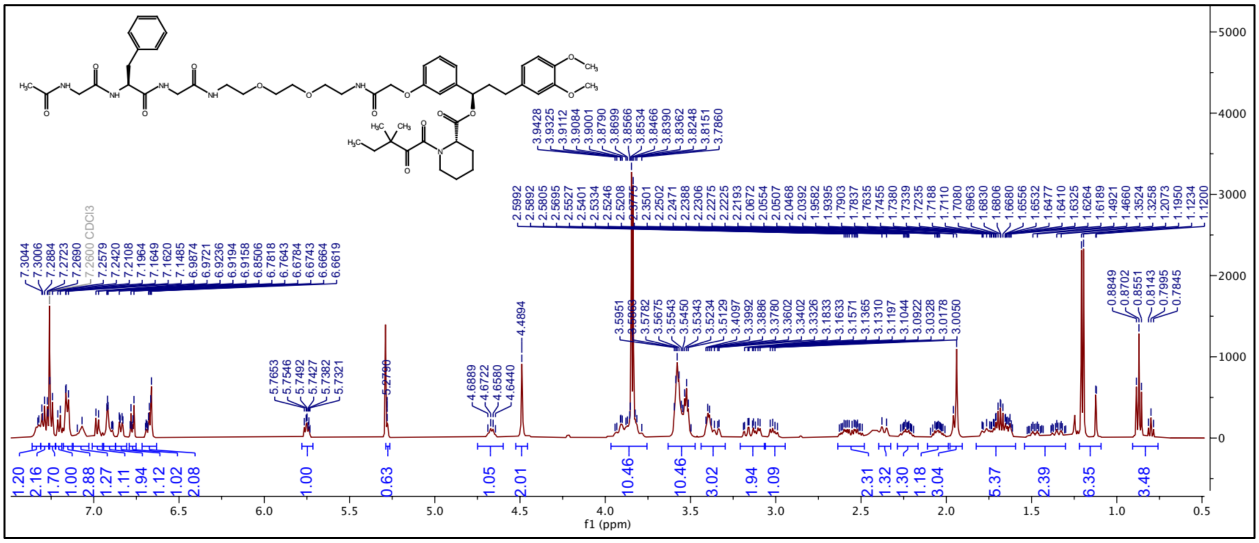

### $^{13}$C NMR spectrum of ZA1

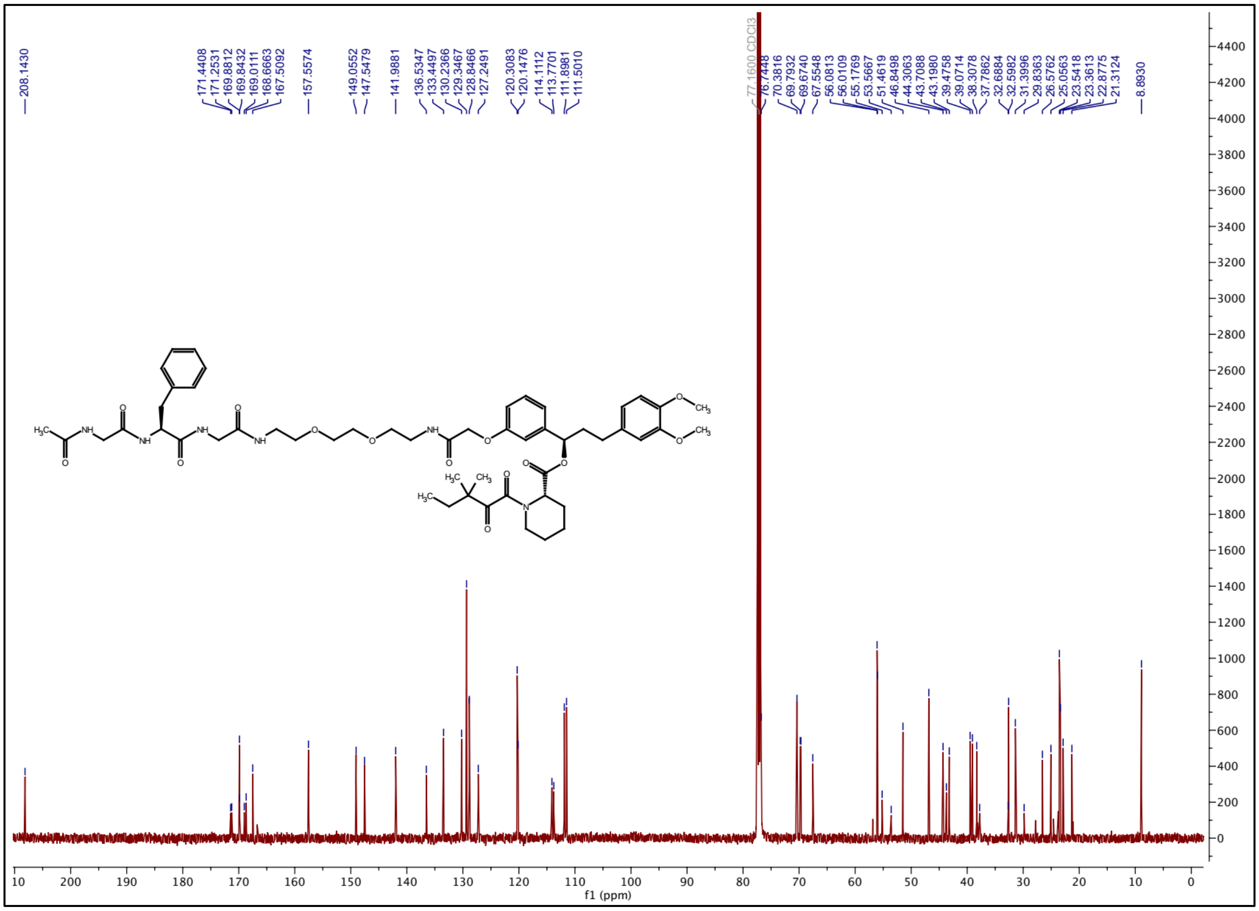

### $^{1}H$ NMR spectrum of ZA2

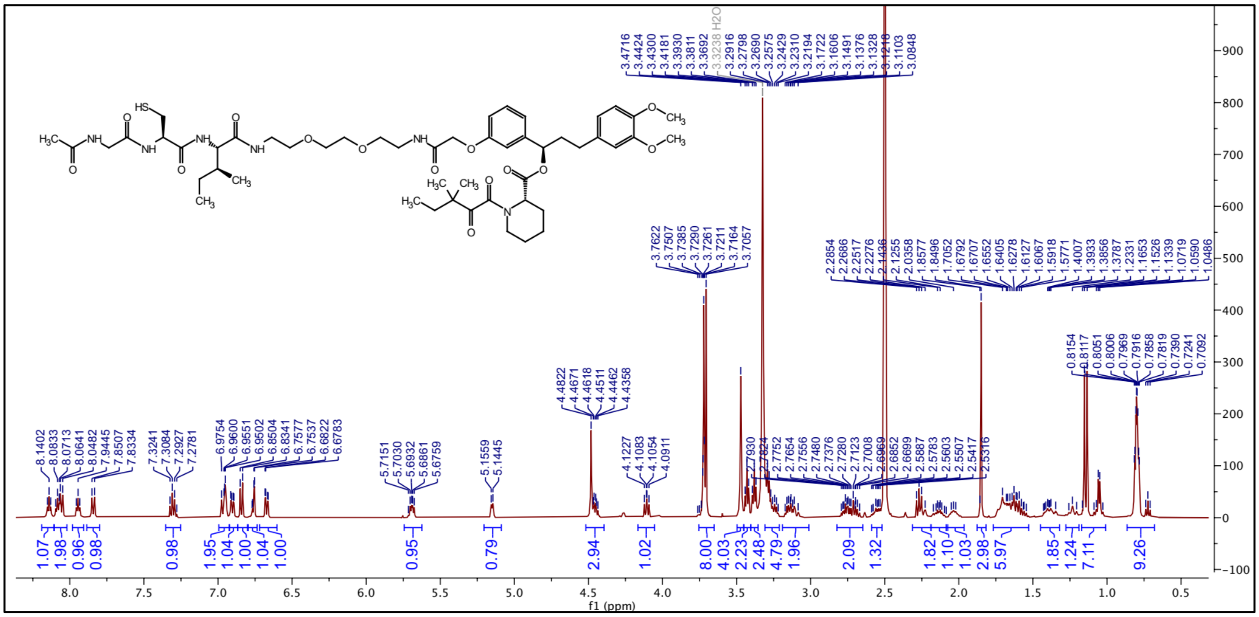

### $^{13}C$ NMR spectrum of ZA2

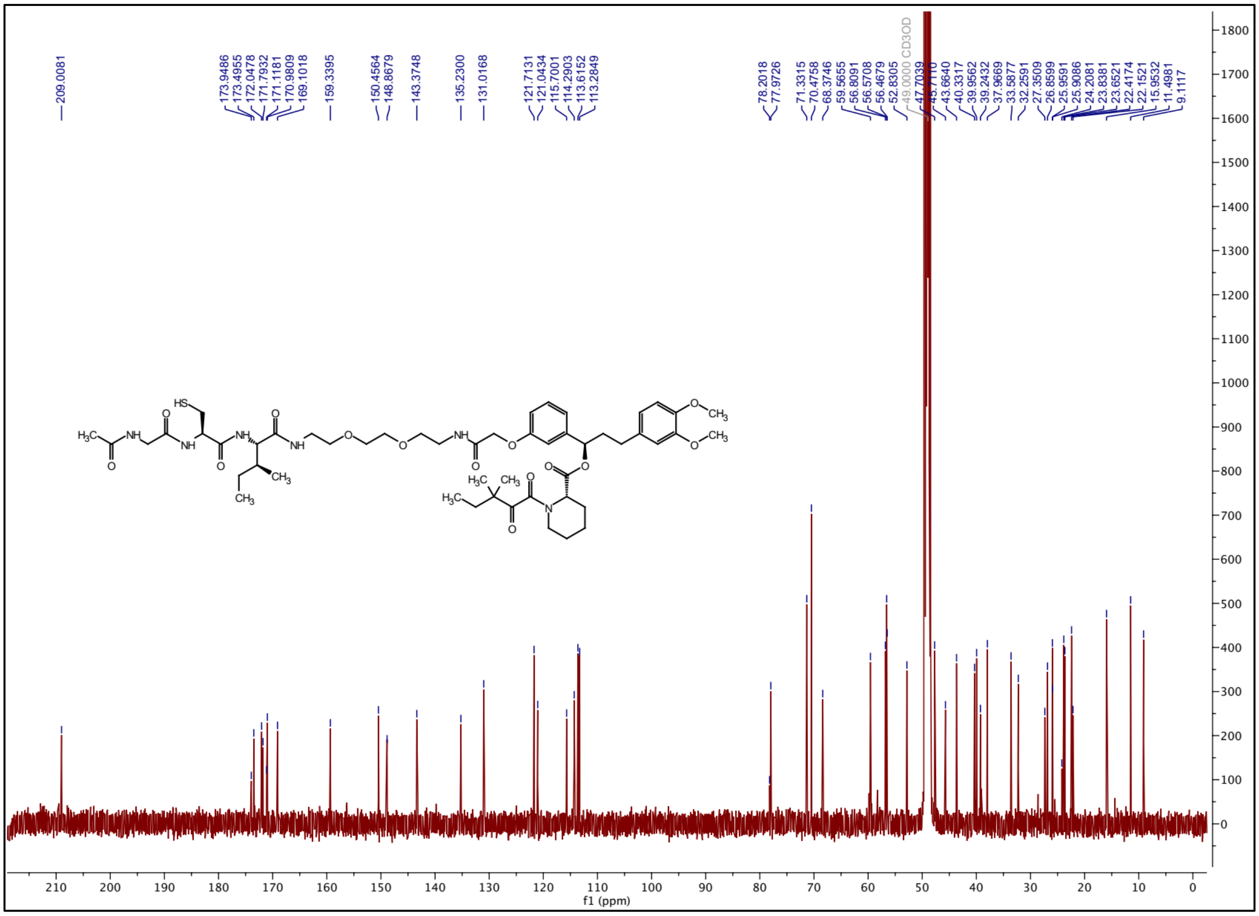

### $^{1}$H NMR spectrum of ZA3

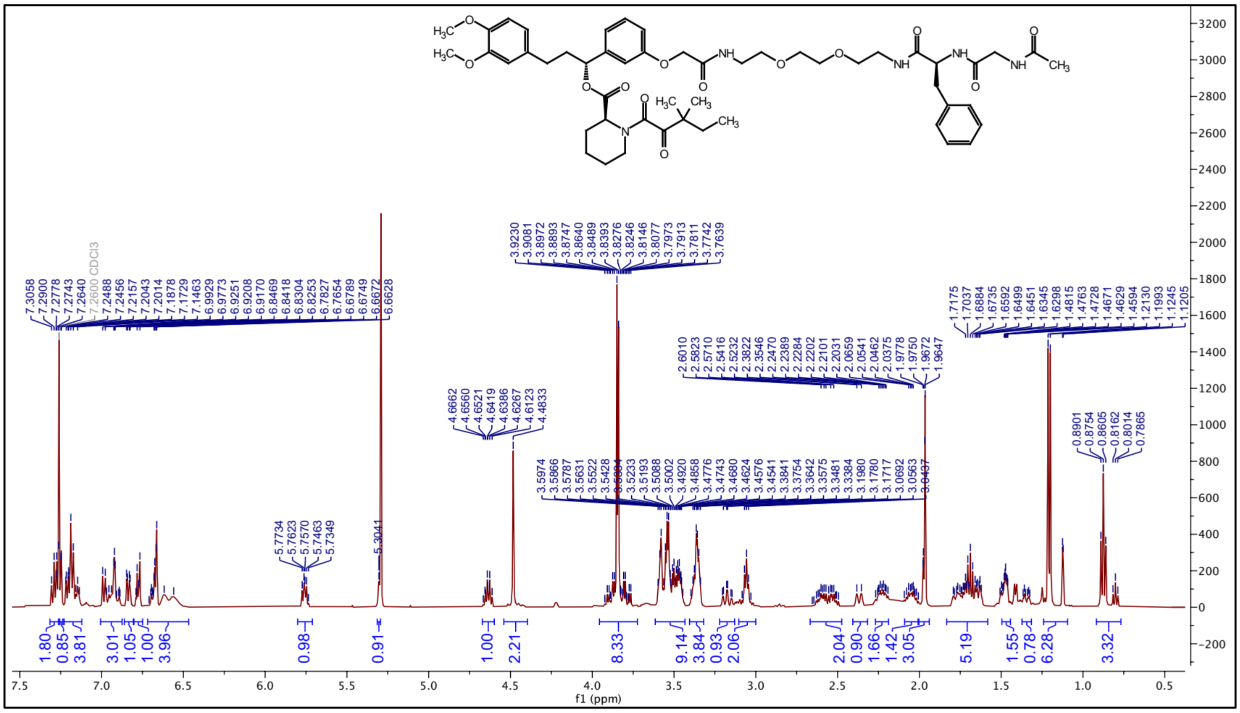

### $^{13}$C NMR spectrum of ZA3

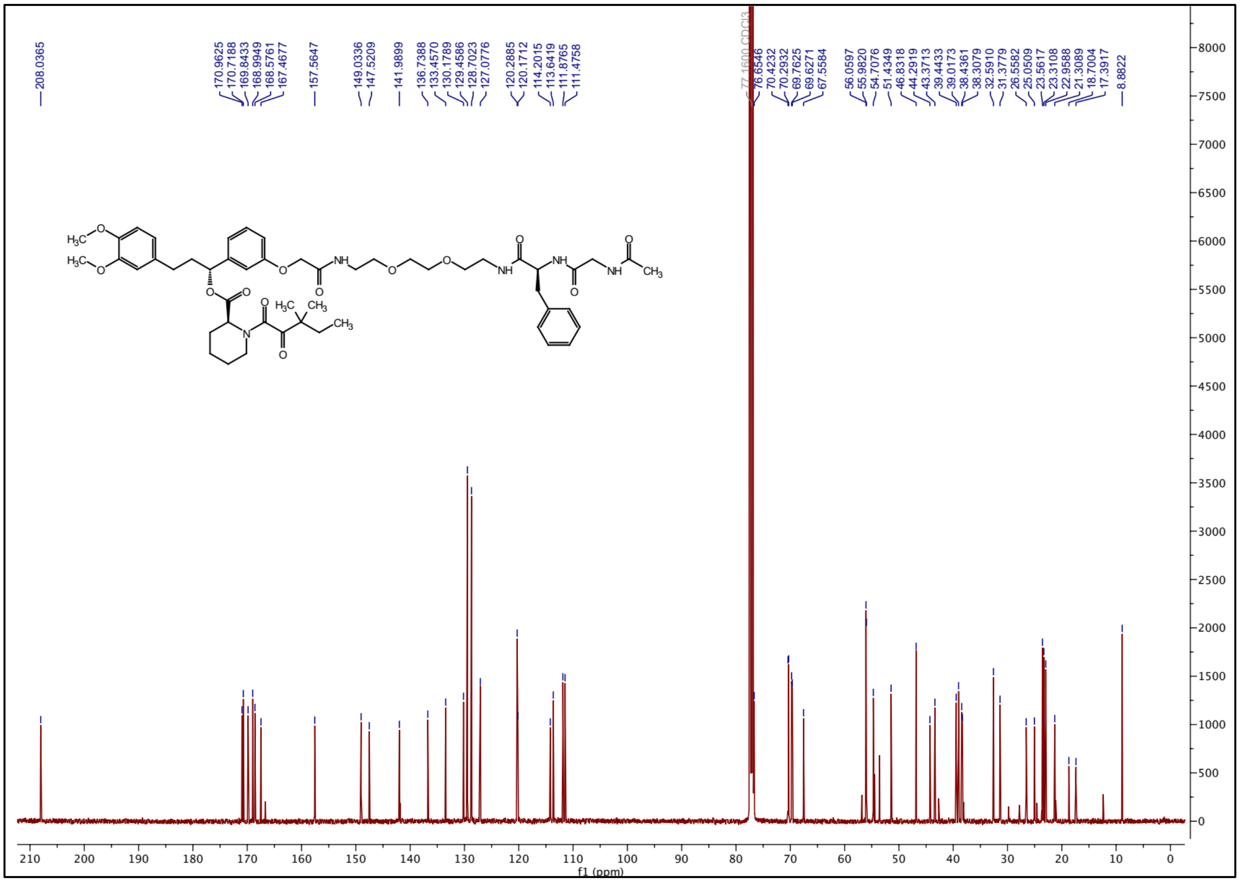

### $^{1}H$ NMR spectrum of ZA4

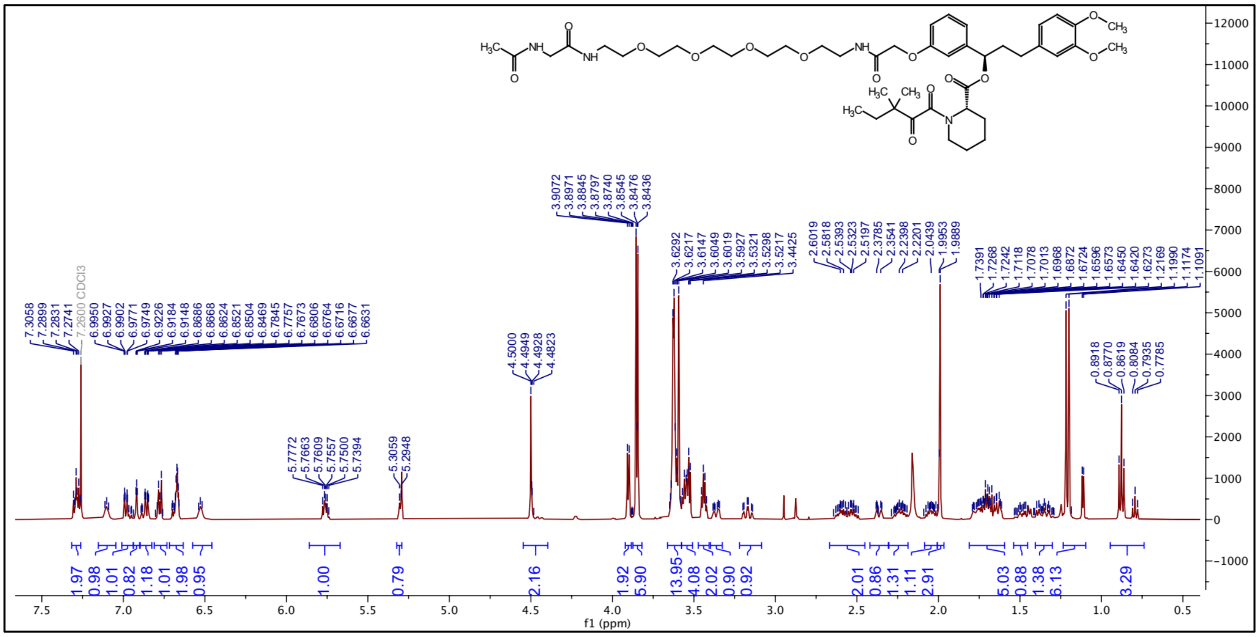

### $^{13}C$ NMR spectrum of ZA4

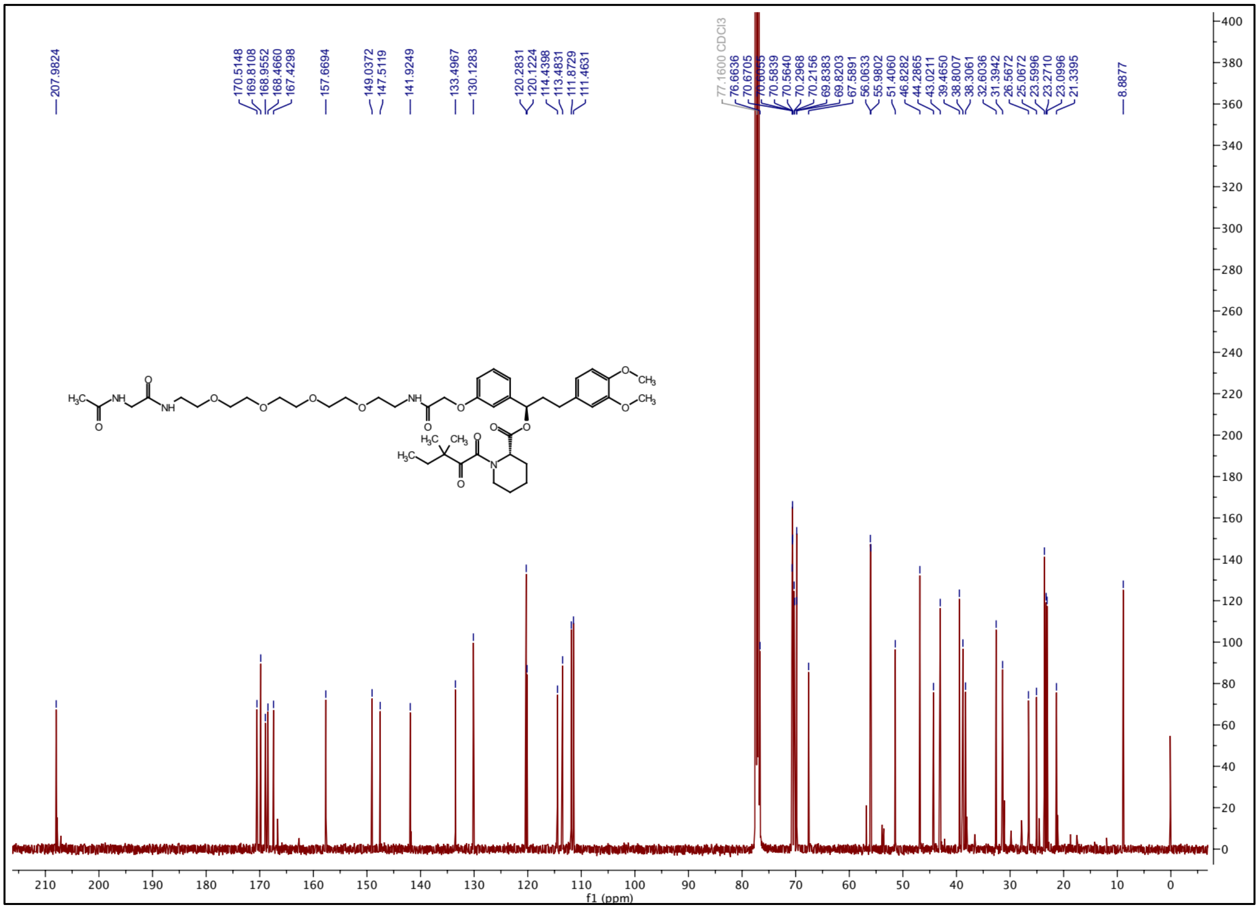

### $^{1}H$ NMR spectrum of ZA5

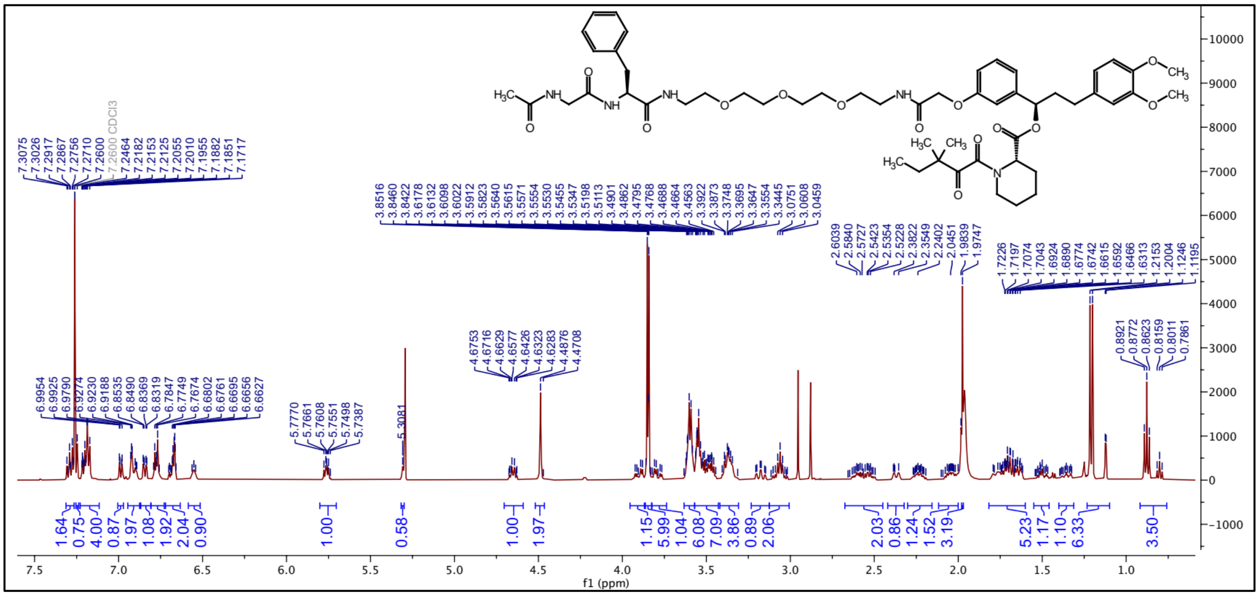

### $^{13}C$ NMR spectrum of ZA5

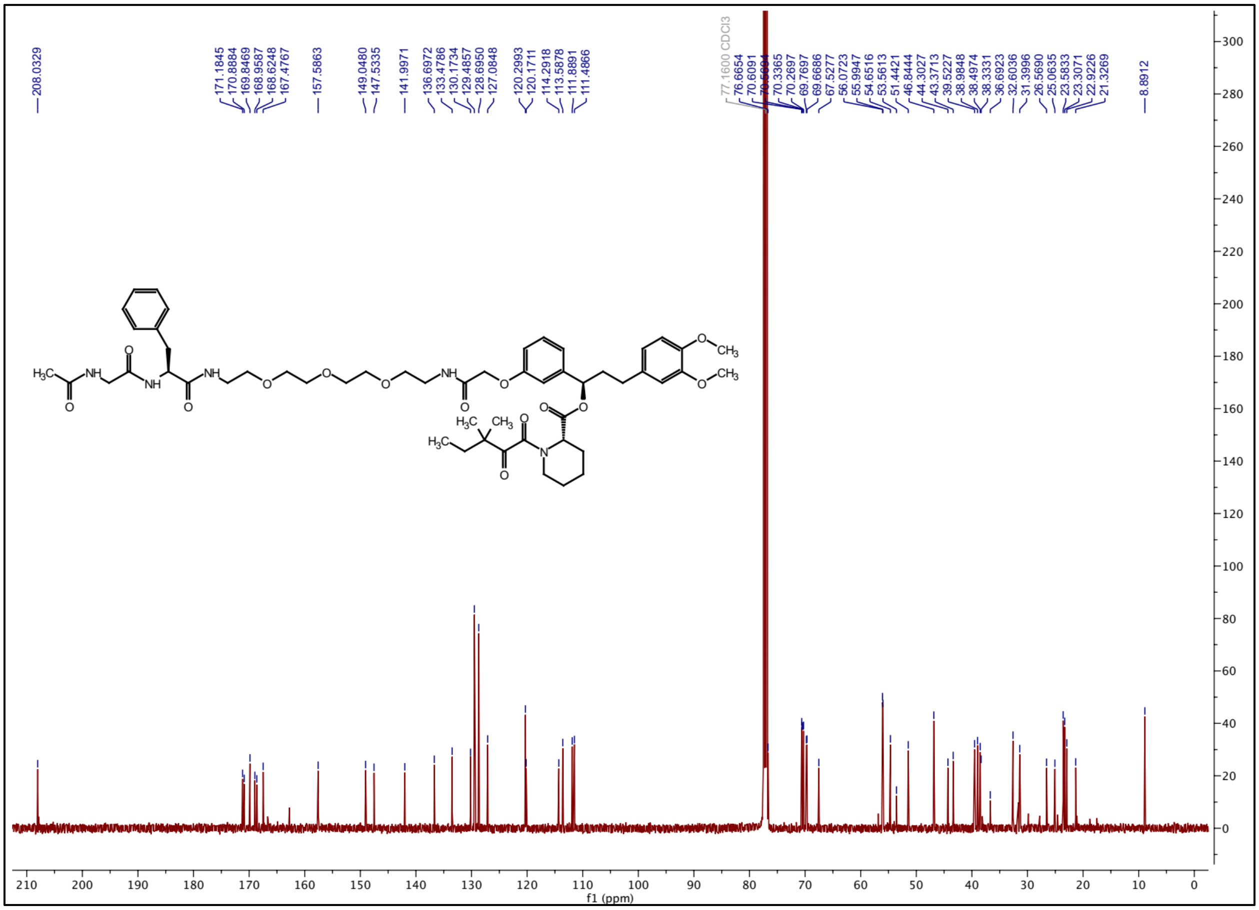

### $^{1}$H NMR spectrum of ZA6

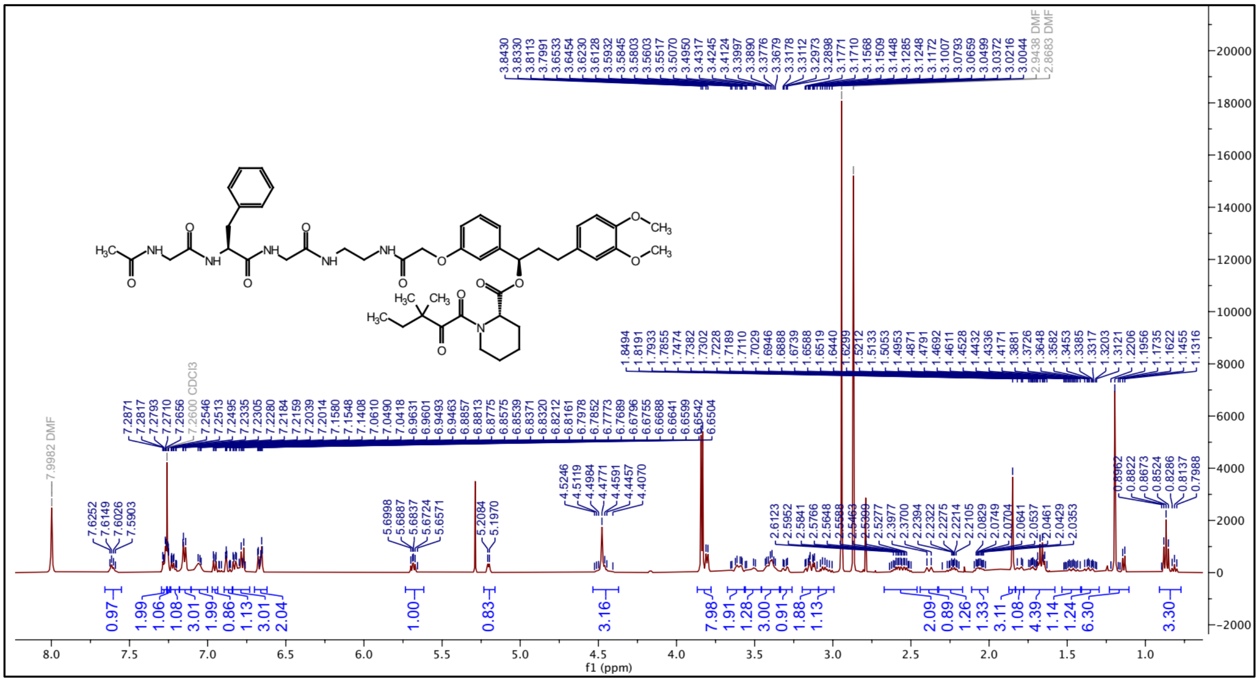


### $^{13}$C NMR spectrum of ZA6

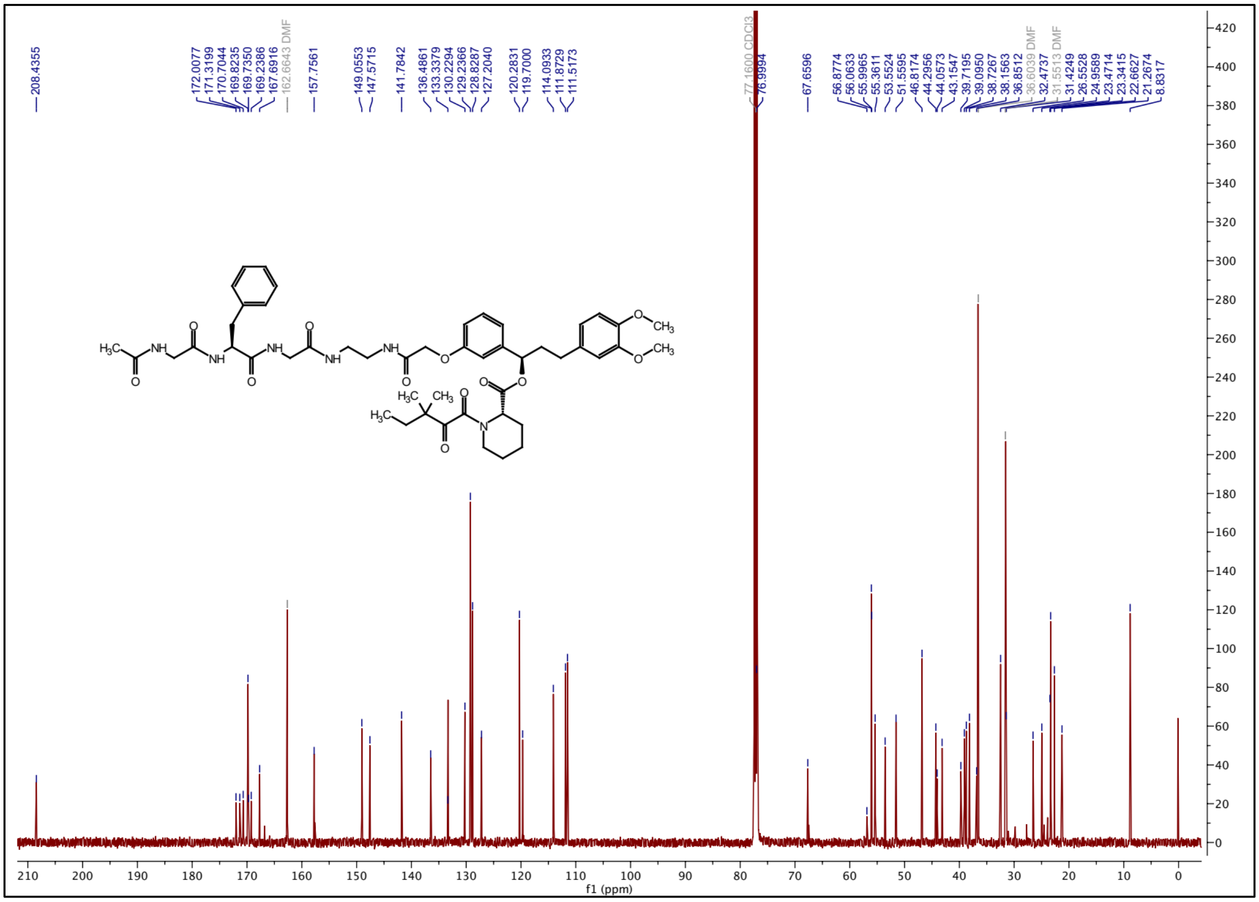

### $^{1}$H NMR spectrum of ZA7

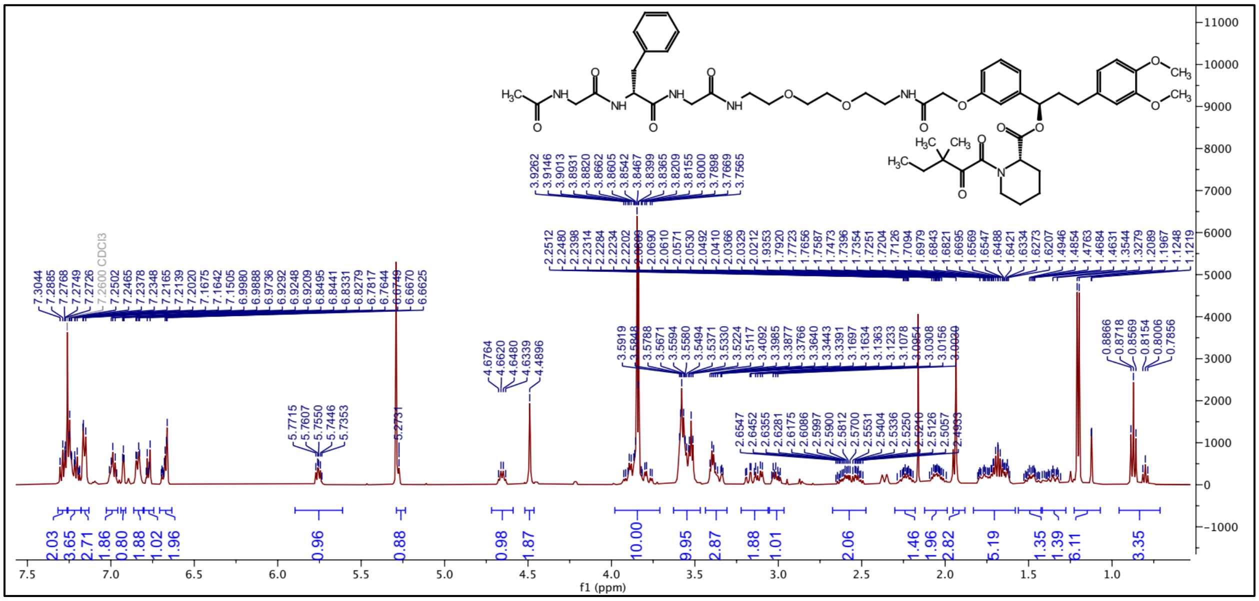


### $^{13}$C NMR spectrum of ZA7

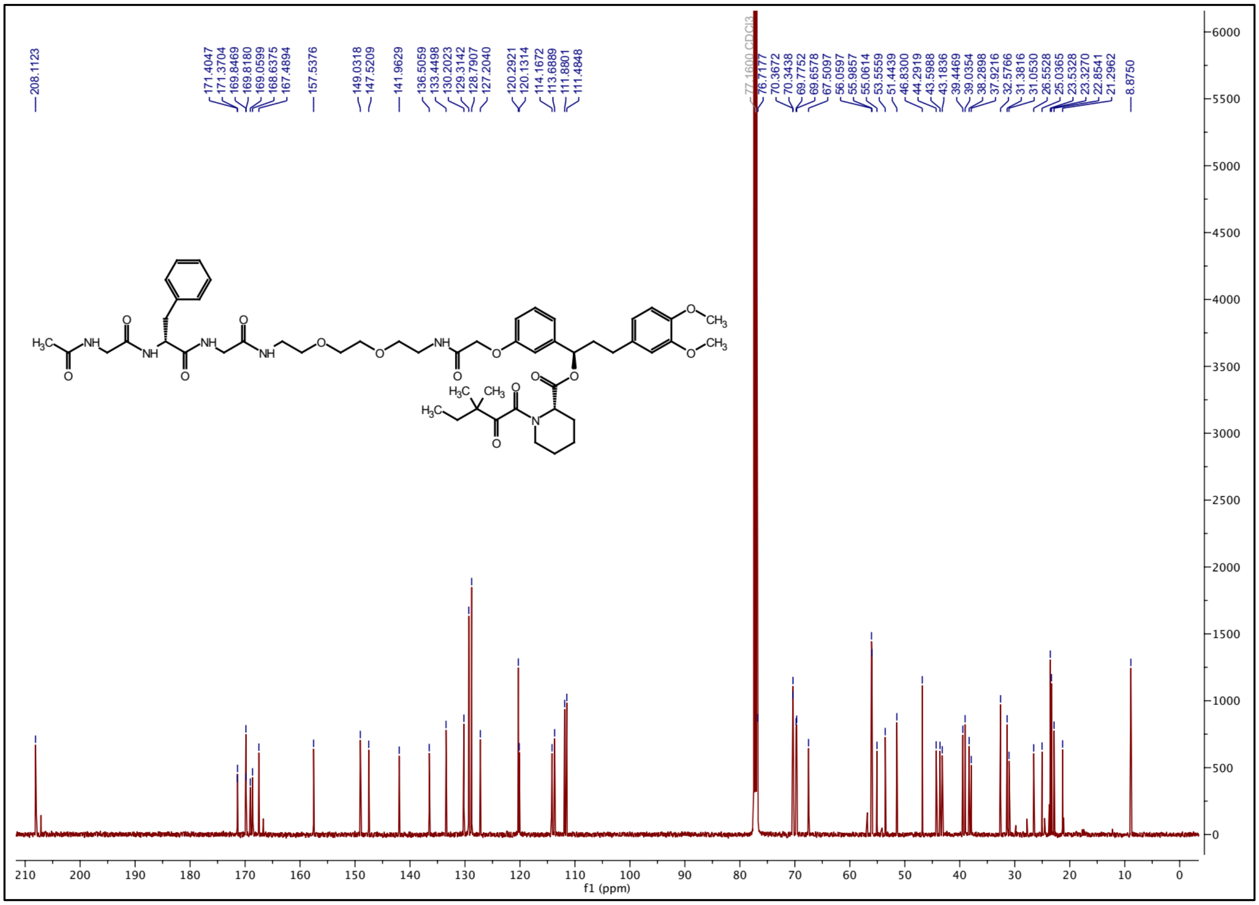

### $^1H$ NMR spectrum of ZA8

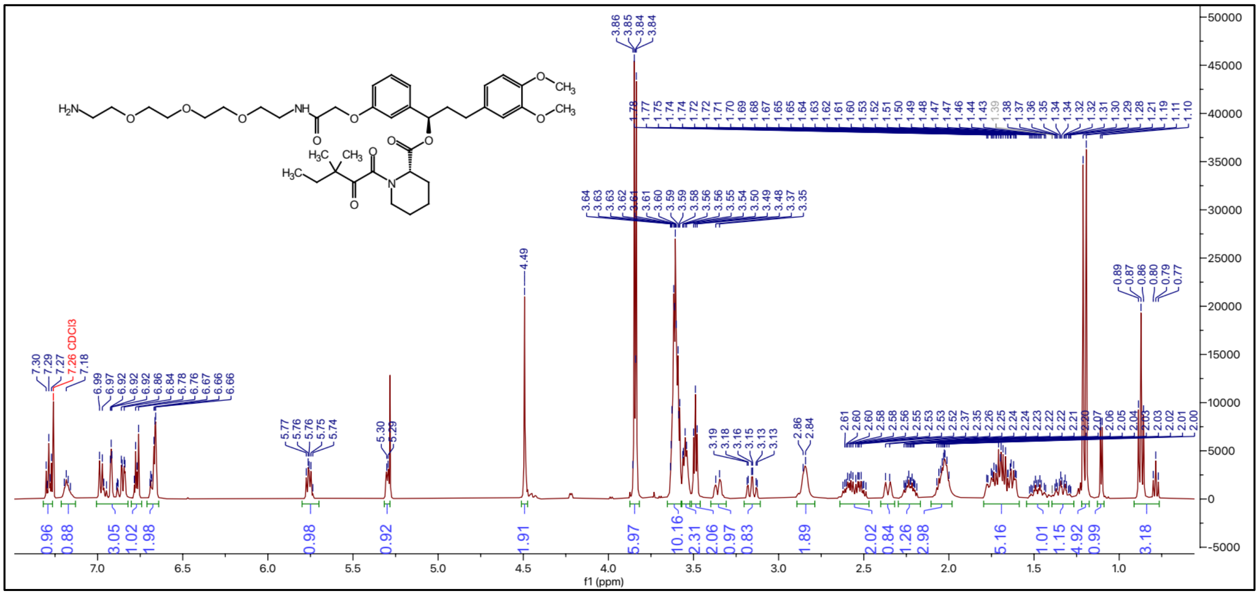

### $^{13}C$ NMR spectrum of ZA8

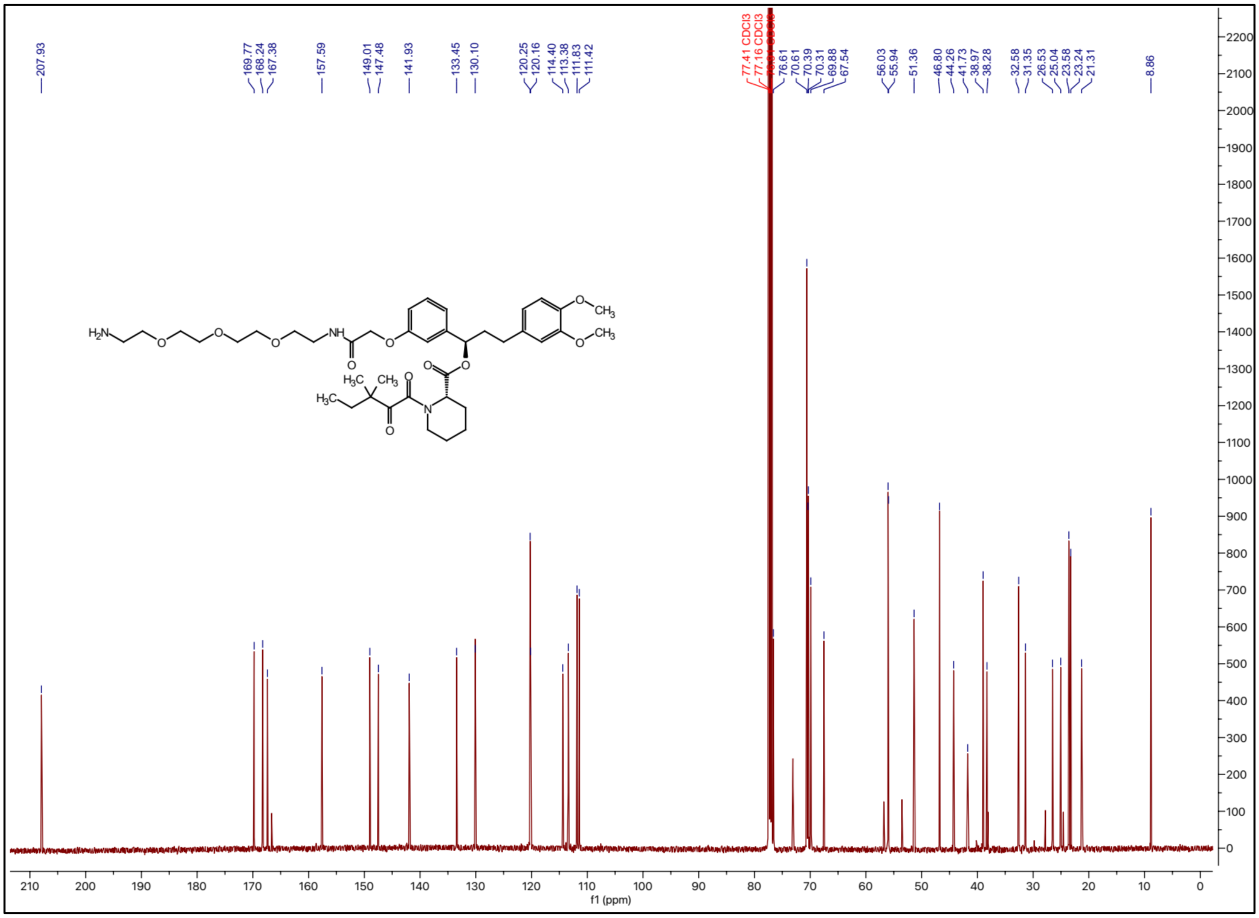

### [1]H NMR spectrum of ZA9

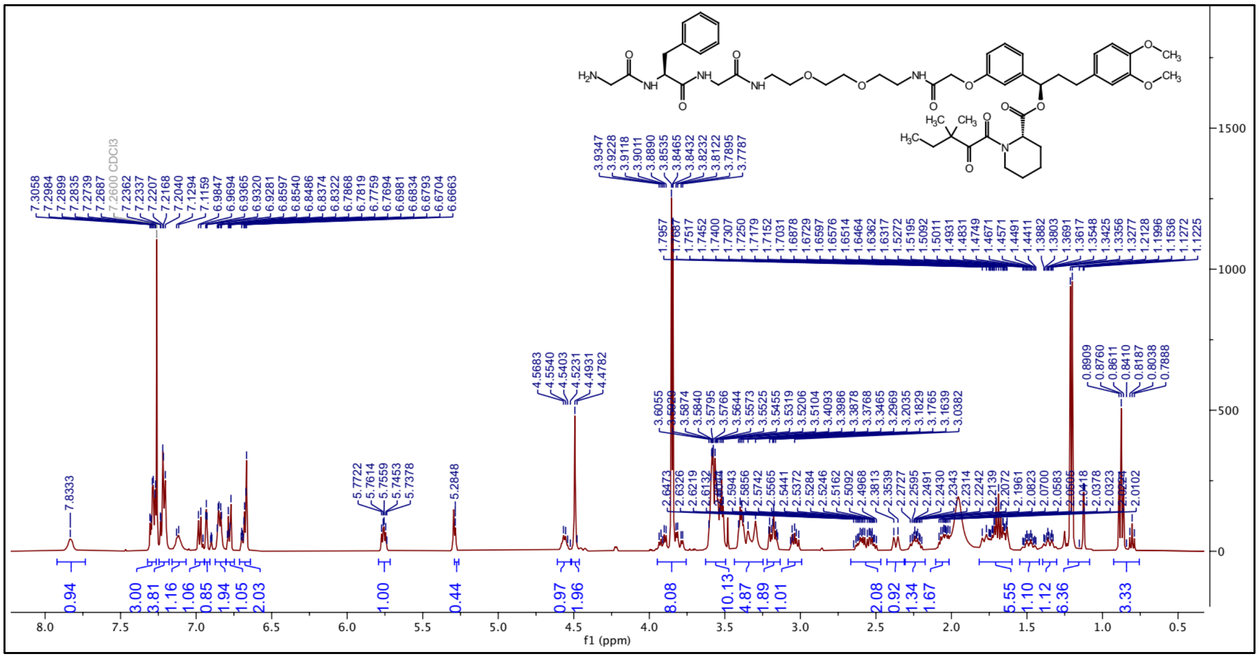

### [13]C NMR spectrum of ZA9

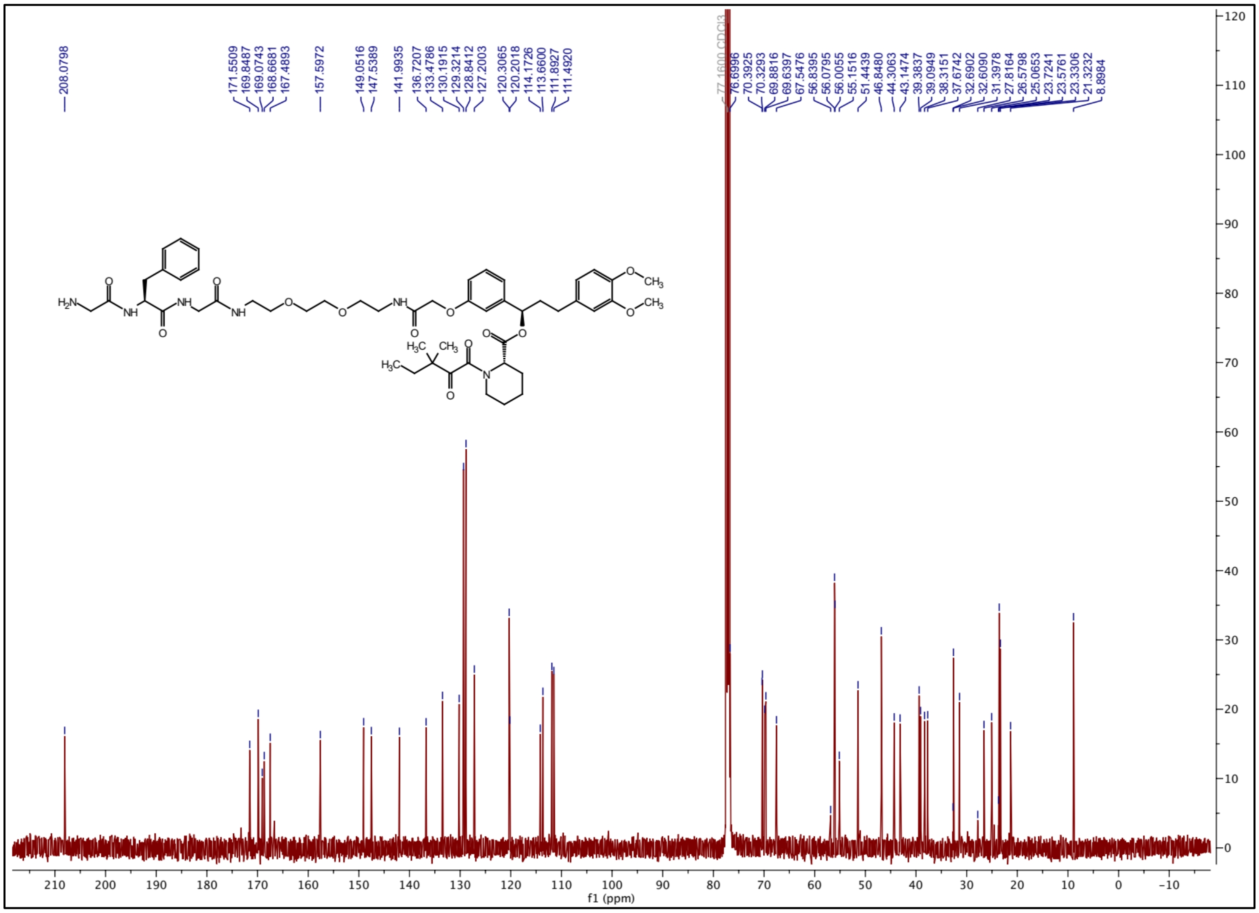

### $^{1}H$ NMR spectrum of ZA10

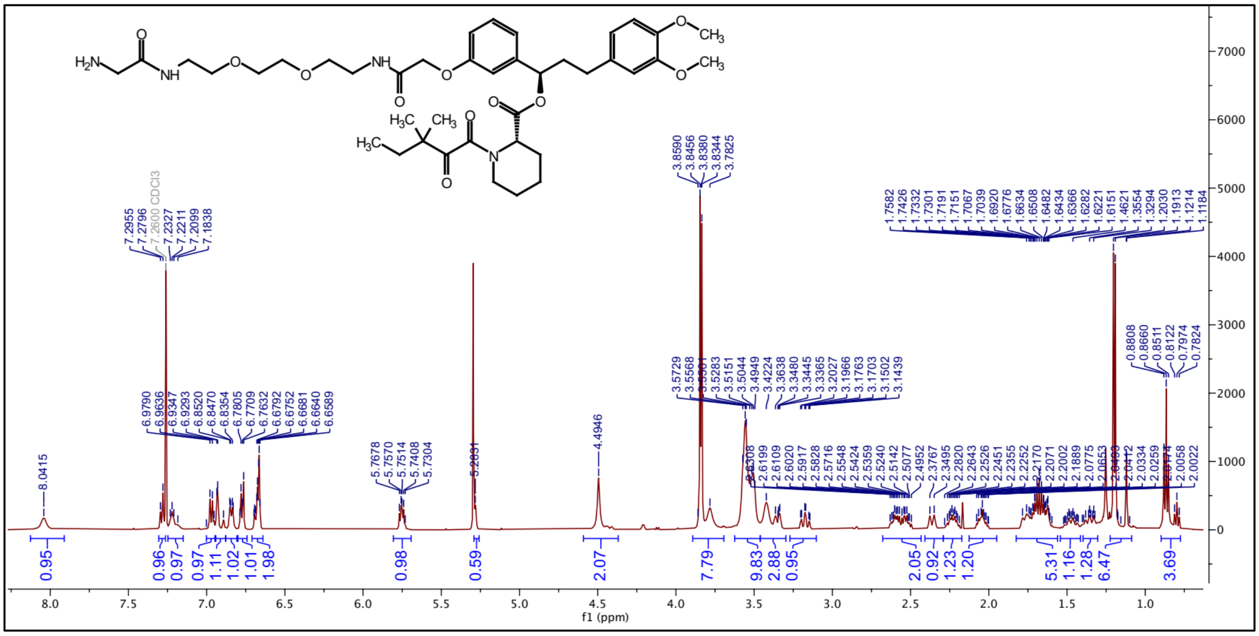

### $^{13}C$ NMR spectrum of ZA10

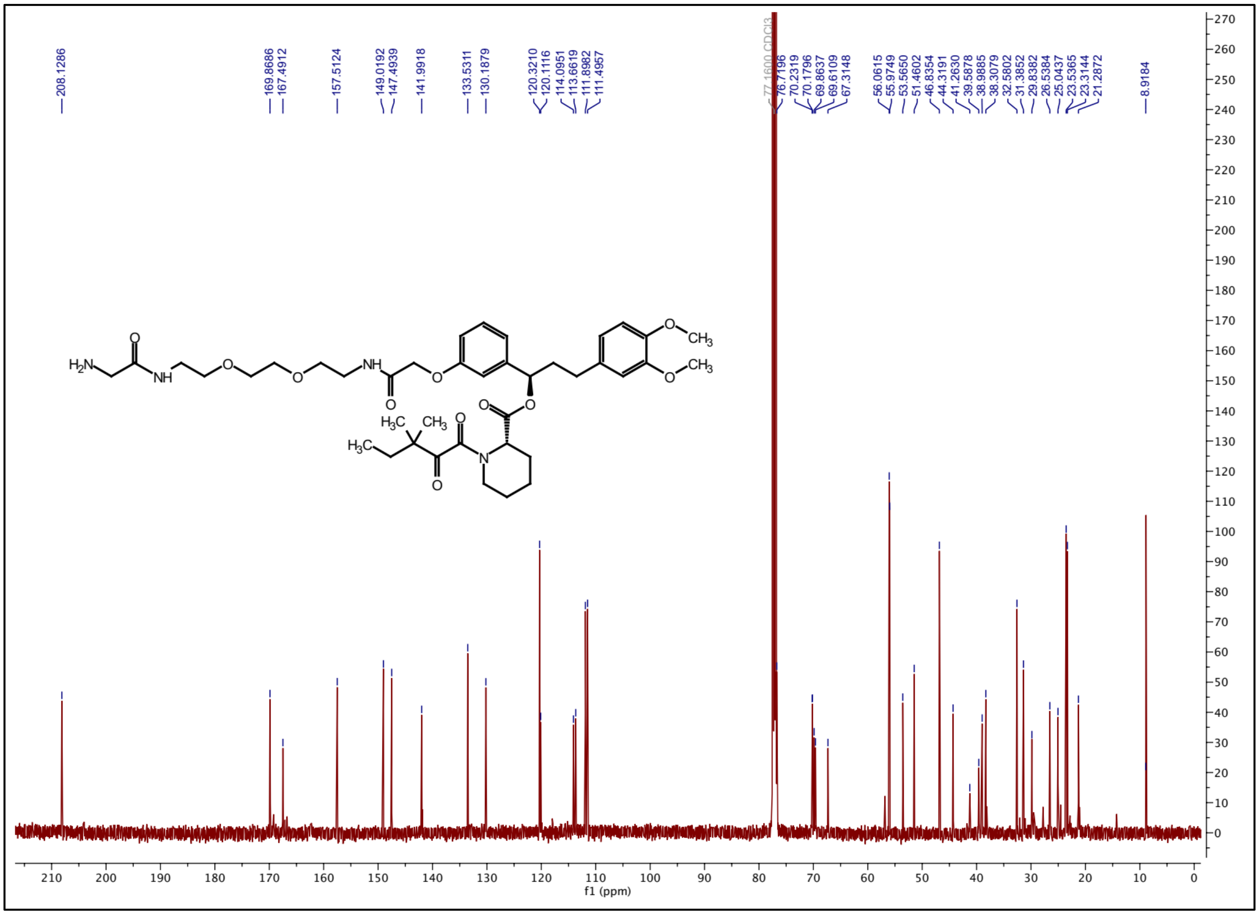

### [1]H NMR spectrum of ZA11

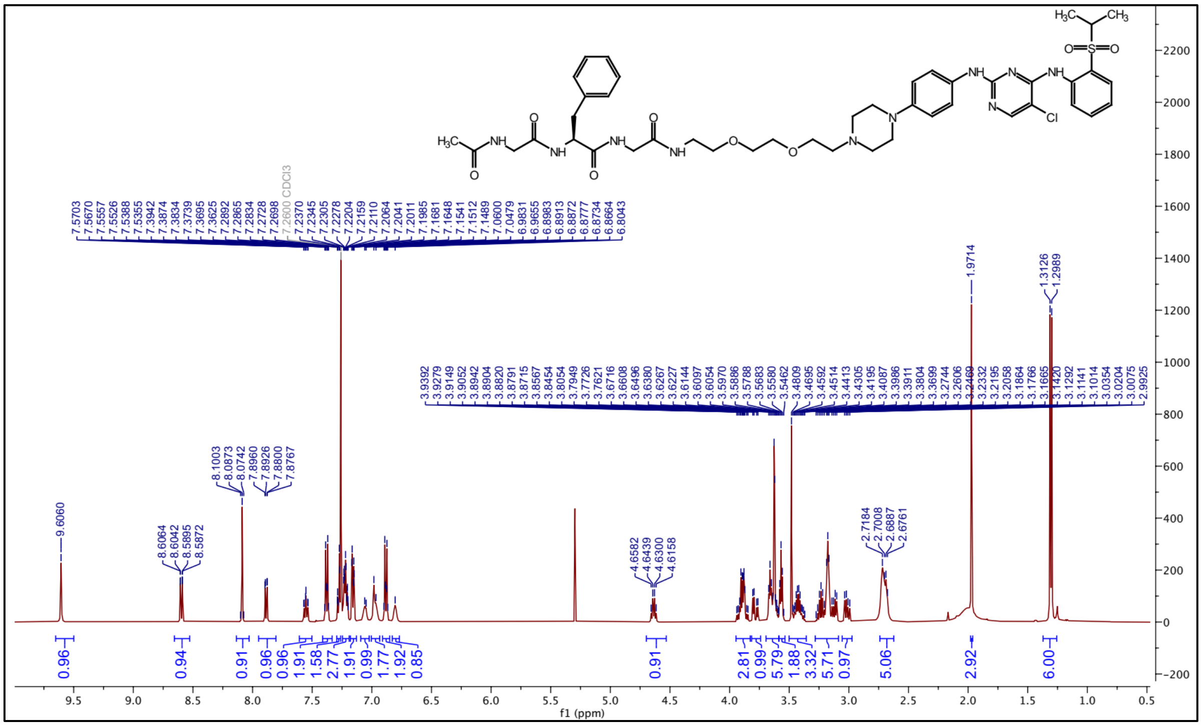

### [13]C NMR spectrum of ZA11

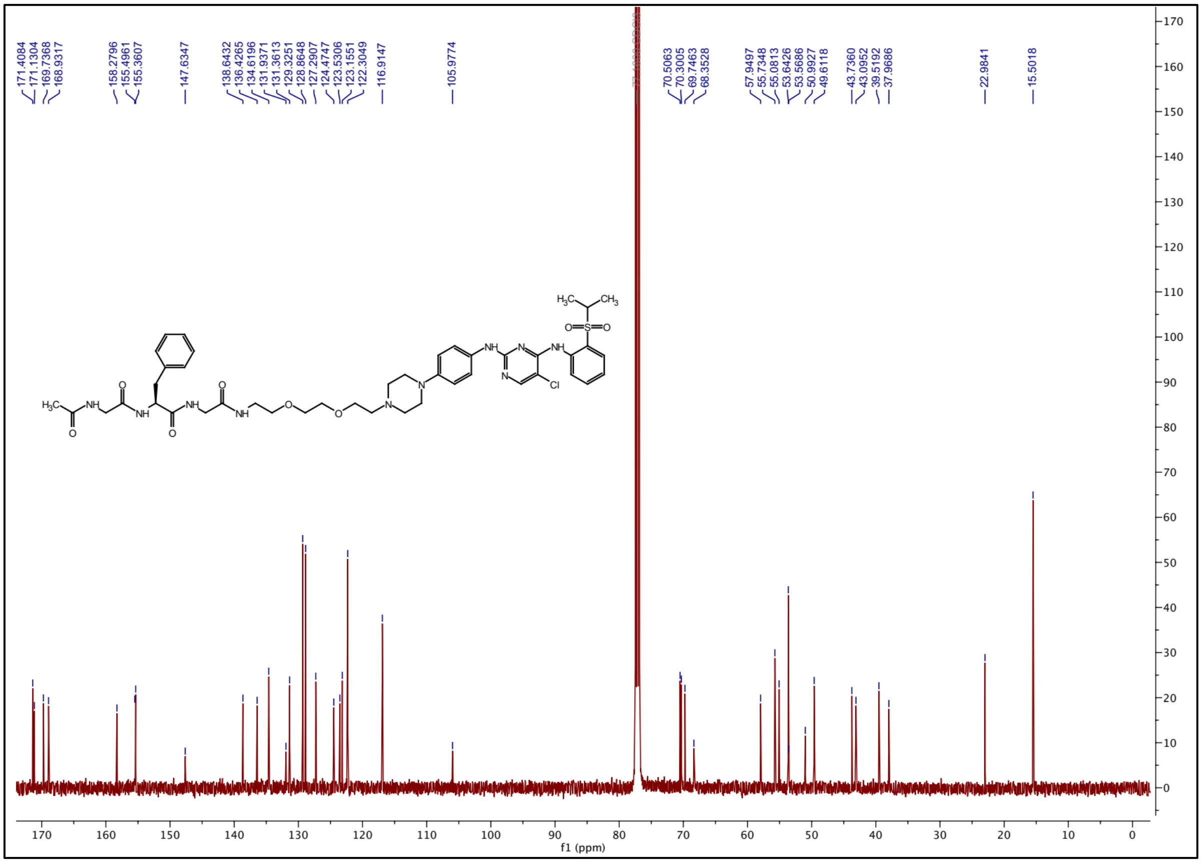

### $^{1}H$ NMR spectrum of ZA12

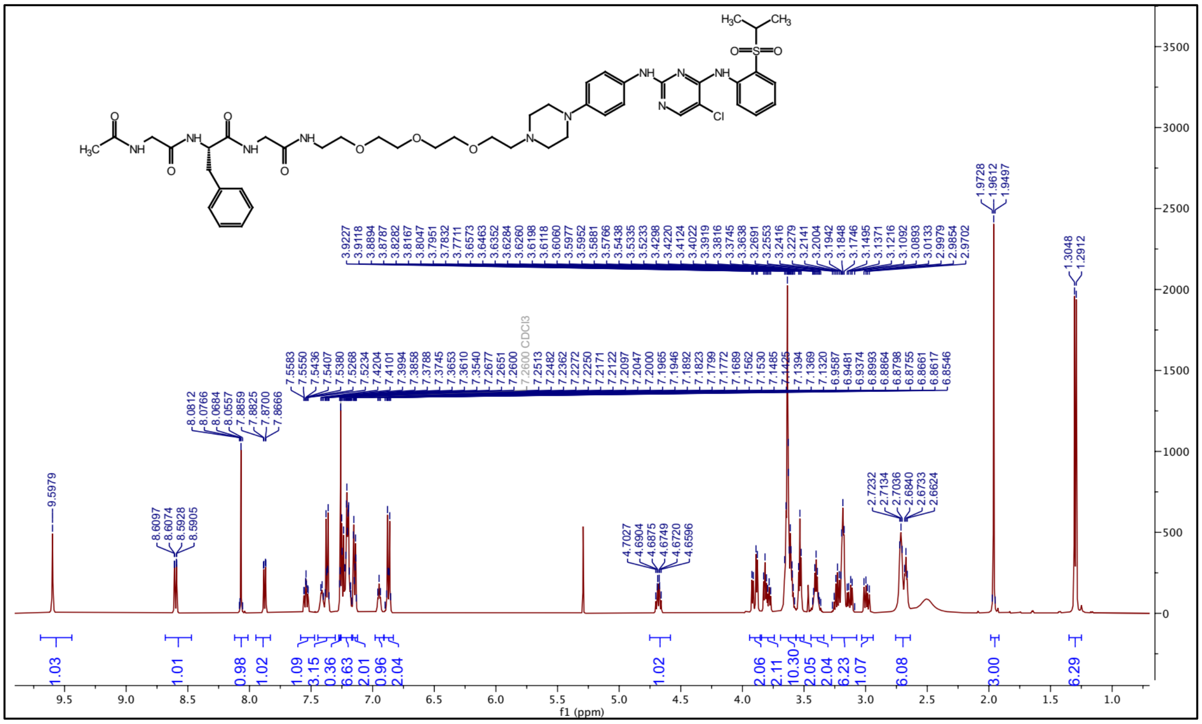

### $^{13}C$ NMR spectrum of ZA12

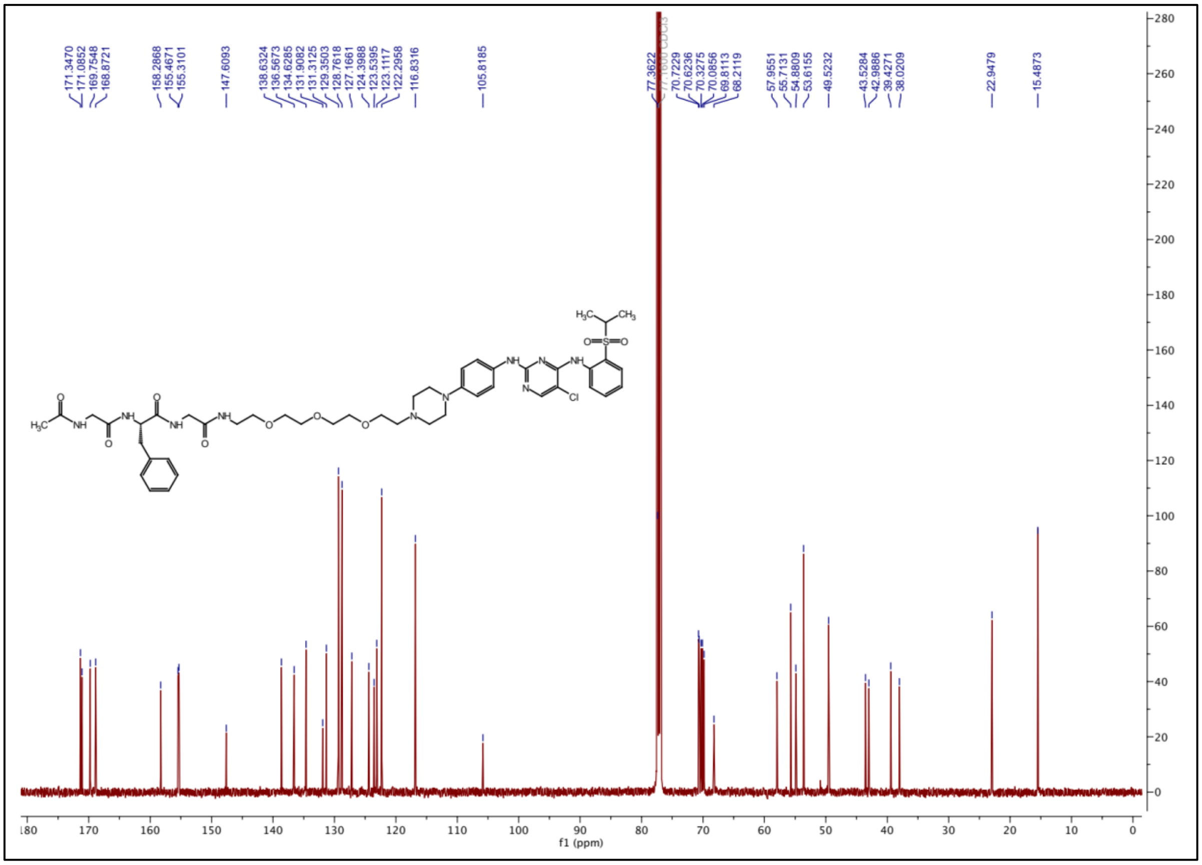

### $^1$H NMR spectrum of ZA13

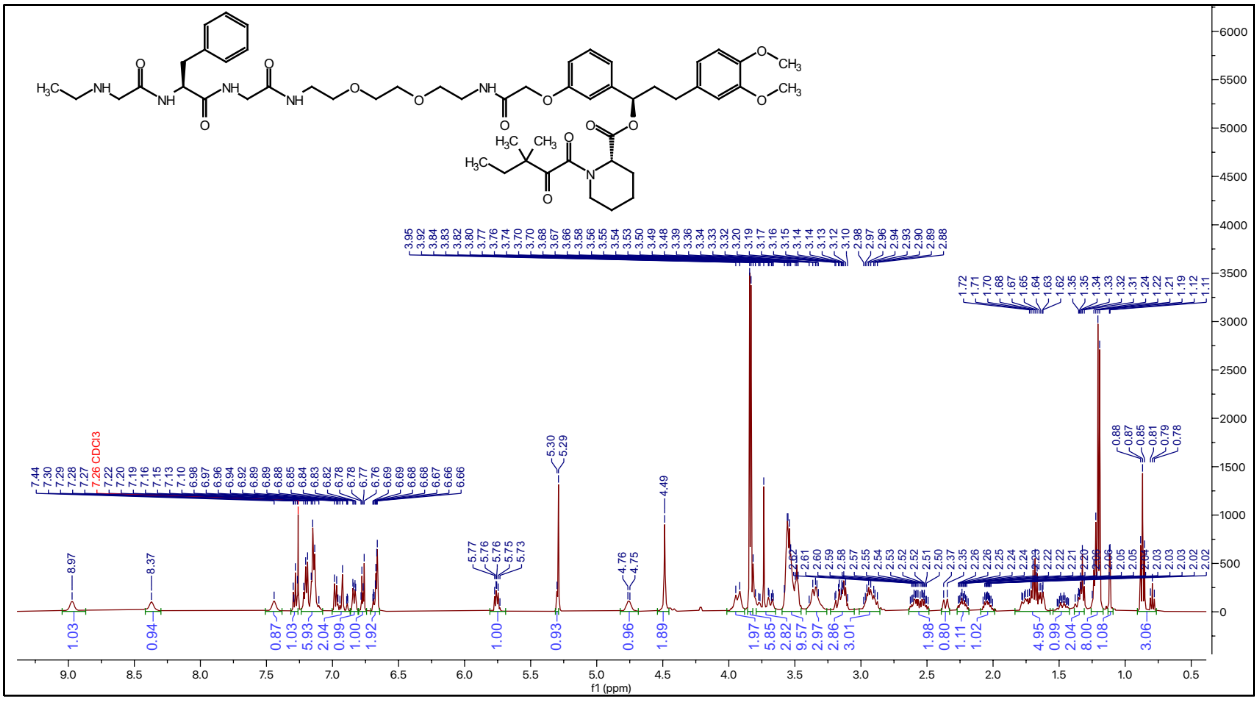

### $^{13}$C NMR spectrum of ZA13

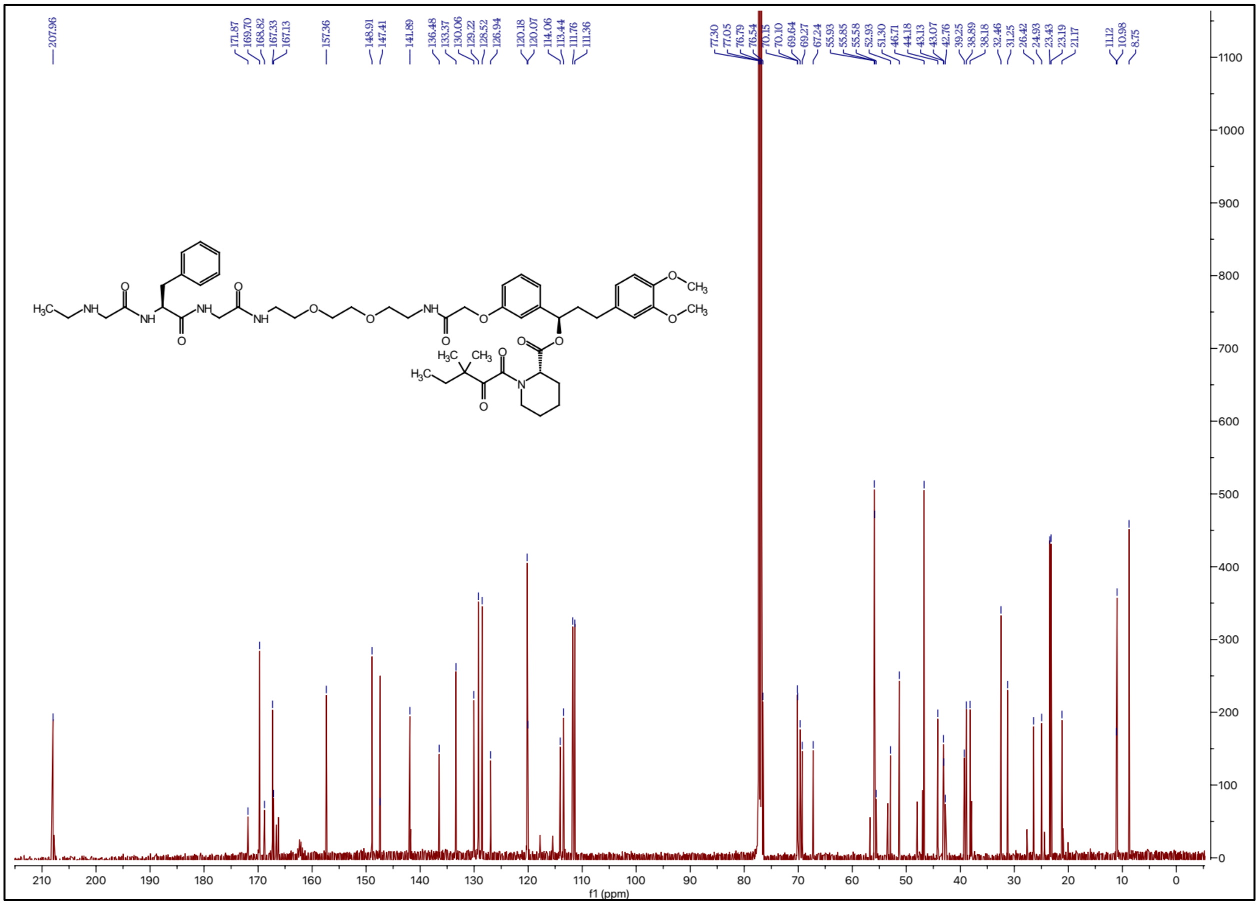

### $^{1}$H NMR spectrum of ZA14

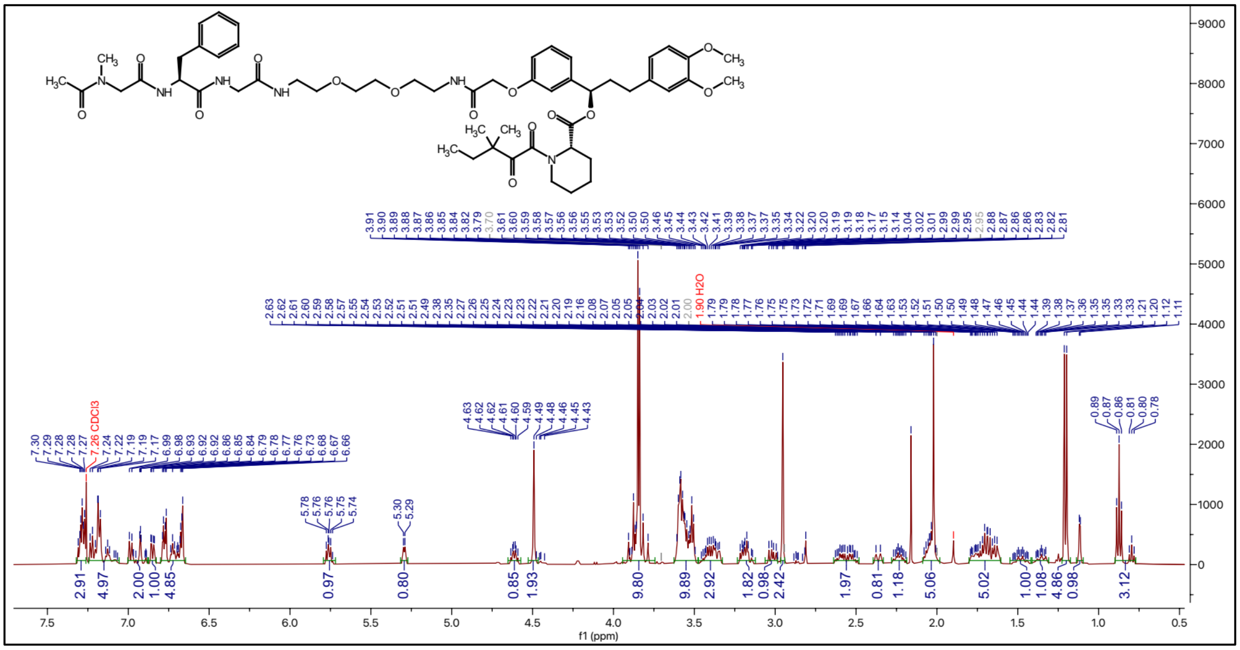

### $^{13}$C NMR spectrum of ZA14

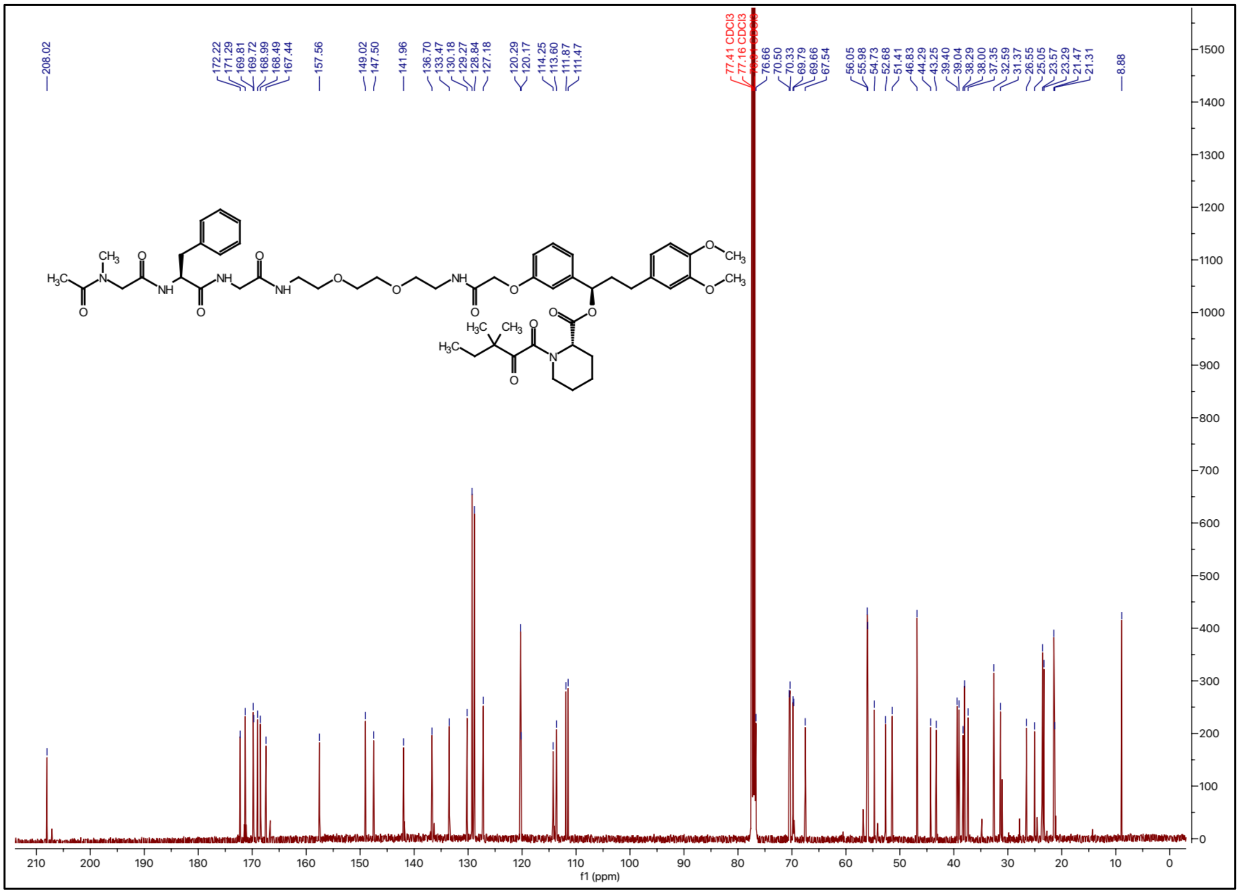

### $^{1}H$ NMR spectrum of ZA15

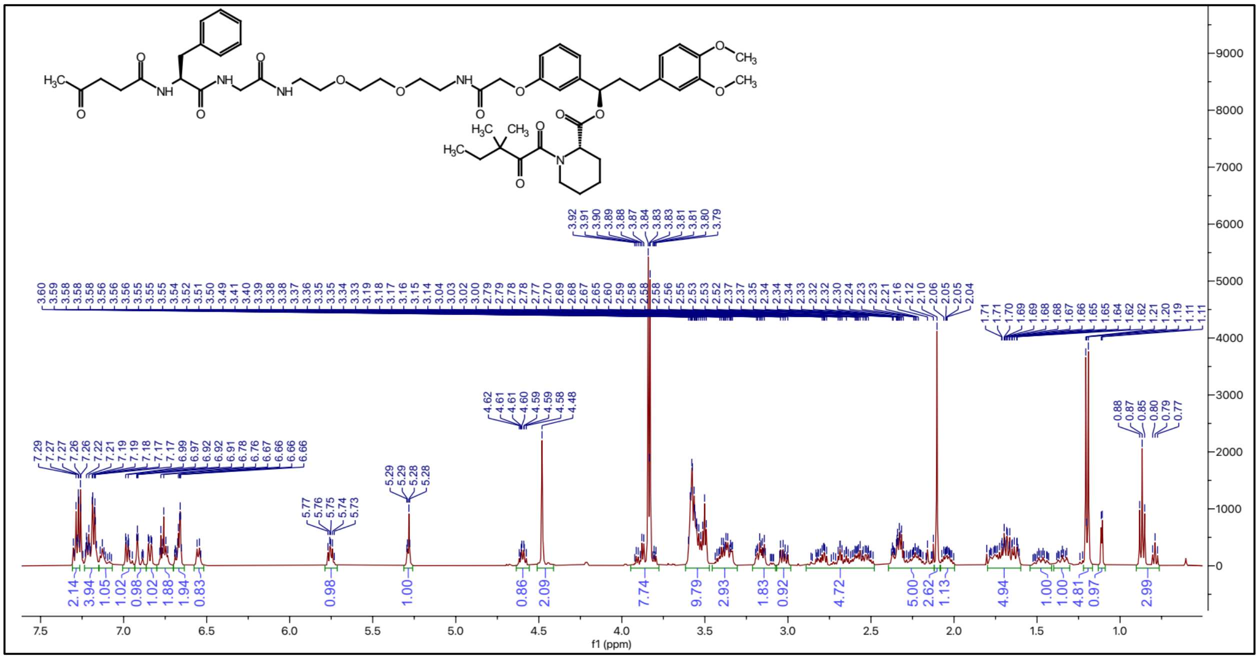

### $^{13}C$ NMR spectrum of ZA15

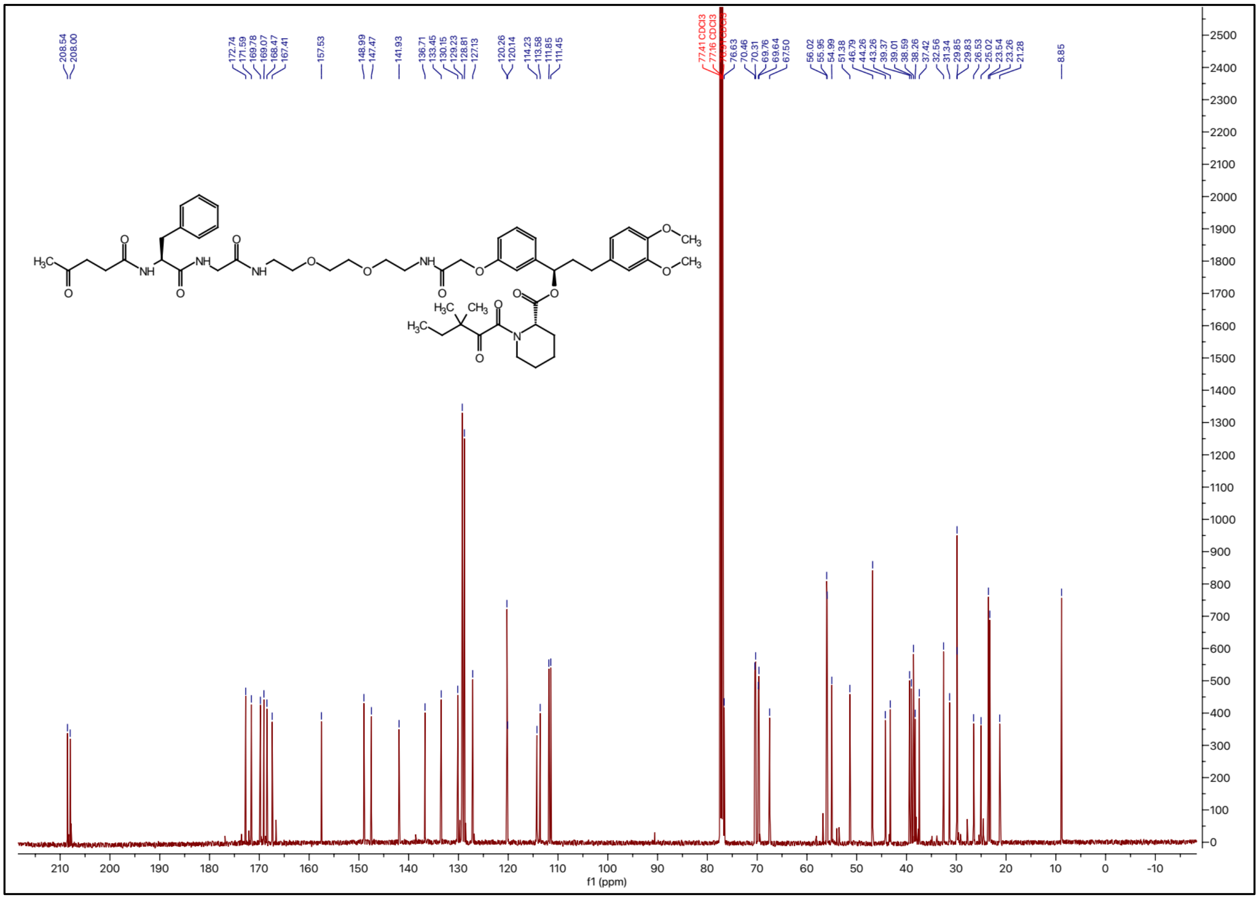

### $^{1}H$ NMR spectrum of ZA16

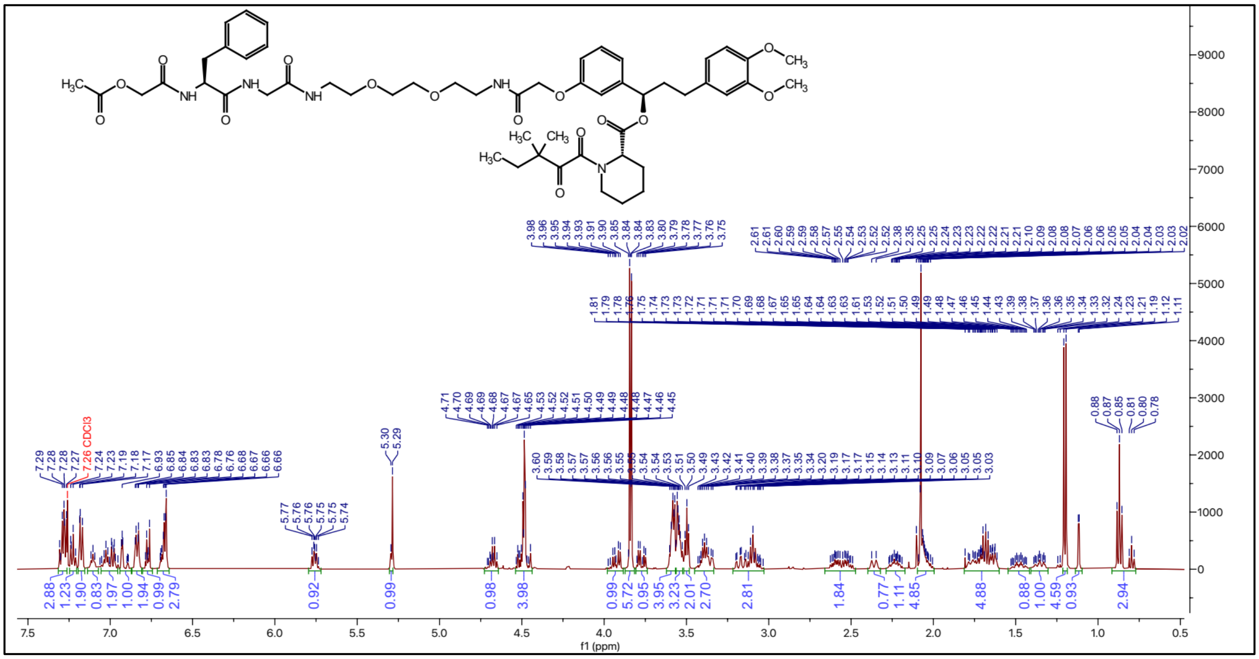

### $^{13}C$ NMR spectrum of ZA16

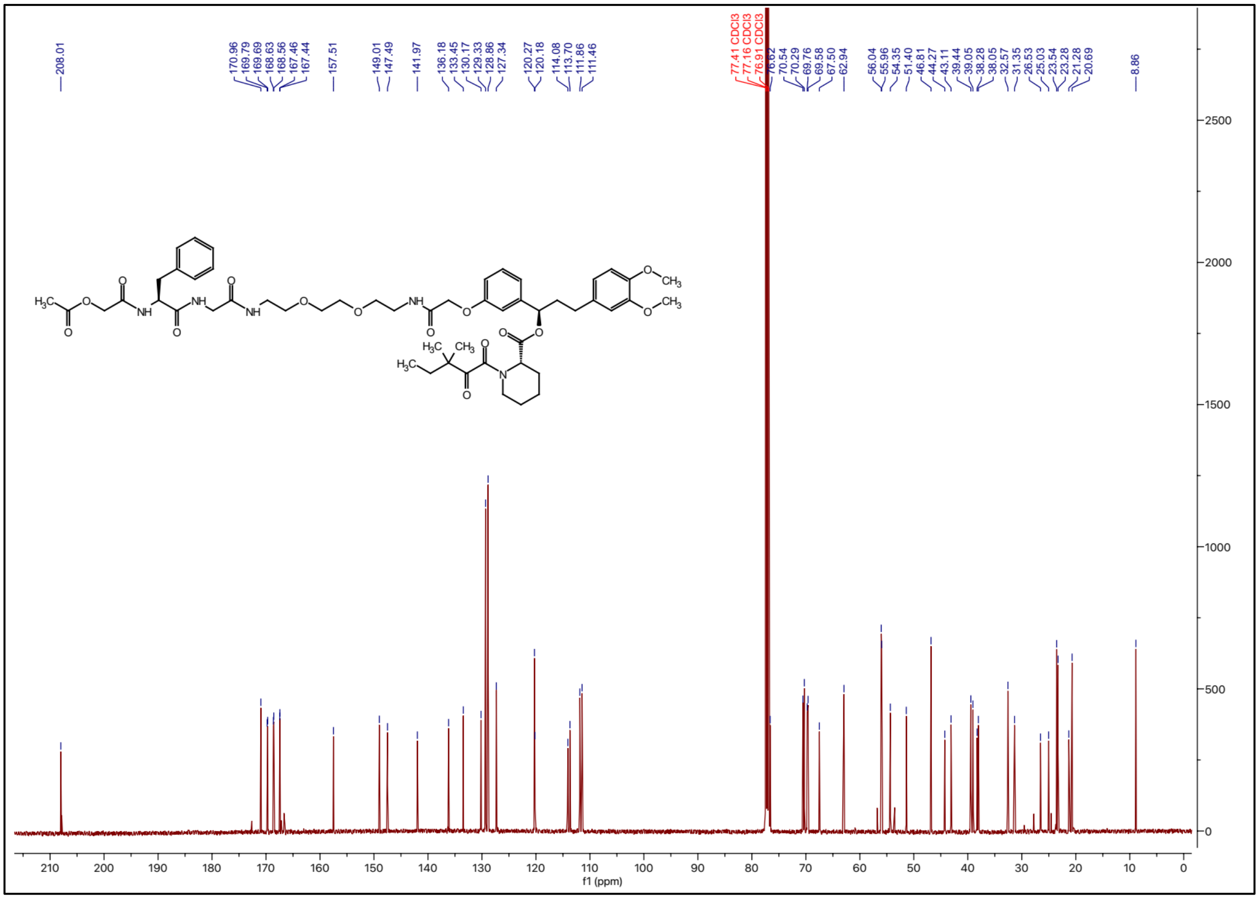

### $^{1}$H NMR spectrum of ZA17

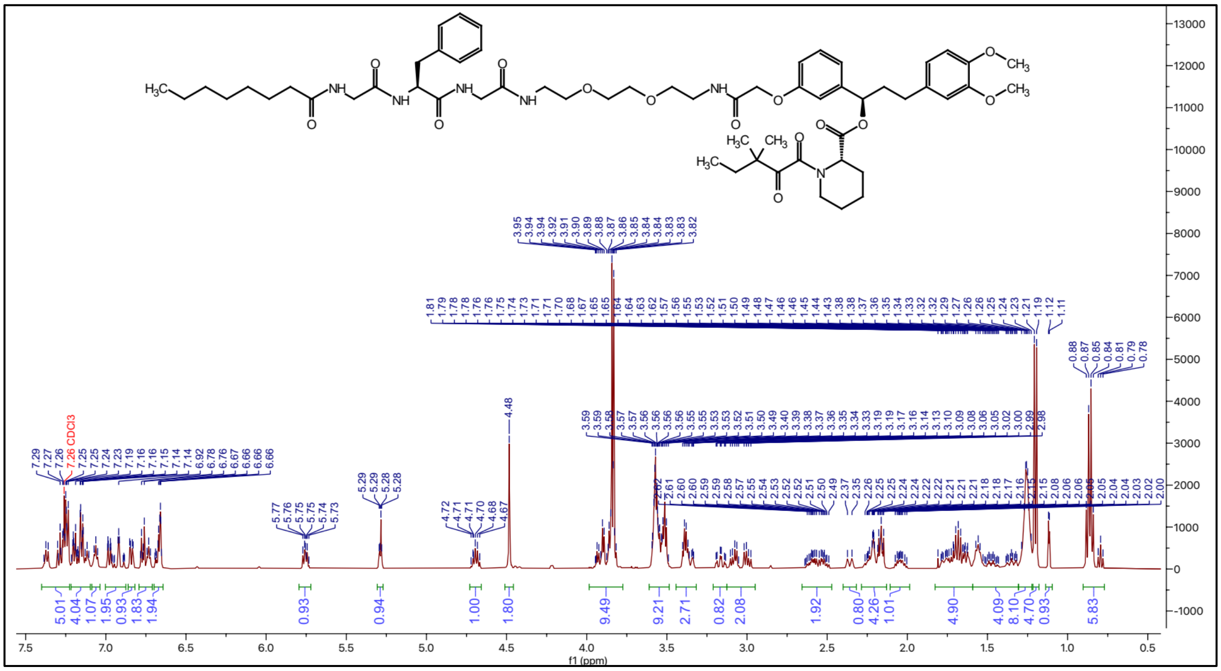

### $^{13}$C NMR spectrum of ZA17

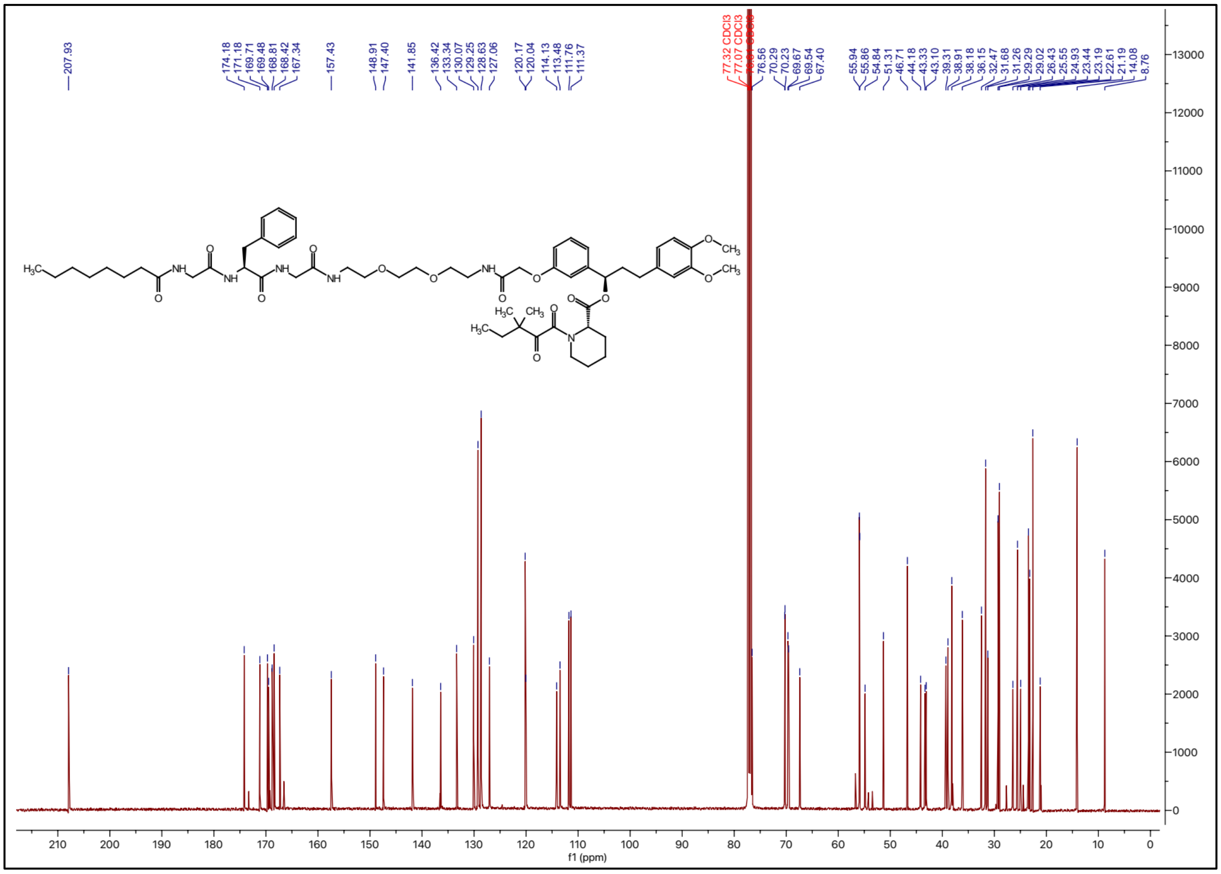

**$^{1}$H NMR spectrum of ZA18**

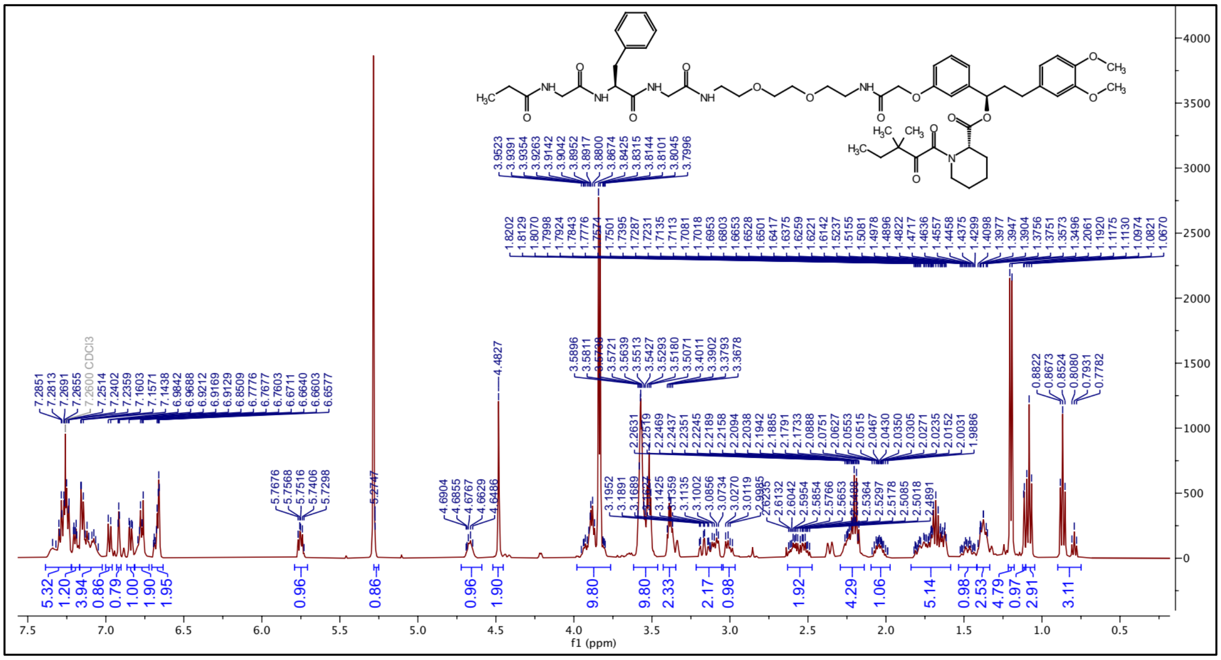

**$^{13}$C NMR spectrum of ZA18**

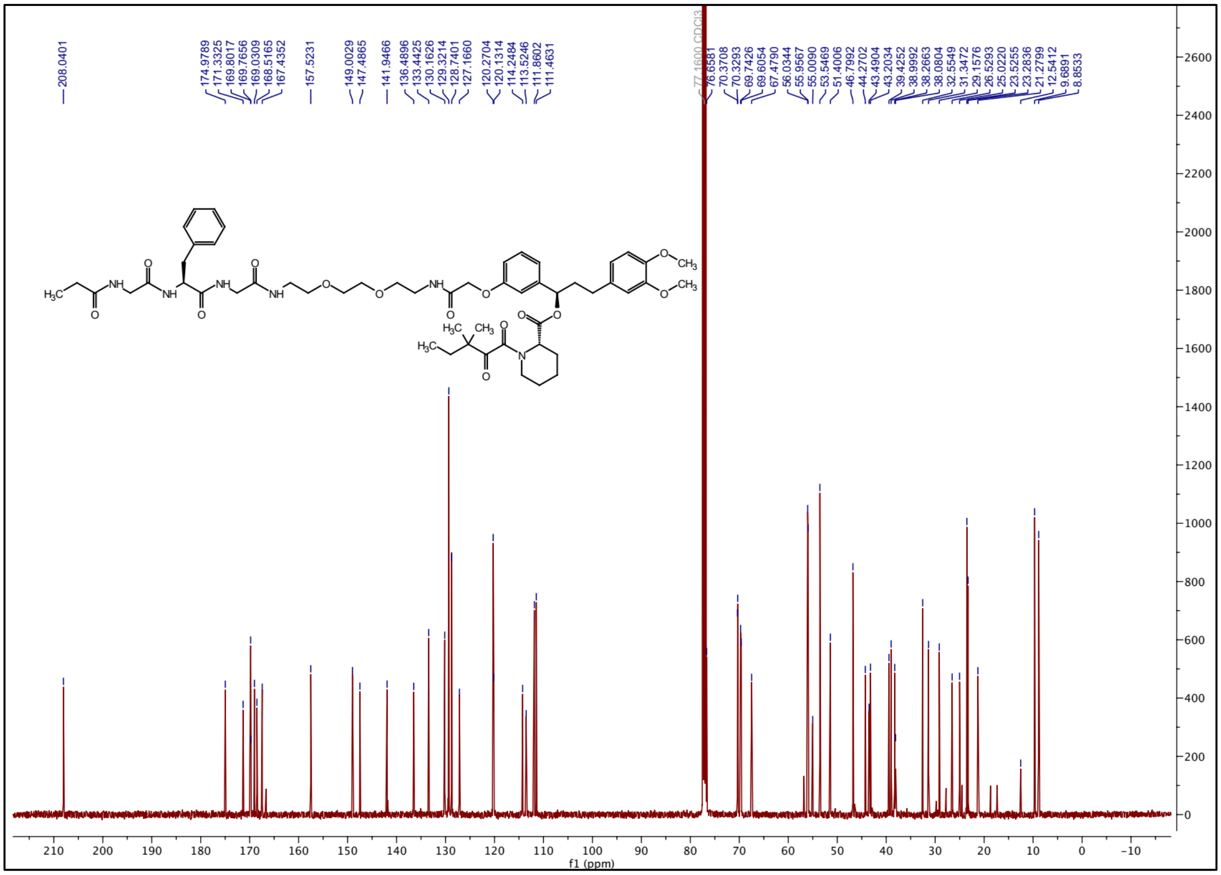

### $^{1}H$ NMR spectrum of ZA19

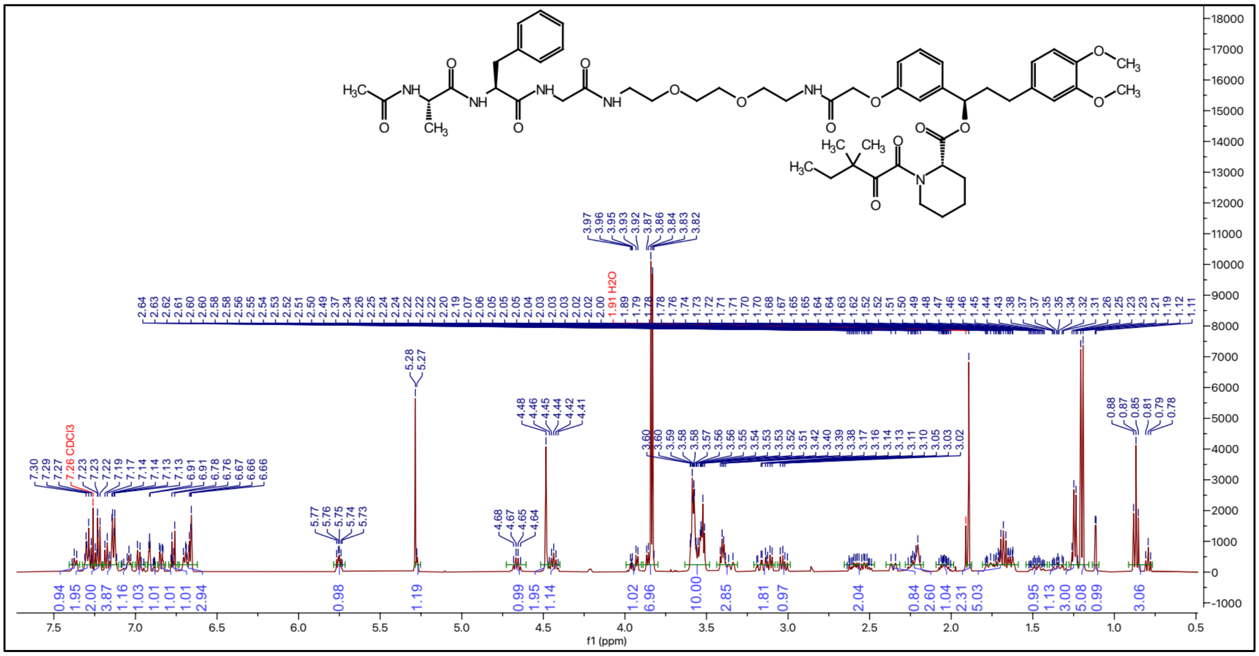

### $^{13}C$ NMR spectrum of ZA19

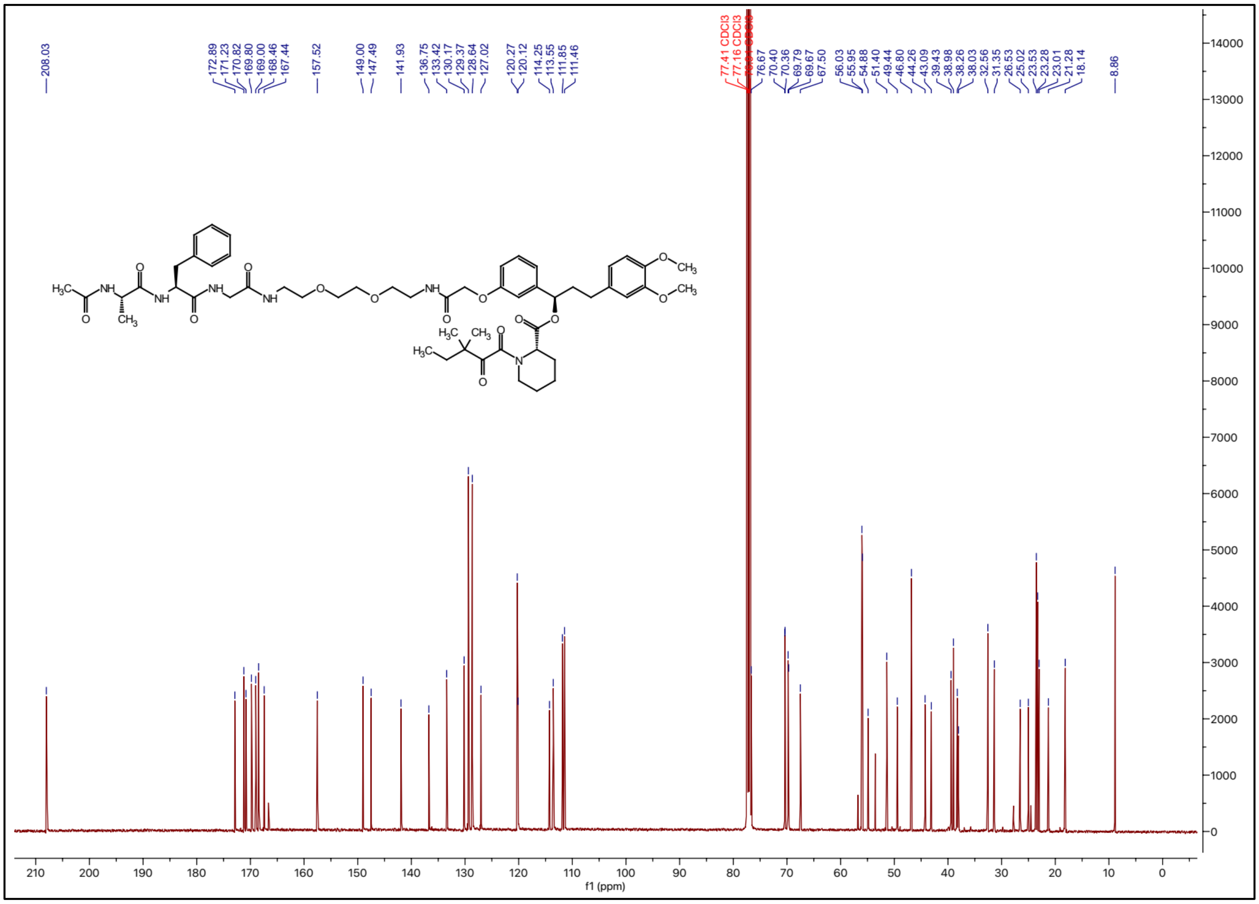

### [1]H NMR spectrum of ZA20

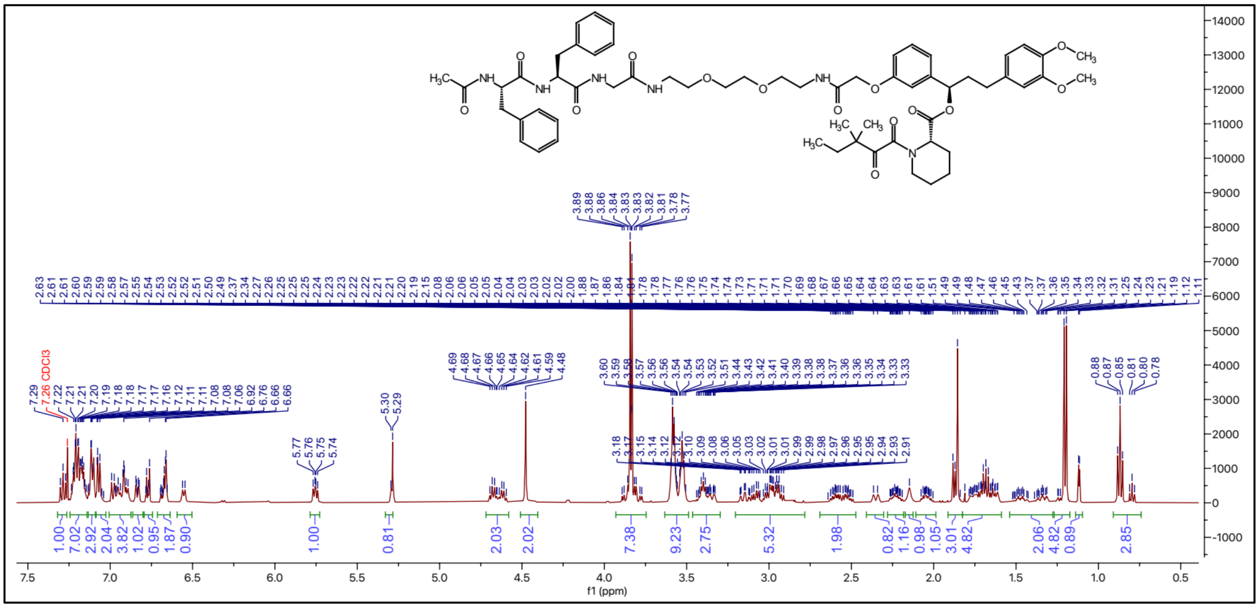

### [13]C NMR spectrum of ZA20

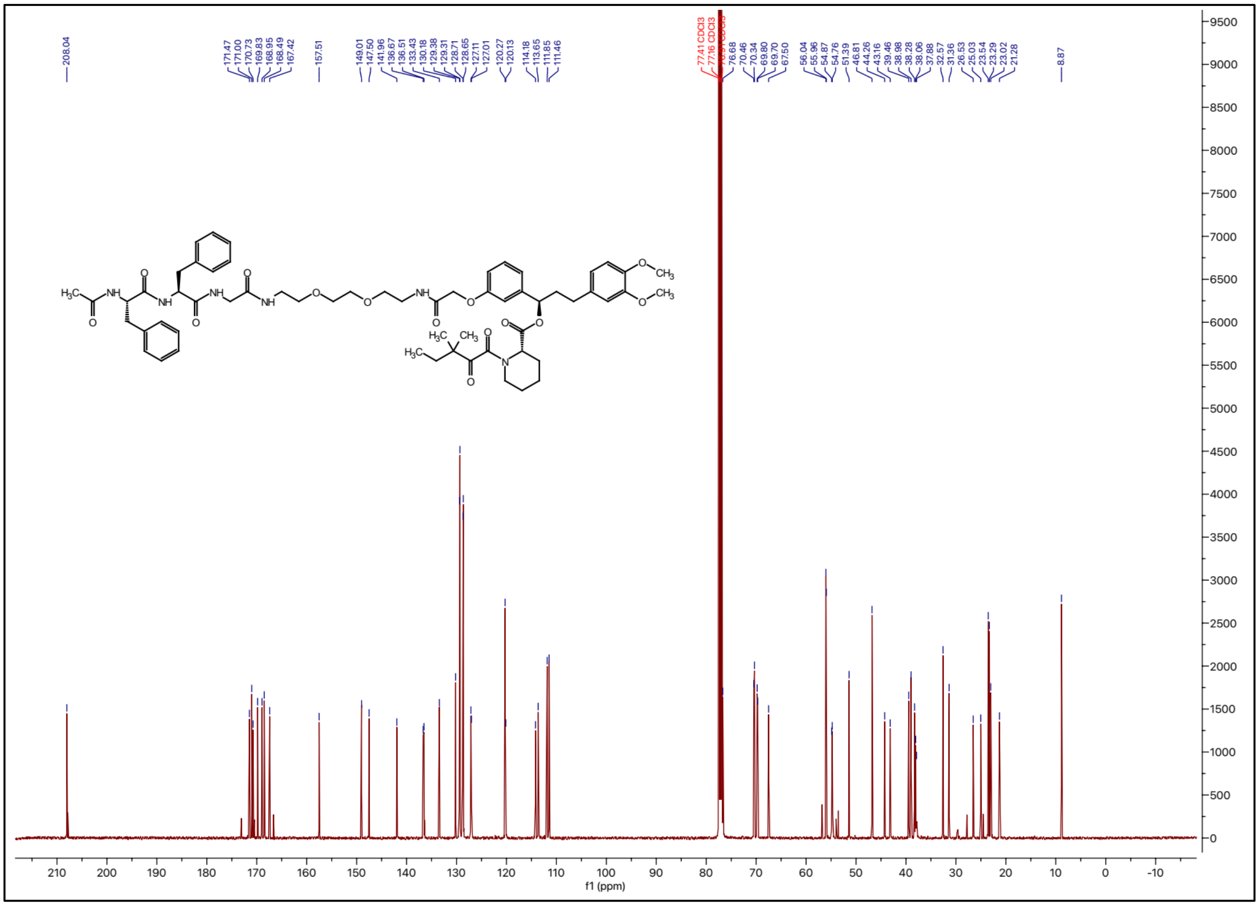

### $^{1}$H NMR spectrum of ZA21

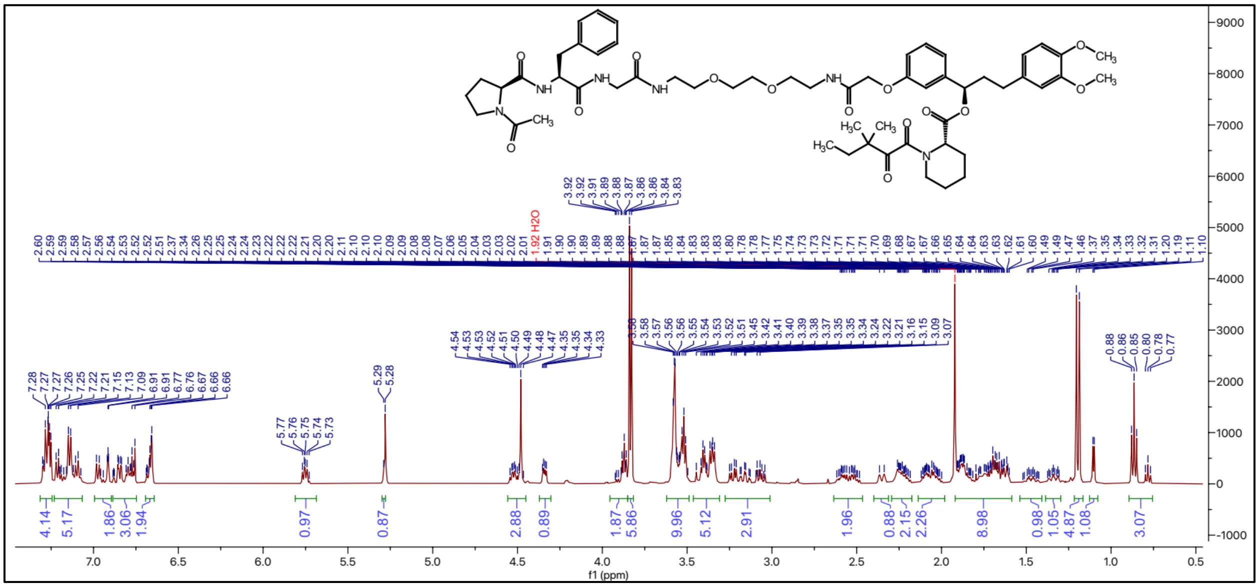

### $^{13}$C NMR spectrum of ZA21

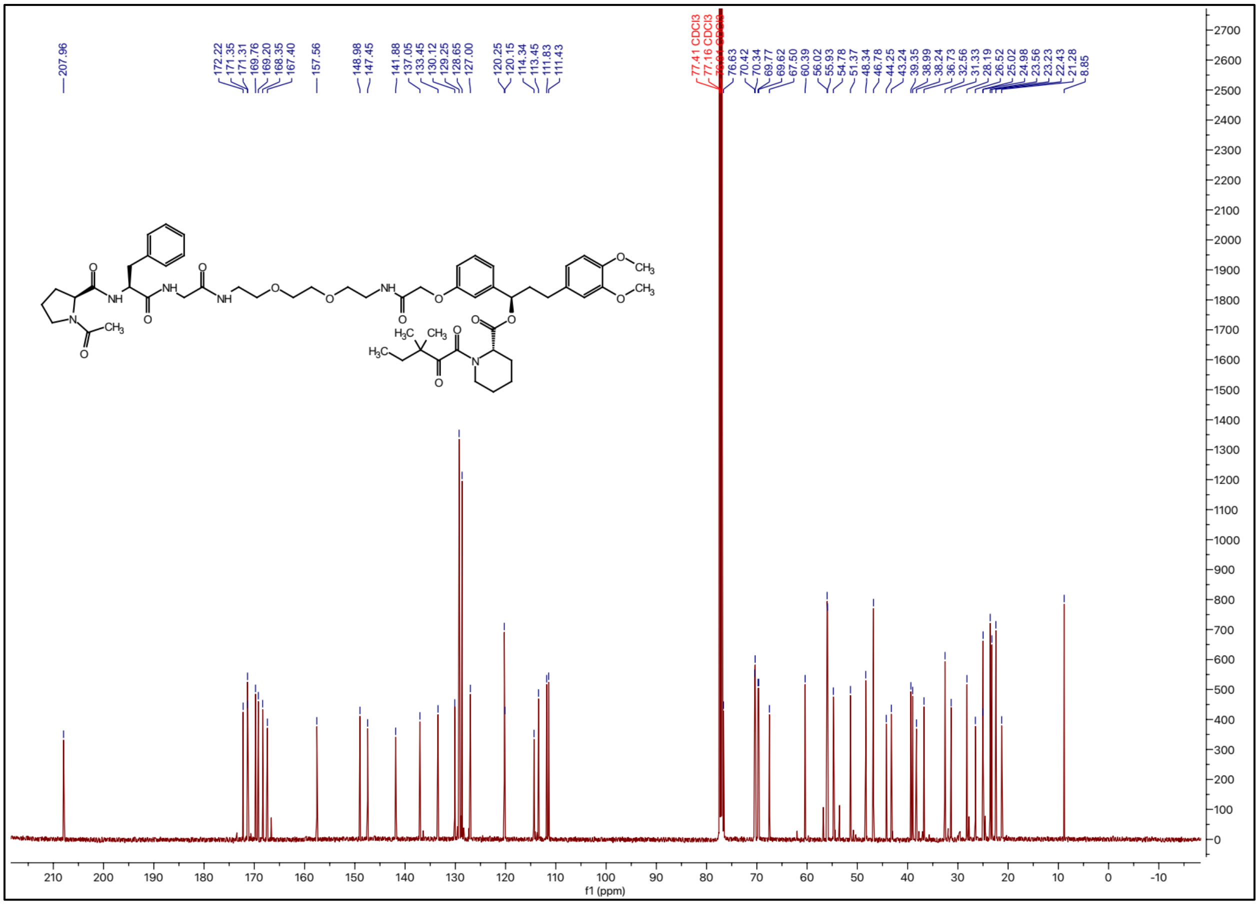

### $^{1}$H NMR spectrum of ZA22

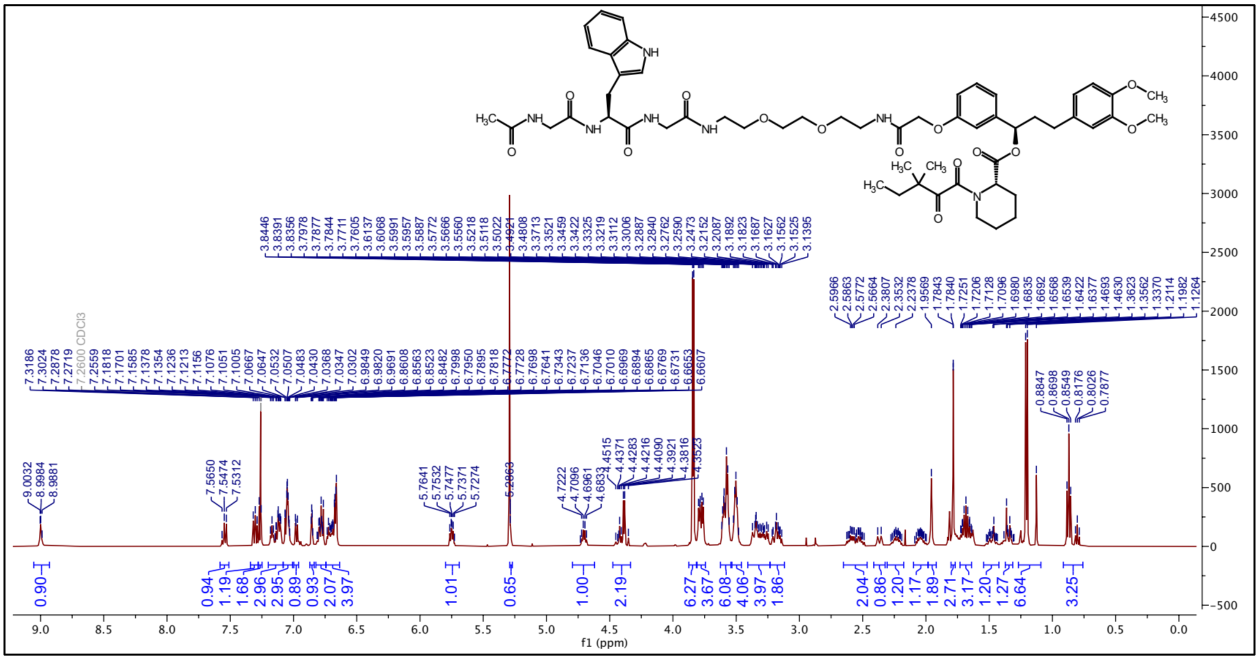

### $^{13}$C NMR spectrum of ZA22

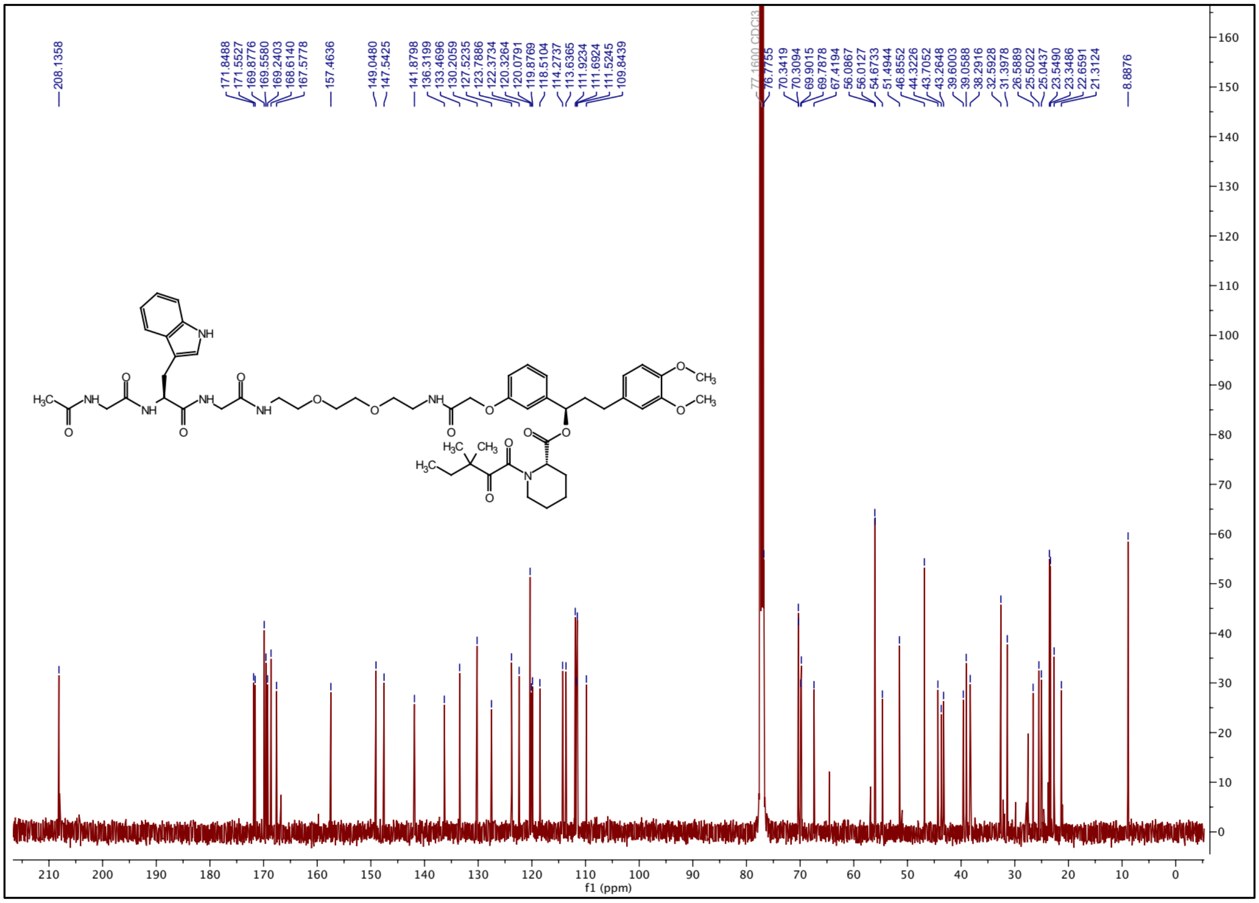

**[1]H NMR spectrum of ZA23**

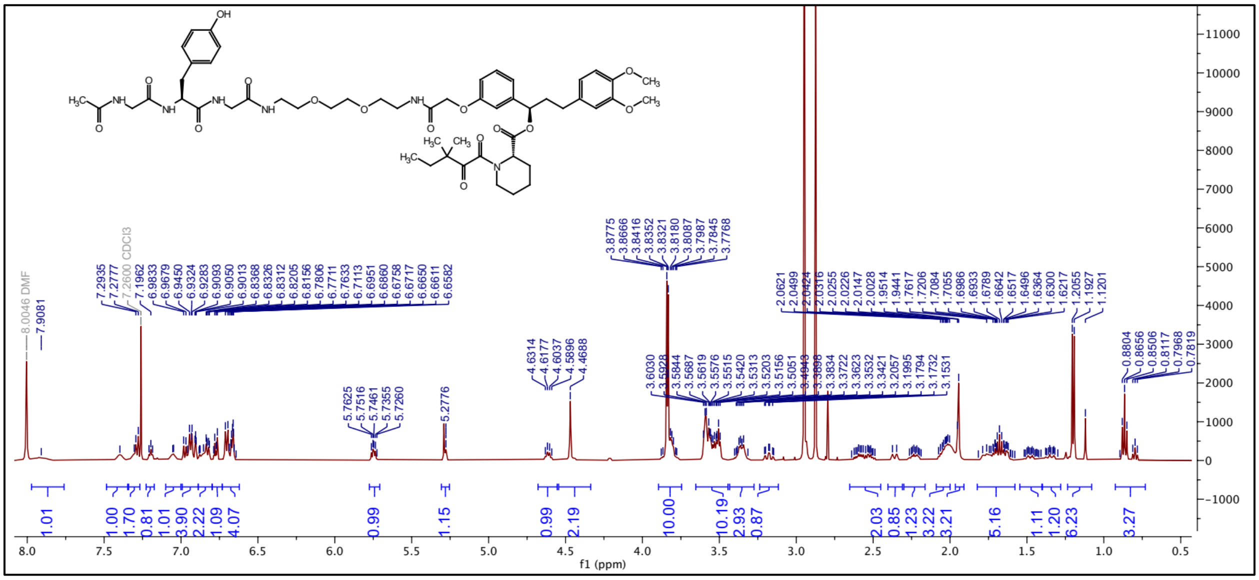

**[13]C NMR spectrum of ZA23**

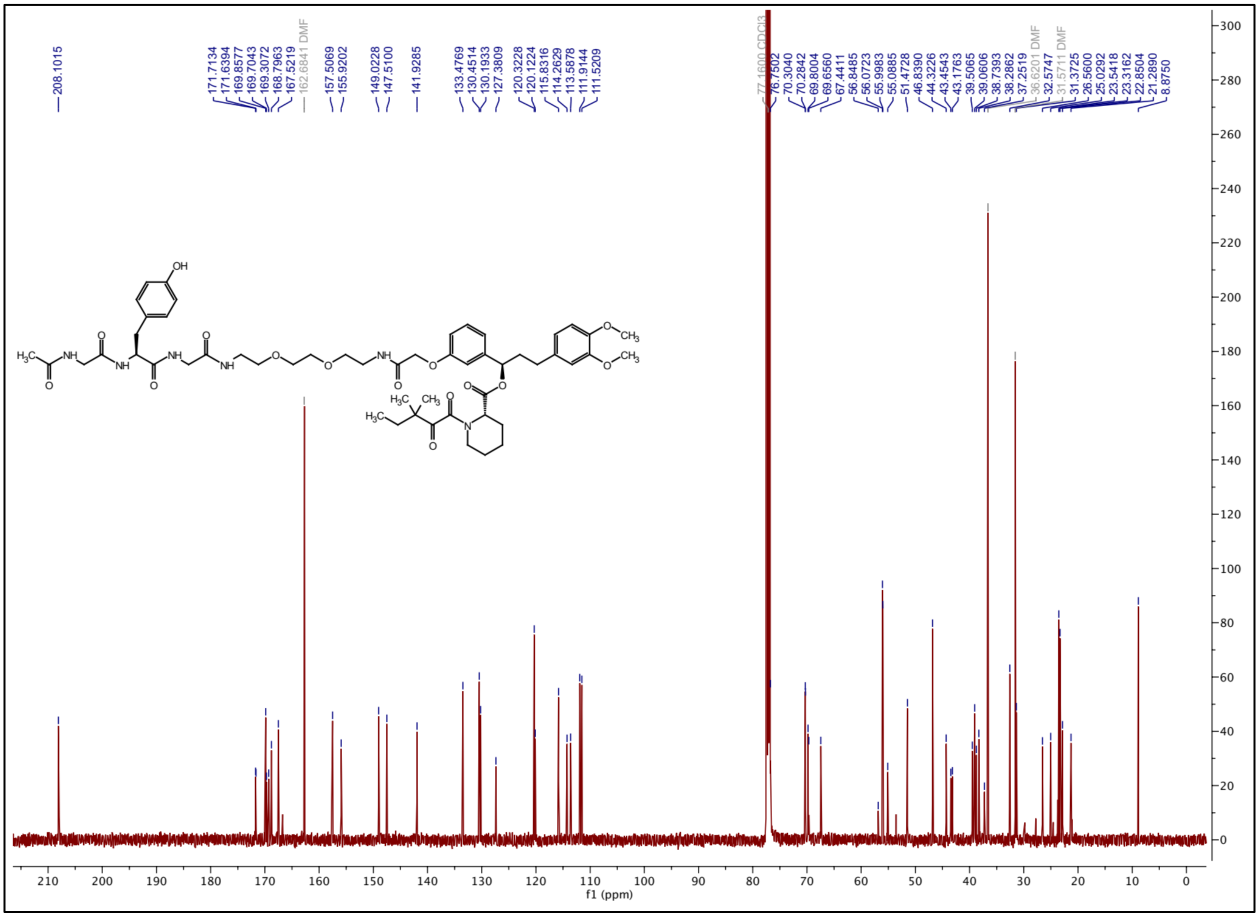

### $^{1}$H NMR spectrum of ZA24

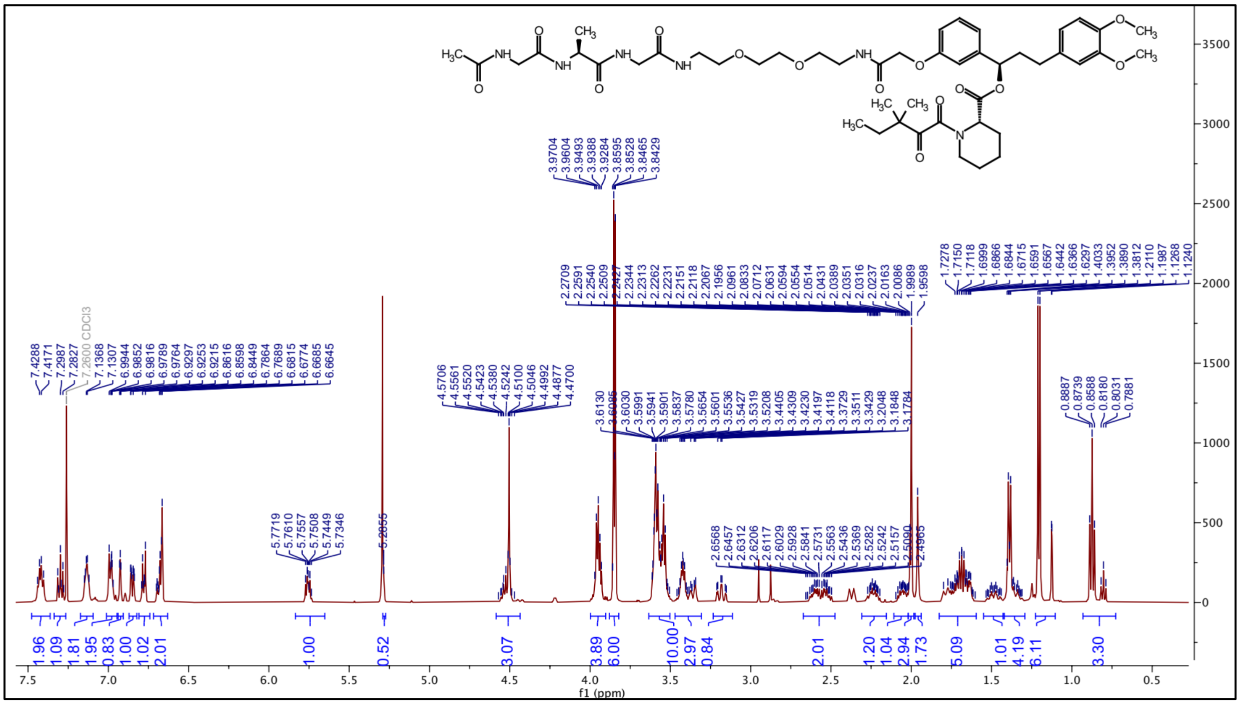

### $^{13}$C NMR spectrum of ZA24

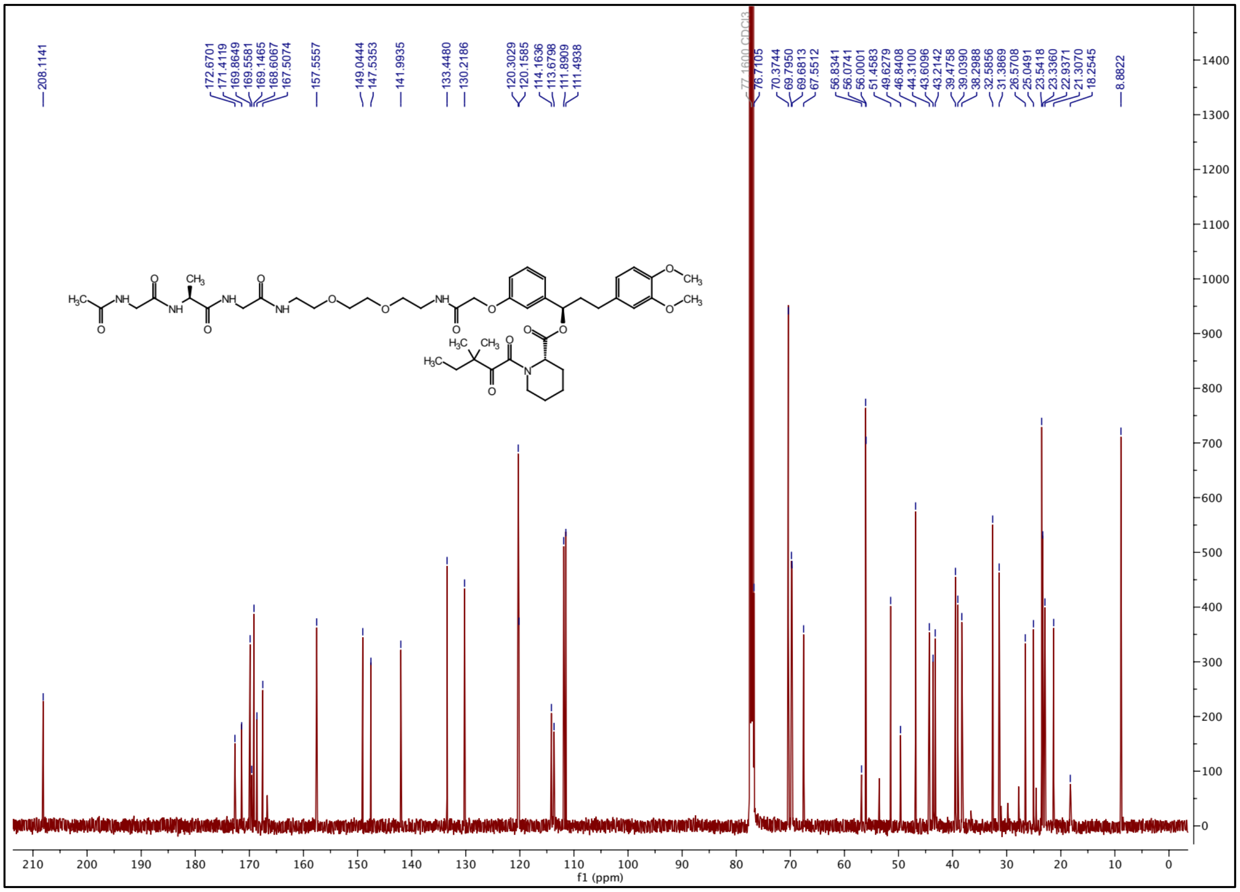

**¹H NMR spectrum of ZA25**

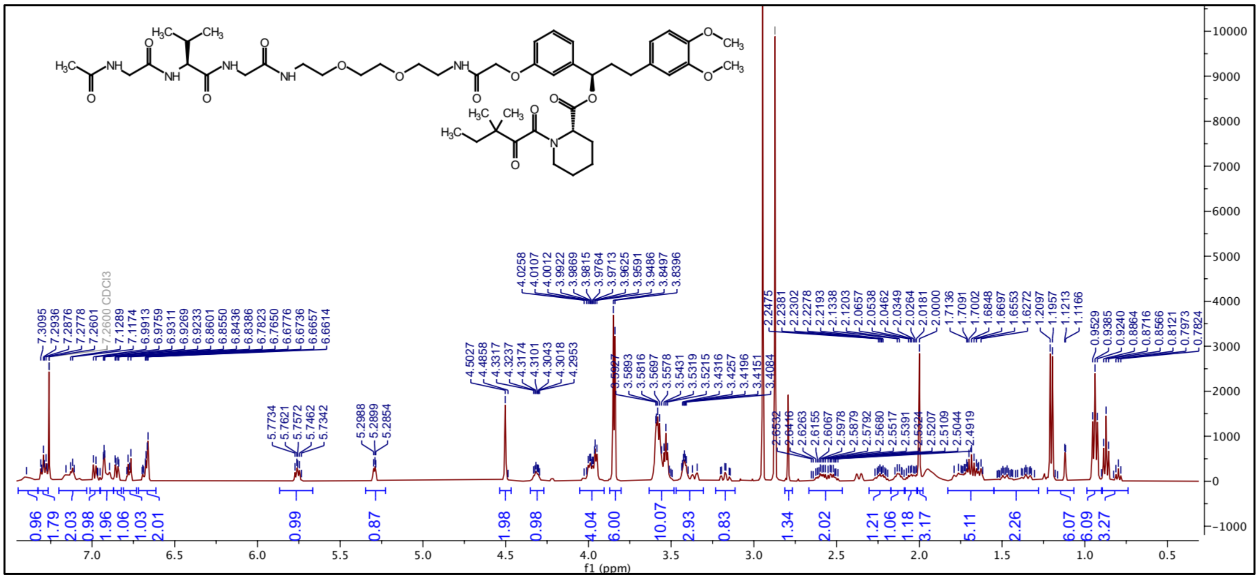

**¹³C NMR spectrum of ZA25**

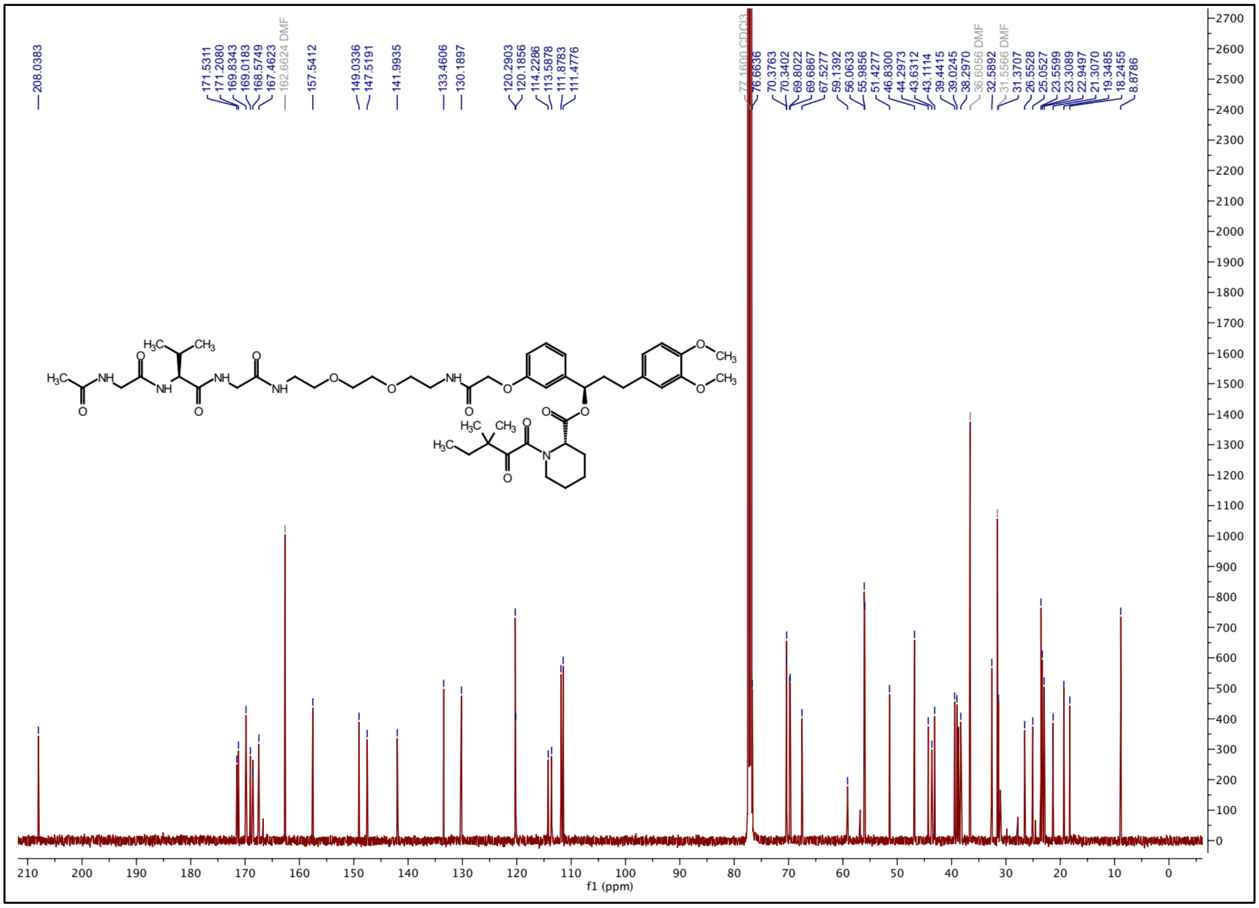

$^1$H NMR spectrum of ZA26

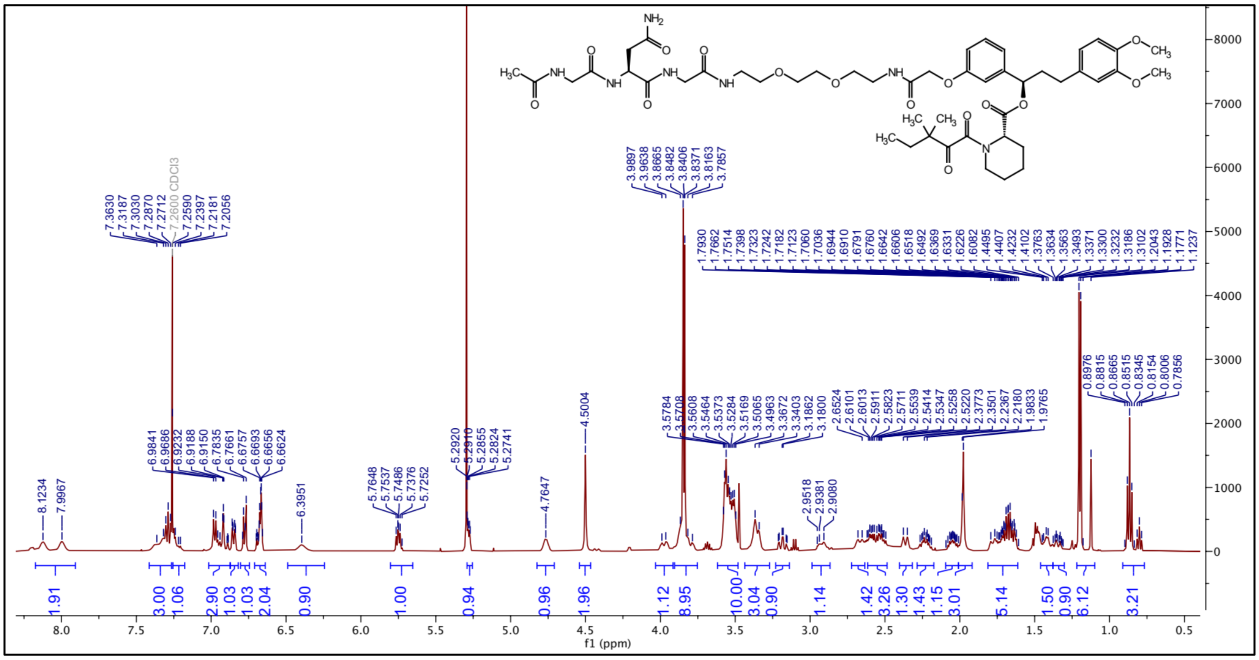

$^{13}$C NMR spectrum of ZA26

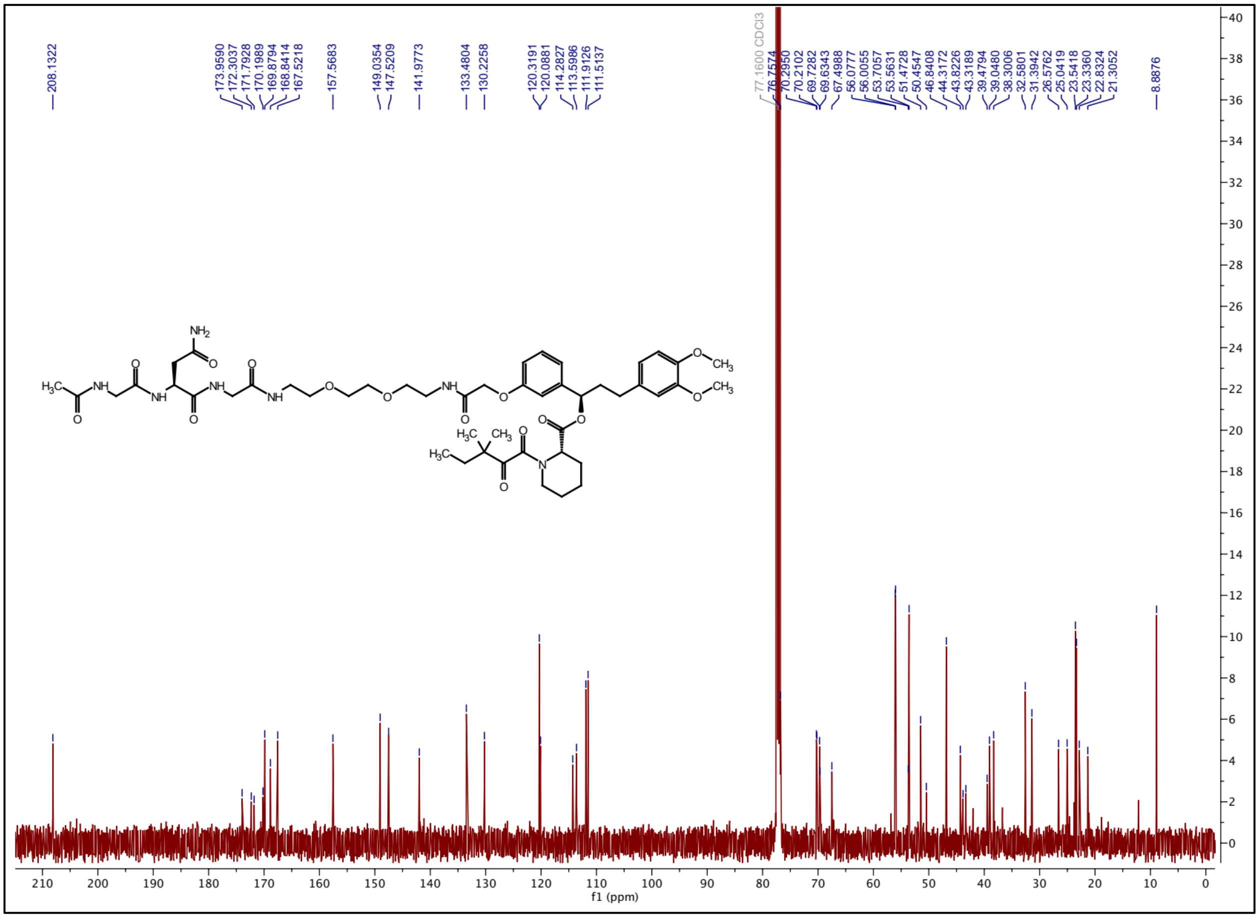

### 1H NMR spectrum of ZA27

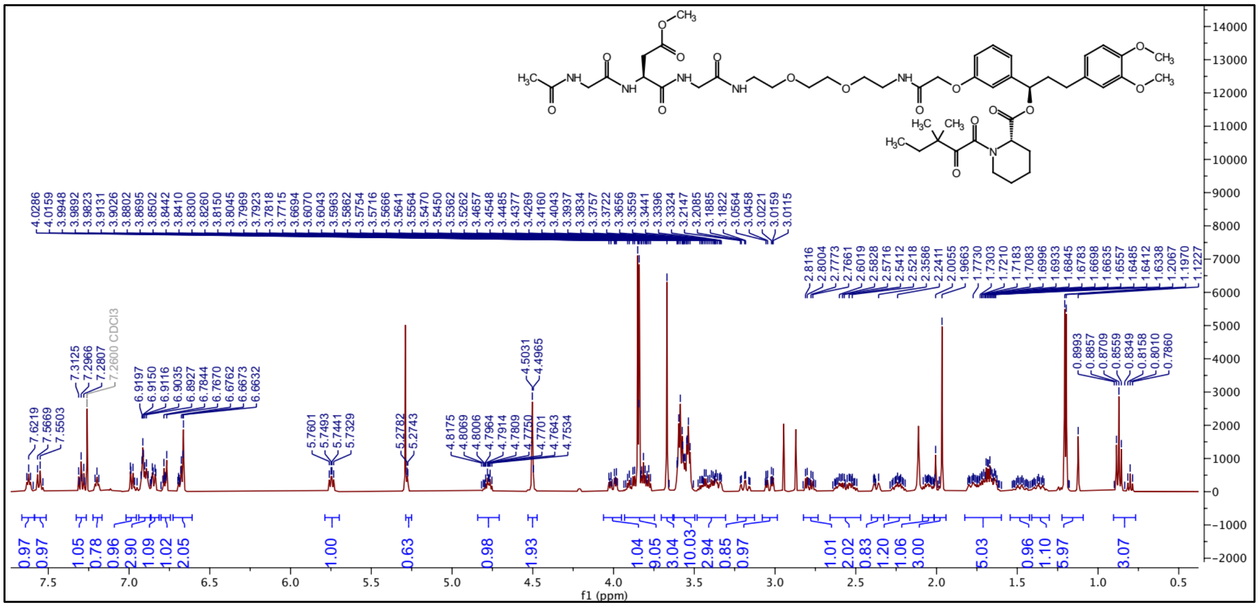

### 13C NMR spectrum of ZA27

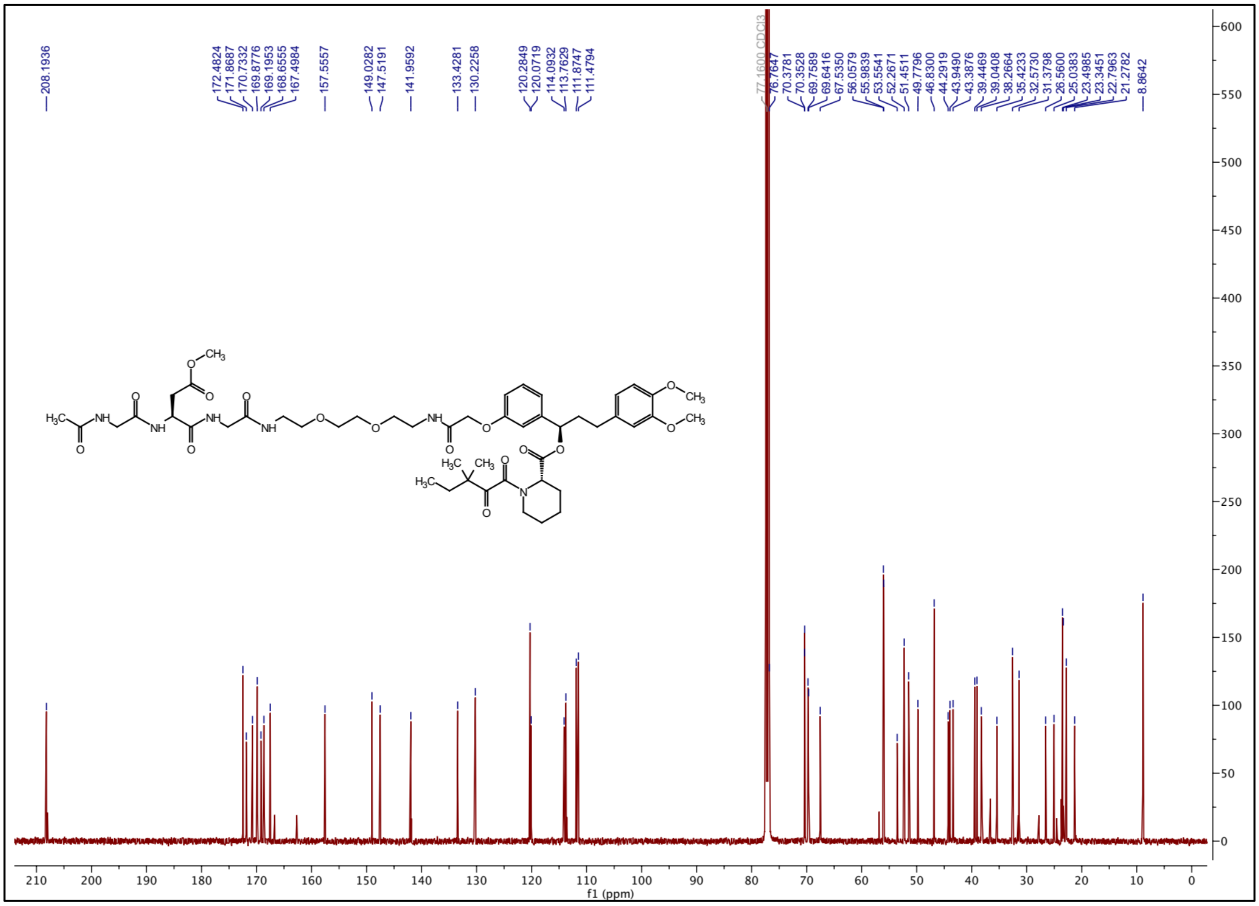

### $^{1}H$ NMR spectrum of ZA28

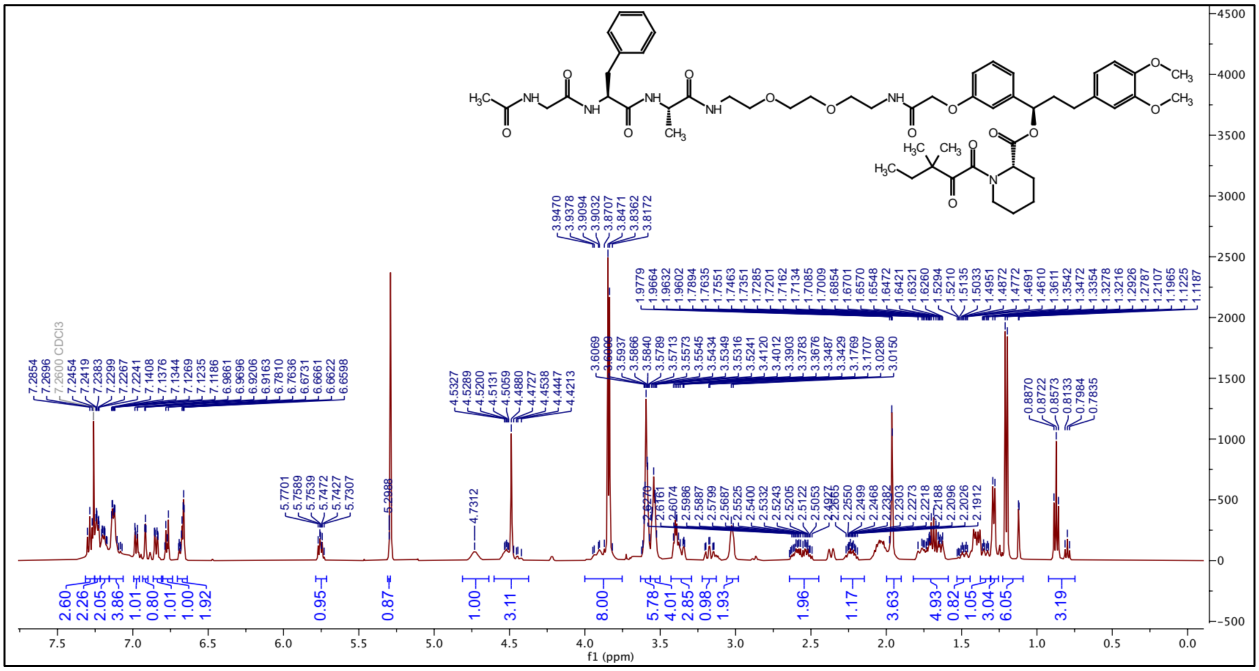

### $^{13}C$ NMR spectrum of ZA28

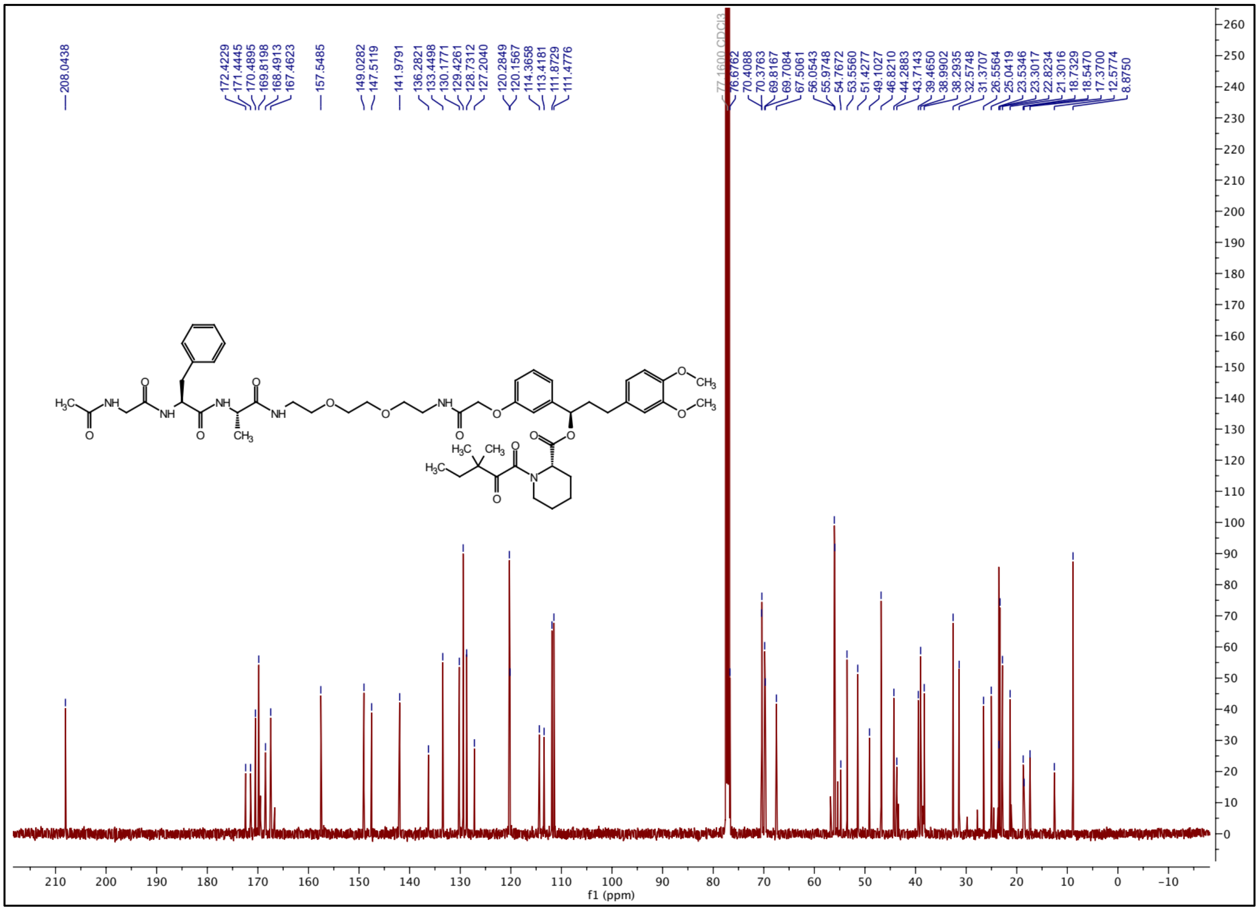

### $^{1}H$ NMR spectrum of ZA29

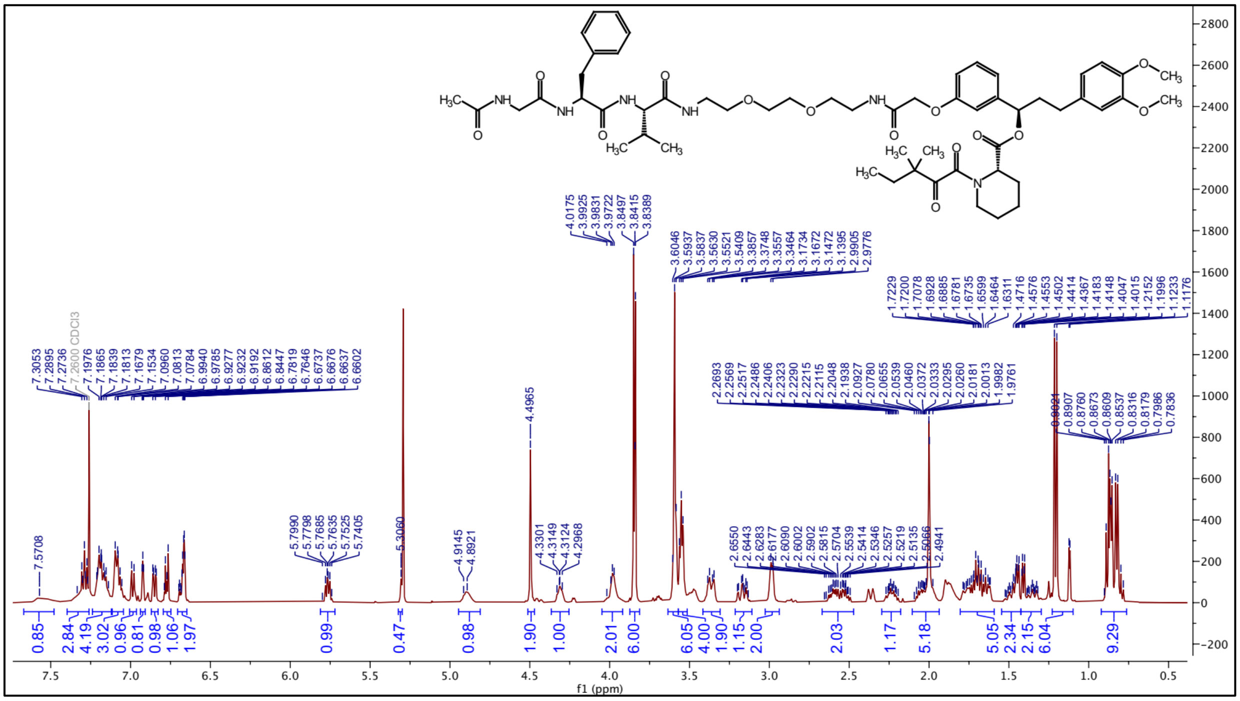


### $^{13}C$ NMR spectrum of ZA29

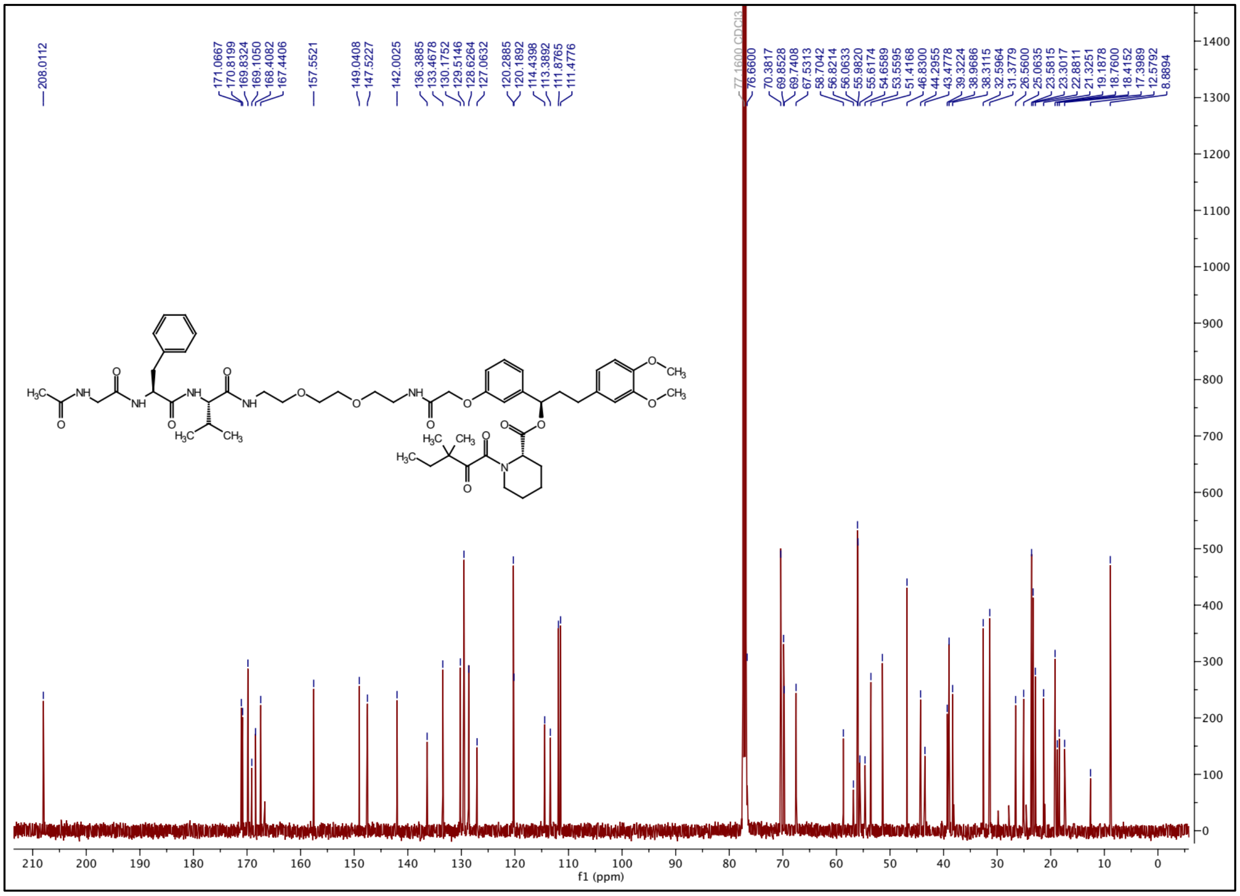

### $^{1}$H NMR spectrum of ZA30

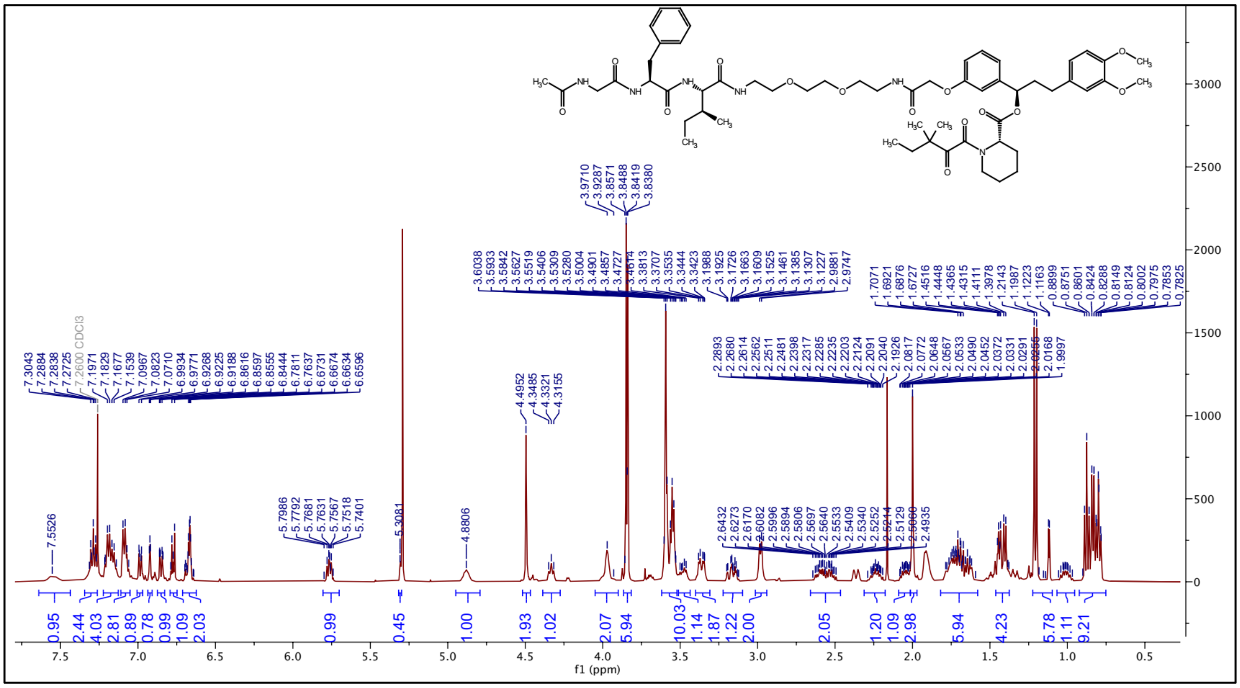

### $^{13}$C NMR spectrum of ZA30

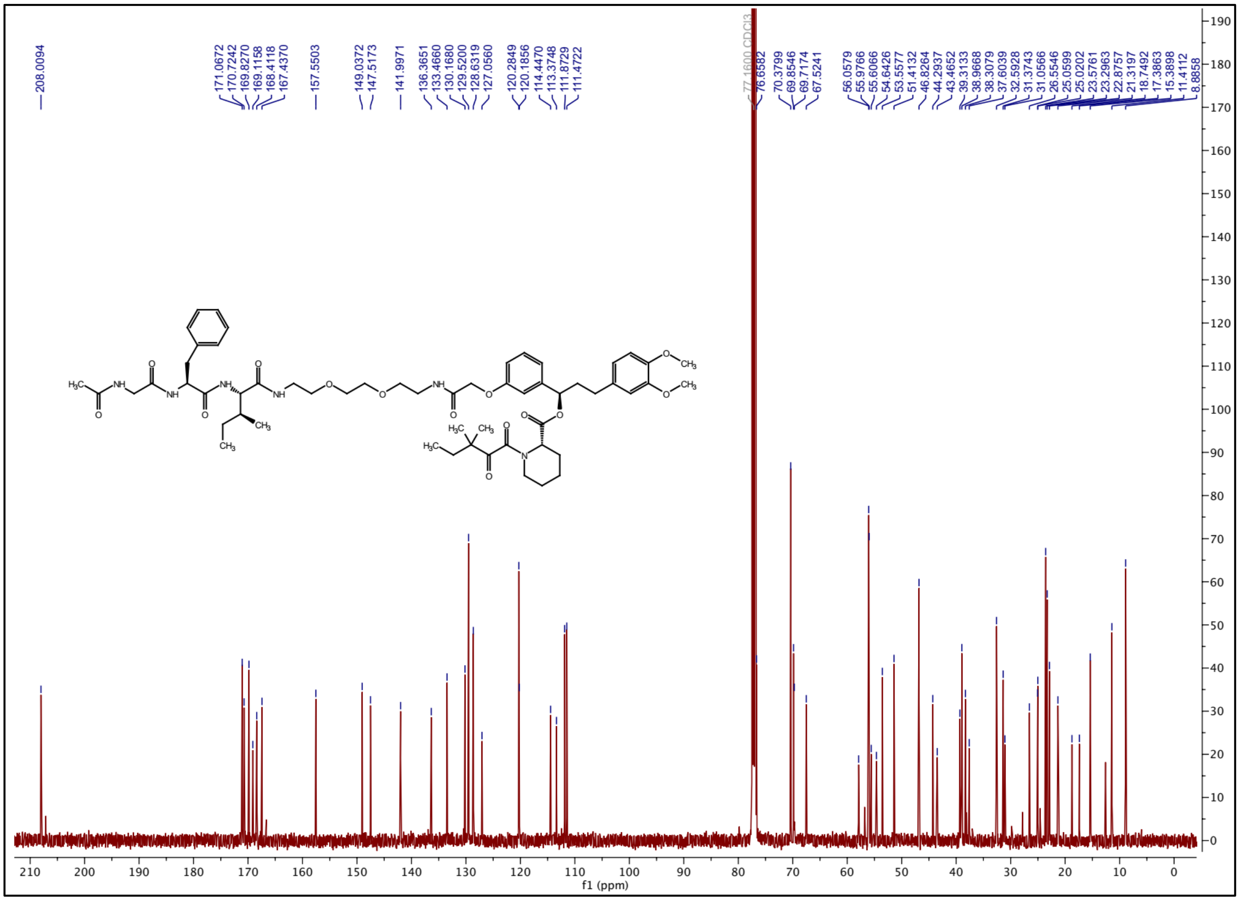

### $^1$H NMR spectrum of ZA31

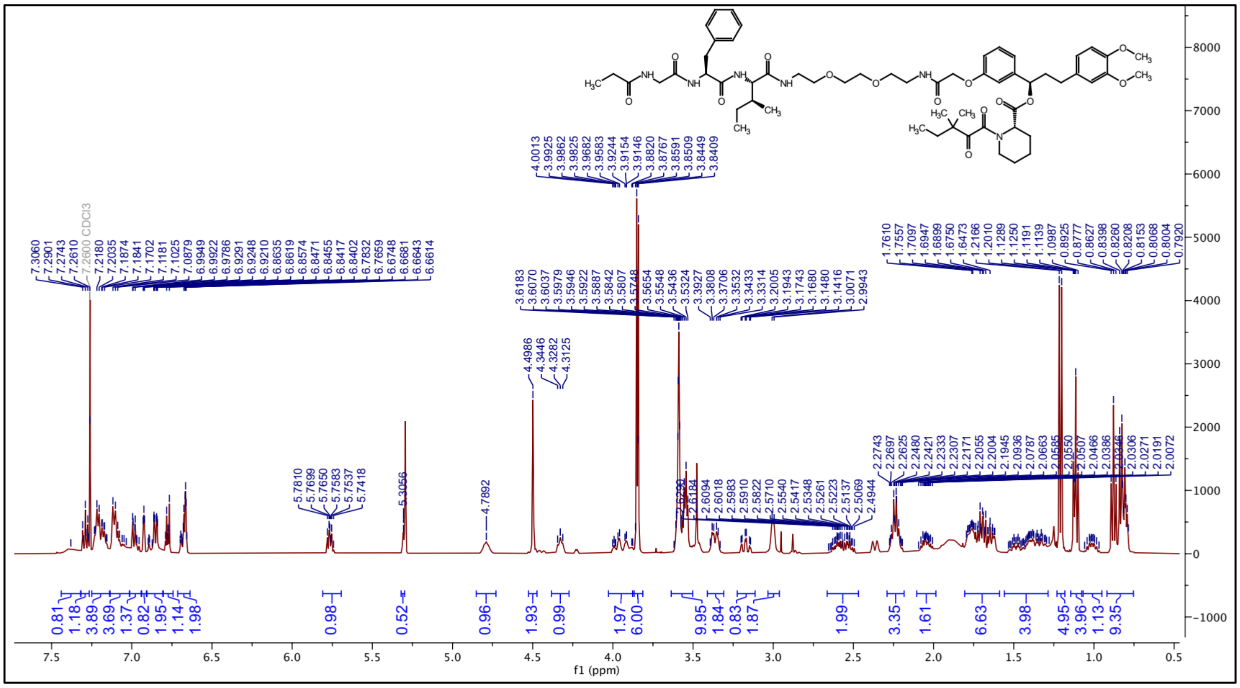

### $^{13}$C NMR spectrum of ZA31

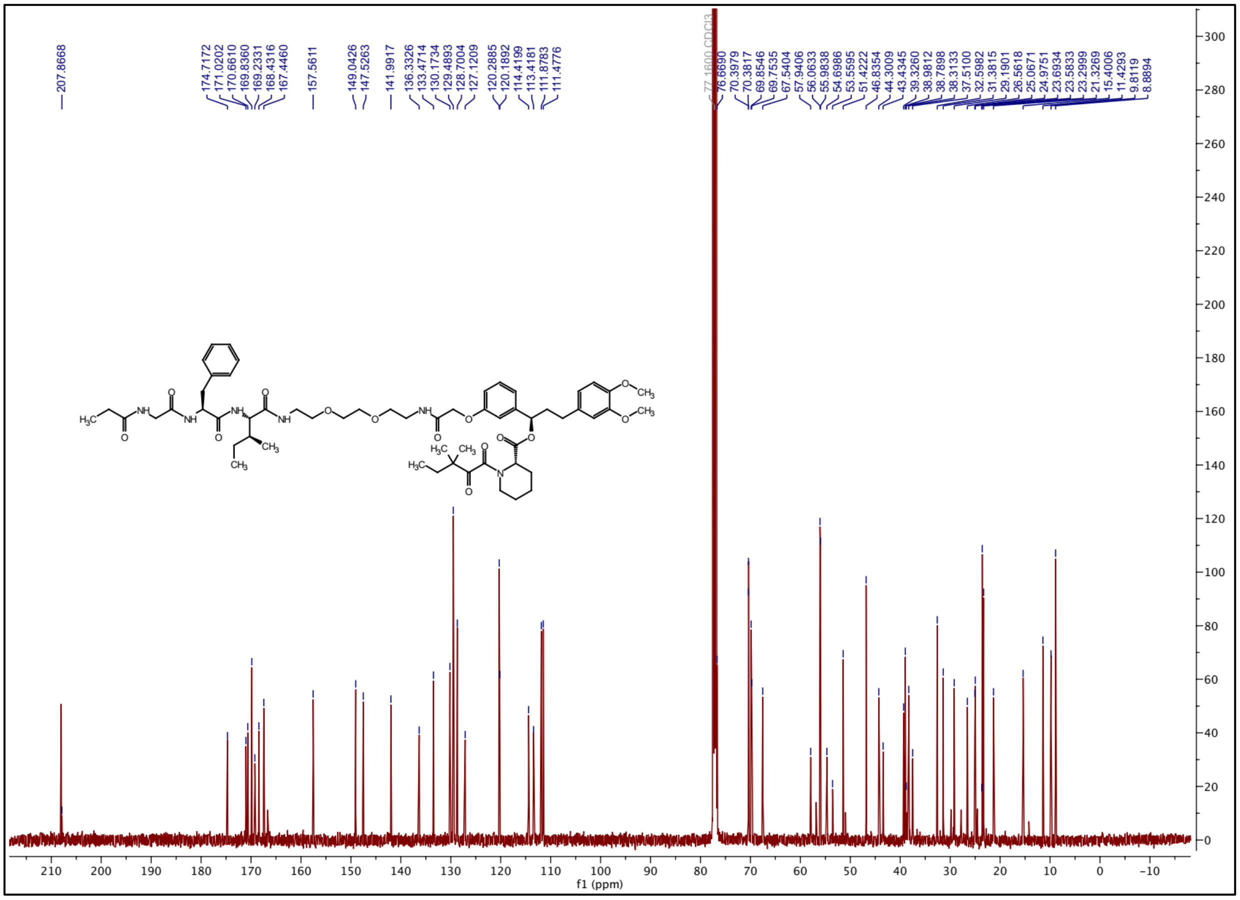

### [1]H NMR spectrum of ZA32

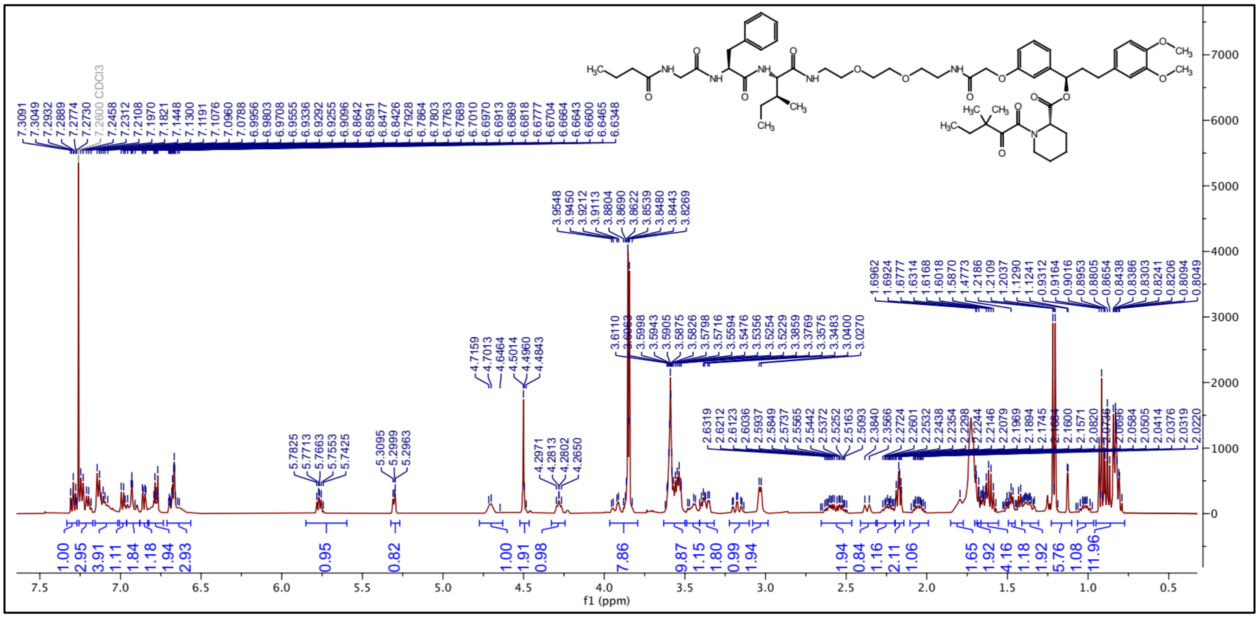

### [13]C NMR spectrum of ZA32

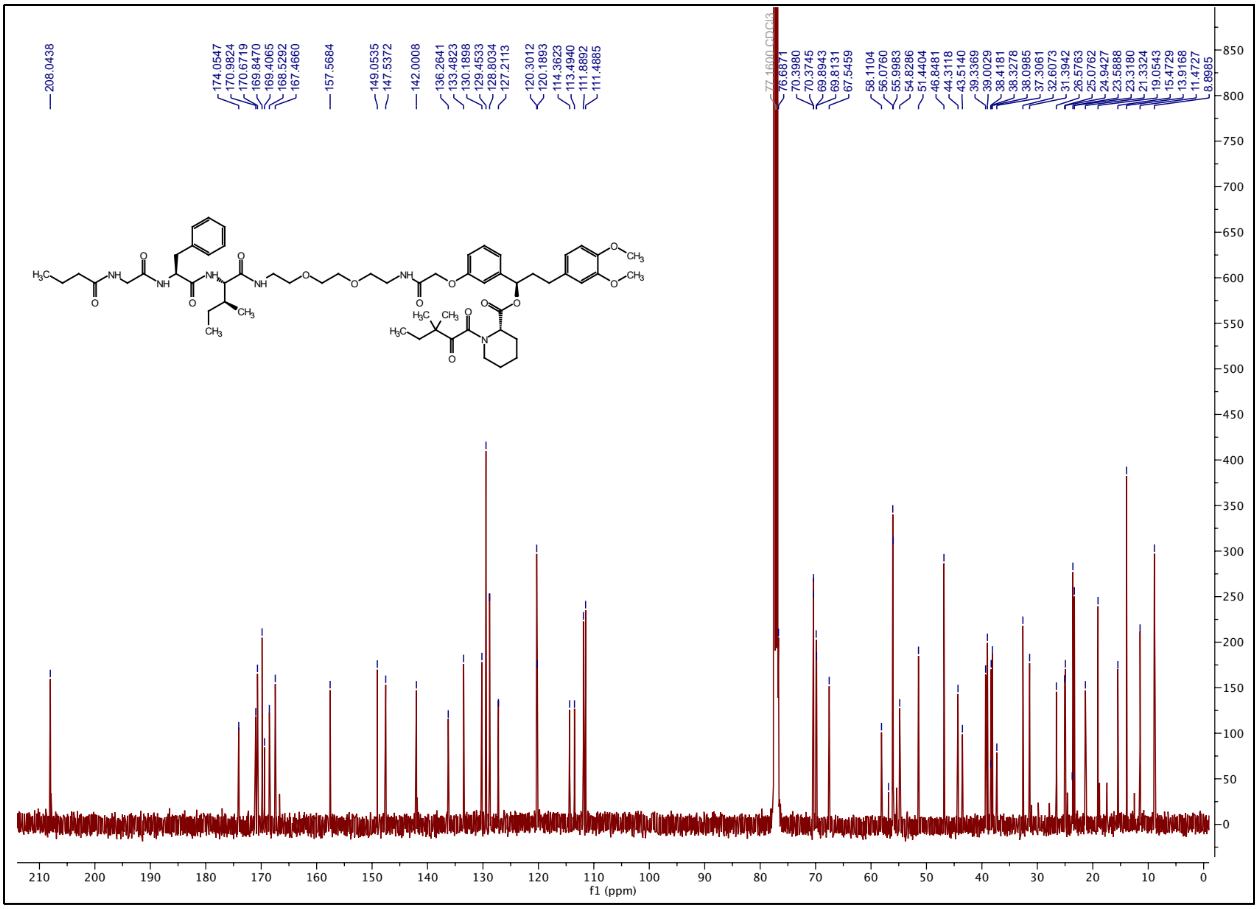

### [1]H NMR spectrum of ZA33

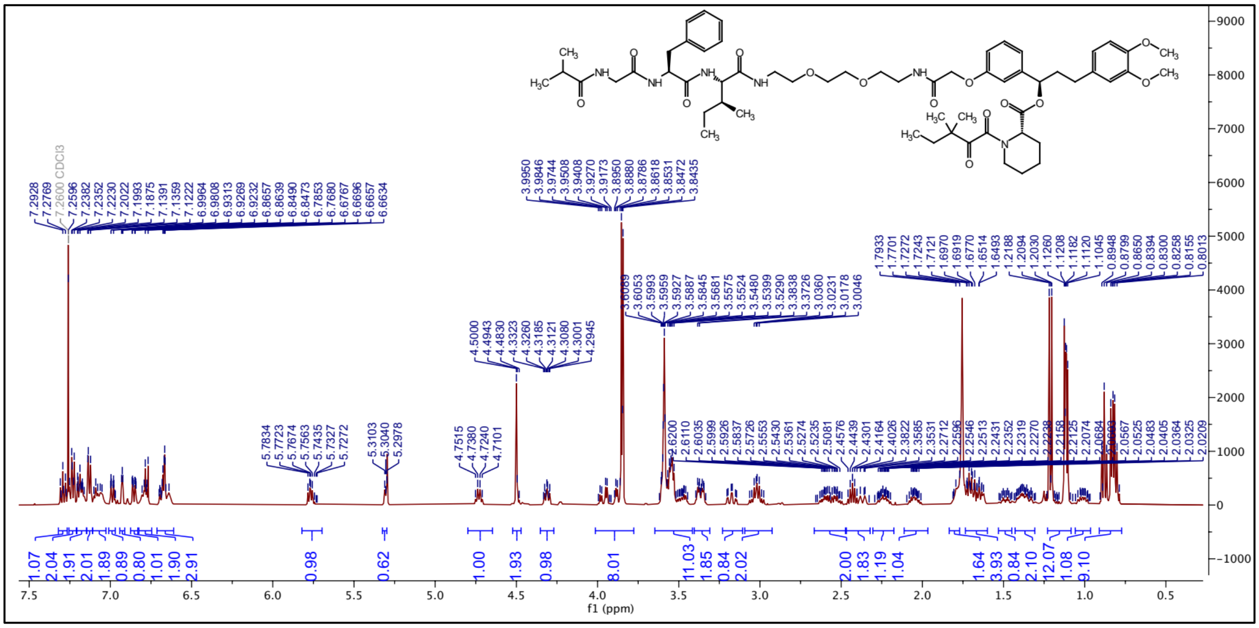

### [13]C NMR spectrum of ZA33

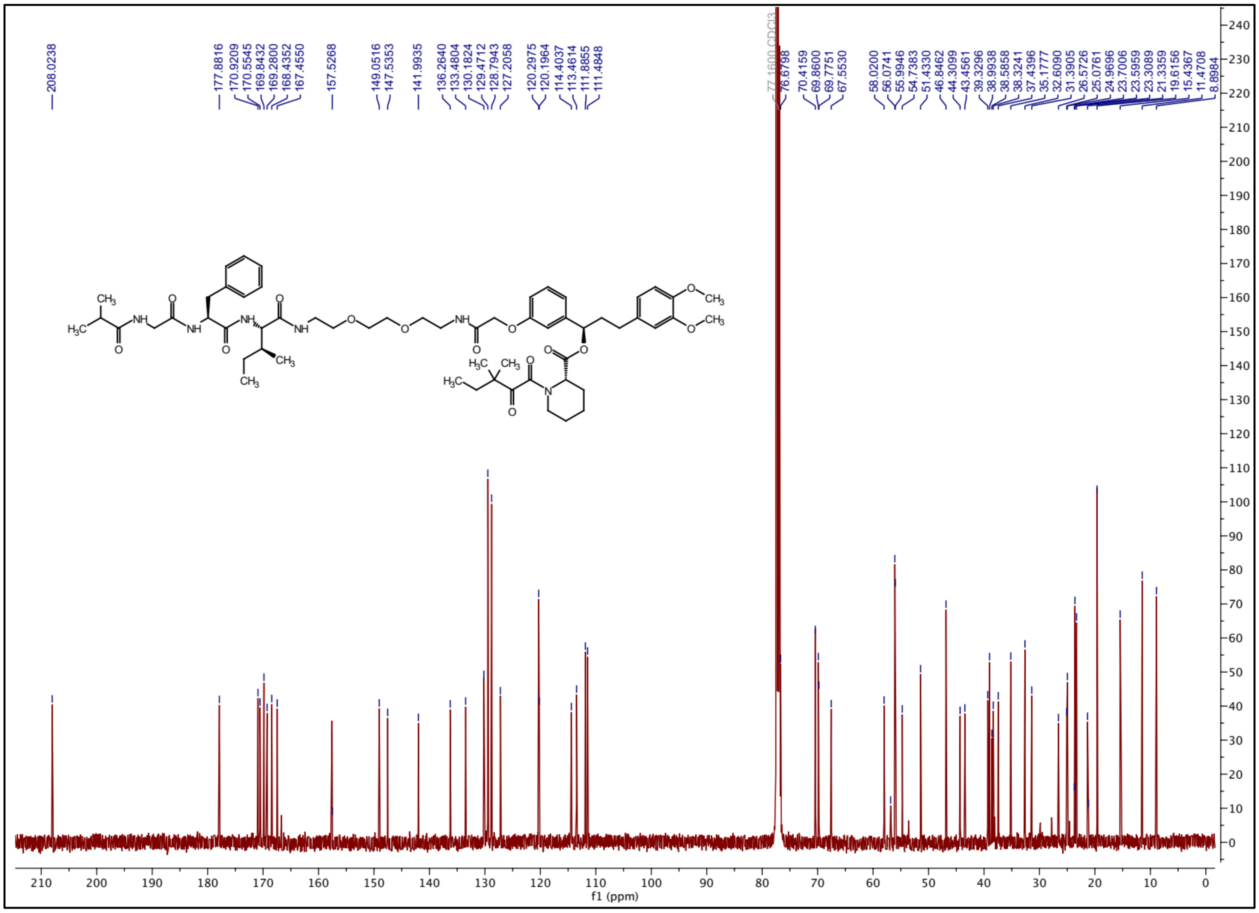

### $^{1}H$ NMR spectrum of ZA34

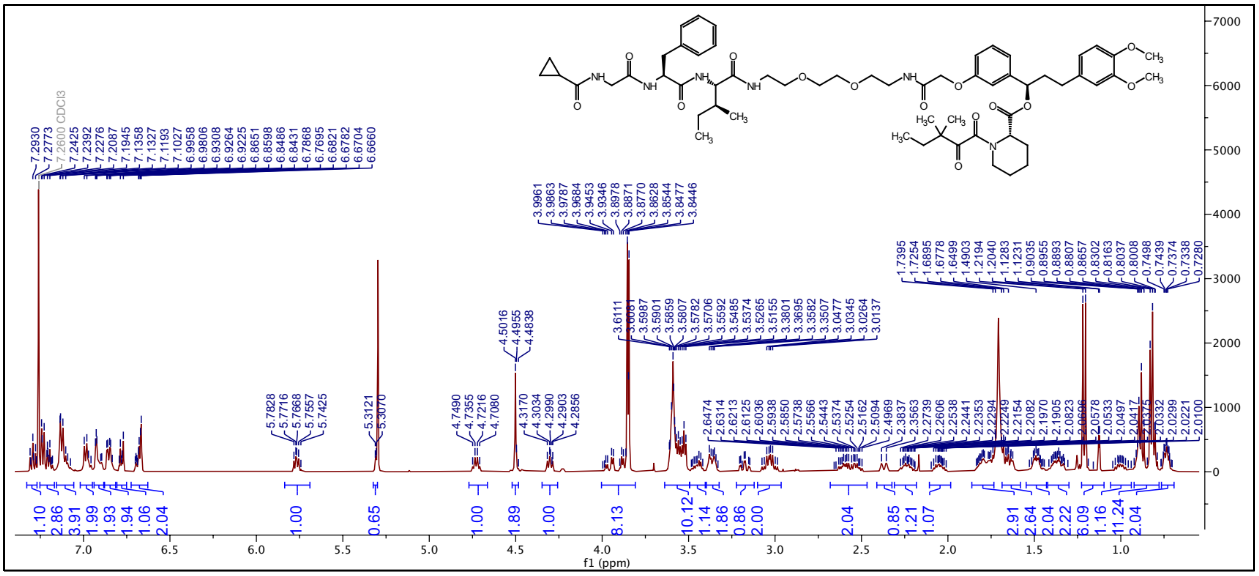

### $^{13}C$ NMR spectrum of ZA34

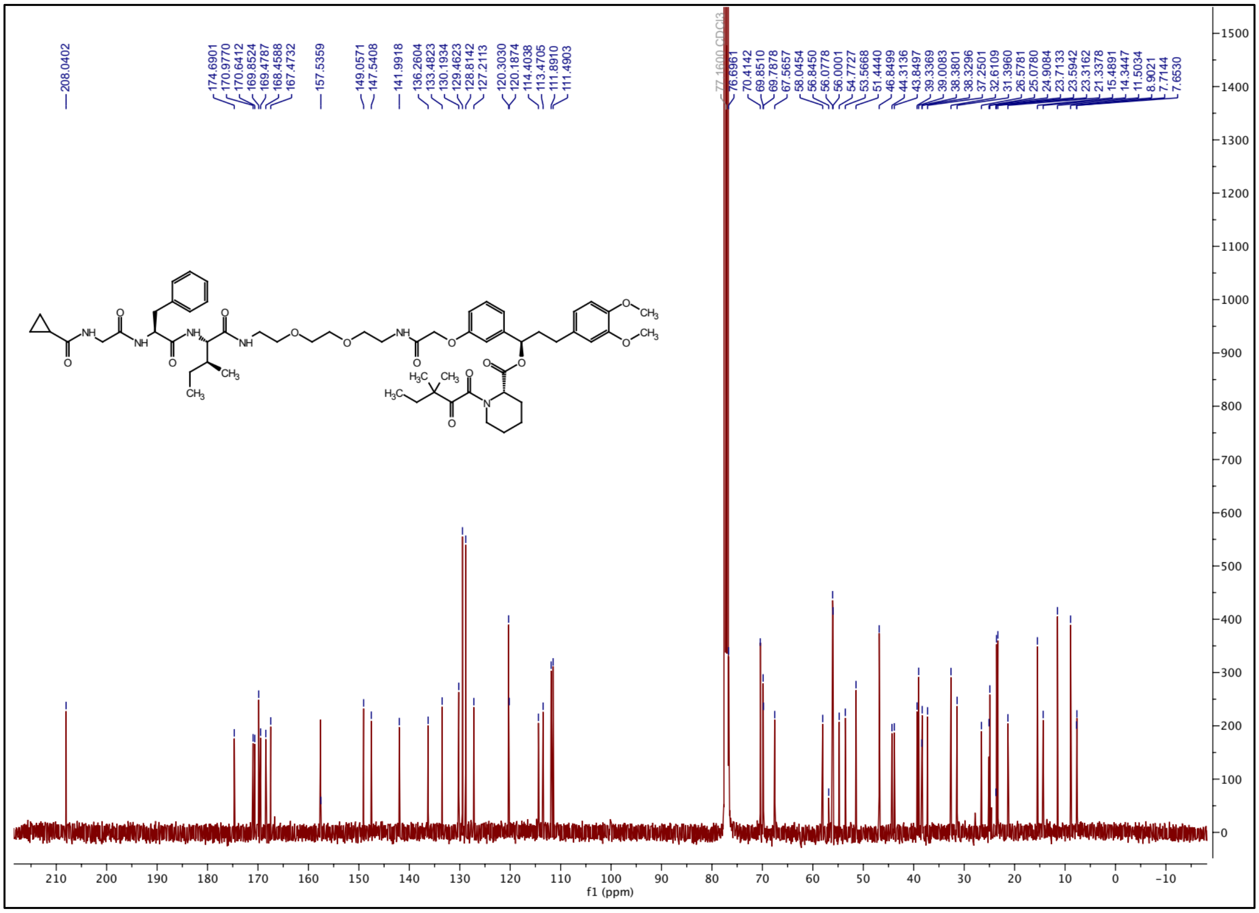

### $^{1}$H NMR spectrum of ZA35

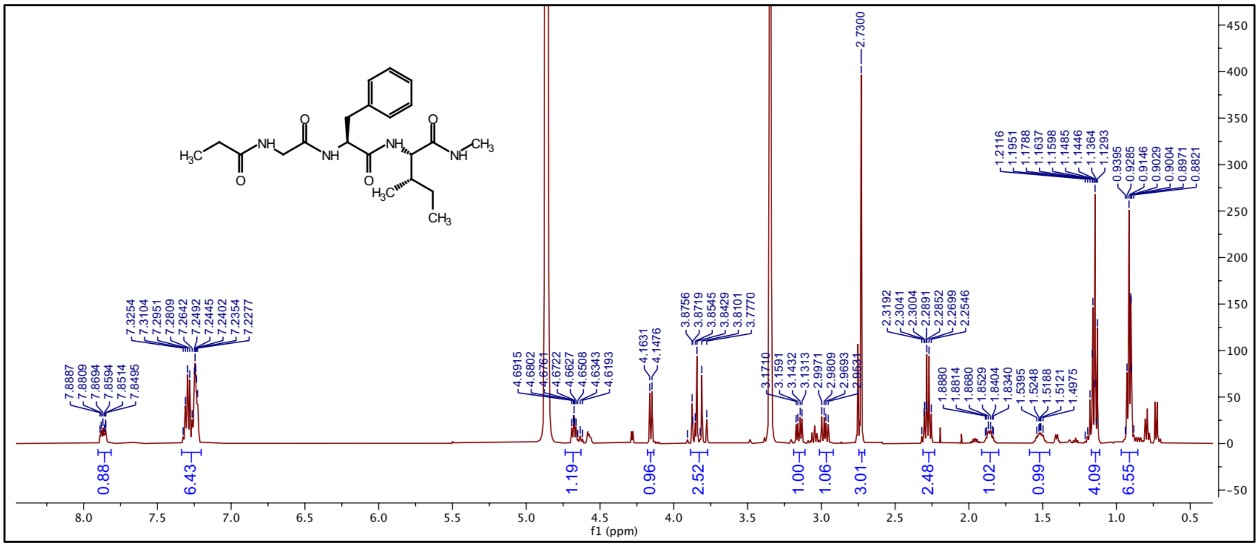

### $^{13}$C NMR spectrum of ZA35

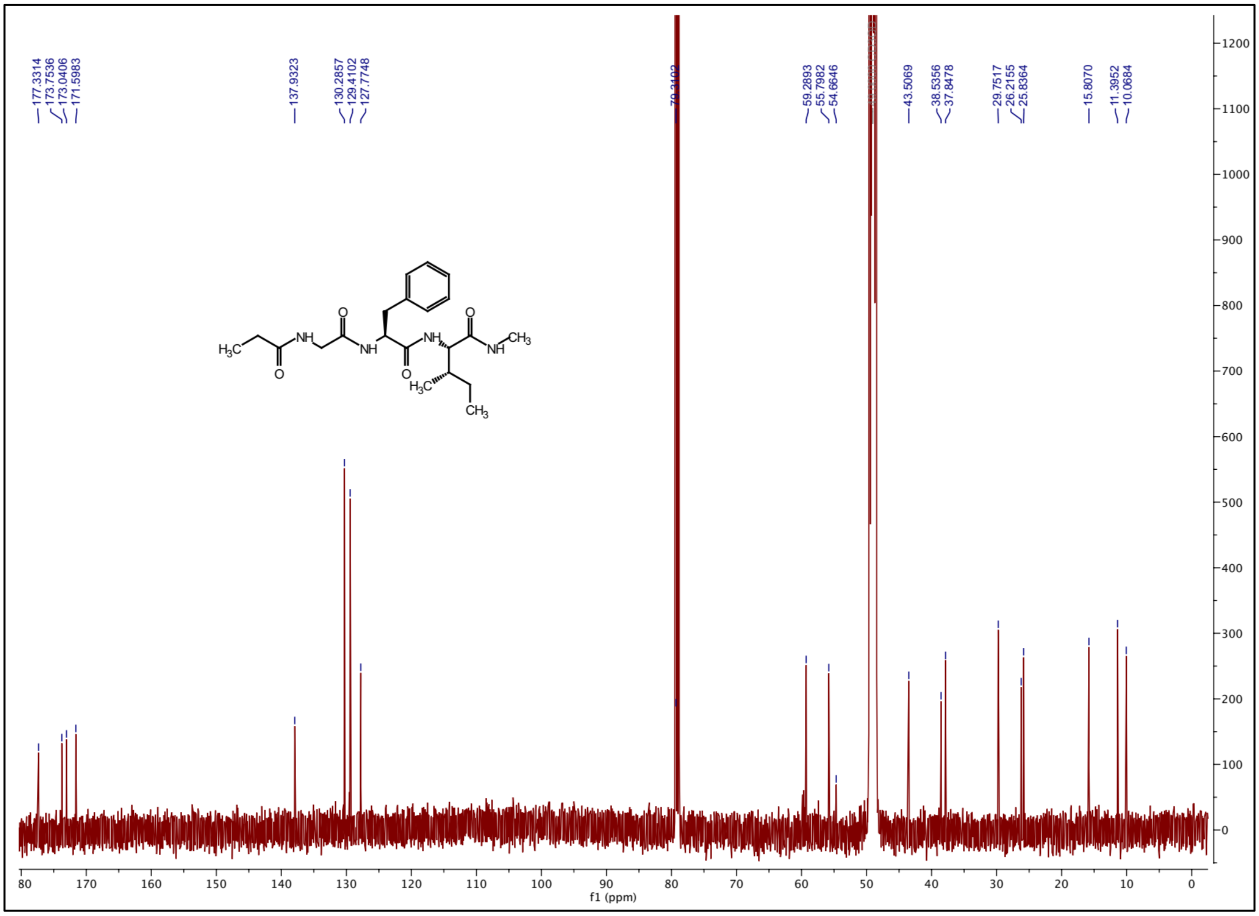

### $^{1}H$ NMR spectrum of ZA36

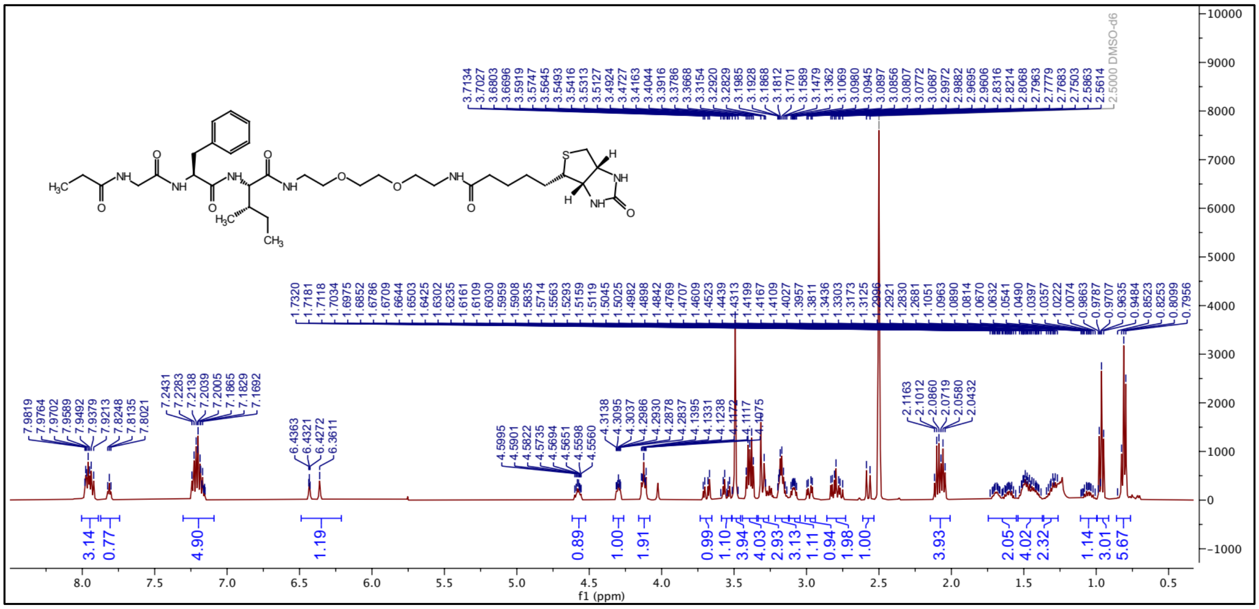

### $^{13}C$ NMR spectrum of ZA36

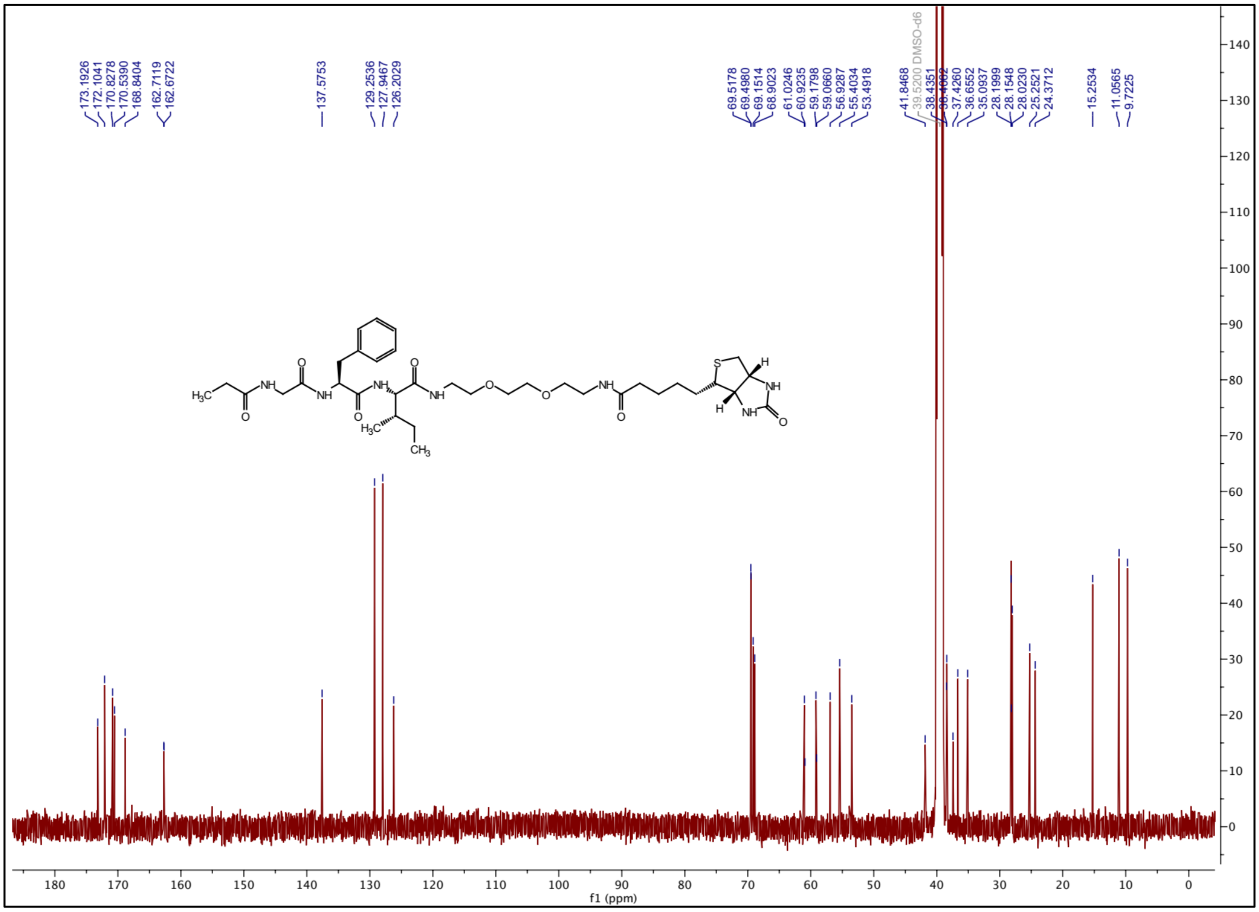

### [1]H NMR spectrum of ZA37

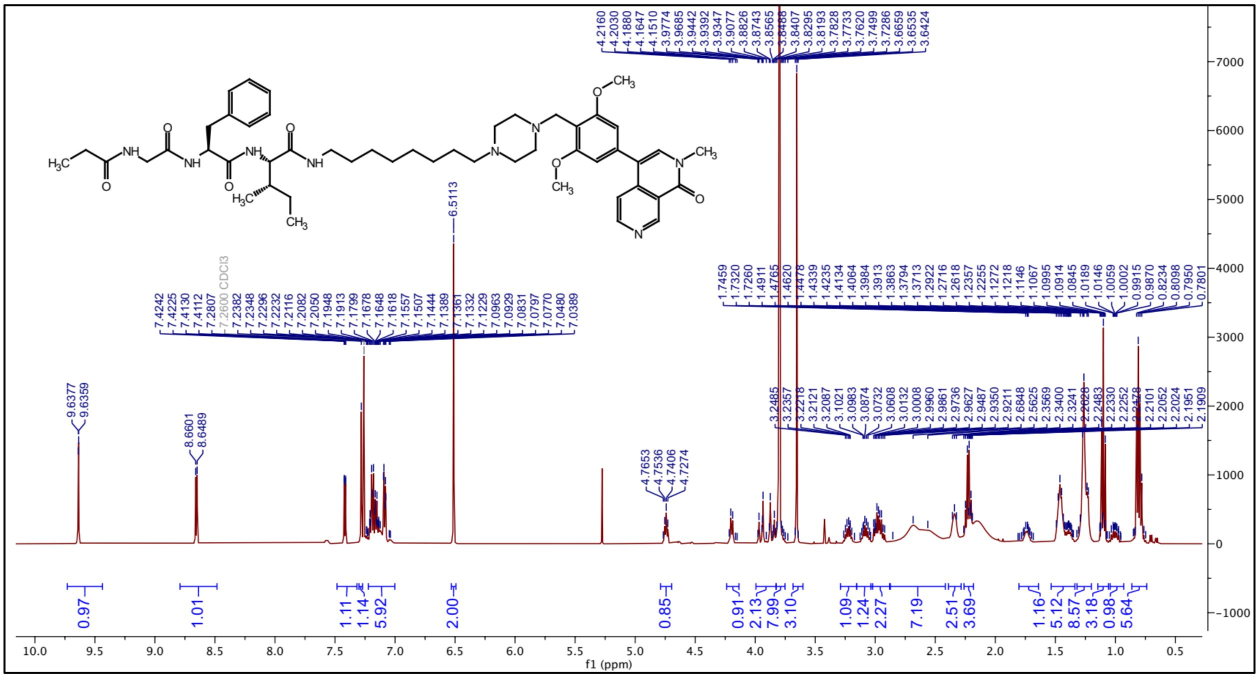

### [13]C NMR spectrum of ZA37

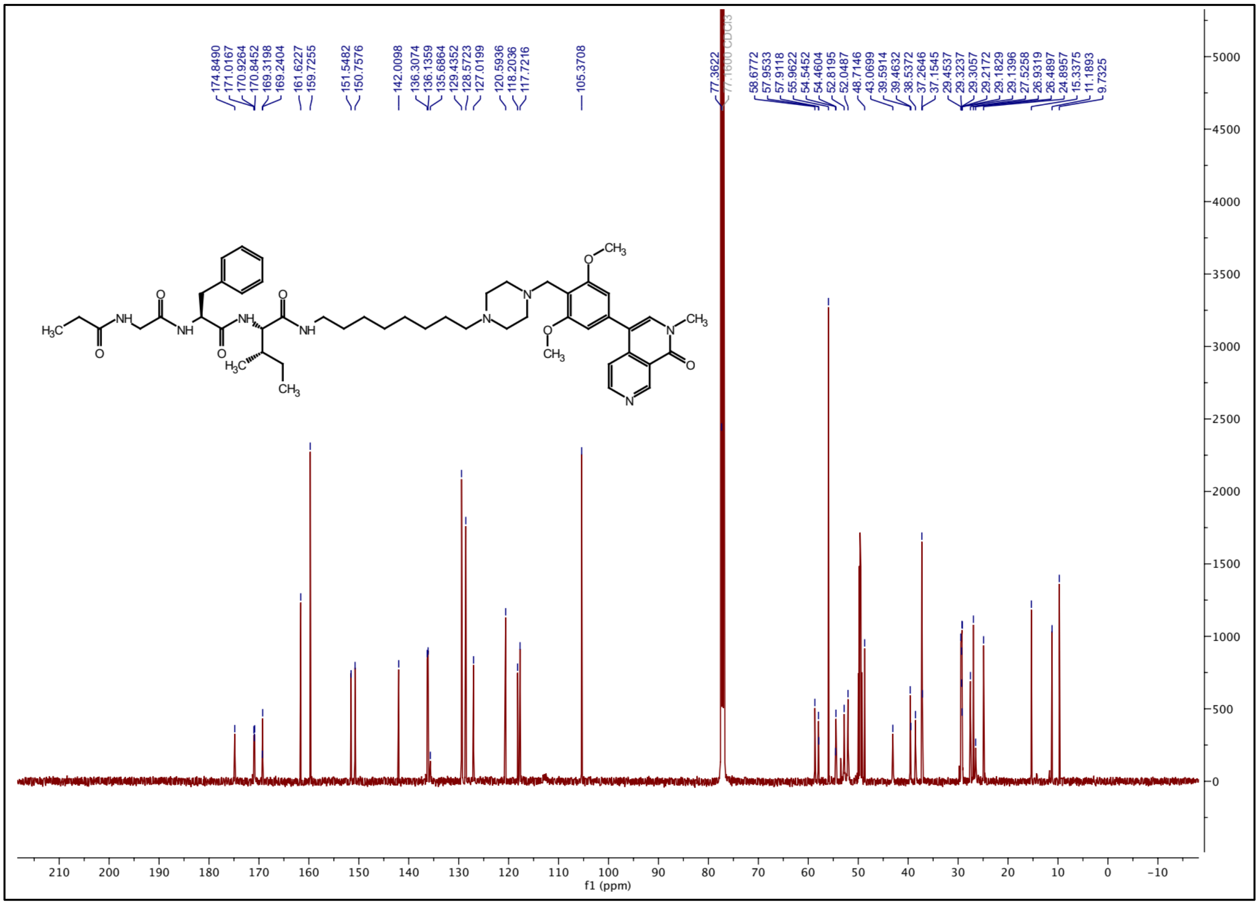
